# The auxin gatekeepers: Evolution and diversification of the YUCCA family

**DOI:** 10.1101/2025.04.11.648386

**Authors:** Mallika Vijayanathan, Amna Faryad, Thanusha D Abeywickrama, Joachim Møller Christensen, Elizabeth H Jakobsen Neilson

**Affiliations:** Section for Plant Biochemistry, Department of Plant and Environmental Sciences, University of Copenhagen, Thorvaldsensvej 40, 1871 Frederiksberg C

**Keywords:** Auxin, Evolution, Flavin-containing monooxygenase, FMO, YUCCA

## Abstract

The critically important YUCCA (YUC) gene family is highly conserved and specific to the plant kingdom, primarily responsible for the final and rate-limiting step for indole-3-acetic acid (IAA) biosynthesis. IAA is an essential phytohormone, involved in virtually all aspects of plant growth and development. In addition, IAA is involved in fine-tuning plant responses to biotic and abiotic interactions and stresses. While the YUC gene family has significantly expanded throughout the plant kingdom, a detailed analysis of the evolutionary patterns driving this diversification has not been performed. Here we present a comprehensive phylogenetic analysis of the YUC family, combining YUCs from species representing key evolutionary plant lineages. We identify and hierarchically classify the YUC family into six distinct classes and 41 subclasses. YUC diversity and expansion is explained in the context of protein sequence conservation, as well as spatial and gene expression patterns. The presented YUC gene landscape offers new perspectives on the distribution and evolutionary trends of this crucial family, which facilitates further YUC characterization within plant development and response to environmental change.

**Short summary:** Comprehensive phylogenetic and sequence analysis of the YUC gene family presents new insights into factors driving evolutionary diversification.

**Highlights:**

1. A phylogeny-based classification system for the YUCCA gene family is presented
2. YUCCA evolution and structural diversification is described, supporting a fine-tuned spatial and temporal control of auxin biosynthesis, but also holds potential for alternative capabilities

## Introduction

Auxins are essential phytohormones that orchestrate almost every facet of plant growth and development (Friml, 2022; Zhao, 2010; Sauer *et al*, 2013) such as cell elongation, vascular bundle differentiation, embryonic polarization, phototrophic and gravity response, and apical dominance (Biedroń & Banasiak, 2018; Pan *et al*, 2015; Han *et al*, 2021; Muday, 2001; Li *et al*, 2022; Mueller-Roeber & Balazadeh, 2014). In addition, auxins are critically important in how plants respond and survive in response to biotic and abiotic stresses (Cohen & Strader, 2024; Salehin, 2024; Korver *et al*, 2018). At least five endogenous plant auxins are reported (**Fig 1A**) (Simon & Petrasek 2011; Ludwig-Muller 2020). Tryptophan (Trp)-derived indole-3-acetic acid (IAA) is the most established of the auxins. IAA is widely and interchangeably referred to as ‘auxin’, a term derived from the Greek word “auxein”, meaning “to grow/increase” (Kögl & Haagen-Smit, 1931). IAA is also present in algae, bacteria and fungi, and considered an important cross-kingdom signaling molecule (Cheng et al, 2023; Fu et al, 2015; Morffy & Strader, 2020). Other auxins include 4-chloroindole-3-acetic acid (4-Cl-IAA); a bioactive form present in many Fabaceae species (Reinecke, 1999; Tivendale *et al*, 2012), indole-3-butyric acid (IBA), and indole-3-propionic acid (IPPA), as well as the Phenylalanine (Phe)-derived auxin phenylacetic acid (PAA)(Sugawara *et al*, 2015; Cook *et al*, 2016). A tyrosine-derived auxin-like molecule (hydroxy-PAA or OH-PAA) has also been identified in algae (Abe *et al*, 1974; Fries & Iwasaki, 1976), but an analogous role in plants has not been established.

**Figure 1.**
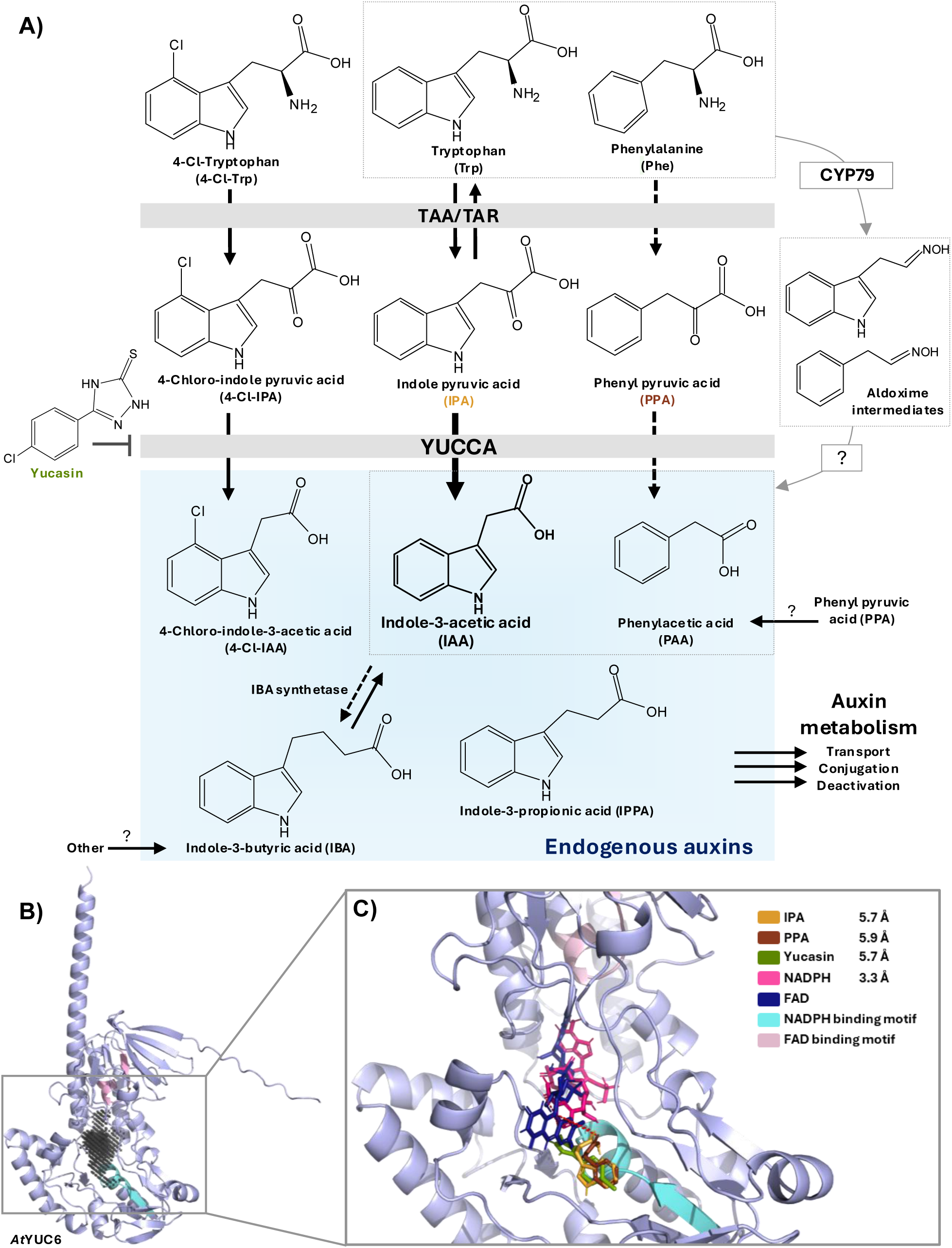
Diversity and biosynthesis of endogenous plant auxins. **A)** Plants produce several endogenous auxins (blue box). Indole-3-acetic acid (IAA) is the best described auxin. *In planta* (solid arrows) and *in vitro* (dashed arrows) show that the primary biosynthetic route for IAA operates via a TAA/TAR and YUC gene members, where YUCs perform the rate-limiting and irreversible step for IAA production. Biosynthesis of halogenated 4-Cl-IAA also operates via this pathway, while *in vitro* studies indicate involvement of TAA/TAR and YUCCAs in the biosynthesis of indole-3-butyric acid (IBA) and phenylacetic acid (PAA). Parallel production of IAA and PAA via a cytochrome P450 CYP79-catalyzed aldoxime pathway is established in several species (dashed grey boxes), while an alternative pathway via PPA for PAA production is also apparent but uncharacterized. **B)** Predicted Alphafold structure of *At*YUC6 depicts two main domains corresponding to the co-factor FAD binding domain (pink) and NADPH binding domain (aqua), with the binding pocket (charcoal) identified within the cleft between these domains. **C)** Molecular docking of NADPH, IPA, PPA and the YUC inhibitor Yucasin to FAD shows close binding distances (red dash lines) within the substrate binding pocket.

Tryptophan-dependent biosynthesis of IAA is well established (**Fig 1A**). The predominant route follows conversion of Trp into indole-3-pyruvate (IPA) through the action of Tryptophan Aminotransferase of Arabidopsis1/Tryptophan Aminotransferase-Related proteins (TAA1/TAR; EC 2.6.1.27) (Zhao, 2010, 2012). This reaction is reversible, with IPA recently shown to be the preferred substrate for the Arabidopsis TAA1 enzyme, implicating TAA1 as a key regulator for IPA accumulation (Sato *et al*, 2022). The conversion of IPA to IAA is then catalyzed by a class B flavin-containing monooxygenase (FMO) belonging to the large YUCCA (YUC) gene family. YUCs performs an irreversible oxidative decarboxylation reaction (Mashiguchi *et al*, 2011); the rate-limiting and gate-keeping step for IAA biosynthesis (Dai *et al*, 2013). The establishment of the YUC pathway as the dominant route for IAA production is strongly supported by significant in vitro, in vivo, and genetic studies, including the use of inhibitors such as Yucasin, which binds tightly within the YUC binding pocket (**Fig**. **1C**). A parallel YUC pathway for 4-Cl-IAA biosynthesis has also been established in pea (*Pisum sativum*) (Tivendale *et al*, 2012), where Trp stands at the point of chlorination. The YUC pathway for IAA biosynthesis is unique to plants, as other pathways are in operation for IAA production in bacteria, algae and fungi (Morffy & Strader, 2020). While TAA gene members are also present in green algae, the evolutionary origin of YUCs is currently unknown, likely derived from bacteria via horizontal gene transfer (Bowman *et al*, 2021). Recent analysis also proposes a class of flavin-containing monooxygenases (FMOs) sister to the YUCs (designated ‘sister YUCCAs’ or sYUCs), but detailed comparisons between this clade and the YUCs has not yet been addressed (Carrillo-Carrasco *et al*, 2023). Furthermore, a detailed comparison of what differentiates YUCs from other plant major class B FMOs [i.e. clade II (S-OX) and clade IV (N-OX) FMOs; (Nicoll & Mascotti, 2023)] has not been performed.

Biosynthesis of the other plant auxins IBA and PAA may also occur via YUC pathway (Sugawara et al, 2015; Ludwig-Müller, 2020), **Fig1A**). IBA biosynthesis from IAA is evidenced by enzymatic assays derived from maize (*Zea mays*), but corresponding genes remain unidentified (Ludwig-Müller *et al*, 1995). YUC involvement in PAA biosynthesis is supported by overexpressing YUC lines (Sugawara *et al*, 2015) and *in vitro* enzyme activity studies (Dai *et al*, 2013). Widespread biochemical characterization of YUC substrate specificity and functionality is severely limited, however, with only Arabidopsis YUC6 (AtYUC6) characterized in detail (Dai *et al*, 2013) (**Fig 1B**). AtYUC6 is capable of forming both IAA and PAA from their respective amino acids. Other routes for the biosynthesis of different auxins have also been proposed (Zhao, 2012; Cook *et al*, 2016; Günther *et al*, 2023), but to date, only one other alternative pathway for IAA and PAA biosynthesis has been firmly established in mono- and di-cotylendous species. This alternative pathway operates via cytochromes P450 from the CYP79 family, which perform two successive *N-*hydroxylations of the precursor amino acids Trp and Phe to form their respective aldoxime intermediates (Irmisch *et al*, 2015; Perez *et al*, 2021, 2023). Given that CYP79s are not present in bryophytes, lycophytes and ferns, this pathway is likely an evolutionary later adaptation for auxin biosynthesis compared to the YUC pathway.

In general, the YUC gene family has widely expanded in plants, with species possessing multiple gene copies. For example, Arabidopsis has 11, rice (*Oryza sativa*) has 14, the Norwegian Spruce (*Picea abies)* has seven, the moss *Physcomitrium patens* has six while the liverwort *Marchantia polymorpha* has two (Cao *et al*, 2019) (**Fig 2**). The diversification of YUCs has been linked to their gate-keeping role in auxin biosynthesis, whereby YUCs control production in a highly fine-tuned spatial and temporal manner (Brumos *et al*, 2018). At present, the evolutionary patterns of YUC diversification remain highly unclear with previous phylogenic analyses separating this large and critical plant family anywhere between 4-9 different classes (Poulet & Kriechbaumer, 2017; Yang *et al*, 2021; Shao *et al*, 2023; Hao *et al*, 2022), without a uniform naming system. A clear resolution of phylogenetic positioning in these studies is likely due to a restricted number of input gene sequences, and/or a bias towards angiosperms and model plants. Without a common representation of the YUC evolutionary landscape, it remains a highly challenging task for the community to compare cross-species gene diversification events, evaluate and assign linage-specific YUC structure, identify specific evolutionary clades potential holding functional diversification beyond auxin biosynthesis, and perform comparative analyses with other gene families. It is therefore highly relevant to realize the evolutionary diversity of the YUC family and establish a common classification that encompasses this rich diversity.

**Figure 2.**
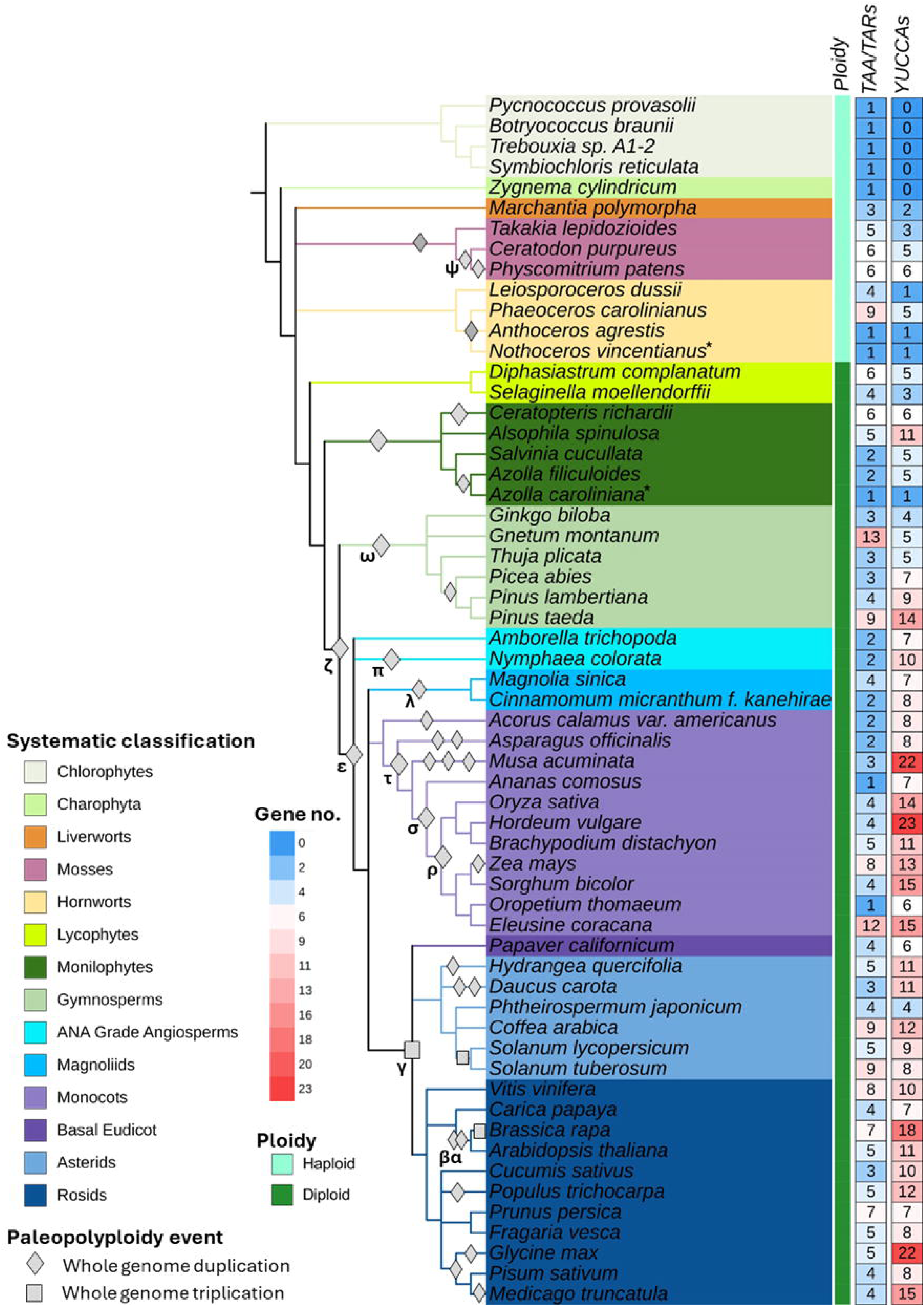
TAA/TAR and YUCCA distribution across the plant kingdom. Representative species tree was derived from taxonomic IDs using NCBI Lifemap (de Vienne, 2016) showing the species topology and number of putatively identified TAA/TAR and YUCs from each species. Paleopolyploidy events are summarized from (Van De Peer *et al*, 2017; Qiao *et al*, 2022), and updated according to (Gao *et al*, 2022; Wang *et al*, 2022b; Stull *et al*, 2021; Marchant *et al*, 2021; Catania *et al*, 2022; Zhang *et al*, 2024). The major named paleo duplication events are marked. *Indicates transcriptome dataset. The annotated tree can be visualized at iTOL v6 (<u>iTOL:</u> Figure 2<u>(embl.de)</u>).

Here we present a comprehensive phylogenetic analysis of the YUC gene family, combining YUCs from 54 different plant species, representing key evolutionary plant lineages. We identify and hierarchically classify the YUC family into six distinct classes and 41 subclasses based on evolutionary position and amino acid sequence similarity, following an expandable naming system comparable to other gene families (i.e. UDP-glucosyltransferases and cytochromes P450, (Ross et al, 2001; Nelson, 2006)). We show that the YUC family is highly diverse, having undergone multiple gene duplication and retention events throughout plant evolution, especially in angiosperms. The rich diversity of YUCs is discussed in the context of protein structure and sequence conservation, as well as localization and gene expression patterns. The presented YUC landscape offers new perspectives on the distribution and evolutionary trends which facilitate further functional characterization of this crucial gene family across the green lineage.

## Results

### TAA/TAR and YUC distribution in Viridiplantae

The main route for auxin biosynthesis in plants operates via the TAA/TAR and YUC pathway (**Fig 1A**). To determine distribution for both enzyme families, homologues were acquired from the sequenced genomes representing major lineages throughout the plant kingdom and green algae (59 species. **Fig 2**). All the identified and retrieved sequences were subsequently verified and manually inspected to eliminate redundant sequences or pseudogenes. In total, 254 TAA/TARs and 469 YUCs were identified as potential homologues from these 59 species, and they are evolutionary conserved in all plant species (**Fig 2**; **Data S1**). Chlorophyte and streptophyte algae also possess TAA/TARs in line with previous reports (Carrillo-Carrasco *et al*, 2023; Romani, 2017; Bowman *et al*, 2021). Gene copy numbers ranged between 1-13 for the TAA/TAR family, and between 1-23 for the YUC family. Angiosperms typically possess a higher ratio of YUCs compared to TAA/TARs, but similar copy numbers are present in bryophyte, fern and gymnosperm genomes (**Fig 2**). *Phaeoceros carolinianus* and *Gnetum montanum* stand out as exceptions for possessing a high TAA/TAR to YUC ratio.

### Phylogenetic distribution and classification of the YUC and TAA/TAR gene families

To infer the molecular basis of YUC evolution and diversity, phylogenetic analysis was performed on 469 YUCs identified from 54 plant species (**Fig 3**). The unrooted tree positions non-vascular plant YUCs (i.e. bryophytes and lycophytes) at the center, with vascular plant YUCs [ferns (monilophytes), gymnosperms and angiosperms] radiating in two major directions (**Fig 3A**). Further large evolutionary diversification events are apparent, with separation into five additional and distinct classes observed. Based on the evolutionary position of these classes, and an amino sequence identity cut-off of ∼40%, we propose a formal classification of six primary YUCs Classes hereby named: Early Diverging (ED), 1 – 5 (**Fig. 3**). Diversification within each major YUC class has occurred, and using a combination of evolutionary position, bootstrap support and a sequence identify cut-off of ∼50%, 41 subclasses were assigned (**Fig S1; Data S2; Fig 3B**). Comparison of YUCs with other plant class B FMOs (sYUC, N-OX and S-OX), showed a maximum sequence identity of 23% with sYUCs, and < 20% sequence identity with N-OXs and S-OXs. The average pairwise distances between YUCs and sYUCs, N-OXs and S-OXs are 0.58, 0.64, and 0.52, respectively (**Data S2, Fig. S2**). Pairwise diversity among plant classes was also assessed, indicating that angiosperms demonstrate markedly greater YUC diversity compared to other lineages (**Fig S3**). To compare the overall evolutionary pattern of YUCs to the TAA/TAR family, phylogenetic analysis of 254 TAA/TARs from the same 54 plant species, plus five algal TAA/TARs were also performed (**Fig. 4; Data S1**). YUCs show significantly more divergence compared to TAA/TARs, whereby TAA/TARs diversity has only occurred along two major axes common across all plant lineages, with algal TAA/TARs positioned at the center, which agrees with previous studies (Carrillo-Carrasco *et al*, 2023; Matthes *et al*, 2019).

**Figure 3.**
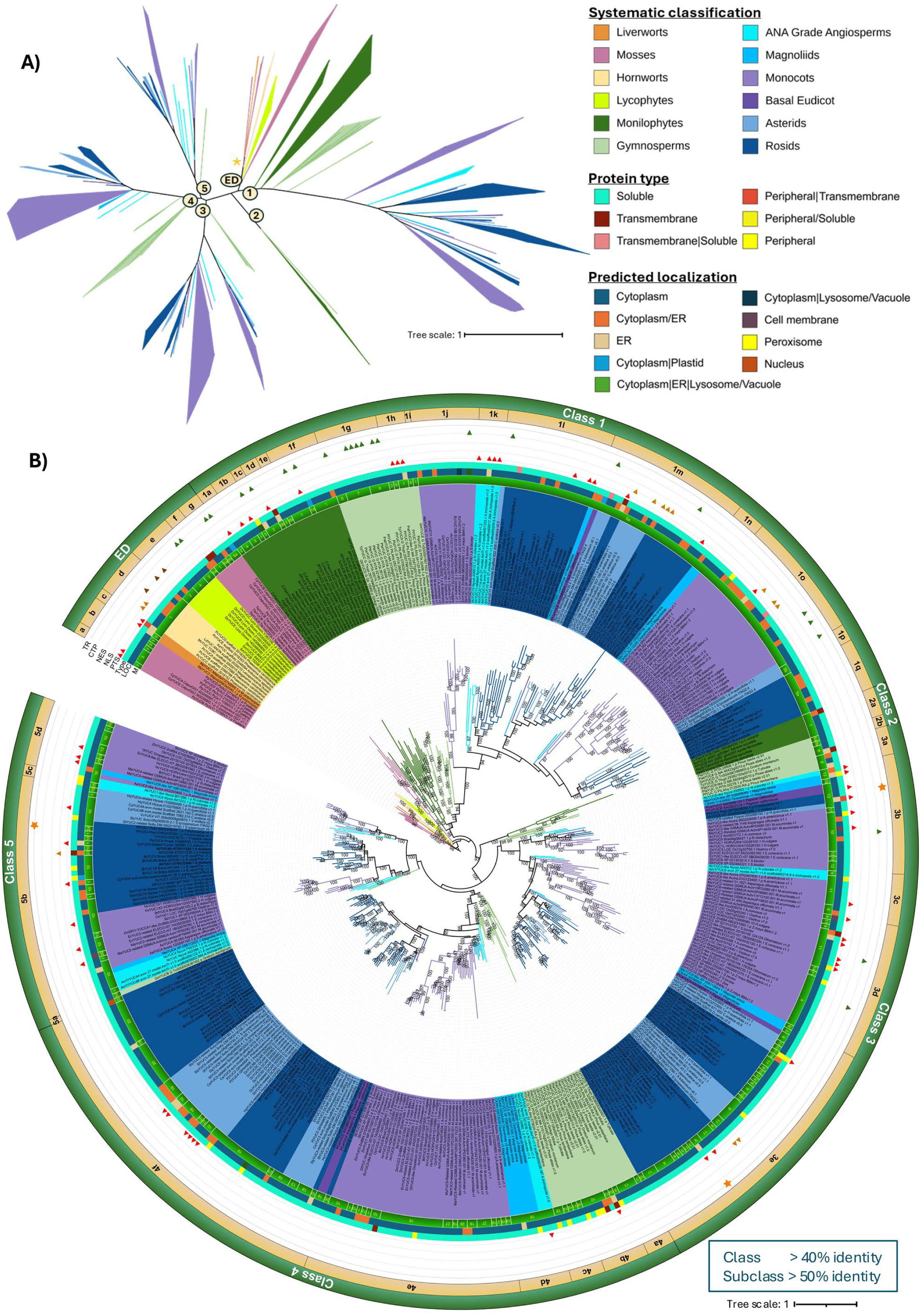
Phylogenetic analysis and classification of the YUC gene family. **A**) Maximum Likelihood unrooted phylogenetic tree exhibit of 469 YUCs representing plant evolution. Annotated unrooted tree can be visualized at iTOL v6 (Letunic & Bork, 2024b). **B)** Rooted phylogram shows the divergence of YUCs from 54 plant species. Circular maximum likelihood phylogenetic tree rooted by selected bryophyte YUCs (rooting location marked with an asterisk in the unrooted tree) showing YUC classification. YUCs are categorized into six major classes [Early Diverging (ED), Classes 1-5; sequence identify cutoff of 40%] and additional sub-classes (sequence identity cut-off of 50%). Bootstrap values ≥ 80 are included. YUC protein characteristic features are mapped: M - motif pattern type, LOC - subcellular localization, Type - membrane type, PTS - Peroxisomal targeting signal, NLS - Nuclear localization signal, NES - Nuclear export signal, CTP - Chloroplast transit peptide, TR – Thiol reductase activity. An expanded version of the tree (**Fig EV3**) is available at the iTOL repository (<u>iTOL:</u> Figure 3B<u>(embl.de)</u>)

**Figure 4.**
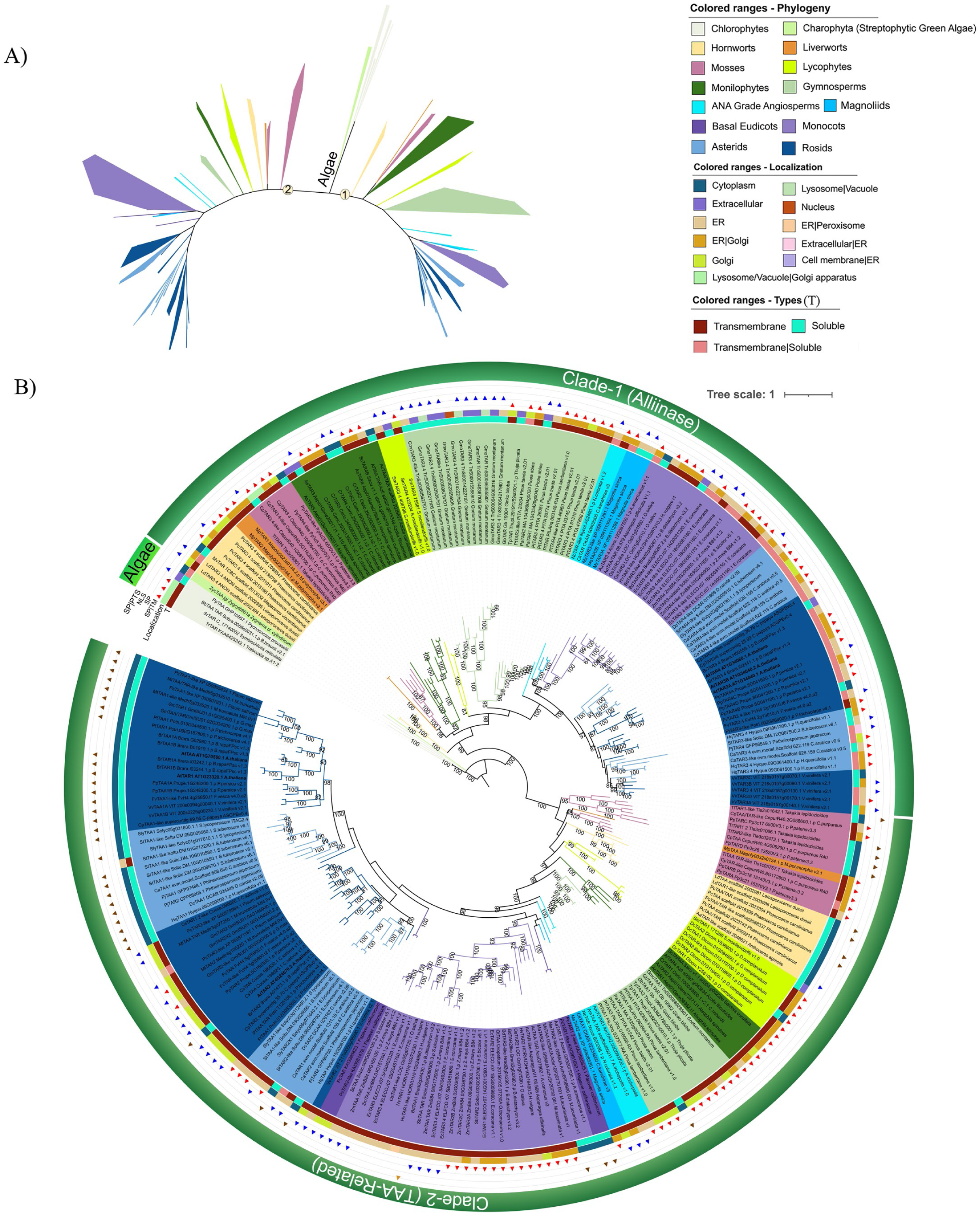
Phylogenetic analysis TAA/TAR gene family. **A**) Maximum Likelihood (ML) unrooted phylogenetic tree showing 254 TAA/TARs. **B)** Rooted phylogram, where proteins from the algal lineage are grouped at the base of the tree. In addition to the algal TAA/TAR (Early diverging forms) clade, they are further classified into two main groups: Clade-1 and Clade-2. Proteins from *A. thaliana* i.e. AtTAR3 and AtTAR4 are clustered in Clade-1 (Alliinase-related); while AtTAA, AtTAR1 and AtTAR2 are grouped in Clade-2 (TAA-Related). The rooted tree was also mapped with the predicted characteristic features [SP-Signal peptide; TM - Transmembrane domain; PTS - Peroxisomal targeting signal, NLS - Nuclear localization signal, T-Types, Localization]. The expanded version of this tree (**Fig EV4**) is available in the iTOL depository (https://itol.embl.de/tree/1302262366206311729598277).

YUC Class ED is categorized into seven subclasses (EDa-g) and encompasses all non-vascular YUC members. Liverworts, hornworts and lycophytes fall into lineage-specific sub-classes (EDc, d and e, respectively), while moss YUCs exhibit a wider evolutionary divergence, separating into four moss-specific sub-classes (EDa, b, f and g). Class 1 represents one major direction of YUC evolution and is the largest and most diverse class assigned. Class 1 consists of 17 sub-classes (1a-o) with vascular species representing ferns, gymnosperms and angiosperms. This class has diversified in multiple directions, with three major duplications apparent for Class 1 angiosperm YUCs (represented by subclasses j-q). Similar diversification is also clear for fern YUCs, exemplified with all fern species genomes possessing members from multiple sub-classes (e.g. *C. richardii* and *A. spinulosa* have members from classes 1b, c, d & f while *A. filiculoides* and *S. cucullate* possess members from classes 1c, e & f). The distribution of Class I YUCs generally follows the expected evolutionary positioning, with the exception of a monophyletic monocot subclass specific to Poaceae (class 1j), which diverges earlier than basal angiosperms. Classes 2-5 represent the second major direction of YUC evolution. YUC Class 2 consists of only two subclasses; one that is specific to ferns (2a) and one specific to gymnosperms (2b). Classes 3-5 are represented by gymnosperms and angiosperms, with angiosperms in classes 3 and 5 exhibiting a further evolutionary duplication event following the split from gymnosperms. Few dicot species retain representative YUCs from these duplications, with only *P. californicum*, *P. trichocarpa*, *V. Vinifera* and *H. querifolia* possessing YUCs from Class 3b, and no dicot species retained members from the Class 5d. Classes 3b and 5d are instead retained by a greater number of monocot species.

Overall, the high copy number of angiosperm YUCs is primarily attributed to paleoploidy genome duplication and triplication events (denoted as γ, α, β, π, Ψ, ω etc. in **Fig 2**) and subsequent gene retention. Tandem gene duplications also contribute to the high gene copy number in some species, such as *H. vulgare, M. truncatula*, *O. sativa, P. trichocarpa* and *V. vinifera*, but not all (e.g. *A. thaliana*; **Fig 6**).

**Figure 5.**
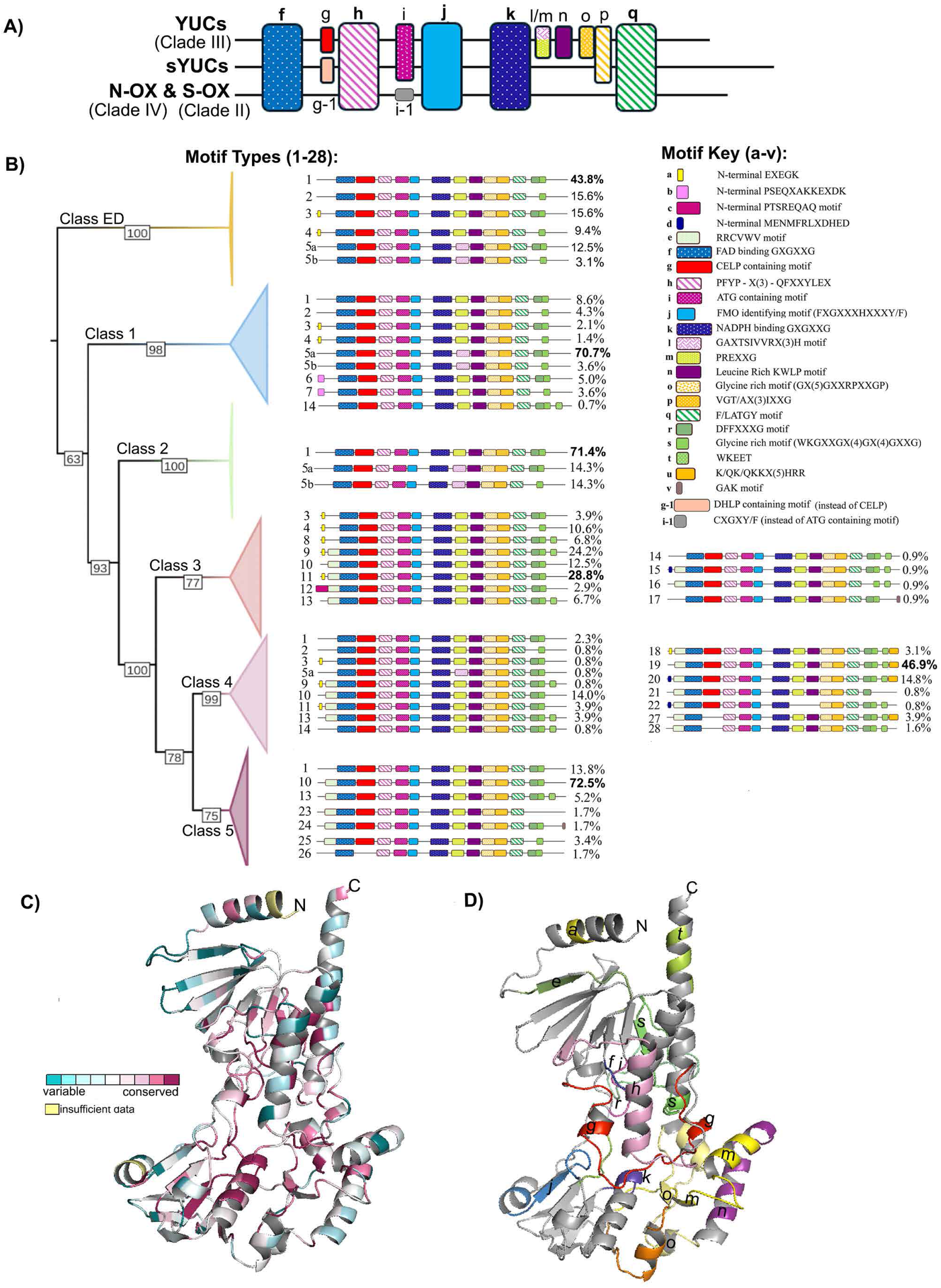
Domain architecture, highlighting motif patterns and structural conservation. **A**) Overall motif conservation patterns within class B FMOs. YUCs belonging to Clade III (clade-based classification adapted from (Nicoll & Mascotti, 2023).The sYUC data is obtained from (Carrillo-Carrasco et al, 2023). B) Collapsed Maximum Likelihood phylogenetic YUC tree with bootstrap values showing class-based (Class-ED to Class-5) motif organization pattern. The motif type most prevalent in each class is indicated in bold. C) YUC structural model and its conservation pattern (based on AtYUC6, accession: AT5G25620.1). The model was developed in the ConSurf Server (Ashkenazy *et al*, 2016; Yariv *et al*, 2023a) using the Modeller (Webb & Sali, 2016). The *N* - and *C -* terminals are marked. D) Predicted motif’s locations are mapped in the structural model (AtYUC6). The motif number and color are according to the motif key.

**Figure 6.**
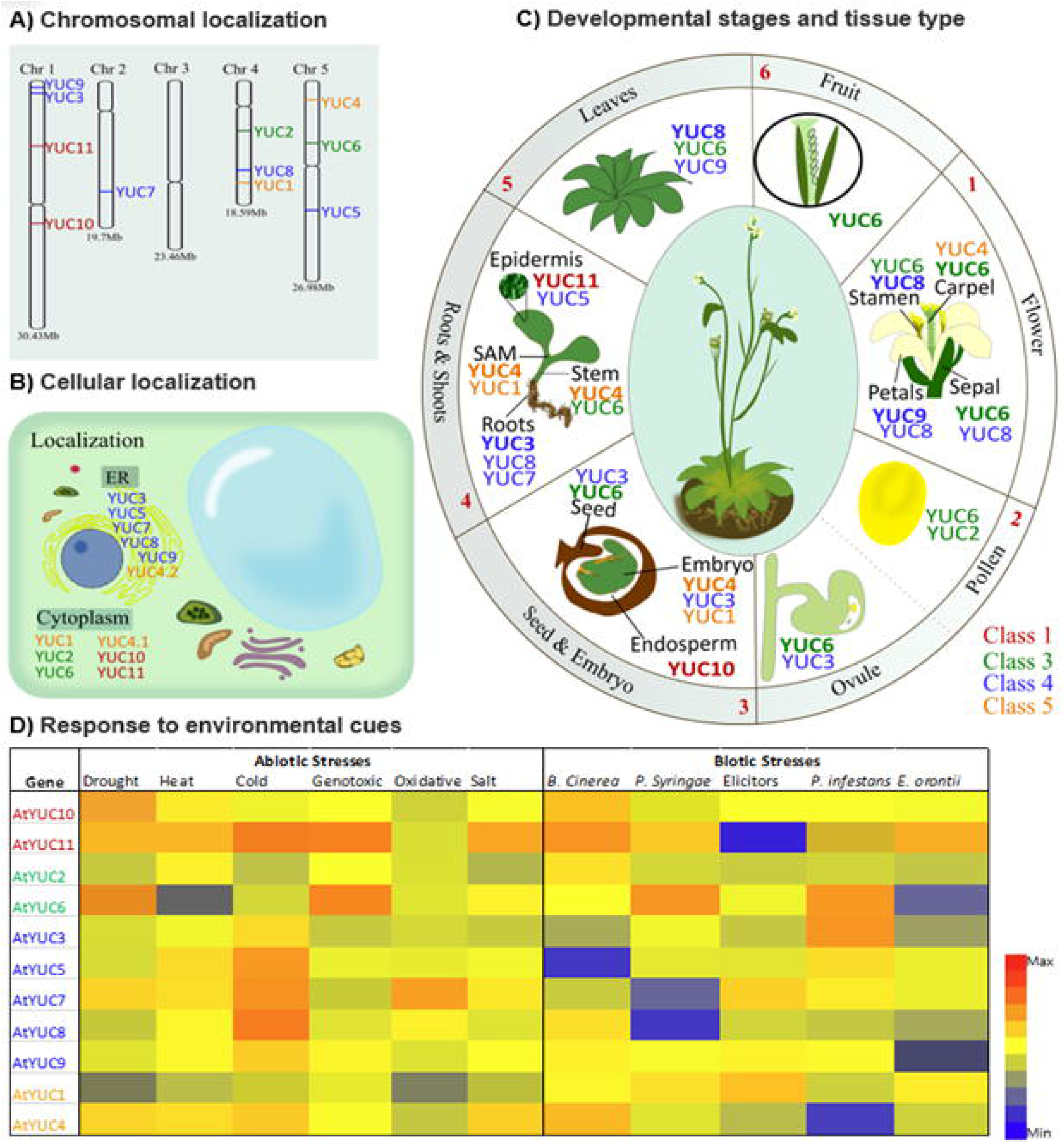
Multi-scale organization and gene expression of the 11 Arabidopsis YUCs. **A)** Chromosomal mapping of AtYUCs, showing broad distribution in the genome. **B)** Published experimental data (Kriechbaumer *et al*, 2016) show all class 4 AtYUCs localize to the ER, while other classes localize to the cytoplasm with the exception of a class 5 splice variant. **(C)** AtYUCs are differentially expressed across different tissues and in various developmental stages. Only the top 2-3 expressed AtYUCs are listed with the highly expressed ones marked as bold for each tissue type. Tissue expression data was retrieved from the Plant Expression Omnibus database (Koh *et al*, 2024). **D)** Heatmap of AtYUC differential gene expression in leaves in response to various abiotic and biotic stress factors. Environmental heatmap data represents relative average temporal gene expression response. Environmental stress expression data was retrieved from the eFP browser (Winter *et al*, 2007) representing average log2 values of up- and down-regulated YUC genes. SAM: Shoot apical meristem.

### Sequence-structure conservation, motif analysis and predicted sub-cellular localization

Multiple-sequence alignment demonstrates considerable sequence similarity among the studied YUCs, with an overall minimum identity of ∼40% (**Data S2-S4**). Detailed motif analysis of the YUC family was done in MEME suite (Bailey *et al*, 2015). To broadly compare the YUC motif patterns with other plant class B FMO families, 20 ‘S-OX’, ‘N-OX’ and ‘sYUC’ representatives were also included. MEME-derived motif analysis, followed by multiple sequence alignment comparisons, showed that the class B families examined share five conserved motifs. The common motifs include the two signature Rossmann folds (GXGXXG) corresponding to FAD-binding and NADPH-binding motifs, an FMO identifying ‘FXGXXXHXXXY/F’ motif, and a ‘F/LATGY’ motif (**Figure 5A, Data S4 and S5**). This work identified a fifth conserved common motif ‘PFYP-X(3)-QFXXYLEX’ in all the studied class B FMOs, which is positioned in between the FAD binding Rossmann fold and FMO identifying motif.

Twenty-two distinct sequence motifs (labelled ‘a-v’) were identified in various YUCs, and based on their occurrences, were classified into 28 motif organization pattern types (labelled as Type 1-28 in **Fig 5B-C**). Four motifs are unique to the YUC family and are present in all YUCs analysed. Three YUC-specific motifs are located between the NADPH-binding and F/LATGY motif and are denoted i) GAXTSIVVRX(3)H or ‘PREXXG’, ii) ‘Leucine-rich KWLP’ and iii) ‘Glycine-rich GX(5)GXXRPXXGP’ (motifs l or m, n & o; respectively; **Fig 5A**). The PREXXG motif (motif m) exhibits some sequence divergence whereby 70% of Class 1 YUCs instead possess a GAXTSIVVRX(3)H-motif (motif l) located on the periphery of the substrate binding cavity (**Fig S5-S6**). The fourth YUC-specific CELP motif is located between the FAD-binding motif and the PFYP-X(3)-QFXXYLEX motif (motif g; **Fig 5**). Two motifs are shared between the YUCs and sYUC: an ATG-containing motif situated between the PFP(X)nYY/WLE and FMO-identifying motifs and a VGT/AX(3)IXXG motif immediately after the first glycine-rich motif, preceding the F/LATGY motif (**Fig 5A**). For all species investigated with multiple YUC copies, they could be classified into different motif types. For example, *P. patens* has six YUCs which classify into three different motif types (Types: 1, 3, 5a), while the 11 Arabidopsis YUCs are organized into seven different motif types (Types: 5, 9, 10, 11, 19, 20 and 23) (**Fig S5**). More details on species-specific motifs and their corresponding sequences are available in **Table S1-S2** and **Data S5-S6**.

Motif patterns related to YUC classes were also observed, with some motifs more prevalent in some classes compared to others, and most sequence divergence occurring in the carboxyl *(C)* or amino (*N)*-terminal area. For example, the *N*-terminal motif EXEGK (motif a) is predominantly found in class 3 YUCs, RRCVWV motif (motif e) is exclusively found in YUC classes 3-5 while ‘PSEQXAKKEXDK’ motif (motif b) is found only in Class 1 gymnosperm YUCs. The *C*-terminal ‘K/QK/QKKX(5)HRR’ motif (motif u) is restricted to Class 4 YUCs. In a few instances, some key motifs were absent which may indicate sub-functionalization, neo-functionalization or pseudogenization. For example, *P. sativum* YUC5 (class 4f) lacks the ‘PREXXG’ motif and the leucine-rich KWLP motif, while *Musa acuminata* YUC (class 4e), lack the ‘CELP’ motif (**Data S4**).

All 469 YUC sequences were combined and analysed for overall structural conservation patterns and motifs mapped. Using the best-known Arabidopsis YUCs as a model system we aimed to map conserved and variable regions in more detail. With Arabidopsis YUC6 (*At*YUC6) as a template, highly conserved regions were found to associate with co-factor binding and substrate pocket areas as expected (**Fig 5C, D; Fig S4**). We then superimposed the structures of all 11 *At*YUCs and show a highly similar structural fold (RMSD 0.33-0.47 Å) with a highly positively charged substrate binding pocket in all (**Fig S5**). *At*YUCs are categorized into four different classes (Class 1: *At*YUC10 & 11, Class 3: *At*YUC2 & 6, Class 4: *At*YUC3, 5, 7-9, Class 5: *At*YUC1 & 4), with the most structural variation exhibited at the *C*-terminal and/or *N*-terminal regions. *At*YUC10 and *At*YUC11 (Class I) also show sequence divergence around the substrate binding pocket area (i.e. lack motif ‘m’), with this region predicted to be under negative selection pressure (**Fig S6, Data S7-S8**). To assess if this sequence divergence altered the binding pocket surface area (SA) and volume (V), these parameters were calculated. While AtYUC10 was predicted to be the smallest of the AtYUCs (SA: 1001.7 Å; V: 758.6 Å; **Fig S5**), it cannot be fully attributed to Class I sequence divergence as AtYUC11 (Class I) is similar to other AtYUCs, with a SA and V of 1102 Å and 854 Å, respectively. Class 3 YUCs (i.e. AtYUC2 and AtYUC6) possess the highest surface area and cavity volume parameters of the AtYUCs (**Fig S5**).

To assess whether overall YUC sequence divergence is linked to sub-cellular positioning (**Fig 3, Fig S7-S8**), *in silico* localization analysis of all 469 YUCs was conducted using the DeepLoc server (Ødum *et al*, 2024). This analysis predicted approximately 80% of the analysed YUCs have cytoplasmic localization, ∼15% to the cytoplasm or endoplasmic reticulum (ER), and ∼4% are localized to the ER alone. The remaining ∼1% of YUCs possess either plastid, cell membrane, lysosome, vacuole, nucleus and peroxisomal-predicted localizations. In addition, we observe instances of nuclear and peroxisomal target signals as well as chloroplast transit peptides. Peroxisomal target signals are dispersed across all YUC classes except for Class 2, with the caveat that Class 2 has a significantly lower number of YUC representatives. Chloroplast target peptides and nuclear localization signals are prevalent in YUC classes ED, 1 and 3. DeepLoc analysis also predicted that the majority (∼91%) of YUCs as soluble, with approximately 9% predicted to be transmembrane proteins.

TAA/TAR localization was also predicted and found to be equally distributed to the ER, cytoplasm, and ER/Golgi apparatus (∼25% in each location). The remaining 25% of TAA/TARs are located in the Golgi apparatus (∼15%) or extracellularly (∼6%), with less than 1% each in other cellular components, including the nucleus, lysosomes, or vacuoles. Most TAA/TARs possess either a signal peptide or a transmembrane domain (∼73%), although few members of Clade-2 have a nuclear localization signal (∼17%). Approximately ∼50% of TAA/TARs are predicted to be transmembrane proteins, while ∼35% are soluble and ∼15% are predicted to be either transmembrane or soluble (**Data S9**). The sequence and structural conservation pattern of Arabidopsis TAA/TARs is given in **Fig S9**.

### Multi-scale coordination of YUC gene expression and interactions; a focus on Arabidopsis

To examine patterns defining YUC classes, the 11 *A. thaliana* YUCs were broadly compared across different scales: genomic and cellular localization, as well as gene expression and co-expression at different developmental stages, tissues and in response to environmental cues. Genomic mapping observations from the ePlant database (Waese *et al*, 2017), indicate that AtYUC gene locations are dispersed across various chromosomes, i.e., chromosomes 1, 2, 4, and 5 (**Fig 6A**). *At*YUC3 and *At*YUC9 from class 4 are in close proximity; however, they are not tandem duplicates as they fall into separate phylogenetic branches (**Fig 3B**). *In silico* analysis indicates that *At*YUC6, *At*YUC10 and the splice variant *At*YUC4.2 have predicted ER localization, while the rest are predicted to localize to the cytoplasm (Fig 3B). *In planta* studies, however, do not support *in silico* predictions, with all class 4 *At*YUCs (*At*YUC3, *At*YUC5, *At*YUC7-9) and *At*YUC4.2 (class 5) localized to ER while the other AtYUC classes localized in the cytoplasm (Kriechbaumer *et al*, 2012, 2016) (**Fig 6B).** In a separate *in planta* study, AtYUC6 (class 3) was reported to localize in both the cytoplasm and an unidentified endomembrane compartment (Kim *et al*, 2007).

*At*YUC gene expression patterns across various tissues and developmental stages were summarized according to the Plant Expression Omnibus (PEO) database (Koh *et al*, 2024). *At*YUCs classes 3, 4 and 5 show broad gene expression patterns, expressed across different ontogenetic stages and tissues (**Fig 6C, Data S10**). This is exemplified by *AtYUC6* which is expressed in the seed, stem, leaves, flower and fruit tissue, and at different developmental stages. In contrast, class 1 *At*YUCs (*AtYUC10*, *AtYUC11*) show a relatively narrow expression pattern, with *AtYUC10* expressed in the endosperm and *AtYUC11* expressed in the epidermis of cotyledons. Differential gene expression data in response to diverse biotic and abiotic stress factors was summarized from the TAIR database generating a heat map based on temporal leaf-specific average expression values across various environmental treatments. The expression of individual *At*YUCs appears highly variable and fine-tuned in response to different environmental stressors, with no obvious class-specific gene expression patterns identified (**Fig 6D**). For example, there is a trend where *At*YUC1 (class 5) is consistently down-regulated in response to abiotic stresses, but shows a variable gene expression upon different biotic stress treatments. In contrast, AtYUC6 (class 3) exhibits a variable gene expression response upon different abiotic stresses.

To examine different proteins that may interact with the various AtYUCs, the top interacting proteins were predicted using the STRING database(Szklarczyk *et al*, 2023) with a confidence cut-off > 0.7. Proteins related to auxin metabolism are commonly shared across all AtYUCs including TAA1, TAR1, TAR2, Amidase 1(AMI1), Aromatic aminotransferase ISS1 (ISS1), Indole-3-acetaldehyde oxidase (AAO2), Auxin Response Factors (ARFs), as well as different Nitrilases (NIT1-3) and PIN genes (**Fig S10, Data S10**). The list of interaction partners also included different transcription factors (e.g. LEAFY COTYLEDON2 (LEC2), PHYTOCHROME INTERACTING FACTOR (PIF), WUSCHEL (WUS)). The stress-related Aldehyde dehydrogenases (ALDHs – ALDH2B7, ALDH3H1, ALDH3F1) are also predicted as interacting proteins, especially for *At*YUC11, where it lies within the top 10 list of predicted proteins.

## Discussion

### Phylogenetic analysis and classification of the YUC family

The YUC family plays a fundamental role in plant auxin biosynthesis, and has attracted significant attention since their role in auxin metabolism was initially identified (Zhao *et al*, 2001). To date, all characterised YUCs catalyze the conversion of IPA into IAA (Cohen & Strader, 2024), a critical plant hormone. In addition, YUCs are involved in the biosynthesis of other auxins, such as 4-Cl-IAA, and possibly PAA and IBA (**Fig 1**). Thus, YUCs represent a plant enzyme family that is extremely important for growth, development and response to environmental stimuli (Cao et al, 2019; Martin-Arevalillo & Vernoux, 2023). The YUC family is relatively large, showcasing significant diversification in angiosperms, represented by the higher ratio of YUCs compared to the precursor biosynthetic enzyme for IAA production, the TAA/TARs family (**Fig 2**). With an average of 11 gene copies per angiosperm species, the YUC family is in a similar range to other denoted large gene families, such as the transcription factor families Aux/IAA and ARF (Li *et al*, 2016; Wu *et al*, 2017; Hernández-García *et al*, 2024) whereby high copy number is attributable to environmental stress response (more details below).

To aid the classification and interpretation of YUC diversification and evolution we established an expandable nomenclature system for the YUC family consisting of six distinct classes (Classes ED, 1 – 5), and 41 subclasses (A, B, C etc.). This expandable naming system is based upon phylogenic positioning and sequence identity cut-offs following similar naming systems of other large gene families (e.g. cytochromes P450 and UDP-glucuronosyltransferases (Nelson, 2006; Ross et al, 2001). The presented YUC classification builds upon previous attempts to organize the YUC family, whereby YUCs were grouped anywhere between 4-9 different clades (Poulet & Kriechbaumer, 2017; Yang *et al*, 2021; Shao *et al*, 2023). Here, the inclusion of diverse plant lineages improves delineation between phylogenetic clades and aids the interpretation of how the family has expanded through evolutionary time. For example, Class 2 YUCs appear unique to ferns and gymnosperms, and given their evolutionary position, may possibly indicate it arose following an ancestral genome duplication event prior to the zeta (ζ) WGD event (Qiao *et al*, 2022; Li *et al*, 2015) in seed plants (**Fig 2**). In a few cases, phylogenetic subclasses did not fall within the expected evolutionary position, such as subclasses 1J and 1L. This might suggest lineage-specific diversification, exemplified by an unusual substitution of serine (S) and alanine (A) in the otherwise conserved and YUC-specific CELP motif of the Poales-specific subclass 1J (**Data S4**). Further characterization of 1J clade could illuminate possible functional diversification or neo-functionalization. Overall, the comprehensive phylogenetic analysis of the YUC family presented here shows that many YUC genes are retained following paleoduplication events. In contrast to YUCs, much fewer TAA/TARs have been retained throughout plant evolution, with these genes only diversifying in two major directions in line with other TAA/TAR-specific studies (**Fig 4**) (Poulet & Kriechbaumer, 2017; Matthes *et al*, 2019; Carrillo-Carrasco *et al*, 2023).

### Origin, Expansion and Diversification of the YUC family

Our homology-based BLAST did not retrieve any potential YUCs from core chlorophyte and streptophyte algae. This result is in line with the consensus that YUCs have originated from bacteria via horizontal gene transfer (HGT), then neo-functionalized to acquire IAA functionality and expanded throughout the Plantae kingdom (Bowman *et al*, 2021). In contrast, the TAA/TAR family has a different evolutionary origin, arising in green algae (**Fig. 2, 4**; (Wang *et al*, 2014)). In general, HGT helps plants to acquire novel genes from bacteria or fungi and adapt to diverse surroundings (Soucy *et al*, 2015), and YUCs represent a key trait for successful terrestrialization. Following the shift from algae to pioneer land plants, a whole genome duplication (WGD) event occurred (Leebens-Mack *et al*, 2019; Qiao *et al*, 2022). This evolutionary event is apparent in the TAA/TAR tree, with two clear directions of evolutionary expansion, but is not evidenced in the YUC evolutionary tree where bryophyte and lycophyte YUCs clade together (i.e. class ED).

The YUC family shows great expansion and high copy number variation (CNV), especially within angiosperms. Evolutionary duplication events – including WGDs, local and small-scale duplications – promote CNV (Soltis & Soltis, 2021; Defoort *et al*, 2019). Arabidopsis, for example, possesses > 60% of genes with at least one paralog corresponding to a syntenic block derived from one of the three α, β, or γ WGD events (Bowers et al, 2003; Panchy et al, 2016; Fig. 2). Here we show that YUC gene expansion across monocots and dicots is largely attributed to both whole genome duplication (WGD) and whole genome triplication (WGT) events (**Fig 2**), but also local, segmental and small-scale duplications. For example, tandem and segmental gene duplicates explain YUC CNV in *Medicago* species, in addition to paleoduplication events (Shao *et al*, 2023). Following a duplication event, most duplicated genes are typically impacted by negative (purifying) selection (Lye & Purugganan, 2019) and become inactive through non-functionalization, pseudogenization and gene loss. Analysis of *Medicago* and *Isatis indigotica* YUCs shows that they are subject to strong negative selection (Shao *et al*, 2023; Qin *et al*, 2020), whereby evolution will hinder the spread of deleterious mutations. Evidence of pseudogenization is also present within the YUCs analysed here, exemplified by banana (*Musa acuminata*) which possesses three TAA/TAR and 22 YUCs. The high YUC copy number in banana is presumably a result of three WGD events following diverging from other monocot lineages (**Fig. 2**; (D’hont *et al*, 2012)), and motif analysis shows several YUCs may be pseudogenized, due to a loss of the YUC-specific CELP motif in several homologs (**Fig 3**, **Fig 5**, **Data S5**).

Following duplication events, genes may also be retained driven by sub-functionalization or neo-functionalization (Nowak *et al*, 1997; Kondrashov & Kondrashov, 2006; Kuzmin *et al*, 2022). High CNV of YUCs relative to TAA/TARs may be linked to their specific biochemical functionalities within auxin production. The TAA/TARs catalyze the conversion of Trp to IPA (Zhao, 2010, 2012), a reversible reaction. Recent characterization of the Arabidopsis TAA1 enzyme shows IPA as the preferred substrate compared to Trp, implicating TAA1 as a key regulator for IPA accumulation (Sato et al, 2022). In contrast, the YUC-catalyzed conversion of IPA to IAA is irreversible and rate-limiting. Accordingly, YUCs stand as the critical gatekeepers for auxin production, where YUC sub-functionalization is strongly linked with gene expression and the tightly controlled, highly fine-tuned production of auxin across time, space and environmental condition (**Fig 6**). High YUC CNV and the complex regulation of auxin production in response to environmental change, mirrors other instances in angiosperms, where CNV of other gene families (e.g. Nucleotide-Binding Leucine-Rich Repeat (NB-LRR) genes) is associated with elevated protein expression, enzymatic activity, and environmental tolerance in plants (Zmieńko et al, 2014; Panchy et al, 2016), providing a mechanism to facilitate rapid adaptation (Sakurai et al, 2005; Lian et al, 2004). Notably, in a recent population genomics assessment of the endangered dipterocarp *Hopea chinensis*, a YUC was identified as a key gene under positive selection, associated with cold and drought tolerance (Xiang *et al*, 2025).

YUC diversification corresponds with highly varied spatial and temporal regulation of auxin synthesis, evidenced by diverse expression patterns of different Arabidopsis *YUC* genes across developmental stages and tissues (**Fig 6C**). For example, AtYUC1 and YUC4 are specifically expressed at the boundary layer between floral whorl 3 and 4, regulated by the transcription factor *SUPERMAN,* which increases local auxin accumulation and leads to the formation of extra primordia of reproductive organs (Xu *et al*, 2018). In maize (*Zea mays),* loss of the *spi1* YUC (class 5B) results in severe vegetative and reproductive tissue phenotypic defects (Gallavotti *et al*, 2008). The relatively high CNV of angiosperm YUCs may suggest a higher order of auxin biosynthetic regulation in flowering plants, providing the ability to generate novel plant morphologies (Gallavotti *et al*, 2008).

Beyond auxin biosynthesis, a small handful of YUCs and ‘YUC-like’ proteins have also been reported to possess extended or alternative functionalities and may contribute to a high degree of YUC gene retention and diversification via neo-functionalization. For example, *in vitro* biochemical analysis of *At*YUC6 reports some substrate promiscuity such that *At*YUC6 can catalyze the conversion of both IPA and PPA to their respective auxins IAA and PAA, respectively (Dai et al 2013; **Fig. 1**). While a role in PAA biosynthesis via the YUC pathway has not been confirmed *in vivo*, Arabidopsis lines overexpressing either YUC1, 2 or 6, all show increased levels of PAA conjugates compared to control lines (Sugawara *et al*, 2015). Arabidopsis YUCs display variability in predicted substrate-binding surface area and volume (**Fig S5**), and in the case of Pea (*P. sativum*), YUCs can successfully take the larger halogenated 4-Cl-IPA substrate to form the highly bioactive 4-Cl-IAA (**Fig. 1**; (Tivendale *et al*, 2012)). This may suggest that YUCs may act on a wider range of substrates but to date, no large screens of potential YUC substrates has been performed, likely hindered by challenges relating to FMO expression and purification (Thodberg & Neilson, 2020).

Dual YUC functionality has been reported for *At*YUC6 (Class 3E) and two poplar (*Populus trichocarpa*) YUCs, *Pt*YUC4 (Class 5B) and *Pt*YUC5 (Class 3B). These YUCs can produce IAA, but also moonlight as NADPH-dependent thiol-reductase enzymes, increasing plant resistance under different abiotic stress conditions (Cha *et al*, 2015; Wang *et al*, 2022a; Cha *et al*, 2022). Lastly, a neo-functionalized ‘YUC-like’ *Bs3* gene is important for plant immunity. Specifically, the pepper (*Capsicum annuum*) *Bs3* gene (Class 4) lacks approximately 70 aa in the region between the NADPH-binding and F/LATGY motifs (Krönauer *et al*, 2019). This neo-functionalized *Bs3* gene encodes an enzyme without auxin capabilities but instead exhibits NADPH oxidase activity (Krönauer & Lahaye, 2021) which aids effector protein recognition against the pathogenic bacterium *Xanthomonas campestris* pv. *vesicatoria* (Römer *et al*, 2007).

### Diversification of YUCCA structure and function

The evolution of protein motifs is influenced by many factors including insertions, deletions, point mutations and selective pressures, and the variation in YUC motif patterns shown here may influence factors such as localization, expression, oligomeric state and protein-protein interaction patterns. YUCs are evolutionarily related to other Class B FMOs (i.e. N-OX, S-OX, sYUC families) and share five major sequence motifs (labelled as f, h, j, k, q; **Fig 4**). When compared to the other class B FMOs, YUCs possess some family-specific motifs (**Fig 5A**) between the NADPH-binding and F/LATGY motifs [i) ‘PREXXG’, ii) ‘Leucine-rich KWLP’ and iii) ‘Glycine-rich GX(5)GXXRPXXGP’ (motifs l or m, n & o; respectively; Fig 5A)] which may indicate important structural regions for their auxin-specific functionality.

Overall, Arabidopsis YUCs possess highly similar structural folds, with many sites under negative selection (**Fig S6**, **Data S7-S8**). Motif pattern variation, however, does exist (**Fig S4**), which could contribute to extended or dual protein functionality. For example, further analysis of different AtYUCs for thiol-reductase activity would identify specific residues responsibility for this moonlighting functionality. Variation in predicted YUC subcellular localization (i.e., cytoplasm, endoplasmic reticulum, plastid, peroxisome, cell membrane, and nucleus) may be important for specific protein-protein interactions based on compartmentalization (**Fig S10**, **Data S10**), especially given that plants TAA/TARs predicted localization is evenly distributed in the endoplasmic reticulum, cytoplasm and also the golgi (**Fig 4**). Comprehensive YUC localization analysis is well documented within Arabidopsis, showing localization to the cytoplasm and ER (Kriechbaumer *et al*, 2012, 2016)**(Fig 6B**). An earlier study also identified *At*YUC6 to occur in an unknown endomembrane compartment (Jeong *et al*, 2007), which here we speculate may be the peroxisome given AtYUC6 (Cha *et al*, 2016, 2015) is predicted to have a peroxisomal targeting signal (**Fig 3B).** Conclusions on YUC localization, however, require empirical support, as the *At*YUC *in silico* predictions do not align with experimental outcomes. Alternative gene splicing may also contribute to YUC sub-functionalization, diversity and evolution. A key example is *At*YUC4 which occurs as two splice variants, experimentally shown to have distinct intracellular localization (Kriechbaumer *et al*, 2012): AtYUC4A is localized in cytoplasm and AtYUC4B in the endoplasmic reticulum (**Fig 6B**). Computational studies also reveal a nuclear localization signal for AtYUC4A, consistent with experimental support showing AtYUC4A in the nucleoplasm (Kriechbaumer et al, 2012), and common for cytosolic proteins.

### Conclusions and Future Perspectives

The critically important YUC family plays a key gate-keeping role in auxin biosynthesis. Here we perform a comprehensive phylogenetic analysis of the YUC family, representing diverse members across the plant kingdom. YUCs are ubiquitous across all plant species, with diversification especially evident in vascular plants. In contrast to the other auxin biosynthetic family TAA/TAR which have evolved in two major directions, vascular YUCs have evolved in multiple. To facilitate classification and interpretation of YUC diversification and evolution we establish and present an expandable nomenclature system for the YUC family, consisting of six distinct classes (Classes ED, 1 – 5), and 41 subclasses (A, B, C, etc.). This expandable naming system is based upon phylogenic positioning and sequence identity cut-offs following similar naming systems of other large gene families. YUC expansion can be attributed to many factors, including the fine-tuned control of auxin biosynthesis in time, space and environmental condition, possible catalytic promiscuity and novel functionalities (i.e. thiol reductase moonlighting capabilities). Experimental evidence of extended functionalities is currently limited and further investigations into the multifunctionality of this family will provide important insights into how and why these genes are retained. Our analysis of YUC sequence diversity revealed variable motif regions which may affect protein folding, stability, localization, and interaction with other proteins. Biochemical characterization of YUCs is extremely limited and a major bottleneck and hinders a more complete understanding of the YUC catalysis mechanisms and substrate selectivity. The evolutionary and structural perspective presented here provides offers new perspectives on the distribution and evolutionary trends of this crucial family, and facilitates future YUC characterization to better understand their role within plant development and environmental change.

## Methodology

### Homologue identification of YUCs and TAA/TARs in Plants

To identify plant YUCs and TAA/TAR homologs, did BLAST search (default parameters) against databases including NCBI (Sayers *et al*, 2023), OneKP (Leebens-Mack *et al*, 2019), Phytozome (Goodstein *et al*, 2012), Phycocosm (Nordberg *et al*, 2014), CNGBdb (Chen *et al*, 2020), PlantGenIE (Sundell *et al*, 2015), Hornwortbase (https://www.hornwortbase.org/), FernBase (https://fernbase.org/) and Takakia database (Hu *et al*, 2023). Arabidopsis YUC and TAA were used as queries (accessions: AtYUC6-AT5G25620.1; AtTAA-AT1G70560.1). We also used the plaBi database’s cladogram view (https://www.plabipd.de/plant_genomes_pa.ep) to get genomic data availability details and primary insights into their phylogenetic relationship. To provide a more comprehensive picture of the evolutionary position of YUC within the green lineage, we included several species from angiosperms, bryophytes, lycophytes, hornworts, liverworts, gymnosperm and green algae. Further, to classify as a prospective hit and to avoid the chance of any false positives, each retrieved putative hit was further validated by reciprocal blast. A limited number of other class B FMOs (S-OX, N-OX and sYUCs) were also retrieved for comparative study. Selected plant species, homologues sequence data and their respective source database details are given in **Data S1**. Ploidy levels were checked and confirmed using PloiDB_web (https://taux.evolseq.net/PloiDB_web) and PhyloGenes (Zhang *et al*, 2020). The species tree was generated using Lifemap (de Vienne, 2016) using taxonomy identifier (TaxId) as input.

### Sequence analysis and phylogenetic studies

The sequence Identity matrix for 469 YUC sequences was constructed using Sequence Demarcation Tool Version 1.3 (SDTv1.3) (Muhire *et al*, 2014) by employing the MAFFT (Katoh & Standley, 2013) aligning algorithm with the Neighbour-joining (NJ) tree function enabled. Percent Identity Matrix for each YUC class and subclass were done in Clustal Omega (Sievers & Higgins, 2018). All the matrix details are provided in **Data S2**. The retrieved and validated YUC and TAA/TAR homologs were aligned in MAFFT v.7 (Katoh & Standley, 2013; Katoh *et al*, 2019) and visualized in ESpript 3.0 (Robert & Gouet, 2014). ClustalW from MEGA11 (Tamura *et al*, 2021) was also used to check the percentage-level-based sequence conservation. Further, to investigate the evolutionary relationships of YUCs and TAA/TARs, phylogenetic trees were constructed in IQ-TREE (both web server and standalone versions; Maximum Likelihood (ML) method with 1000 bootstraps) (Trifinopoulos *et al*, 2016; Hoang *et al*, 2018; Minh *et al*, 2020). The MAFFT-aligned sequences were used as input. Very short/partial fragments were omitted based on the sequence length. IQ-TREE-generated trees were visualized using iTOL v.6(Letunic & Bork, 2024a). The YUC phylogram includes 469 sequences retrieved from 54 diverse plant species, and the TAA/TAR tree includes 254 sequences from the same species. The YUC tree was rooted using bryophytes (Early Diverging (ED) clade) (Harris *et al*, 2022; Qiu & Mishler, 2024). Further, YUCs were classified into specific classes based on the clustering pattern in the phylogram and sequence identity matrix. The DIVEIN server was also used to calculate and confirm the diversity of different YUC classes, and subclasses (and between different plant groups) using the Pairwise Hamming distance (Deng *et al*, 2010). For class-specific sequence conservation/motif studies, class-based YUCs were extracted from the phylogram using the fasta extractor program in the FaBox (1.61) toolbox (Villesen, 2007). Selection pressure and diversification analysis were conducted on the Datamonkey server with the MEME (Mixed Effects Model of Evolution) and SLAC (Single-Likelihood Ancestor Counting) modules (Weaver *et al*, 2018; Murrell *et al*, 2012; Kosakovsky Pond & Frost, 2005), using AtYUCs (both amino acid and nucleotide sequences) as the models. Codon alignment was generated in PAL2NAL program (Suyama *et al*, 2006) MEME analysis (v4.0) was executed on the AtYUC codon alignment (excluding stop codons) using HyPhy v2.5.64 (Kosakovsky Pond *et al*, 2020).

### Structural conservation pattern, motif analysis and subcellular localization

To examine the pattern of sequence conservation in the structure, ConSurf analysis (Yariv *et al*, 2023b) was done using all the chosen YUC sequences. Results were subsequently mapped into AtYUC6 as the representative model generated in the built-in Modeller (Webb & Sali, 2016) within the Consurf. Besides, to check the domain architecture of YUC/TAA sequences, a SMART search (Letunic *et al*, 2021) was done. Further, MEME Suite 5.5.5 (Bailey *et al*, 2015) was employed to identify and examine the conserved sequence-specific motifs of YUCs with 30 cut-offs. DeepLoc 2.1 (Ødum *et al*, 2024) was used for predicting the subcellular localization and all localization details were mapped into the phylogram.

### Gene localization, expression and interaction analysis

The chromosomal locations of all Arabidopsis *YUC* genes were obtained from the ePlant database (Waese *et al*, 2017). To explore the organ-specific expression of Arabidopsis YUCs the Plant Expression Omnibus web-based tool (PEO)(Koh *et al*, 2024) was used. The expression pattern of all 11 *AtYUCs* were estimated by analysing the mean value of TPM values (transcripts per million) of distinctively expressed samples in different stages of Arabidopsis development. Gene expression data for various environmental stresses was obtained from the eFP browser (Winter *et al*, 2007). The expression levels of *YUC* genes under different abiotic and biotic stresses were analysed using relative values of Log2 Ratios. Expression levels were examined under multiple abiotic stress conditions, including heat, cold, drought, salt, genotoxic, and oxidative stress. For biotic stress analysis, the Log2 Ratios of one representative treatment was selected for each stress type to ensure consistent and comparable data across all AtYUC genes. For *Botrytis cinerea*, *Phytophthora infestans*, and *Erysiphe orontii*, gene expression data was analysed using the only representative treatment available for each pathogen. For *Pseudomonas syringae*, Log2 ratios derived from triplicate infiltrated leaf samples treated with a virulent strain were estimated. Elicitor stress was assessed using Log2 ratios from a 10 µM HrpZ treatment, a bacterial-derived elicitor. The overall expression level was determined as the mean of Log2 ratios for both abiotic and biotic stress conditions, based on visual inspection of the data. Further, to check YUC interaction patterns, protein-protein interactions were analyzed in the STRING predicted database (Szklarczyk *et al*, 2023) using AtYUCs as models. [Parameters used for STRING search: High confidence (interaction score >0.700); active interaction sources used: Text mining, expression, Databases (IntAct, BioGRID etc), co-expression].

### Molecular modelling and docking

To investigate structural features and probable variations of YUCs between phylogenetic classes, protein models were generated for representative AtYUCs (AtYUCs) using AlphaFold (Jumper *et al*, 2021). Structural refinement was done in GalaxyWEB using GalaxyRefine option (Ko *et al*, 2012), and all models were validated using Verify 3D, PROCHECK, ERRAT and ProSA-web (Eisenberg *et al*, 1997; Laskowski *et al*, 1993; Wiederstein & Sippl, 2007; Dym *et al*, 2006). Molecular docking was done in AutoDock 4.2.6 and PyRX software (Garrett M. Morris, 2010; Dallakyan & Olson, 2015). FAD, NADPH, IPyA, PPA, and the auxin inhibitor yucasin were used as ligands. Energy minimization was done using uff (United Force Field). Further, the cavity parameters of AtYUC models were computed in CASTpFold (Ye *et al*, 2024), and the electrostatic surface potential (ESP) was calculated using ASPB electrostatics plugin (version 2.1) in PyMOL 2.5.4 (Academic licensed version) (DeLano, 2002; Schrodinger LLC, 2015; Jurrus *et al*, 2018).

## Structured Methods - Tools Table

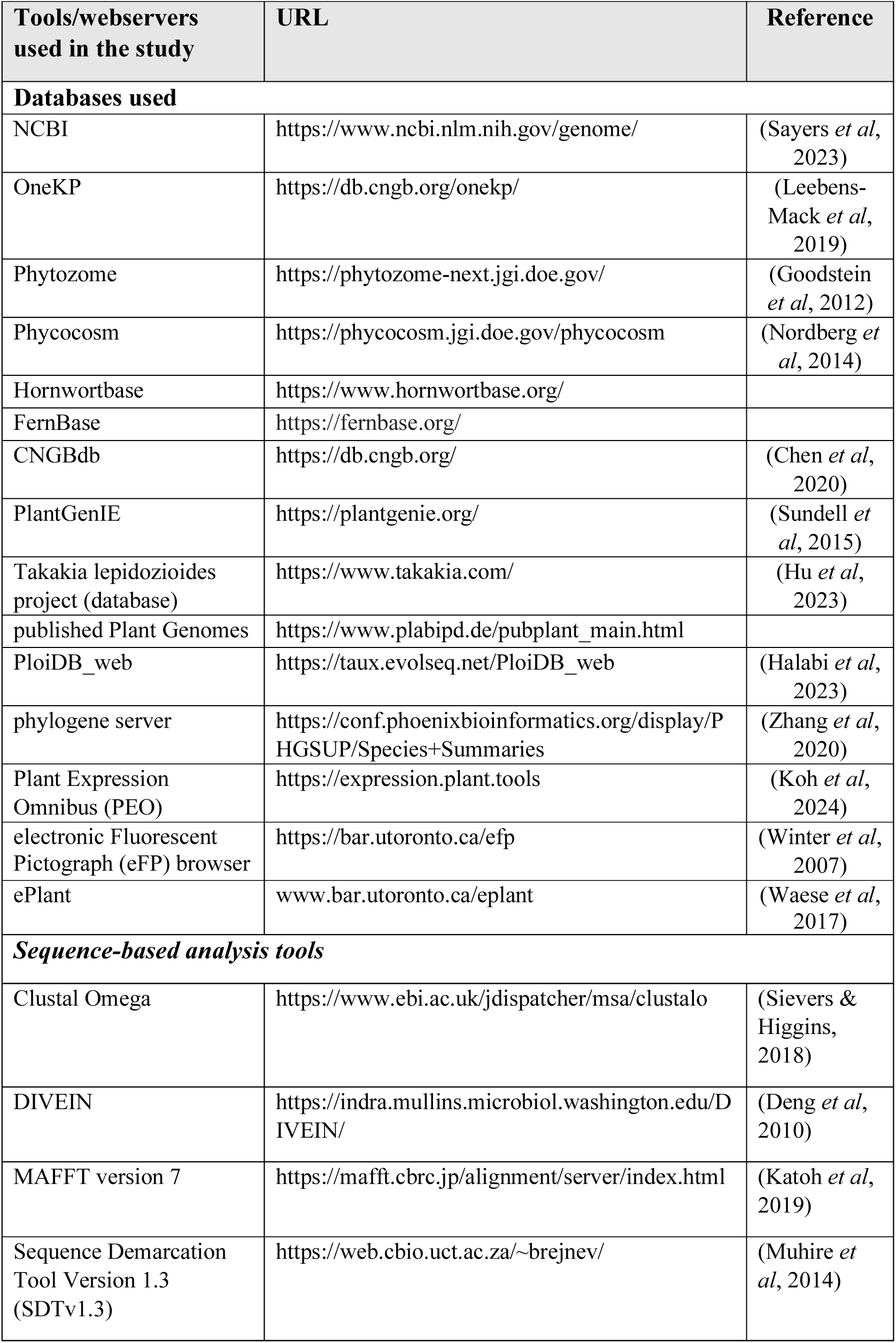
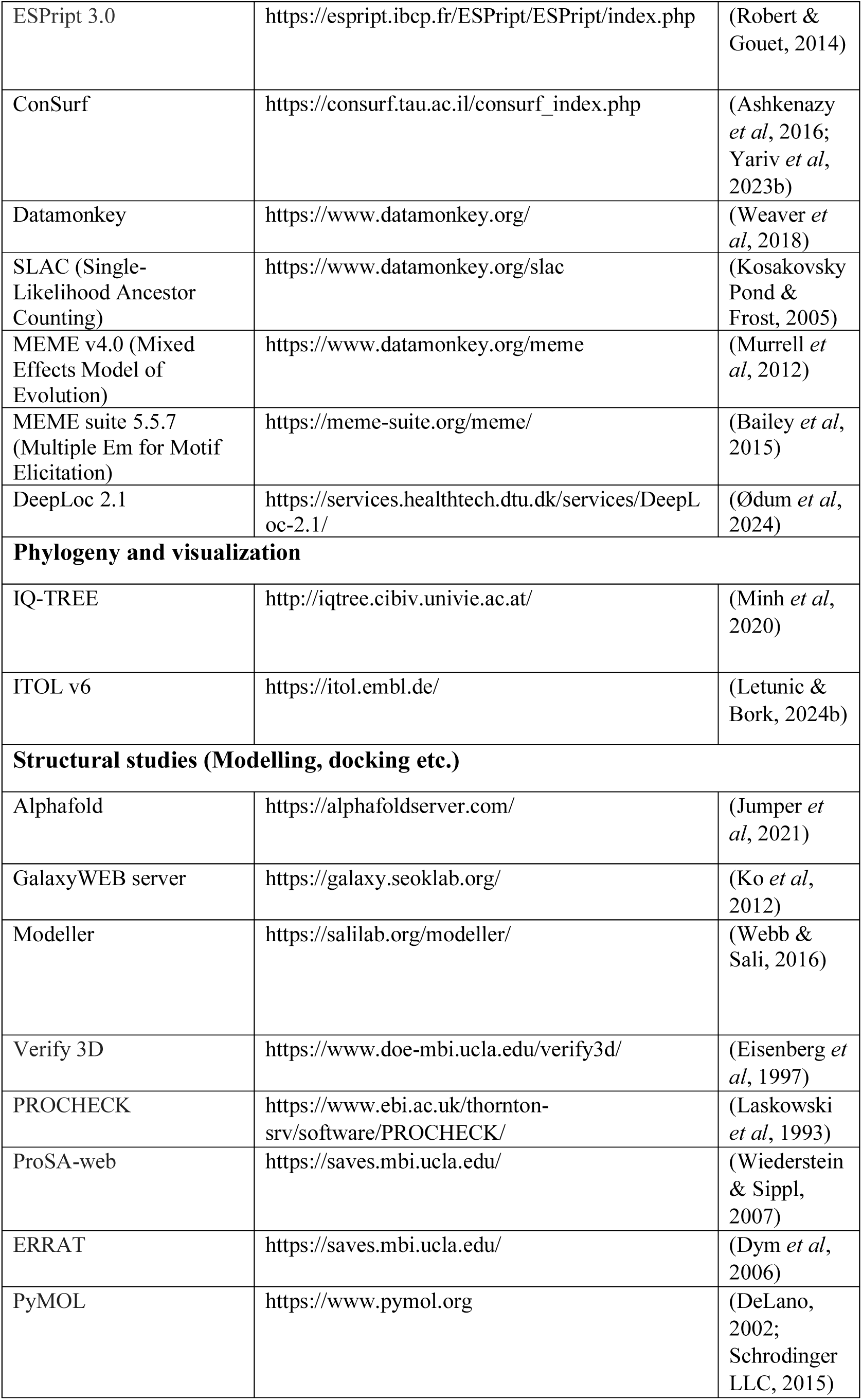
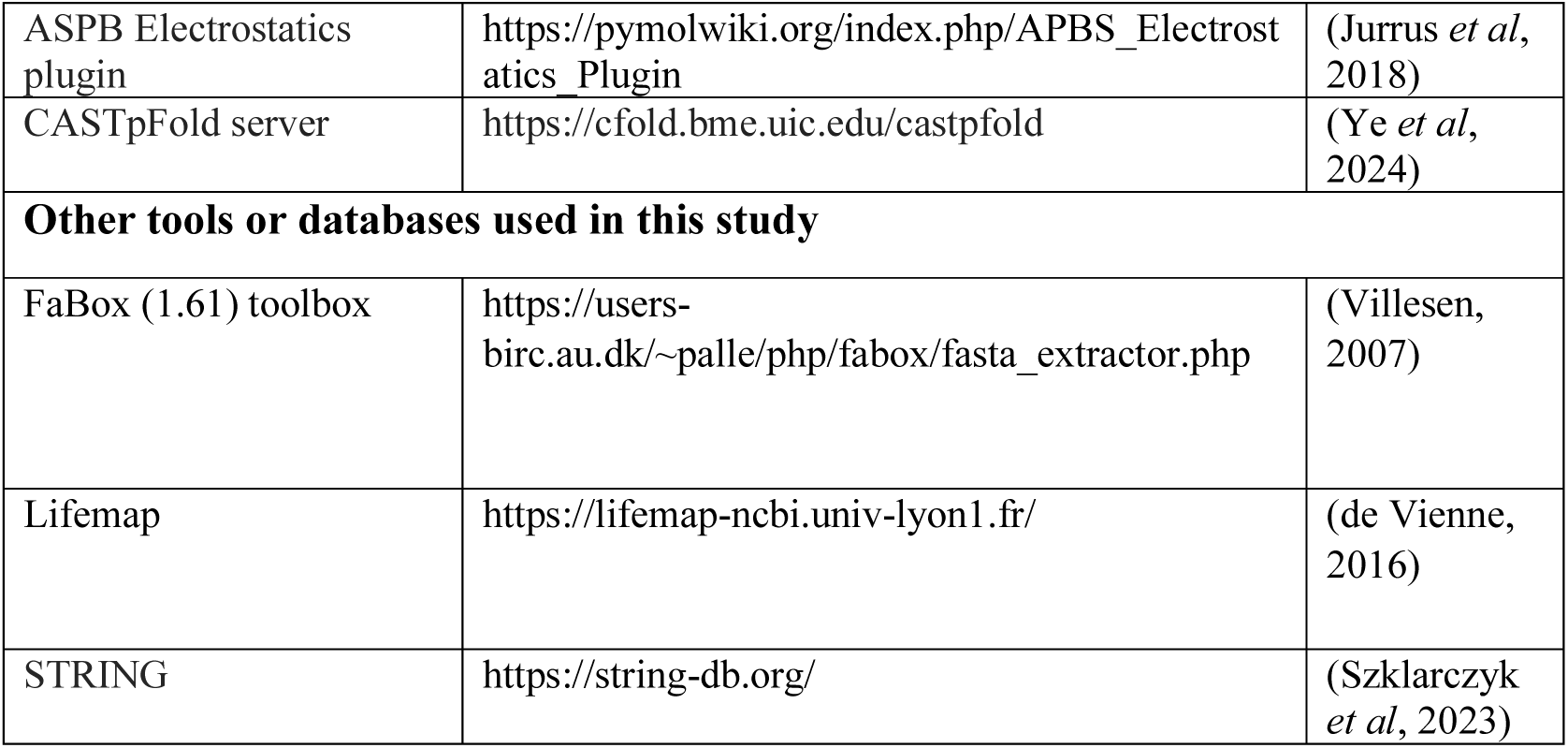

## Author Contribution

**Mallika Vijayanathan**: Conceptualization; Data curation; Investigation; Methodology; Writing—original draft; Writing—review and editing. **Amna Faryad**: Investigation, Writing —review and editing. **Thanusha D Abeywickrama:** Investigation, Writing—review and editing. **Joachim Møller Christensen**: Investigation, Writing—review and editing. **Elizabeth Heather Jakobsen Neilson**: Conceptualization; Supervision; Funding acquisition; Project administration; Writing—review and editing.

## Acknowledgments

This work was supported by the Independent Research Fund Denmark (Grants 1051-00083B & 1131-0002B) and the Novo Nordisk Foundation (Grant No. 0054890) awarded to EHJN. MV acknowledges the Marie Skłodowska-Curie Individual Fellowship (MSCA grant agreement No. 101110417). The authors also thank Kristian Fønlev Nicolajsen for the help in collating the motifs presented in figure 5.

## Supplementary Materials

**Supplementary Information** – Figures and Tables (S1-S2) (PDF document)

**Supplementary Data** (**Data S1-S10**) (Excel format, zip files, pdf)

**Data S1**: Sequence data (YUCs, sYUCs, other FMOs) used in the study & Respective source database details (Excel & fasta format files – Zip file)

**Data S2**: Percent Identity Matrix for YUCs in each Classes and Subclasses (All details used for YUC enzyme classification strategies)- Excel

**Data S3**: Multiple sequence alignment files – all source files (Zip)

**Data S4:** Sequence alignment – Graphical representation (for YUCs, sYUCs and comparison with other class B FMOs)- pdf

**Data S5:** MEME motif analysis result files for YUCs with other class B FMOs (S-OX, N-OX, sYUCs) pdf

**Data S6:** MEME Extracted motif sequence area for YUCs (Excel)

**Data S7:** AtYUC protein and codon alignment with selection pressure details (Pdf)

**Data S8:** Evolutionary statistics calculation results (for AtYUCs) from Datamonkey (Excel)

**Data S9**: Subcellular localization data for YUCs, TAA/TARs, signal peptide, disorder predictions etc. (Excel)

**Data S10**: Gene expression and interaction data for Arabidopsis YUCs (Excel)

**Expanded View Figure legends**

**Fig EV1. Expanded View of YUC phylogram**

**Fig EV2. Expanded View of TAA/TAR phylogram**

