## Supplementary Data (Data S1-S10) for "The auxin gatekeepers: Evolution and diversification of the YUCCA family": Data S4.pdf

### Sequence alignment files

1. Sequence alignment for selected S-OX, N-OX, sYUCs and YUCs - to get an insight on conservation pattern (page 1 - 5)
2. Sequence alignment for all YUCs- Class based (Page 6 onwards)

| | |
|---|---|
| ED class - Page 6 -8 | Class 3 - Page 21-29 |
| Class 1 - Page 9-19 | Class 4 - Page 30-40 |
| Class 2 - Page 20 | Class 5 - Page 41-44 |

3. Sequence alignment for all sister YUCCAs (sYUCs) (Page 45 - 55)

#### 1. Sequence alignment (selected FMOs – S-OX, N-OX, Sister YUCs, and ÝUCs).

Alignment was done in MAFFT version 7 (https://mafft.cbrc.jp/alignment/server/, July 2024) and visualized in ESPript 3.0 (https://espript.ibcp.fr/ESPript/cgi-bin/ESPript.cgi). All used sequences were manually curated. Sequence details and alignment given below.

**Selected FMO sequences details**

| Name | Accession | Species |
|---|---|---|
| AsFMO (S-OX) | BAS32646 | *Allium sativum* |
| AtFMO (S-OX) | NP001321123.1 | *Arabidopsis thaliana* |
| OsFMO (S-OX) | XP015612837 | *Oryza sativa* |
| HvFMO (S-OX) | XP044971712.1 | *Hordeum vulgare* |
| SmFMO (S-OX) | XP2973683.1 | *Selaginella moellendorffii* |
| MpFMO (S-OX) | PTQ49826.1 | *Marchantia polymorpha* |
| PcFMO (S-OX) | scaffold2006551 | *Phaeoceros carolinianus* |
| AtFMO1(N-OX) | AT1G19250.1 | *Arabidopsis thaliana* |
| OsFMO1(N-OX) | LOCOs03g08410.1 | *Oryza sativa* |
| HvFMO1(N-OX) | HORVU4Hr1G077170 | *Hordeum vulgare* |
| SmFMO1(N-OX) | XP002967925.1 | *Selaginella moellendorffii* |
| PcFMO1(N-OX) | scaffold2088977 | *Phaeoceros carolinianus* |
| SisterYUCCA-Pc | HW-WCZB-2015184 | *Phaeoceros carolinianus* |
| SisterYUCCA-Pc | HW-WCZB-2016004 | *Phaeoceros carolinianus* |
| SisterYUCCA-Mp | Mapoly0052s0062 | *Marchantia polymorpha* |
| SisterYUCCA-Mp | Mapoly0122s0026 | *Marchantia polymorpha* |
| SisterYUCCA-Sm | Sm113792 | *Selaginella moellendorffii* |
| SisterYUCCA-Sm | EFJ16321.1 | *Selaginella moellendorffii* |
| Ed-c-MpYUC1 | Mapoly0127s0005.1.p | *Marchantia polymorpha* |
| Ed-d-PcYUCA | scaffold 2017088 | *Phaeoceros carolinianus* |
| ED-e-SmYUCA | 80431 | *Selaginella moellendorffii* |
| Class-1L-AtYUC11 | AT1L21430.1 | *Arabidopsis thaliana* |
| Class-1Q-AtYUC10 | AT1G48910.1 | *Arabidopsis thaliana* |
| Class-3e-AtYUC2 | AT4G13260.1 | *Arabidopsis thaliana* |
| Class-3e-AtYUC6A | AT5G25620.1 | *Arabidopsis thaliana* |
| Class-3e-AtYUC6B | AT5G25620.2 | *Arabidopsis thaliana* |
| Class-4f-AtYUC3 | AT1G04610.1 | *Arabidopsis thaliana* |
| Class-4f-AtYUC5 | AT5G43890.1 | *Arabidopsis thaliana* |
| Class-4f-AtYUC7 | AT2G33230.1 | *Arabidopsis thaliana* |
| Class-4f-AtYUC8 | AT4G28720.1 | *Arabidopsis thaliana* |
| Class-4f-AtYUC9 | AT1G04180.1 | *Arabidopsis thaliana* |
| Class-5b-AtYUC1 | AT4G32540.1 | *Arabidopsis thaliana* |
| Class-5B-AtYUC4A | AT5G11320.1 | *Arabidopsis thaliana* |
| Class-5B-AtYUC4B | AT5G11320.2 | *Arabidopsis thaliana* |
| Class-2a-AfYUCB | s0087.g042257 | *Azolla* |

Alignment is given below:-

```
                              1        10
AsFMO_S-OX                    MVSSSCSSIPK.....................................................................
AtFMO_S-OX                    M...............................................................................
OsFMO_S-OX                    MKT.............................................................................
HvFMO_S-OX                    ................................................................................
SmFMO_S-OX                    M...............................................................................
MpFMO_S-OX                    MIIQAAFRPPF....SSARRTACTSALNR.....SGAKS...QLVDRSRRLFDED..........FVPTLSSRGGREL..
PcFMO_S-OX                    ................................................................................
AtFMO1_N-OX                   MASNYDK.........................................................................
OsFMO1_N-OX                   MAAAQQQQKE...........................................................GGARRTREE..
HvFMO1_N-OX                   MAMAQGQ...............................................................AARATREA..
SmFMO1_N-OX                   MGDDLGF....................................DL...............................EF..
PcFMO1_N-OX                   ................................................................................
SisterYUCCA_P.carolinianus    ................................................................................
SisterYUCCA-P.carolinianus    MAATDS.............VRAWLADFS......DALQK...QDVNAVVELFQEEECFWRDLLAFTWNIYTAESKDEVAE
SisterYUCCA_Marchantia        MASNGTLENGSS..DNSKIVKAWLSYLD......EALQK...QDIAHLVTLFDEEECFWRDMLAFTWNLYTAESRDEIAA
SisterYUCCA_Marchantia        MGSEELSAGQK........ITSWLSEFD......TVLQD...EDVDQILELFDDQECFWRDLLPFTWNIETFESKDEILK
SisterYUCCA_S.moellendorffii  MAT....VNGSPKIDPDALVDAWLARLD......SSLAS...GDASAAANLFEESTSFWRDLIAFTWNIVTVEGRSQIHA
SisterYUCCA_S.moellendorffii  M.......NGSPKVDPDALVDAWLARLD......SSLAS...GDASAAANLFEESTSFWRDLIAFTWNIVTVEGRSQIHA
ED_c_MpYUC1                   MAARLGFCLGSL...........LSFLDIGWKLCSGLTRAVLTNLCNAVGIFSDR.CL...MSPTLTDVYDISASSEVNN
ED_d_PcYUCA                   EAE.........................................................................DEF..
ED-e-SmYUCA                   ................................................................................
Class-1L-AtYUC11              ................................................................................
Class-1Q-AtYUC10              ME..............................................................................
Class-3e-AtYUC2               ME..............................................................................
Class-3e-AtYUC6A              MDF.............................................................................
Class-3e-AtYUC6B              MDF.............................................................................
Class_4f-AtYUC3               MYG.............................................................................
Class_4f-AtYUC5               MEN...............................................MFR...........................
Class_4f-AtYUC7               MCN.............................................................................
Class_4f-AtYUC8               MEN...............................................MFR...........................
Class_4f-AtYUC9               MEN...............................................MFR...........................
Class-5b_AtYUC1               ME..............................................................................
Class-5B-AtYUC4A              MGT.............................................................................
Class-5B-AtYUC4B              MGT.............................................................................
Class-2a-AfYUCB               MMKAFDLW........................................................................

AsFMO_S-OX                    ................................................................................
AtFMO_S-OX                    ................................................................................
OsFMO_S-OX                    .....................................................................PQNDKLQYHTP
HvFMO_S-OX                    ................................................................................
SmFMO_S-OX                    ................................................................................
MpFMO_S-OX                    .....................................................................LRNEGVSDRKP
PcFMO_S-OX                    ................................................................................
AtFMO1_N-OX                   ................................................................................
OsFMO1_N-OX                   ................................................................................
HvFMO1_N-OX                   ................................................................................
SmFMO1_N-OX                   ................................................................................
PcFMO1_N-OX                   ................................................................................
SisterYUCCA_P.carolinianus    ................................................................................
SisterYUCCA-P.carolinianus    MLKATLSEVKPDLWEVDGEA..VEVGGTVDAMLKFETAVGRGRGHVRLKGNK.....CWTMFTALRELKGHEEKVGKARP
SisterYUCCA_Marchantia        MMNSTLATVKPSGWELDGEV..EEKGGALQVFLKFETSIARGRGHLRLKGGK.....CWTLLTAMTELKGYEEHLGSARP
SisterYUCCA_Marchantia        MLTENLARVKPRAFVIDGDV.IDRPNGLMGSAFKFETSLAWCRGFVWLKGGK.....CWTITTSMWDLKGYEEKISRSRP
SisterYUCCA_S.moellendorffii  MLQSTLESVAPKGWVRNGTASYSADSGVIEAWIKFETRDAVCTGHLRLMATT...SLCRTLLTAMEGLRDFPENKGRTRP
SisterYUCCA_S.moellendorffii  MLQSTLESVAPKGWVRNGTASYSADSGVIEAWIKFETRDAVCTGHLRLVATT...SLCRTLLTAMEGLKDFPEKKGHTRP
ED_c_MpYUC1                   SGQTSQS.........................................KGPENDSSDAPKSICGSKSSQFVDKTSGSKHG
ED_d_PcYUCA                   ............................................................YQQRERGQQQRRRMSSRKQP
ED-e-SmYUCA                   ................................................................................
Class-1L-AtYUC11              .........................................................................MEKEIKI
Class-1Q-AtYUC10              ................................................................................
Class-3e-AtYUC2               ............................................................FVTETLGKRIHDPYVEETRC
Class-3e-AtYUC6A              .........................................................CWKREMEGKLAHDHRGMTSPRRI
Class-3e-AtYUC6B              .........................................................CWKREMEGKLAHDHRGMTSPRRI
Class_4f-AtYUC3               ....................................................NNNKKSINITSMFQNLIPEGSDIFSRRC
Class_4f-AtYUC5               ...................................................................LMGSEDSSDRRRC
Class_4f-AtYUC7               ....................................................NNNTSCVNIS.....SMLQPEDIFSRRC
Class_4f-AtYUC8               ...................................................................LMDQDQDLTNNRC
Class_4f-AtYUC9               ...................................................................LMASEEYFSERRC
Class-5b_AtYUC1               ...................................................................SHPHNKTDQTQHI
Class-5B-AtYUC4A              .........................................................C...............RESEPTQ
Class-5B-AtYUC4B              .........................................................C...............RESEPTQ
Class-2a-AfYUCB               .......................................................................GDEEPMNSL

                                                                      (f)
                                                                     GXGXXG
                                                               20        30        40        50        60
AsFMO_S-OX                    .........................MPVTPLSLVTRHVAIIGAGAAGLVTARELRR..EGH.TTTIFERGSSIGGTWIY.
AtFMO_S-OX                    .........................VPAVNPPTTSNHVAVIGAGAAGLVAARELRR..EGH.SVVFERGNHIGGVWAY.
OsFMO_S-OX                    PLPS..KSQQKS...EAKSRTHRLAMPSP.....SLRLAVVGAGAAGLVAARELRR..EGH.SPVVFERAASVGGTWLY.
HvFMO_S-OX                    .........................MPSP.....SLRLAVVGAGAAGLAAARELRR..EGH.APVVFERAAAVGGTWLYA
SmFMO_S-OX                    ..................................EKRVAVIGAGASGLVAARELLR..EGH.SVVIFEQARRIGGTWVY.
MpFMO_S-OX                    SILV..RFKGVD...EMTREEDGRSGAAE.....IKRVLVIGAGAAGLAAAVELAR..EGH.DVVVYEKSSEIGGVWNY.
PcFMO_S-OX                    .....................................VAVIGAGAAGLVAARELAR..EGH.EVVAFEQSGHVGGVWVY.
AtFMO1_N-OX                   ................................LTSSRVAIIGAGVSGLAAAKNLVH..H...NPTVFEASDSVGGVWR..
OsFMO1_N-OX                   .......VPA........................VGRVAIIGGGISGLAAAKQLAA..H...DPVVFEATPHIGGVWK..
HvFMO1_N-OX                   .......VPL........................VSRVAIIGGGISGLAAAKQMSA..Y...DPVVFEATPSVGGVWK..
SmFMO1_N-OX                   .......SPA........................AARVCVVGAGVSGLCACRHLLK..RGI.RPTVLEGSSHIGGVWR..
PcFMO1_N-OX                   ....................................RVCIVGAGVSGLVACKYLSREAEGW.EATVVEGQAGLGGIWSG.
SisterYUCCA_P.carolinianus    ................................................................................
SisterYUCCA-P.carolinianus    TGLVYGAVPGRKTWREERQEEEGKLGYSE.....QPYCVVVGGGQSGIALGARLRK..LGV.PTIIVEKNERPGDSWR..
SisterYUCCA_Marchantia        MGVKNENVPGRKTWLEERMQERQELGYKS.....QPYCVIVGGGQAGIALAARLRM..LNV.PALIIEKNARPGDSWR..
SisterYUCCA_Marchantia        FGPTLGQIKGRKTAYQEKQQEQHDMGRSK.....QPYCLVVGAGQGGVMLGARLRM..LGV.PAIIVEKNERLGDNWR..
SisterYUCCA_S.moellendorffii  NGVTHGVIRHRASWLDGRKEEERTLGSTV.....QPYCVIVGGGQAGIGLAARLRQ..LGV.PCIVVEKNPRPGDSWR..
SisterYUCCA_S.moellendorffii  KGVTHGVIRHRASWLDGRKEEERTLGSTV.....QPYCVIVGGGQAGIGLAARLRQ..LGV.PCIVVEKNPRPGDSWR..
ED_c_MpYUC1                   TMSR..PVEG...........................AIIVGAGTSGLAAAACLRE..RGV.PITLLEKSGCIGSLWK..
ED_d_PcYUCA                   VL.....VEG...........................PIIVGAGPAGLAAAASLKD..KDI.PSLIIERADCIASLWK..
ED-e-SmYUCA                   MW.....VDG...........................AIIVGAGPSGLATAACLSA..AGIGSSVILEKNSCIGSLWQ..
Class-1L-AtYUC11              LV....................................LIIGAGPAGLATSACLNR..LNI.PNIVVERDVCSASLWK..
Class-1Q-AtYUC10              ...................................TVVVIVGAGPAGLATSVCLNQ..HSI.PNVILEKEDIYASLWK..
Class-3e-AtYUC2               LM.....IPG...........................PIIVGSGPSGLATAACLKS..RDI.PSLILERSTCIASLWQ..
Class-3e-AtYUC6A              CV.....VTG...........................PVIVGAGPSGLATAACLKE..RGI.TSVLLERSNCIASLWQ..
Class-3e-AtYUC6B              CV.....VTG...........................PVIVGAGPSGLATAACLKE..RGI.TSVLLERSNCIASLWQ..
Class_4f-AtYUC3               IW.....VNG...........................PVIVGAGPSGLAVAAGLKR..EGV.PFIILERANCIASLWQ..
Class_4f-AtYUC5               IW.....VNG...........................PVIVGAGPSGLATAACLRE..EGV.PFVVLERADCIASLWQ..
Class_4f-AtYUC7               IW.....VNG...........................PVIVGAGPSGLAVAADLKR..QEV.PFVILERANCIASLWQ..
Class_4f-AtYUC8               IW.....VNG...........................PVIVGAGPSGLATAACLHE..QNV.PFVVLERADCIASLWQ..
Class_4f-AtYUC9               VW.....VNG...........................PVIVGAGPSGLATAACLHD..QGV.PFVVVERSDCIASLWQ..
Class-5b_AtYUC1               IL.....VHG...........................PIIIGAGPSGLATSACLSS..RGV.PSLILERSDSIASLWK..
Class-5B-AtYUC4A              IF.....VPG...........................PIIVGAGPSGLAVAACLSN..RGV.PSVILERTDCLASLWQ..
Class-5B-AtYUC4B              IF.....VPG...........................PIIVGAGPSGLAVAACLSN..RGV.PSVILERTDCLASLWQ..
Class-2a-AfYUCB               CF.....ASG...........................PIIVGAGPSGLAVAASLRL..LNI.PSLILERSDGIASLWR..
```

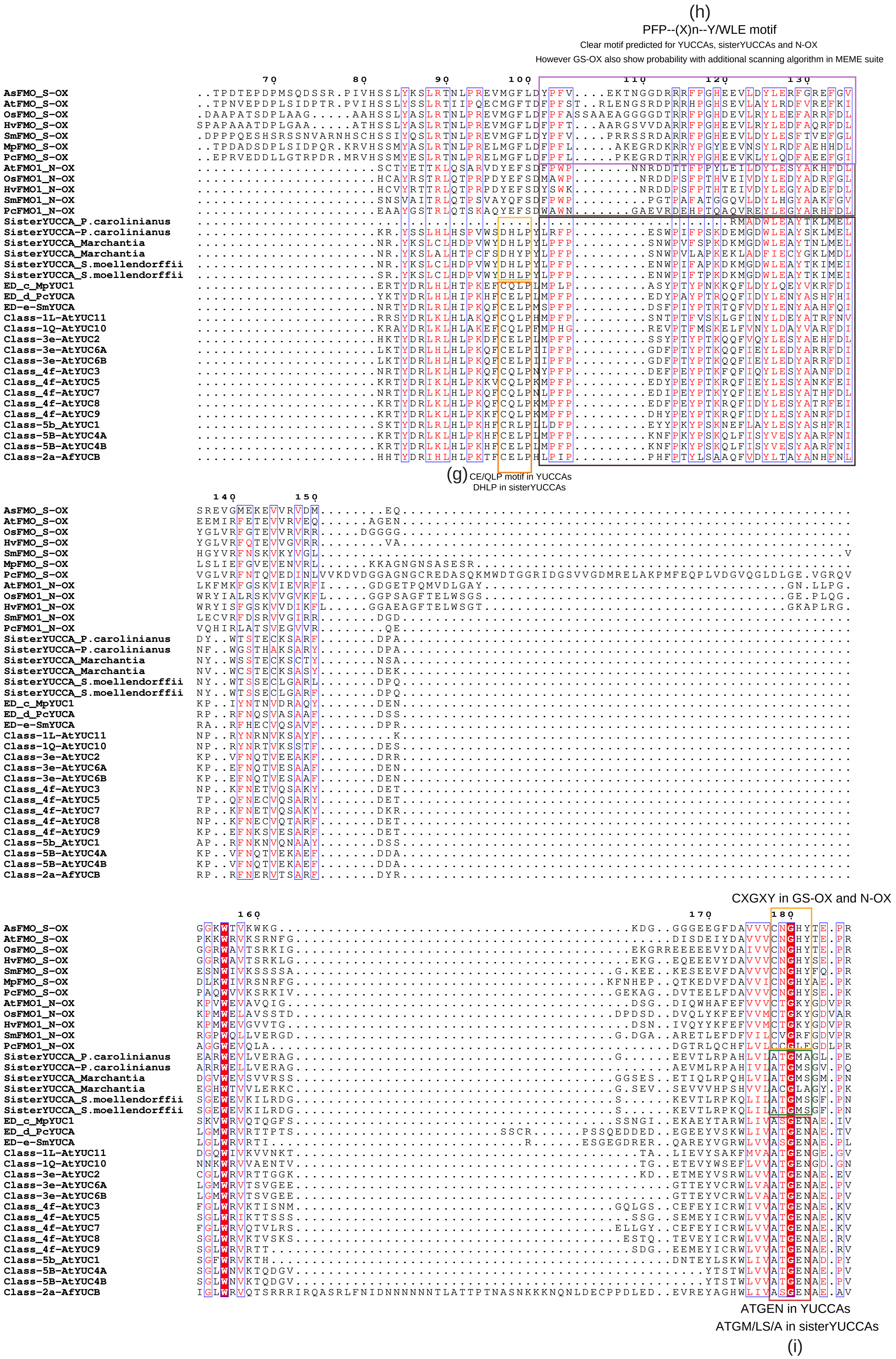
(h)
PFP--(X)n--Y/WLE motif
Clear motif predicted for YUCCAs, sisterYUCCAs and N-OX
However GS-OX also show probability with additional scanning algorithm in MEME suite
(g) CE/QLP motif in YUCCAs
DHLP in sisterYUCCAs
CXGXY in GS-OX and N-OX
ATGEN in YUCCAs
ATGM/LS/A in sisterYUCCAs
(i)

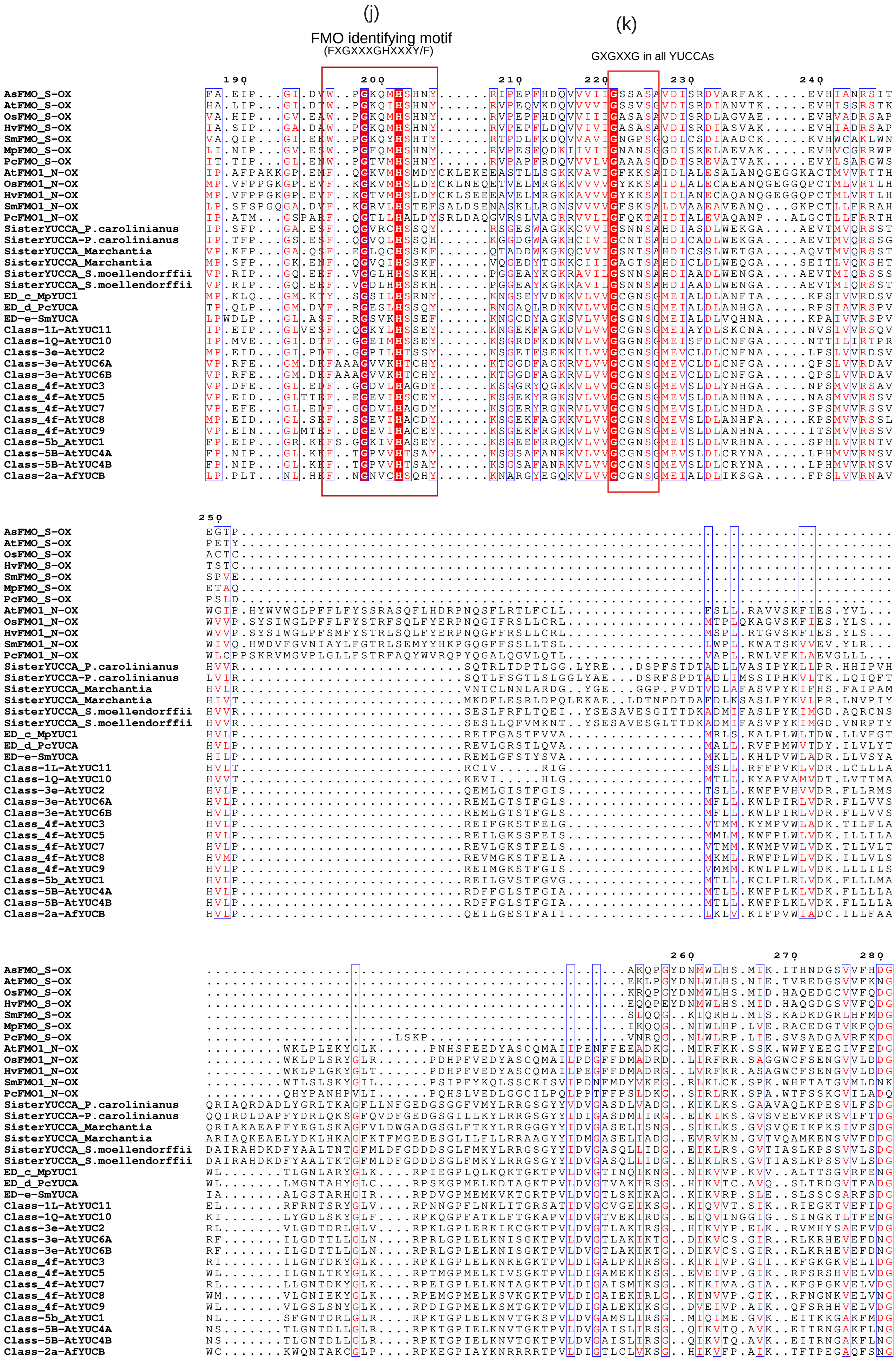
(j)
FMO identifying motif
(FXGXXXGHXXXY/F)
(k)
GXGXXG in all YUCCAs
AsFMO_S-OX
AtFMO_S-OX
OsFMO_S-OX
HvFMO_S-OX
SmFMO_S-OX
MpFMO_S-OX
PcFMO_S-OX
AtFMO1_N-OX
OsFMO1_N-OX
HvFMO1_N-OX
SmFMO1_N-OX
PcFMO1_N-OX
SisterYUCCA_P.carolinianus
SisterYUCCA-P.carolinianus
SisterYUCCA_Marchantia
SisterYUCCA_Marchantia
SisterYUCCA_S.moellendorffii
SisterYUCCA_S.moellendorffii
ED_c_MpYUC1
ED_d_PcYUCA
ED-e-SmYUCA
Class-1L-AtYUC11
Class-1Q-AtYUC10
Class-3e-AtYUC2
Class-3e-AtYUC6A
Class-3e-AtYUC6B
Class_4f-AtYUC3
Class_4f-AtYUC5
Class_4f-AtYUC7
Class_4f-AtYUC8
Class_4f-AtYUC9
Class-5b_AtYUC1
Class-5B-AtYUC4A
Class-5B-AtYUC4B
Class-2a-AfYUCB

(q)

F/LATGY motif

```
                                   290       300                         310                           320       330
AsFMO_S-OX                         CSVHVDVIMHCTGYVYNFP.FLN.TN.................GIVTVDDNRVGP.......LYKHVFPPLLAPSLSFVG
AtFMO_S-OX                         KTVYADTIMHCTGYKYYFP.FLD.TK.................GEVTVEDNRVGP.......LYKHVFPPALSPGLSFIG
OsFMO_S-OX                         SSIKADVIMHCTGYLYDFP.FLE.DD.................SAITVDDNCVDP.......LYKHVFPPEVAPHLSFIG
HvFMO_S-OX                         SSIKADVIMHCTGYLYDFP.FLG.DD.................STITVDDNCVDP.......LYKHVFPIEVAPDLSFIG
SmFMO_S-OX                         TSAVVDAILHCTGYLYDFS.FLD.TK.................NYVKVEDNRVGP.......LFKHVFPPALAPSLSFIG
MpFMO_S-OX                         KSTRADVILHCTGYLYSFP.FLD.TE.................GVVTVKDGRIDP.......LYKHVFVPAFGPTLSFVG
PcFMO_S-OX                         STVFANTIIFCTGYVYSFP.FLH.SH.................GCVSVEENQVGP.......LYQHVFPPALAPTLSFVG
AtFMO1_N-OX                        TTLEADVVILATGYDGKKK..LK.....AIVPEPFRT........WLEFPSGVMP.......LYRGTIHPLI.PNMGFVG
OsFMO1_N-OX                        THVDADLVFLATGFEGKDK..LR.....SVLPEPFRG........LVVNKSSMMP.......LYRGTIHPLI.PNMAFVG
HvFMO1_N-OX                        TKVEADLVFLATGFEGKDK..LR.....EVLPKPFRD........LVVGKSSMMP.......LYRGTIHPLI.PNMAFVG
SmFMO1_N-OX                        RELKADVVILCTGFEGEVK..LK.....AILPERYAE........VLQGPDGVVP.......LYRGMIHPRI.PGMAILG
PcFMO1_N-OX                        TQLEADVIIMCTGYDGVGK..LR.....NLLPPPYRQ........LLTWSSGLLP.......LYRGVIHPDI.PDMAFLG
SisterYUCCA_P.carolinianus         SELQADLIVFATGF.GSMNGWVA.....RLISQEVAA..........................................G
SisterYUCCA-P.carolinianus         TELPADLIVYATGY.TSMNNVAA.....KLISQEVADKVGKCWGLGSGTTRDPGPWERELRNMWKPTHQ....EGLWFHG
SisterYUCCA_Marchantia             SELPADLIVYATGY.GNMNQWAA.....KLISQEVADKVGMVWGLGSNTTNDPGPWEKELRNMWKPTQQ....PGLWFMG
SisterYUCCA_Marchantia             SEVSADVIIYATGY.GNMQEFVE.....KLISKEAAEKVGLCWGLGSGFKGDPGPWVGELKNMYRPLPV....PALWILS
SisterYUCCA_S.moellendorffii       TEIAADVVVLATGY.GSMNQWAA.....KLISQEVADRVGKCWGLGSNTRKDPGPWEGELRNMWKPTAQ....EGLWFHG
SisterYUCCA_S.moellendorffii       TEIAADVVILATGY.GSMNQWAA.....KLISQEVADRVGKCWGLGSNTRKDPGPWEGELRNMWKPTAQ....EGLWFHG
ED_c_MpYUC1                        EFESYDAIIFATGYRSNVPMWLK.GN..FFG..............EDGSPSASMR......DGWKGE......RGLYAIG
ED_d_PcYUCA                        TCGDFDCIIWATGFRSNVPRWLK.DD..TVFG.............ADGLPKAAFP......HGWKGT......RGLYVAG
ED-e-SmYUCA                        QQGDYDAIIFATGYKSNVPQWLK.GEVGNSFS.............ADGFPRC..........GWKGE......RGLYVAG
Class-1L-AtYUC11                   NTKNVDSIVFATGYKSSVSKWLEVDD.GDLFN.............ENGMPKREFP......DHWKGK......NGLYSAG
Class-1Q-AtYUC10                   HKQDFDAIVFATGYKSSVCNWLE.DY.EYVMK.............KDGFPKAPMP......KHWKGE......KNLYCAG
Class-3e-AtYUC2                    RVDNFDAIILATGYKSNVPMWLK.GV..NMFSE............KDGFPHKPFP......NGWKGE......SGLYAVG
Class-3e-AtYUC6A                   KTERFDAIILATGYKSNVPSWLK.EN..KMFSK............KDGFPIQEFP......EGWRGE......CGLYAVG
Class-3e-AtYUC6B                   KTERFDAIILATGYKSNVPSWLK.EN..KMFSK............KDGFPIQEFP......EGWRGE......CGLYAVG
Class_4f-AtYUC3                    RVLEIDSVILATGYRSNVPSWLK.DN..DFFS.............DDGIPKNPFP......NGWKGE......AGLYAVG
Class_4f-AtYUC5                    QRLDLDAVVLATGYRSNVPSWLQ.EN..DLFS.............KNGFPKSPFP......NAWKGK......SGLYAAG
Class_4f-AtYUC7                    RVLQIDSVILATGYRSNVPSWLK.EN..D.LG.............EIGIEKNPFP......KGWKGK......AGLYAVG
Class_4f-AtYUC8                    EQLDVDSVVLATGYRSNVPYWLQ.EN..EFFA.............KNGFPKTVAD.....NNGWKGR......TGLYAVG
Class_4f-AtYUC9                    QKLDIDAVVLATGYRSNVPSWLQ.ES..EFFS.............KNGFPKSPFP......NAWKGK......SGLYAAG
Class-5b_AtYUC1                    QEKDFDSIIFATGYKSNVPTWLQ.GG..DFFT.............DDGMPKTPFP......NGWRGG......KGLYTVG
Class-5B-AtYUC4A                   KEIEFDSIILATGYKSNVPDWLK.EN..SFFT.............KEGMPKTPFP......NGWKGE......KGLYTVG
Class-5B-AtYUC4B                   KEIEFDSIILATGYKSNVPDWLK.V..................................................LLSIY
Class-2a-AfYUCB                    QSDKFDSIILATGYRSNVKDWLI.GQ.DDHFT.............PDGFPSSQGC.....IGGWKGR......NGLYVVG

                                   340       350       360       370       380       390       400       410
AsFMO_S-OX                         IPWKIVPFPLCELQSKWIAAVLSGRISL.PTKKEMMEDVEAYYKQMEAAGIPKRYTHNIG..HNQFDYDDWLANECGYSC
AtFMO_S-OX                         LPWQVIPFPMFELQSKWVAAVLAGRVSL.PSQEEM.EDTKMFYLKLEASCIPKRYTHLMAELDSQFVYNNWLADQCDYPR
OsFMO_S-OX                         LPWKVIPFPLFELQSKWVAGVLSGRVKL.PSSEEMMEDVKAFHSKMEARGWPKRYAHNFS..DCQFEYDDWLAEQCGHPP
HvFMO_S-OX                         LPWKVIPFPLFELQSKWVAGILSGRIKL.PTKDEMMEDVKAMYSRRETRGWPKRYTHNFSG.GYQFEYDDWLAEQCGHPP
SmFMO_S-OX                         LPWKTIPFPLSELQAKWISAVLKGRASL.PPKGEMANDVVGFYRTLEAQRVPNRHTHRLD..LEMFDYAEWLAKQCGGTC
MpFMO_S-OX                         LPQKIVPFPLFQAQSKWISKLLSGKVKL.PSESTMMSDLKQFYDELEANGITTYNTHTMFK.RDQFVYCDWILEQCGAQP
PcFMO_S-OX                         LPYKVVPFPLCELQSKWIAKVLSGKLSL.PSAHQMMVAVQNFYASLKAHGLPKRFTHNAD..TYQFAYVSWLAEQCGCE.
AtFMO1_N-OX                        YVQSSSNLHTSELRSMWLSRLVDEKFRL.PSKEKMLDQFLKEMEVTRNS...SRFYKRHCISTFSIQHADDMCNDMGLNP
OsFMO1_N-OX                        YVESVSNLHTSELRCRWLAGLLGGRFAL.PAVEEMVRHVDGETEAMRRT...TRFYRRHCISTYSIHDSDAMCADLGSRV
HvFMO1_N-OX                        FVESVSNLHTSELRCRWLSGLLEGRFEL.PTVKAMMGHVAGEADAMRRT...TRFYRRHCISTYSIHDSDGMCADLGSAT
SmFMO1_N-OX                        YQEDVSNVRASEMGARWLSYYLTGKVKL.PDSQTMSKSANEWVRYMKWS...TPYFDRACVALARTWHSDELCRDIGWNP
PcFMO1_N-OX                        YQEGLSSIHSSELGARWLAHYLSGKITL.PSTQAMEKDISSWMACVVTA...SRFVSRICISLLHIAHADKLCQDMRWEP
SisterYUCCA_P.carolinianus         ................................................................................
SisterYUCCA-P.carolinianus         .....GTVQENRHFSLVLALQLKARFEGISTPVYGMQPVFHLS......................................
SisterYUCCA_Marchantia             .....GNLHQCRHYSLVVALQLKARLEGIPTPVYALPKVNHA.......................................
SisterYUCCA_Marchantia             .....GGLSHCRFYSRHVALQLKARMEGIPTPVYKLVRHKEPDSPVLASKMSS............................
SisterYUCCA_S.moellendorffii       .....GNLHQARHYSLYLALQLKARMEGIATPVYGMPKVHHLS......................................
SisterYUCCA_S.moellendorffii       .....GNLHQARHYSLYLALQLKARMEGITTPVYGMPKVHHLS......................................
ED_c_MpYUC1                        FTK..KGLVGTFTDATNIAKDIGRAYDH.DMLQTAI............................................
ED_d_PcYUCA                        MSK..RGLKGTSFEARQIAQEI..........................................................
ED-e-SmYUCA                        LSR..KGIFGGSKDAQMIAEDISKEYSL.VHKLVSTNRAKQQQQQ...................................
Class-1L-AtYUC11                   FGK..QGLAGISRDARNIARDIDSLVCG.RSSKNKLSK..........................................
Class-1Q-AtYUC10                   FSR..KGIAGGAEDAMSVADDIRSILAT....................................................
Class-3e-AtYUC2                    FTK..LGLLGAAIDAKKIAEDIEVQRHF....................................................
Class-3e-AtYUC6A                   FTK..RGISGASMDAKRIAEDIHKCWKQ.DEQVKKI............................................
Class-3e-AtYUC6B                   FTK..RGISGASMDAKRIAEDIHKCWKQ.DEQLQCKLGKRMKRKFSESD..........................CGGN.
Class_4f-AtYUC3                    FTR..KGLFGASLDAMSVAHDIANRWKE.ESKQQKKTAAARHRRCISHF...............................
Class_4f-AtYUC5                    FTR..KGLAGASADAVNIAQDIGNVWRE.ETKRQKMRTRVGHRRCISVA...............................
Class_4f-AtYUC7                    FTG..RGLSGASFDAMSVAHDIANSWKE.ETKQQIKTVATRHRRCISHF...............................
Class_4f-AtYUC8                    FTR..KGLSGASMDAVKIAQDIGSVWQL.ETKQPTKRSRGSLRRCISQQF..............................
Class_4f-AtYUC9                    FTR..KGLAGASVDAVNIAQDIGNVWRE.ETKRQKMRRNVGHRRCISVA...............................
Class-5b_AtYUC1                    FTR..RGLLGTASDAVKIAGEIGDQWRD.EIKGSTRNMCSSRFVFTSKS...............................
Class-5B-AtYUC4A                   FTR..RGLSGTAYDAVKIAEDITDQWMKFNGPLSCRNICSSHIIHLHFNK.............................S
Class-5B-AtYUC4B                   YLS..YSLSCYIYVNVYM.......YM.....................................................
Class-2a-AfYUCB                    MSG..KGLAGTKSDAQLIAKDIQLEF......................................................

                                       420       430          440       450
AsFMO_S-OX                         ..IEEWRRLMYKEVSKNRKERPESYR...DEWDDDHLVAQARETFSKFLS.............
AtFMO_S-OX                         ..IEKWREQMFYKVFKRIQSQASTYK...DDWDDDHLIAEAYEDFVKFPSNYPSSLIEREYTS
OsFMO_S-OX                         ..IEQWRKLMYAANSENKAARPESYR...DEWDDDHLVAEAAEDFKKYL................
HvFMO_S-OX                         ..IEEWRKLMYAANAKNKAARPERYR...DEWDDDDLVAQASEDFKKYL................
SmFMO_S-OX                         HGFEPWRKELFLSTRDNRKLNSDSYR...DEWSDNDLHEKVVGVLA.......TSFFMREQ..
MpFMO_S-OX                         ..LDSWREAIYFSAHANKGT.STNYR...DDWKDQDLLDTATKSLVELSER..LSLQVK....
PcFMO_S-OX                         ..VDWWREEMYTAAE..RAKEPATYR...DDWNDQQLM.........................
AtFMO1_N-OX                        WRKSNFLLEAFSPYG.....SQDYRLGQEEKED...........................MTA
OsFMO1_N-OX                        HRKSNFLAELFAPYN.....NQDYK...EE.................................
HvFMO1_N-OX                        LRKANWIAELFAPYN.....NKDYK...EQ.................................
SmFMO1_N-OX                        RRKKNWLQELLSPYS.....NMDYI......D...............................
PcFMO1_N-OX                        RRKRSFLADLFKPYT.....NADY.......................................
SisterYUCCA_P.carolinianus         ...............................................................
SisterYUCCA-P.carolinianus         ...............................................................
SisterYUCCA_Marchantia             ...............................................................
SisterYUCCA_Marchantia             .............................IQMQD.............................
SisterYUCCA_S.moellendorffii       ...............................................................
SisterYUCCA_S.moellendorffii       ...............................................................
ED_c_MpYUC1                        ...............................................................
ED_d_PcYUCA                        ...............................................................
ED-e-SmYUCA                        ...............................................................
Class-1L-AtYUC11                   ...............................................................
Class-1Q-AtYUC10                   ...............................................LKNN............
Class-3e-AtYUC2                    .....................................................LPL.ARPQHC
Class-3e-AtYUC6A                   ...............................................................
Class-3e-AtYUC6B                   ...............................................................
Class_4f-AtYUC3                    ...............................................................
Class_4f-AtYUC5                    ...............................................................
Class_4f-AtYUC7                    ...............................................................
Class_4f-AtYUC8                    ...............................................................
Class_4f-AtYUC9                    ...............................................................
Class-5b_AtYUC1                    ...............................................................
Class-5B-AtYUC4A                   ...............................................................
Class-5B-AtYUC4B                   ...............................................................
Class-2a-AfYUCB                    ...............................................................
```

#### 2. Sequence alignment for YUCCAs (Class based)

##### Early Diverging (ED) -Class YUCCAs

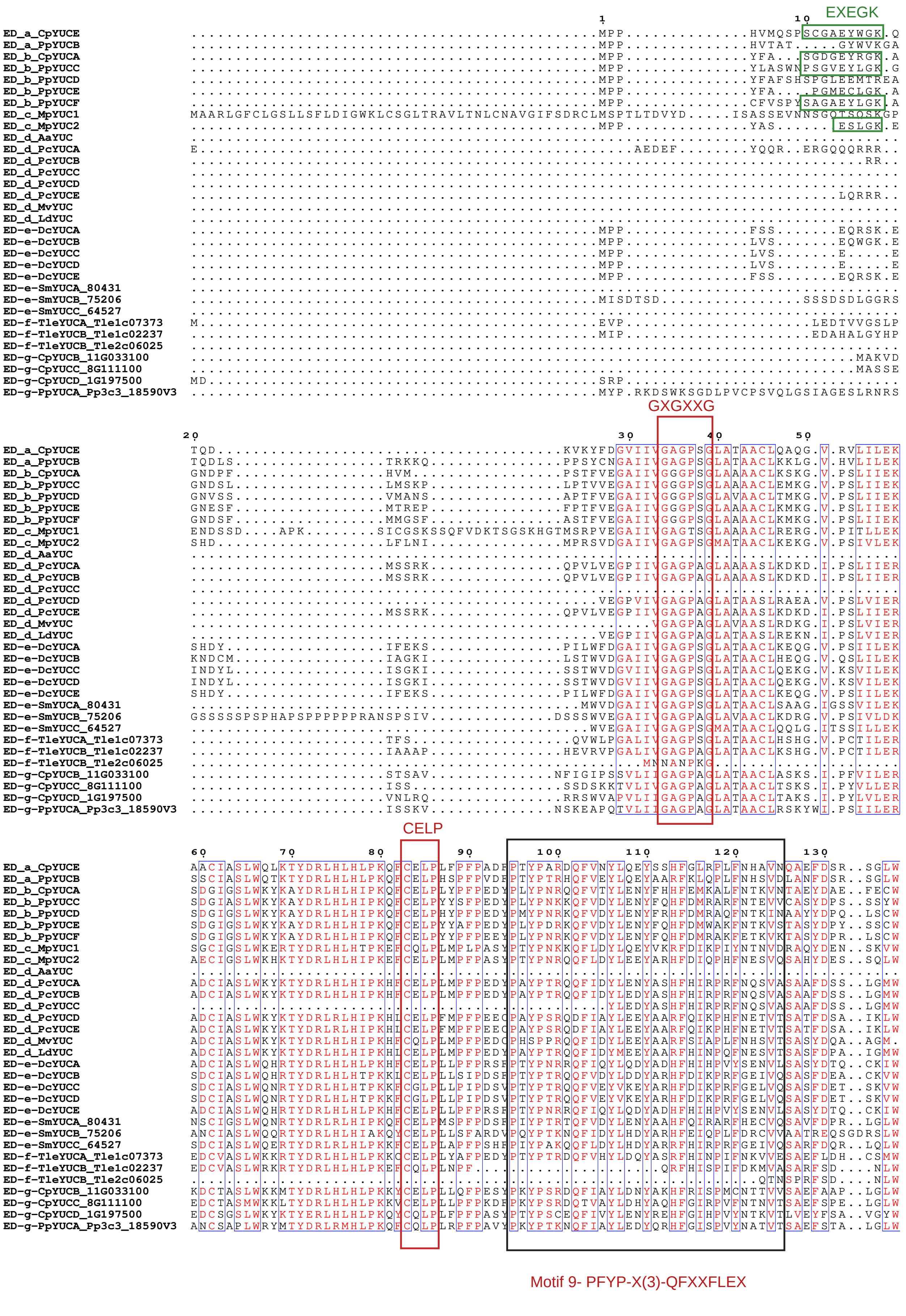

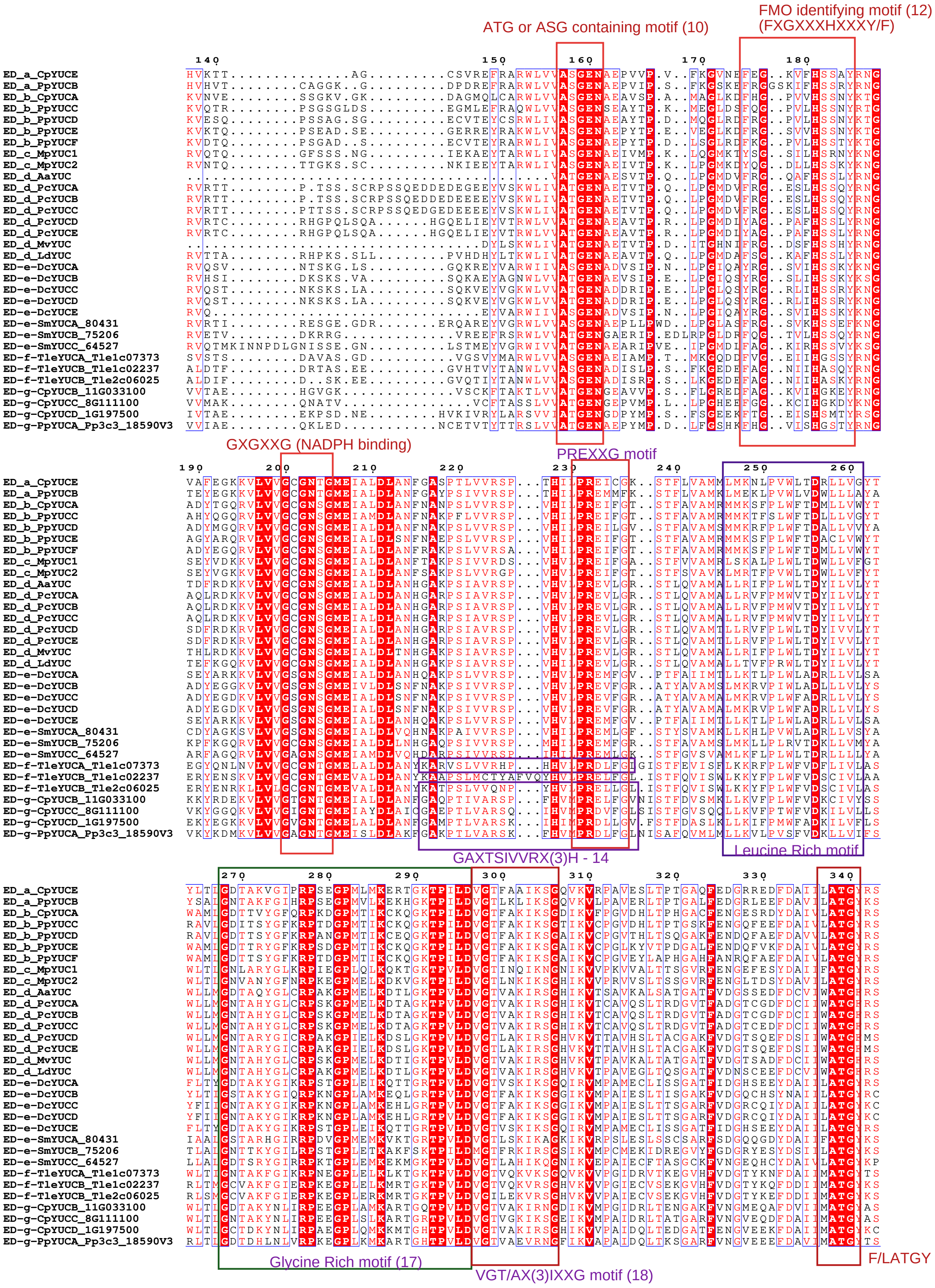
ATG or ASG containing motif (10)
FMO identifying motif (12)
(FXGXXXHXXXY/F)
GXGXXG (NADPH binding)
PREXXG motif
GAXTSIVVRX(3)H - 14
Leucine Rich motif
Glycine Rich motif (17)
VGT/AX(3)IXXG motif (18)
F/LATGY

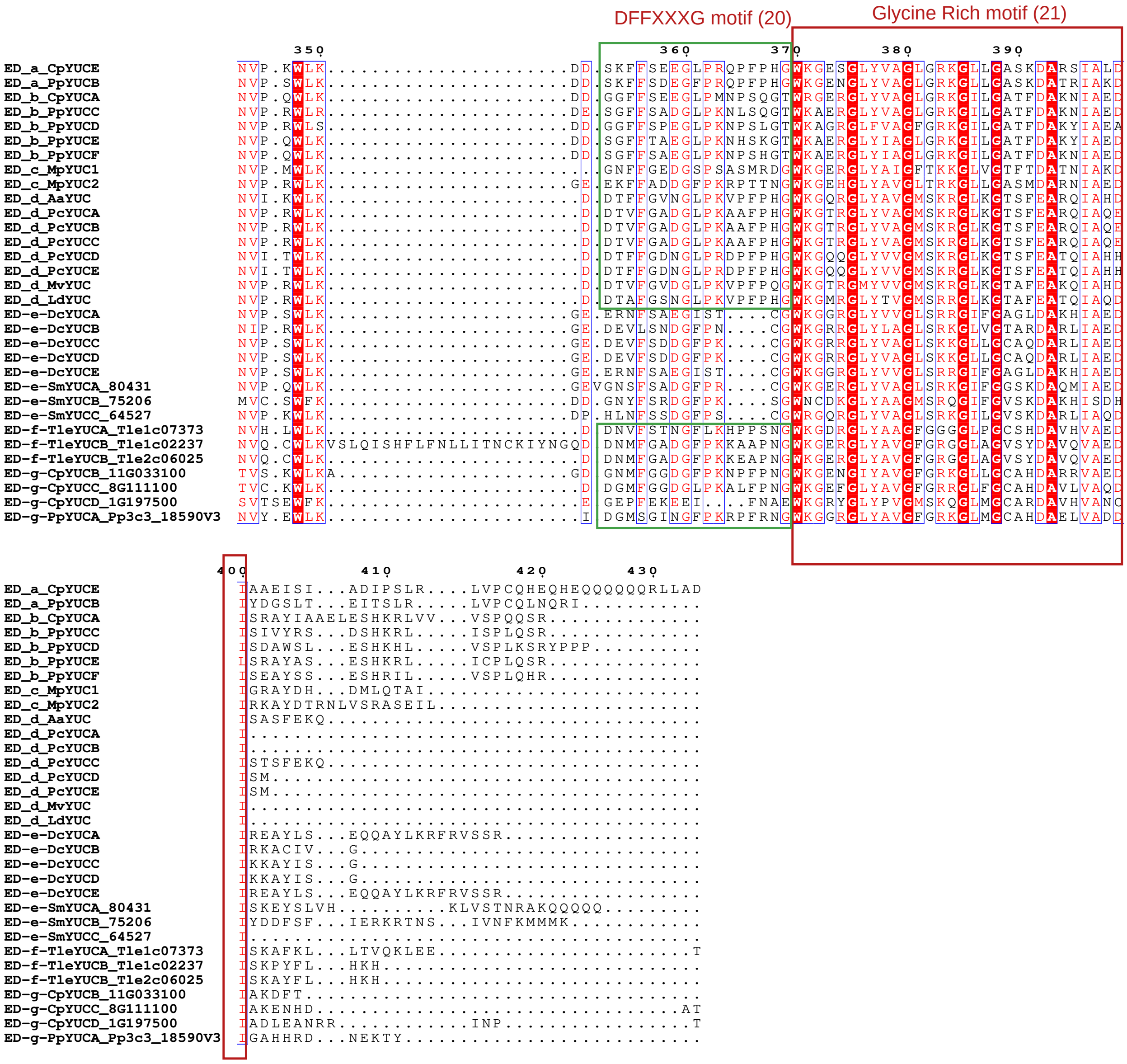
DFFXXXG motif (20)
Glycine Rich motif (21)
ED_a_CpYUCE
ED_a_PpYUCB
ED_b_CpYUCA
ED_b_PpYUCC
ED_b_PpYUCD
ED_b_PpYUCE
ED_b_PpYUCF
ED_c_MpYUC1
ED_c_MpYUC2
ED_d_AaYUC
ED_d_PcYUCA
ED_d_PcYUCB
ED_d_PcYUCC
ED_d_PcYUCD
ED_d_PcYUCE
ED_d_MvYUC
ED_d_LdYUC
ED-e-DcYUCA
ED-e-DcYUCB
ED-e-DcYUCC
ED-e-DcYUCD
ED-e-DcYUCE
ED-e-SmYUCA_80431
ED-e-SmYUCB_75206
ED-e-SmYUCC_64527
ED-f-TleYUCA_Tle1c07373
ED-f-TleYUCB_Tle1c02237
ED-f-TleYUCB_Tle2c06025
ED-g-CpYUCB_11G033100
ED-g-CpYUCC_8G111100
ED-g-CpYUCD_1G197500
ED-g-PpYUCA_Pp3c3_18590V3

### Class-1 YUCCAs

```
                                1
Class1a_Aspi01Gene11003         M...................MMMMSP......................................................
Class1a_Aspi01Gene36448         ....................MMGMAH......................................................
Class1a_CFernYUCD               .......................MEL......................................................
Class1b_Aspi01Gene19346         M....................CVISE......................................................
Class1b_Aspi01Gene11463         .......................MSE......................................................
Class1b_Aspi01Gene53362         M....................CKTSD......................................................
Class1b_CFernYUCC               M....................CRMPE......................................................
Class1c_AfYUCE                  .......................M........................................................
Class1c_ScYUCC                  .......................MDGWKGQGQGQGRKGREGKGKKGRRDRDRDREEGEGAETGTGEEGGEGAGAGAGEEG
Class1d_CFernYUCF               .......................MLN......................................................
Class1d_Aspi01Gene38026         .......................MSP......................................................
Class1d_Aspi01Gene72802         .......................MPP......................................................
Class1e_AfYUCC                  .......................MPLFSNRKQQ...............................................
Class1e_ScYUCD                  .......................MTLFS....................................................
Class1f_Aspi01fene30495         .......................MRP.................................A....................
Class1f_Aspi01fene32088         .......................MP.......................................................
Class1f_AcYUC                   ................................................................................
Class1f_AfYUCA                  .....................MKLPPLS....................................................
Class1f_AfYUCD                  .......................M........................................................
Class1f_ScYUCA                  ................................................................................
Class1f_ScYUCB                  .......................MAA......................................................
Class1f_CFernYUCA               .......................MPASMM...................................................
Class1f_CFernYUCE               .......................MPGVSM...................................................
Class1g_PabYUCA                 .......................MAN......................................................
Class1g_GbYUCD_Gb_15900.p       M....................LKMER......................................................
Class1g_PlYUC_PILAhq_070532     .......................MAN......................................................
Class1g_PlYUC_PILAhq_052127     .......................MAN......................................................
Class1g_PtaYUC_PITA_26624.1     .......................MAN......................................................
Class1g_PtaYUC_PITA_02185.1     .......................MPN......................................................
Class1g_PtaYUC_PITA_00074.1     .......................MPN......................................................
Class1g_PtaYUC_PITA_27422.1     .......................MPN......................................................
UC_PITA_18009.1.p_Pinus_taeda   .......................MPN......................................................
Class1g_PtaYUC_PITA_26118.1     .......................MPN......................................................
Class1g_TpYUCC                  .......................M........................................................
Class1h_PabYUC2                 ................................................................................
Class1h_PlYUC_PILAhq_070572     .......................MPQ......................................................
Class1h_PlYUC_PILAhq_041125     .......................MPP......................................................
Class1h_PtaYUC_PITA_32408       .......................MPS......................................................
Class1h_TpYUCA                  ................................................................................
lass1i_GmYUCE_TnS000915671t02   .......................MAE......................................................
Class1j_HvYUC11_XP_044958834    ........................M.......................................................
1j_SbYUC10_Sobic.003G128700.1   ................................................................................
lass1j_SbYUC10_XP_002455445.1   .......................MAR......................................................
Class1j_ZmYUC11                 ................................................................................
Class1j_BdYUC11                 .......................M........................................................
Class1j_AcYUC10                 .......................MH.......................................................
Class1j_OsYUC9_LOCOs01g16714    .......................M........................................................
Class1j_OsYUC10_LOCOs01g16750   .......................M........................................................
Class1j_MaYUC11_Achr9P17710     .......................M........................................................
lass1j_MaYUC10/11_Achr8P00130   ................................................................................
Class1j_EcYUC10A                ................................................................................
Class1j_EcYUC10B                .......................M........................................................
Class1k_AmTrYUC10               ................................................................................
Class1k_AmTrYUC11A              ................................................................................
Class1k_AmTrYUC11B              ................................................................................
Class1k_NcYUC11                 ................................................................................
Class1k_NcYUC10                 ................................................................................
Class1l_AtYUC11                 ................................................................................
Class1l_PsYUC4_QTZ21217.1       .......................M........................................................
Class1l_GmYUC4                  .......................MNS......................................................
Class1l_GmYUC10                 ................................................................................
Class1l_CsYUC11                 ................................................................................
Class1l_MtYUC10                 ................................................................................
Class1l_MtYUC11                 ................................................................................
Class1l_MtYUC10_Medtr3g088925   ................................................................................
Class1l_MtYUC10_Medtr3g088945   ................................................................................
Class1l_PpYUC10_Prupe8G014100   ................................................................................
Class1l_BrYUC11                 ................................................................................
Class1l_PtYUC9_                 ................................................................................
Class1l_PtYUC11                 ................................................................................
Class1l_PtYUC12                 ................................................................................
Class1l_CaYUC11                 ................................................................................
Class1l_CpYUC11                 .......................M........................................................
Class1l_FvYUC1                  ................................................................................
Class1l_VvYUC10                 ................................................................................
Class1l_VvYUC11                 ................................................................................
Class1m_PsYUC6                  ................................................................................
Class1m_PsYUC8                  ................................................................................
Class1m_CKAN_02337900           ................................................................................
Class1m_PtYUC10                 ................................................................................
Class1m_SlyYUC                  .VISL.......NTNKFSFSILVMATF.....................................................
Class1m_StYUC10                 MYIYIPHLFFFSQHHQIFFSILVMTTF.....................................................
Class1m_StYUC10                 MYIDIPHLFFFSQHHQIFFSILVMTTF.....................................................
Class1m_HqYUC10                 ................................................................................
Class1m_CaYUC10A                ................................................................................
Class1m_CaYUC10B                ................................................................................
Class1m_CaYUC10C                ................................................................................
Class1m_GmYUC2                  ................................................................................
Class1m_GmYUC21                 ................................................................................
Class1m_CpYUC10                 ................................................................................
Class1m_CsYUC10_Cucsa108650.1   ................................................................................
ass1m_CsYUC10X1_Cucsa084650.1   ................................................................................
Class1m_FvYUC10_AFG16920.1      ................................................................................
ass1m_MtYUC10_Medtr7g107710.1   ................................................................................
ass1m_MtYUC10_Medtr7g107690.1   ................................................................................
s1m_VvYUC10_VIT207s0104g01260   ................................................................................
s1m_VvYUC10_VIT207s0104g01250   ................................................................................
ass1m_PpYUC10_Prupe8G252500.1   ................................................................................
Class1m_PCaYUCE_KAI3988924      ................................................................................
Class1m_DcYUC10A                ................................................................................
Class1m_DcYUC10B                ................................................................................
Class1n_CKAN_00715200           ................................................................................
Class1n_MsiYUC10_XP_058099706   ................................................................................
Class1n_AoYUC10_09.284          ................................................................................
Class1n_AoYUC10B_09.282         ................................................................................
ass1n_AaYUC10_Acora.05G103700   ................................................................................
Class1o_HvYUC10_XP_044959882    ................................................................................
Class1o_SbYUC11_XP_021302284    ................................................................................
lass1o_OsYUC11_LOC_Os12g08780   ................................................................................
lass1o_ZmYUC1_ZmB84.10G037700   .......................M........................................................
Class1o_BdYUC10A_Bradi4g40750   ................................................................................
Class1o_BdYUC_Bradi4g40770      ................................................................................
Class1o_OtYUC11_Oropetium       ................................................................................
Class1o_EcYUC11                 .......................MSSSR....................................................
Class1o_HvYUC10_XP044968451     ................................................................................
Class1o_HvYUC10_XP_044960962    ................................................................................
Class1o_HvYUC10_XP_044957826    ................................................................................
ss1o_HvYUC10_HORVU2Hr1G118760   ................................................................................
ss1o_HvYUC10_HORVU1Hr1G022530   ................................................................................
Class1o_HORVU1Hr1G067720        ................................................................................
lass1o_HvYUC_HORVU7Hr1G017630   ................................................................................
Class1o_SbYUC10_XP_002449399    ................................................................................
lass1o_OsYUC12_LOC_Os02g17230   ................................................................................
lass1o_OsYUC12_LOC_Os11g10140   .......................M........................................................
lass1o_OsYUC14_LOC_Os11g10170   .......................M........................................................
1o_ZmYUC3_ZmB84.04G035900.1.p   ................................................................................
lass1o_BdYUC_Bradi2g04257.1.p   ................................................................................
s1o_BdYUC10B_Bradi2g04247.1.p   ................................................................................
ass1p_SlyYUC10_Solyc09g091720   ................................................................................
Class1p_SlyYUC_Solyc09g091870   .....................KNL........................................................
s1p_HqYUC10_Hyque.05G023500.1   ................................................................................
Class1q_AtYUC10_AT1G48910       ................................................................................
Class1q_CsYUC10_Cucsa.249910    ................................................................................
Class1q_FvYUC11_AFG16919.1      ................................................................................
Class1q_BrYUC10_Brara.F00397    ................................................................................
Class1q_PsYUC7_QTZ21220         ................................................................................
lass1q_GmYUC3_Glyma.04G213600   ................................................................................
ass1q_GmYUC11_Glyma.06G152700   ................................................................................
Class1q_MtYUC10_Medtr5g033260   ................................................................................
Class1q_MtYUC10_Medtr8g432640   ................................................................................
```

```
Class1a_Aspi01Gene11003          ................................................................................
Class1a_Aspi01Gene36448          ................................................................................
Class1a_CFernYUCD                .......................................................MRIVLSDI.ISRDLTQMKAAPNNQR
Class1b_Aspi01Gene19346          ..................................................P.....RALI.............APPVRAR
Class1b_Aspi01Gene11463          ..................................................S.....RALL.............APHVRPR
Class1b_Aspi01Gene53362          ..................................................P.....RVPL.............APHACAR
Class1b_CFernYUCC                ..................................................P.....R..N............TATRSQPR
Class1c_AfYUCE                   .......................................................TNVRLGE..................
Class1c_ScYUCC                   EGKGRKGAVILPPSRGPFYAPRTCLLFIHSCFRPRTSFTIPPLTSIPSPFPSLPLFALHFNYVLYMRCAAMPVDPWLPAR
Class1d_CFernYUCF                ........................................................RNRLNHC.......CD........
Class1d_Aspi01Gene38026          ...........................................................Y....................
Class1d_Aspi01Gene72802          ...........................................................Y....................
Class1e_AfYUCC                   ...........................THA..............ELPNLF.........FDEE.EE.....EEEEKNHPR
Class1e_ScYUCD                   ........................................................AVVFSGR.................
Class1f_Aspi01fene30495          ...........................................................F....................
Class1f_Aspi01fene32088          ................................................................................
Class1f_AcYUC                    ................................................................................
Class1f_AfYUCA                   ...........................................................F....................
Class1f_AfYUCD                   ................................................................................
Class1f_ScYUCA                   ................................................................................
Class1f_ScYUCB                   .........................................................................ETGALAR
Class1f_CFernYUCA                .........................................................................ESKYVPR
Class1f_CFernYUCE                ................................................................................
Class1g_PabYUCA                  ...........................................................F....................
Class1g_GbYUCD_Gb_15900.p        ...........................................................F....................
Class1g_PlYUC_PILAhq_070532      ...........................................................F....................
Class1g_PlYUC_PILAhq_052127      ...........................................................F....................
Class1g_PtaYUC_PITA_26624.1      ...........................................................F....................
Class1g_PtaYUC_PITA_02185.1      ...........................................................F....................
Class1g_PtaYUC_PITA_00074.1      ...........................................................F....................
Class1g_PtaYUC_PITA_27422.1      ...........................................................F....................
UC_PITA_18009.1.p_Pinus_taeda    ...........................................................F....................
Class1g_PtaYUC_PITA_26118.1      ...........................................................F....................
Class1g_TpYUCC                   ................................................................................
Class1h_PabYUC2                  ................................................................................
Class1h_PlYUC_PILAhq_070572      ...........................................................F....................
Class1h_PlYUC_PILAhq_041125      ...........................................................L....................
Class1h_PtaYUC_PITA_32408        ...........................................................F....................
Class1h_TpYUCA                   ................................................................................
lass1i_GmYUCE_TnS000915671t02    ................................................................................
Class1j_HvYUC11_XP_044958834     ................................................................................
1j_SbYUC10_Sobic.003G128700.1    ................................................................................
lass1j_SbYUC10_XP_002455445.1    ................................................................................
Class1j_ZmYUC11                  ................................................................................
Class1j_BdYUC11                  ................................................................................
Class1j_AcYUC10                  ................................................................................
Class1j_OsYUC9_LOCOs01g16714     ................................................................................
Class1j_OsYUC10_LOCOs01g16750    ................................................................................
Class1j_MaYUC11_Achr9P17710      ................................................................................
lass1j_MaYUC10/11_Achr8P00130    ................................................................................
Class1j_EcYUC10A                 ................................................................................
Class1j_EcYUC10B                 ................................................................................
Class1k_AmTrYUC10                ................................................................................
Class1k_AmTrYUC11A               ................................................................................
Class1k_AmTrYUC11B               ................................................................................
Class1k_NcYUC11                  ................................................................................
Class1k_NcYUC10                  ................................................................................
Class1l_AtYUC11                  ................................................................................
Class1l_PsYUC4_QTZ21217.1        ................................................................................
Class1l_GmYUC4                   ................................................................................
Class1l_GmYUC10                  ................................................................................
Class1l_CsYUC11                  ................................................................................
Class1l_MtYUC10                  ................................................................................
Class1l_MtYUC11                  ................................................................................
Class1l_MtYUC10_Medtr3g088925    ................................................................................
Class1l_MtYUC10_Medtr3g088945    ................................................................................
Class1l_PpYUC10_Prupe8G014100    ................................................................................
Class1l_BrYUC11                  ................................................................................
Class1l_PtYUC9_                  ................................................................................
Class1l_PtYUC11                  ................................................................................
Class1l_PtYUC12                  ................................................................................
Class1l_CaYUC11                  ................................................................................
Class1l_CpYUC11                  ................................................................................
Class1l_FvYUC1                   ................................................................................
Class1l_VvYUC10                  ................................................................................
Class1l_VvYUC11                  ................................................................................
Class1m_PsYUC6                   ................................................................................
Class1m_PsYUC8                   ................................................................................
Class1m_CKAN_02337900            ................................................................................
Class1m_PtYUC10                  ................................................................................
Class1m_SlyYUC                   ................................................................................
Class1m_StYUC10                  ................................................................................
Class1m_StYUC10                  ................................................................................
Class1m_HqYUC10                  ................................................................................
Class1m_CaYUC10A                 ................................................................................
Class1m_CaYUC10B                 ................................................................................
Class1m_CaYUC10C                 ................................................................................
Class1m_GmYUC2                   ................................................................................
Class1m_GmYUC21                  ................................................................................
Class1m_CpYUC10                  ................................................................................
Class1m_CsYUC10_Cucsa108650.1    ................................................................................
ass1m_CsYUC10X1_Cucsa084650.1    ................................................................................
Class1m_FvYUC10_AFG16920.1       ................................................................................
ass1m_MtYUC10_Medtr7g107710.1    ................................................................................
ass1m_MtYUC10_Medtr7g107690.1    ................................................................................
s1m_VvYUC10_VIT207s0104g01260    ................................................................................
s1m_VvYUC10_VIT207s0104g01250    ................................................................................
ass1m_PpYUC10_Prupe8G252500.1    ................................................................................
Class1m_PCaYUCE_KAI3988924       ................................................................................
Class1m_DcYUC10A                 ................................................................................
Class1m_DcYUC10B                 ................................................................................
Class1n_CKAN_00715200            ................................................................................
Class1n_MsiYUC10_XP_058099706    ................................................................................
Class1n_AoYUC10_09.284           ................................................................................
Class1n_AoYUC10B_09.282          ................................................................................
ass1n_AaYUC10_Acora.05G103700    ................................................................................
Class1o_HvYUC10_XP_044959882     ................................................................................
Class1o_SbYUC11_XP_021302284     ................................................................................
lass1o_OsYUC11_LOC_Os12g08780    ................................................................................
lass1o_ZmYUC1_ZmB84.10G037700    ................................................................................
Class1o_BdYUC10A_Bradi4g40750    ................................................................................
Class1o_BdYUC_Bradi4g40770       ................................................................................
Class1o_OtYUC11_Oropetium        ................................................................................
Class1o_EcYUC11                  ................................................................................
Class1o_HvYUC10_XP044968451      ................................................................................
Class1o_HvYUC10_XP_044960962     ................................................................................
Class1o_HvYUC10_XP_044957826     ................................................................................
ss1o_HvYUC10_HORVU2Hr1G118760    ................................................................................
ss1o_HvYUC10_HORVU1Hr1G022530    ................................................................................
Class1o_HORVU1Hr1G067720         ................................................................................
lass1o_HvYUC_HORVU7Hr1G017630    ................................................................................
Class1o_SbYUC10_XP_002449399     ................................................................................
lass1o_OsYUC12_LOC_Os02g17230    ................................................................................
lass1o_OsYUC12_LOC_Os11g10140    ................................................................................
lass1o_OsYUC14_LOC_Os11g10170    ................................................................................
1o_ZmYUC3_ZmB84.04G035900.1.p    ................................................................................
lass1o_BdYUC_Bradi2g04257.1.p    ................................................................................
s1o_BdYUC10B_Bradi2g04247.1.p    ................................................................................
ass1p_SlyYUC10_Solyc09g091720    ................................................................................
Class1p_SlyYUC_Solyc09g091870    ................................................................................
s1p_HqYUC10_Hyque.05G023500.1    ................................................................................
Class1q_AtYUC10_AT1G48910        ................................................................................
Class1q_CsYUC10_Cucsa.249910     ................................................................................
Class1q_FvYUC11_AFG16919.1       ................................................................................
Class1q_BrYUC10_Brara.F00397     ................................................................................
Class1q_PsYUC7_QTZ21220          ................................................................................
lass1q_GmYUC3_Glyma.04G213600    ................................................................................
ass1q_GmYUC11_Glyma.06G152700    ................................................................................
Class1q_MtYUC10_Medtr5g033260    ................................................................................
Class1q_MtYUC10_Medtr8g432640    ................................................................................
```

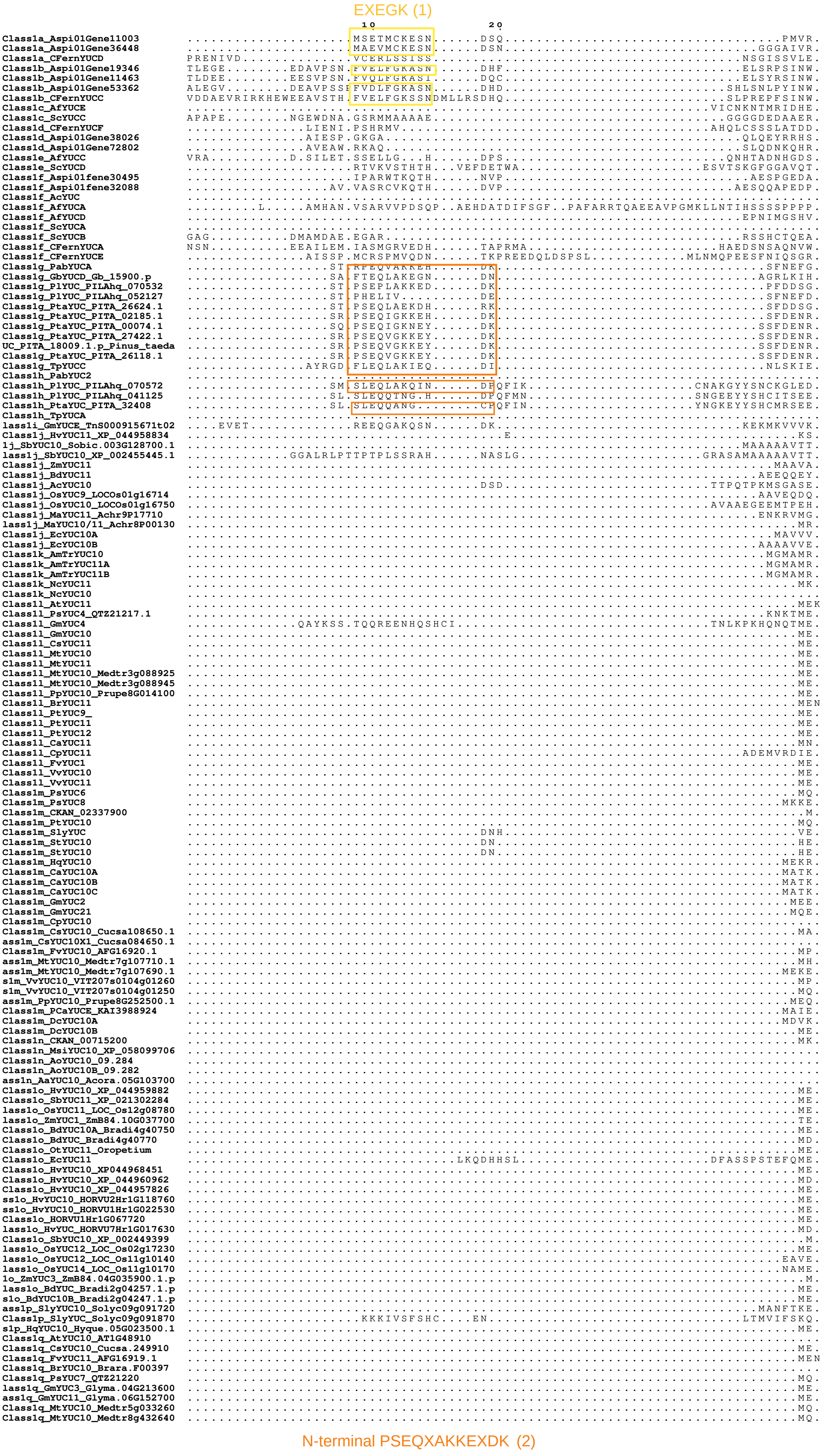


N-terminal PSEQXAKKEXDK (orange highlighted) found only in some species of gymnosperm species.
Could not detect this type in any other species studied

#### Class-1 YUCCAs

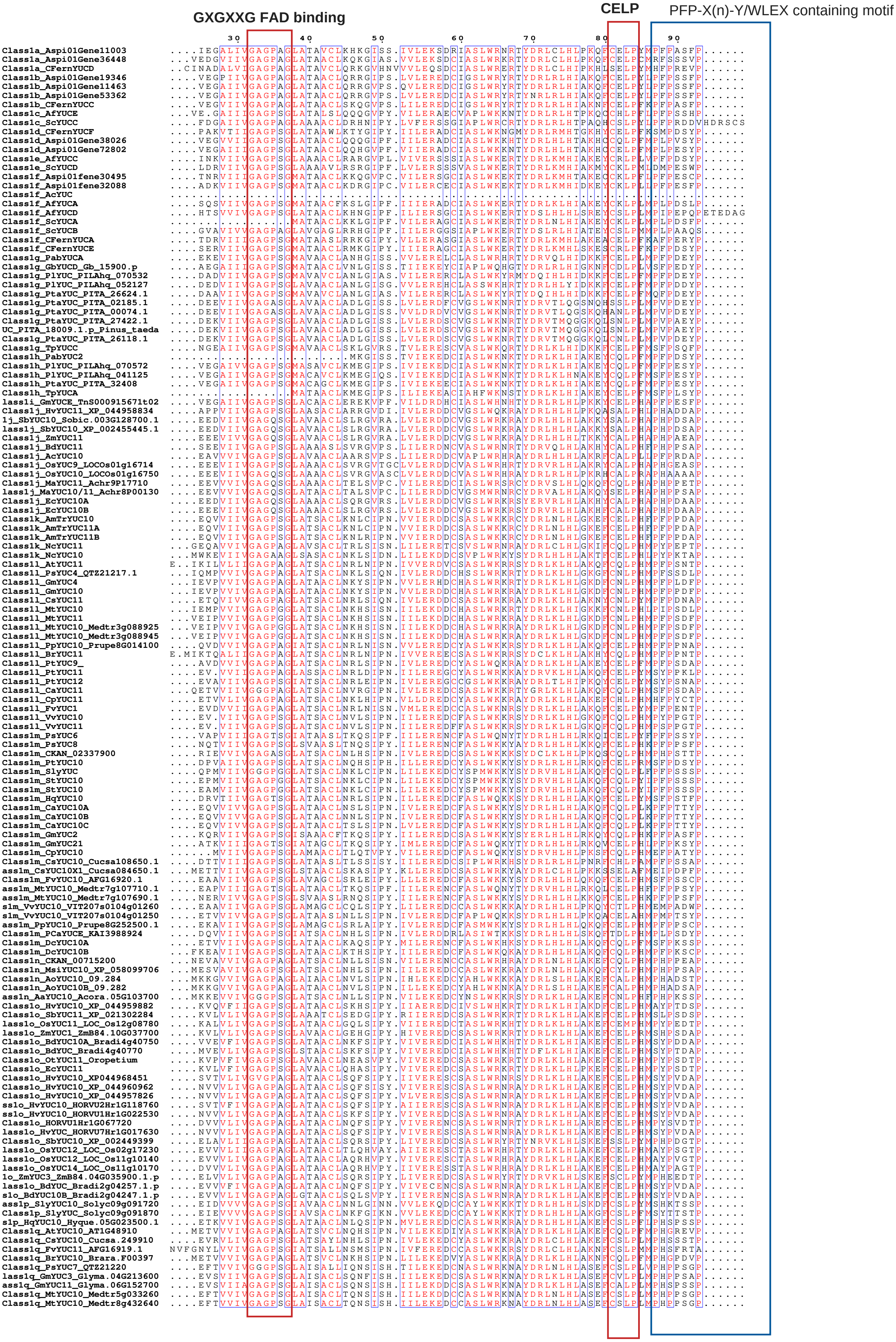

PFP-X(n)-Y/WLEX containing motif

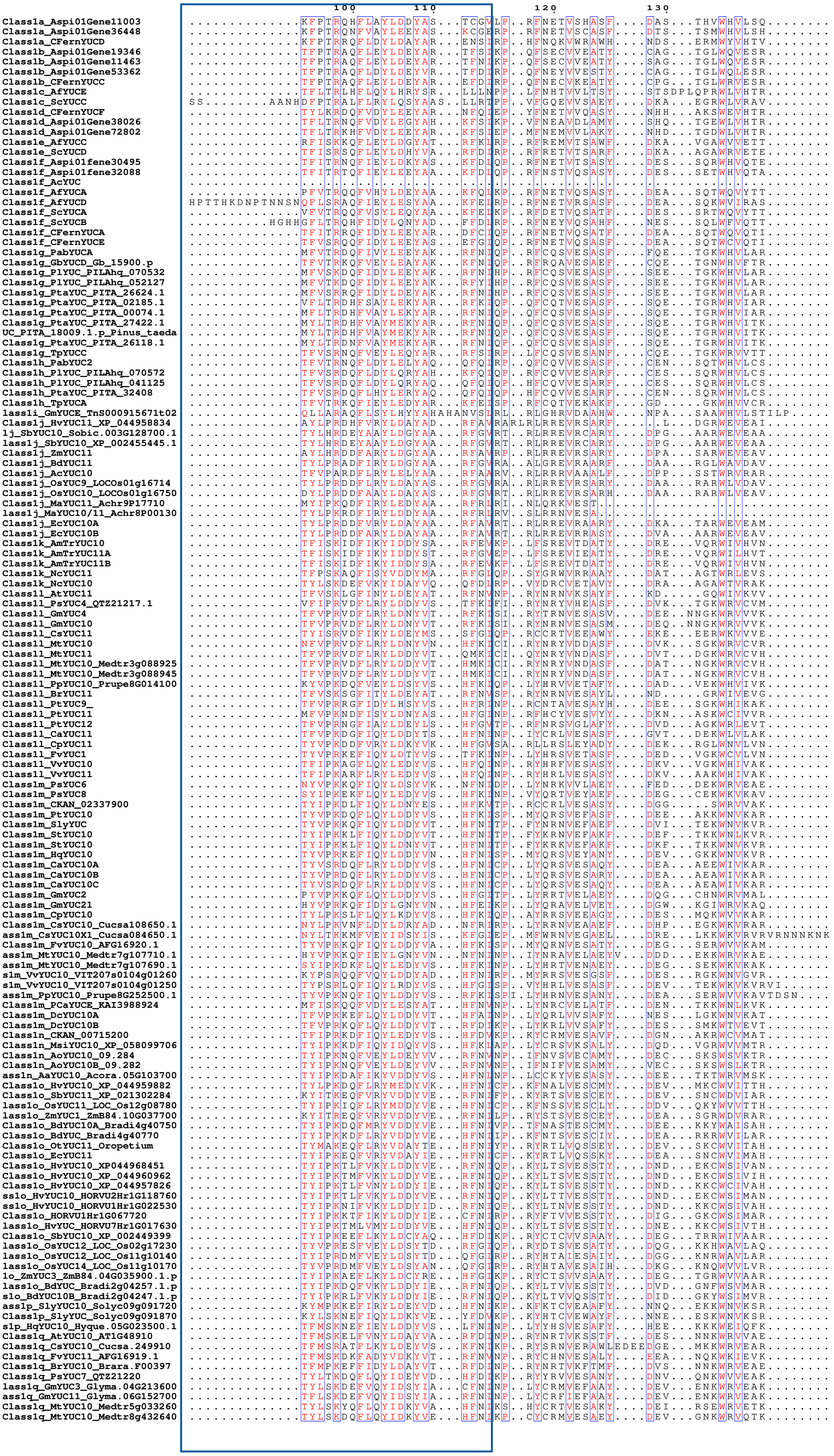

#### Class-1 YUCCAs

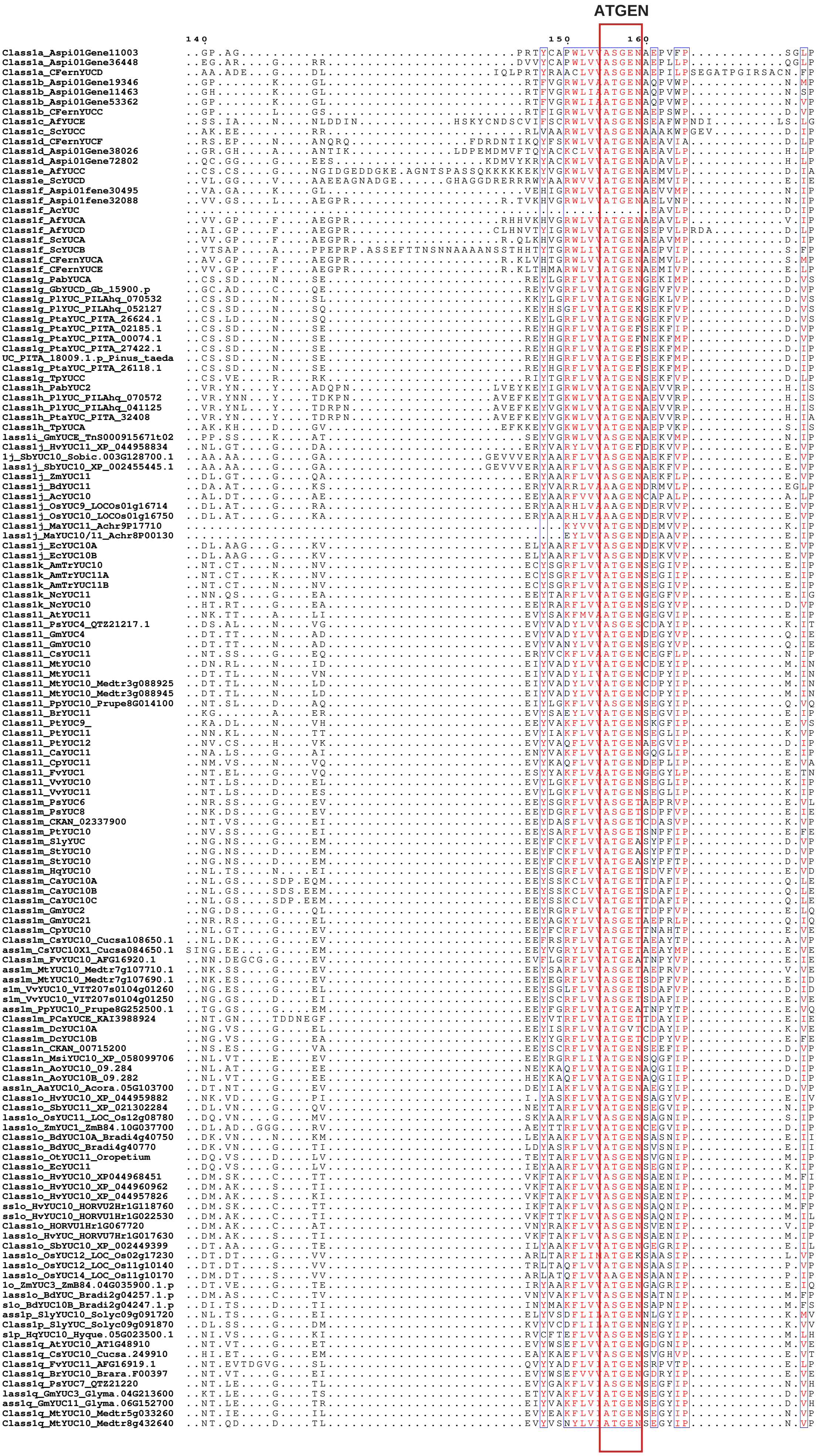

Class-1 YUCCAs

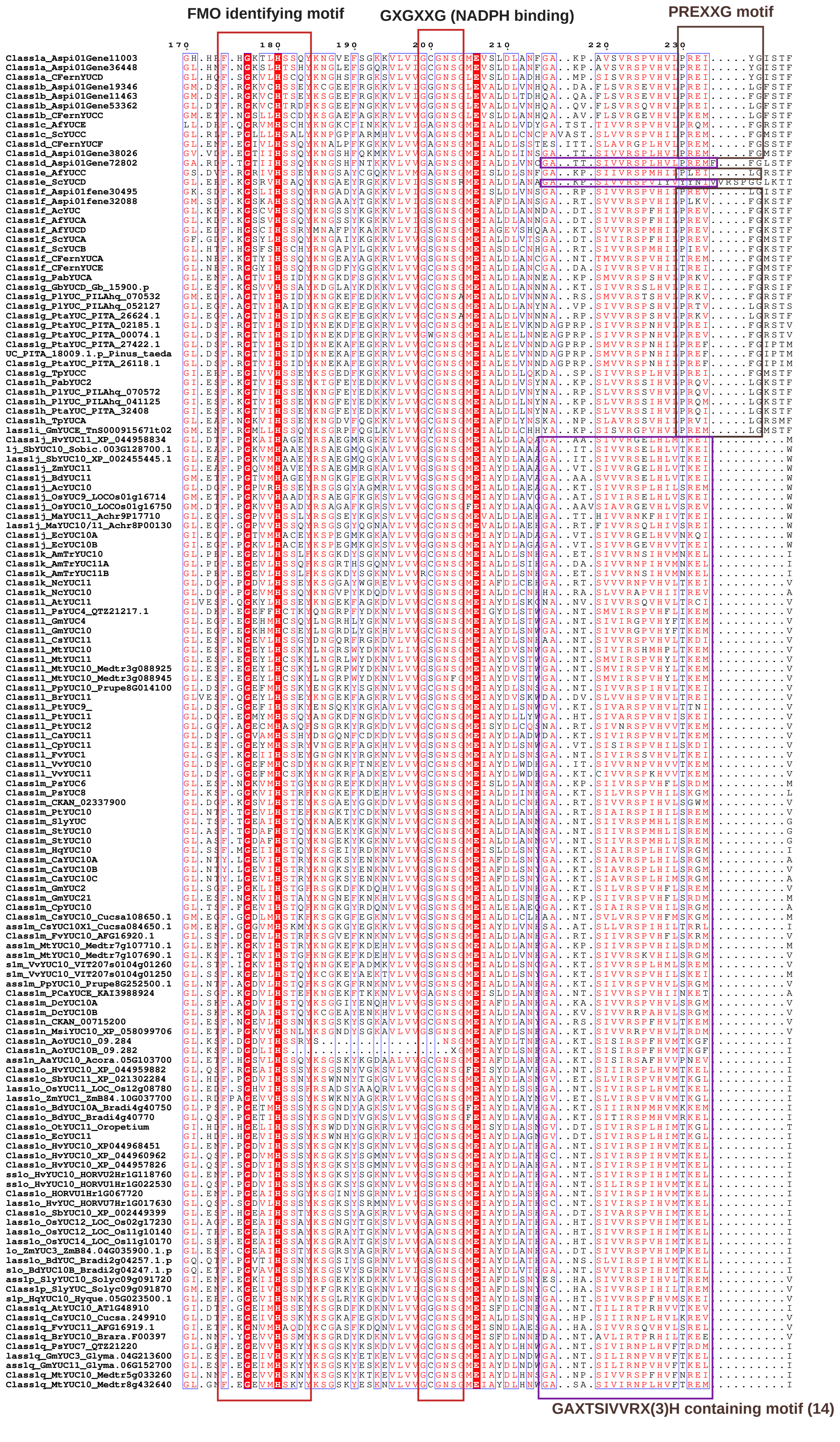

### Class-1 YUCCAs

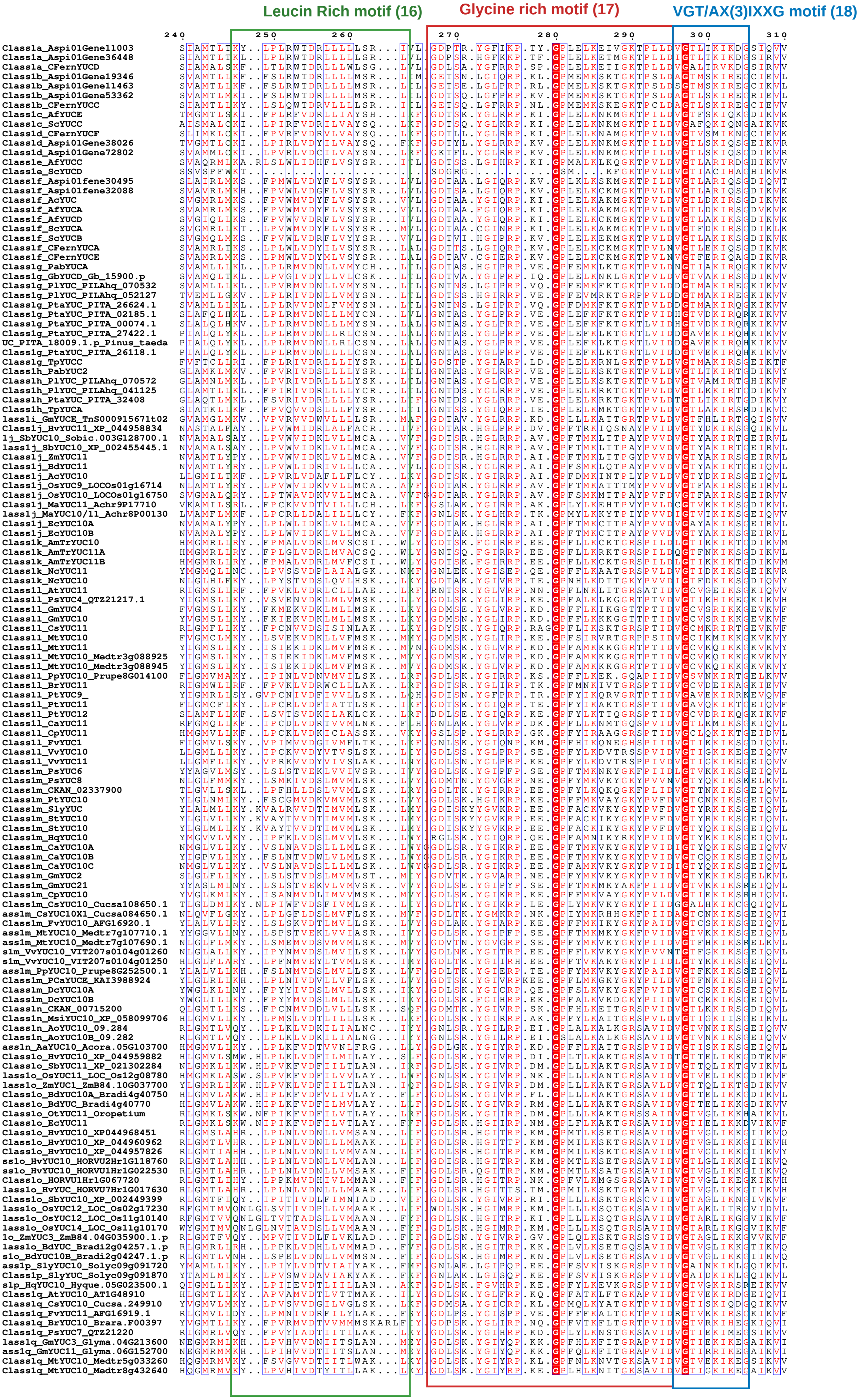

**Class-1 YUCCAs**

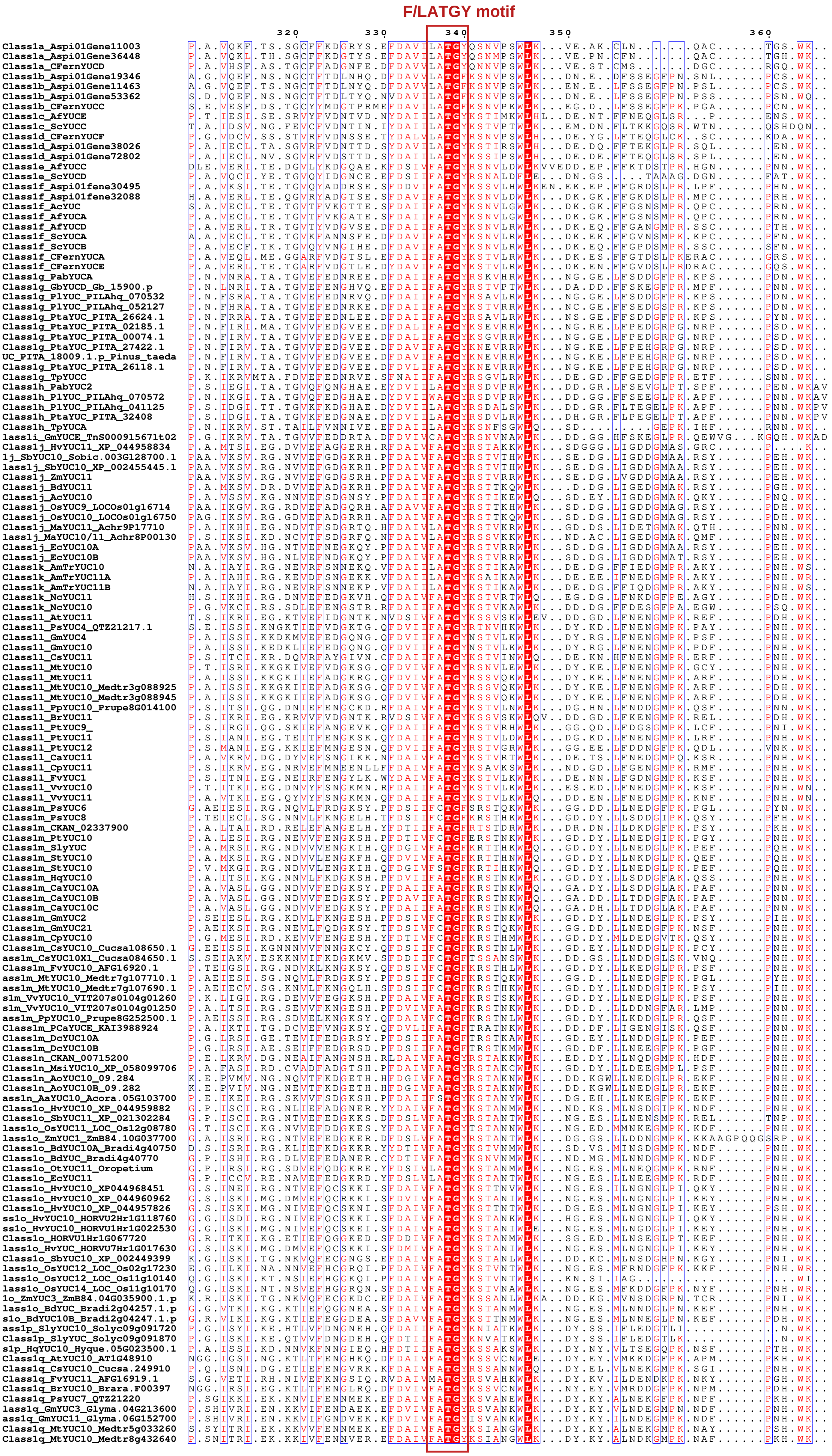

#### Class-1 YUCCAs

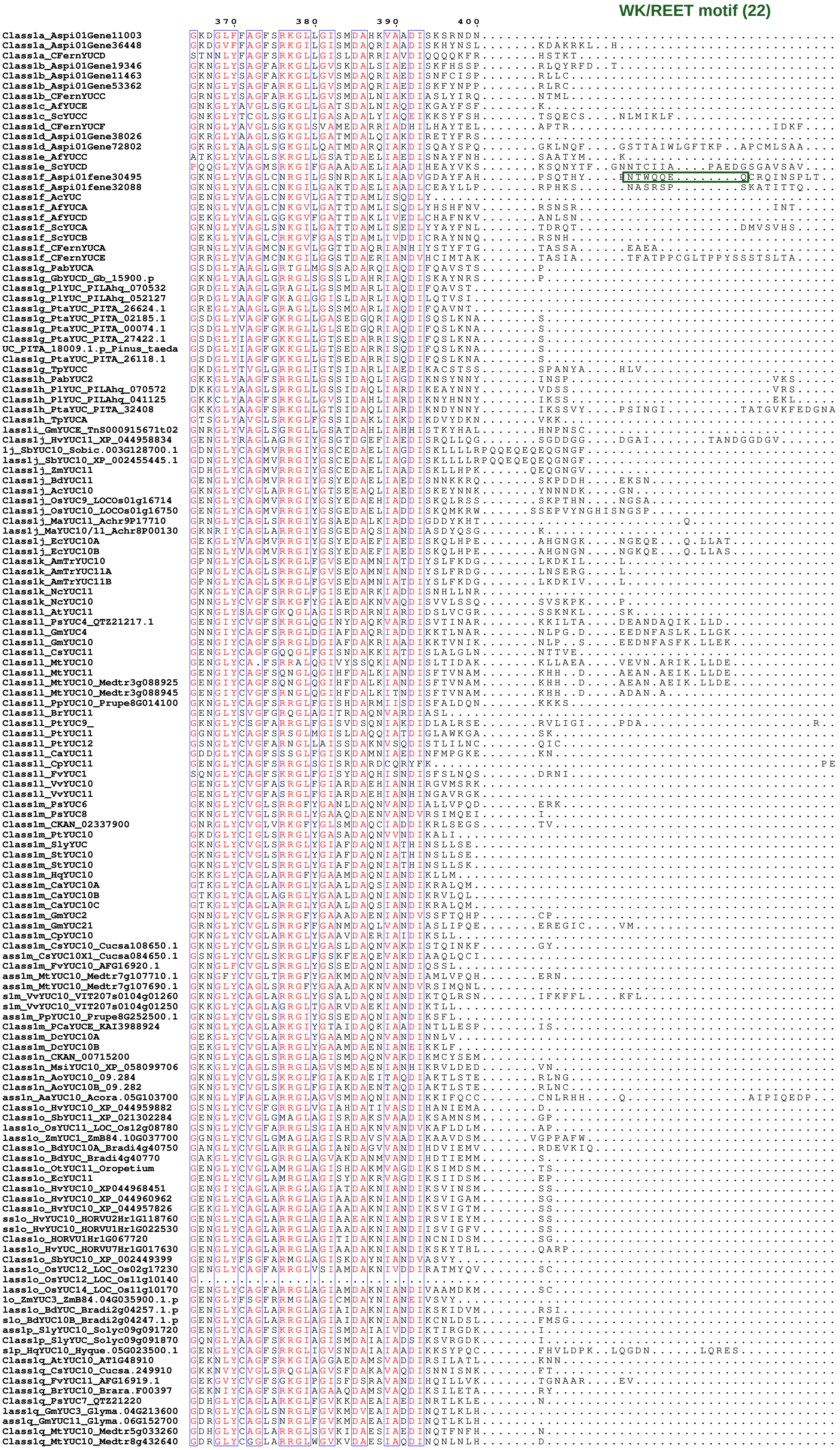

```
Class1a_Aspi01Gene11003          ...............................IV
Class1a_Aspi01Gene36448          ..............................LHV
Class1a_CFernYUCD                .................................
Class1b_Aspi01Gene19346          .................................
Class1b_Aspi01Gene11463          .................................
Class1b_Aspi01Gene53362          .................................
Class1b_CFernYUCC                .................................
Class1c_AfYUCE                   .................................
Class1c_ScYUCC                   .................................
Class1d_CFernYUCF                ....I............................
Class1d_Aspi01Gene38026          .....................HDGR........
Class1d_Aspi01Gene72802          ....VVIVNYIFLAHLFYCCLHMIRGARLSMRN
Class1e_AfYUCC                   .................................
Class1e_ScYUCD                   .................................
Class1f_Aspi01fene30495          .................................
Class1f_Aspi01fene32088          ....IPL..........................
Class1f_AcYUC                    .................................
Class1f_AfYUCA                   .................................
Class1f_AfYUCD                   .................................
Class1f_ScYUCA                   .................................
Class1f_ScYUCB                   .................................
Class1f_CFernYUCA                .................................
Class1f_CFernYUCE                .................................
Class1g_PabYUCA                  .................................
Class1g_GbYUCD_Gb_15900.p        .................................
Class1g_PlYUC_PILAhq_070532      .................................
Class1g_PlYUC_PILAhq_052127      .................................
Class1g_PtaYUC_PITA_26624.1      .................................
Class1g_PtaYUC_PITA_02185.1      .................................
Class1g_PtaYUC_PITA_00074.1      .................................
Class1g_PtaYUC_PITA_27422.1      .................................
UC_PITA_18009.1.p_Pinus_taeda    .................................
Class1g_PtaYUC_PITA_26118.1      .................................
Class1g_TpYUCC                   .................................
Class1h_PabYUC2                  .................................
Class1h_PlYUC_PILAhq_070572      .................................
Class1h_PlYUC_PILAhq_041125      .................................
Class1h_PtaYUC_PITA_32408        EDYDVIILATGYRSDVLRWLKDDGKFLP.....
Class1h_TpYUCA                   .................................
lass1i_GmYUCE_TnS000915671t02    .................................
Class1j_HvYUC11_XP_044958834     .................................
1j_SbYUC10_Sobic.003G128700.1    .................................
lass1j_SbYUC10_XP_002455445.1    .................................
Class1j_ZmYUC11                  .................................
Class1j_BdYUC11                  .................................
Class1j_AcYUC10                  .................................
Class1j_OsYUC9_LOCOs01g16714     .................................
Class1j_OsYUC10_LOCOs01g16750    .................................
Class1j_MaYUC11_Achr9P17710      .................................
lass1j_MaYUC10/11_Achr8P00130    .................................
Class1j_EcYUC10A                 .................................
Class1j_EcYUC10B                 .................................
Class1k_AmTrYUC10                .................................
Class1k_AmTrYUC11A               .................................
Class1k_AmTrYUC11B               .................................
Class1k_NcYUC11                  ..............HK.................
Class1k_NcYUC10                  .................................
Class1l_AtYUC11                  .................................
Class1l_PsYUC4_QTZ21217.1        .................................
Class1l_GmYUC4                   .................................
Class1l_GmYUC10                  .................................
Class1l_CsYUC11                  .................................
Class1l_MtYUC10                  .................................
Class1l_MtYUC11                  .................................
Class1l_MtYUC10_Medtr3g088925    .................................
Class1l_MtYUC10_Medtr3g088945    .................................
Class1l_PpYUC10_Prupe8G014100    .................................
Class1l_BrYUC11                  .................................
Class1l_PtYUC9_                  ..VC.............................
Class1l_PtYUC11                  .................................
Class1l_PtYUC12                  .................................
Class1l_CaYUC11                  .................................
Class1l_CpYUC11                  .................................
Class1l_FvYUC1                   .................................
Class1l_VvYUC10                  .................................
Class1l_VvYUC11                  .................................
Class1m_PsYUC6                   .................................
Class1m_PsYUC8                   .................................
Class1m_CKAN_02337900            .................................
Class1m_PtYUC10                  .................................
Class1m_SlyYUC                   .................................
Class1m_StYUC10                  .................................
Class1m_StYUC10                  .................................
Class1m_HqYUC10                  .................................
Class1m_CaYUC10A                 .................................
Class1m_CaYUC10B                 .................................
Class1m_CaYUC10C                 .................................
Class1m_GmYUC2                   .................................
Class1m_GmYUC21                  .................................
Class1m_CpYUC10                  .................................
Class1m_CsYUC10_Cucsa108650.1    .................................
ass1m_CsYUC10X1_Cucsa084650.1    .................................
Class1m_FvYUC10_AFG16920.1       .................................
ass1m_MtYUC10_Medtr7g107710.1    .................................
ass1m_MtYUC10_Medtr7g107690.1    .................................
s1m_VvYUC10_VIT207s0104g01260    .................................
s1m_VvYUC10_VIT207s0104g01250    .................................
ass1m_PpYUC10_Prupe8G252500.1    .................................
Class1m_PCaYUCE_KAI3988924       .................................
Class1m_DcYUC10A                 .................................
Class1m_DcYUC10B                 .................................
Class1n_CKAN_00715200            .................................
Class1n_MsiYUC10_XP_058099706    .................................
Class1n_AoYUC10_09.284           .................................
Class1n_AoYUC10B_09.282          .................................
ass1n_AaYUC10_Acora.05G103700    ...C.M.....Y.....................
Class1o_HvYUC10_XP_044959882     ...........F..N..................
Class1o_SbYUC11_XP_021302284     ...........F.....................
lass1o_OsYUC11_LOC_Os12g08780    ...........F.....................
lass1o_ZmYUC1_ZmB84.10G037700    ...........F.....................
Class1o_BdYUC10A_Bradi4g40750    ...........L.....................
Class1o_BdYUC_Bradi4g40770       ...........F..N..................
Class1o_OtYUC11_Oropetium        ...........S.....................
Class1o_EcYUC11                  ...........F.....................
Class1o_HvYUC10_XP044968451      .................................
Class1o_HvYUC10_XP_044960962     .................................
Class1o_HvYUC10_XP_044957826     .................................
ss1o_HvYUC10_HORVU2Hr1G118760    .................................
ss1o_HvYUC10_HORVU1Hr1G022530    .................................
Class1o_HORVU1Hr1G067720         .................................
lass1o_HvYUC_HORVU7Hr1G017630    .................................
Class1o_SbYUC10_XP_002449399     .................................
lass1o_OsYUC12_LOC_Os02g17230    .................................
lass1o_OsYUC12_LOC_Os11g10140    .................................
lass1o_OsYUC14_LOC_Os11g10170    .................................
1o_ZmYUC3_ZmB84.04G035900.1.p    .................................
lass1o_BdYUC_Bradi2g04257.1.p    .................................
s1o_BdYUC10B_Bradi2g04247.1.p    .................................
ass1p_SlyYUC10_Solyc09g091720    .................................
Class1p_SlyYUC_Solyc09g091870    .................................
s1p_HqYUC10_Hyque.05G023500.1    .................................
Class1q_AtYUC10_AT1G48910        .................................
Class1q_CsYUC10_Cucsa.249910     .................................
Class1q_FvYUC11_AFG16919.1       .................................
Class1q_BrYUC10_Brara.F00397     .................................
Class1q_PsYUC7_QTZ21220          .................................
lass1q_GmYUC3_Glyma.04G213600    .................................
ass1q_GmYUC11_Glyma.06G152700    .................................
Class1q_MtYUC10_Medtr5g033260    .................................
Class1q_MtYUC10_Medtr8g432640    .................................
```

------------------------------------------------------------------------------------------------------------------------------------------------------

#### Class-2 YUCCAs

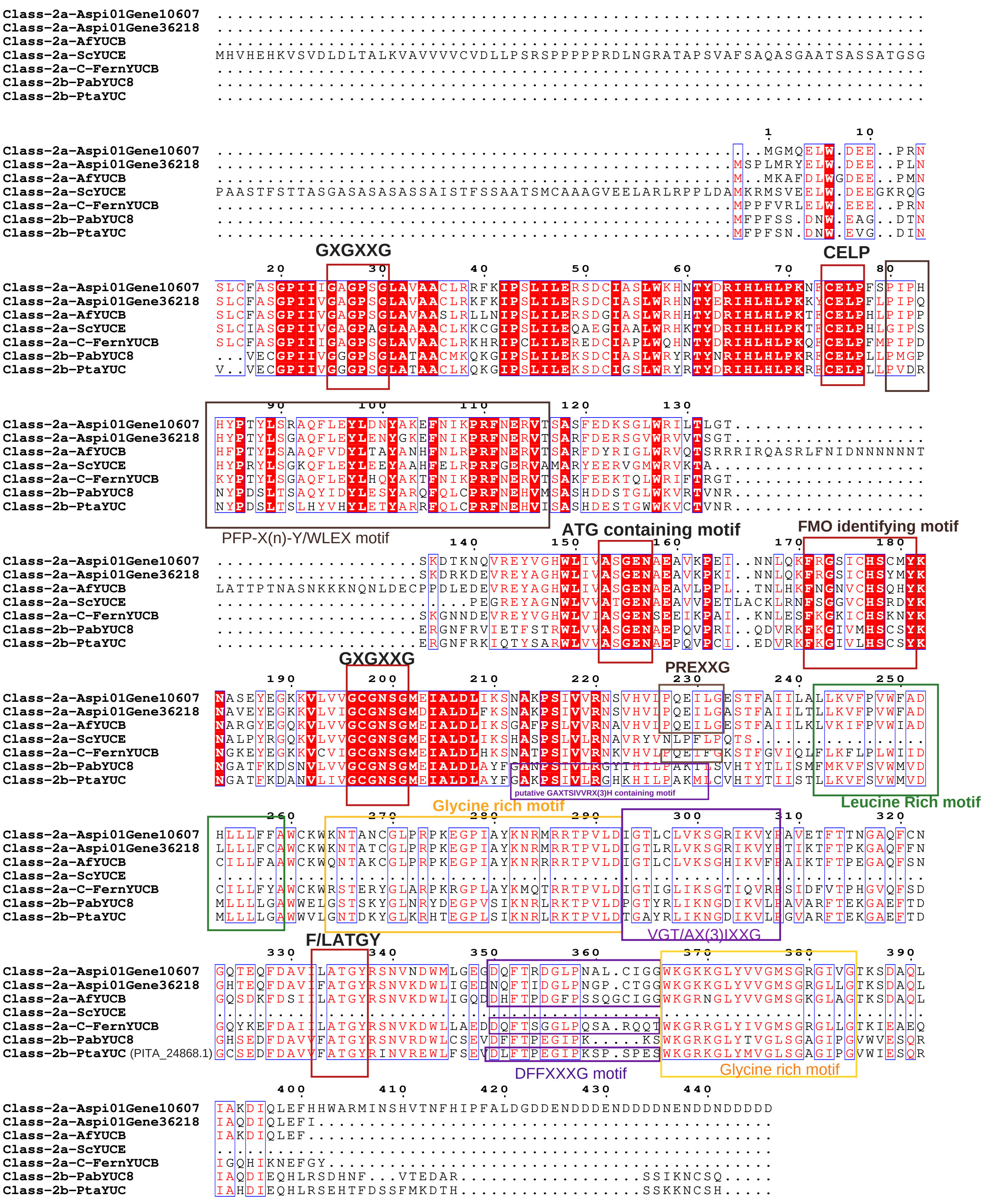

### Class-3 YUCCAs

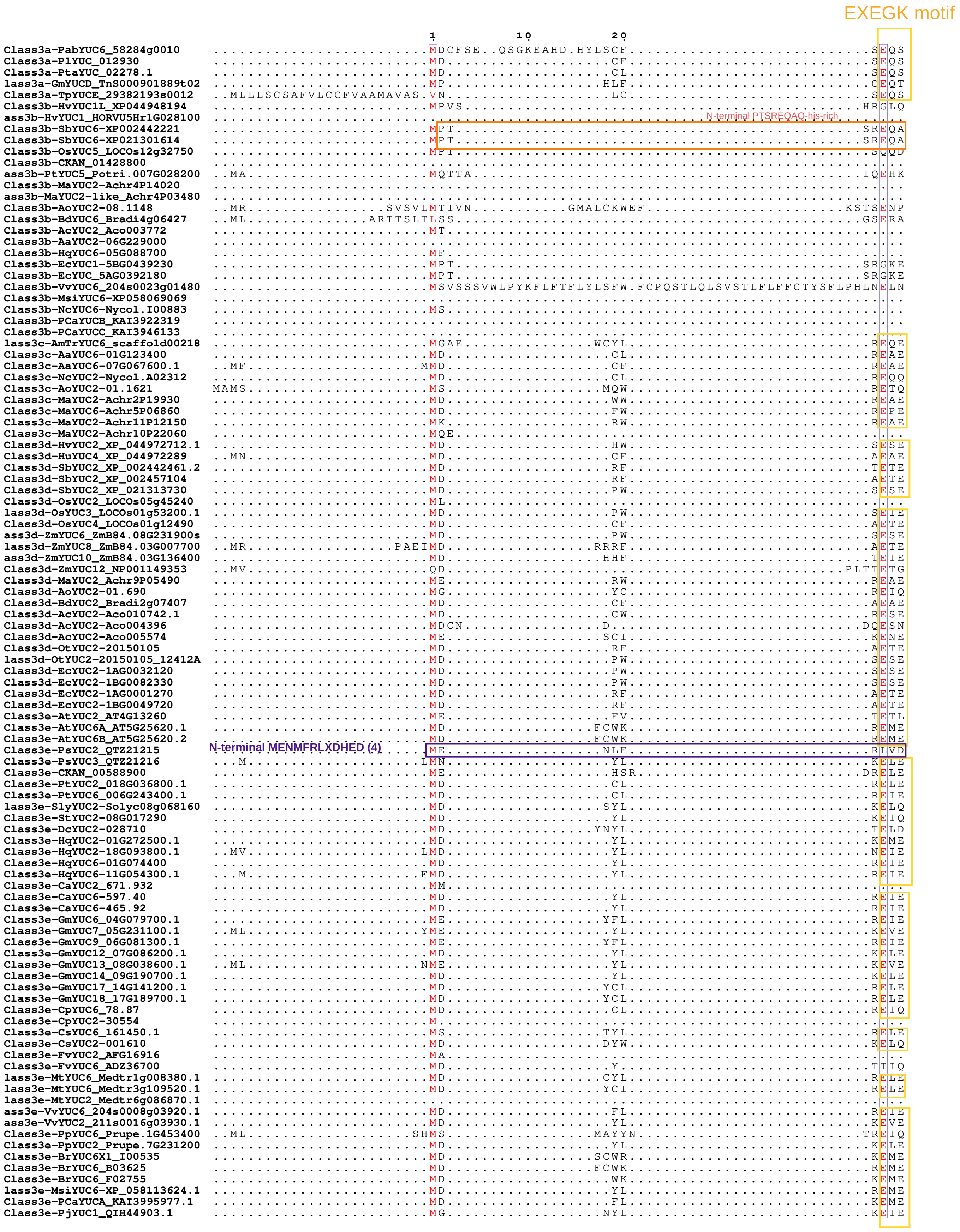


Notes:

N-terminal MENMFRLXDHED present in PsYUC2-Class 3E (and this motif is common in 4f as well)

N-terminal PTSREQAQ-his-rich motif is exclusively present in SbYUC6 alone (in 3b subclass).

EXEGK motif is mostly present

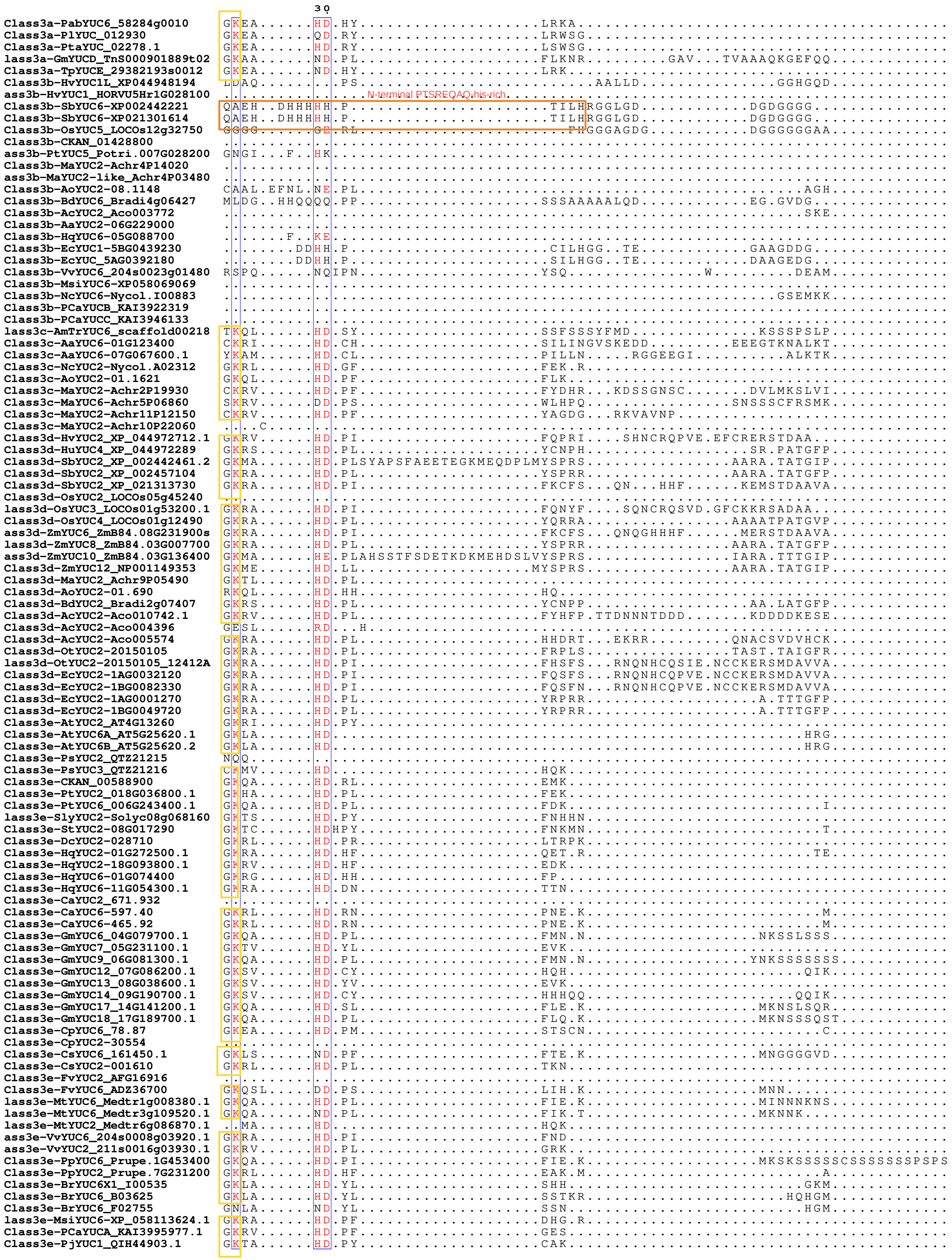

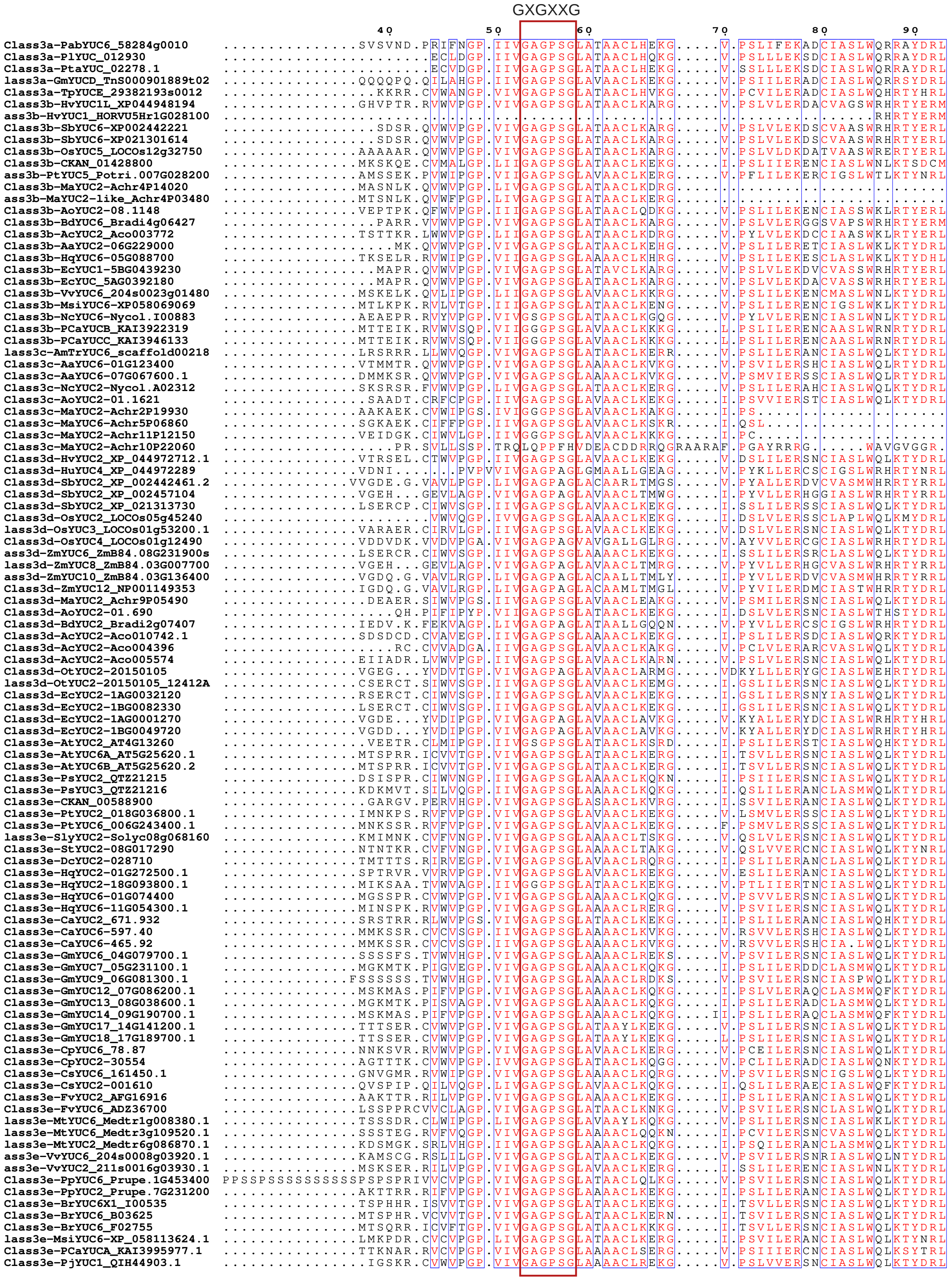
GXGXXG
40
50
60
70
80
90

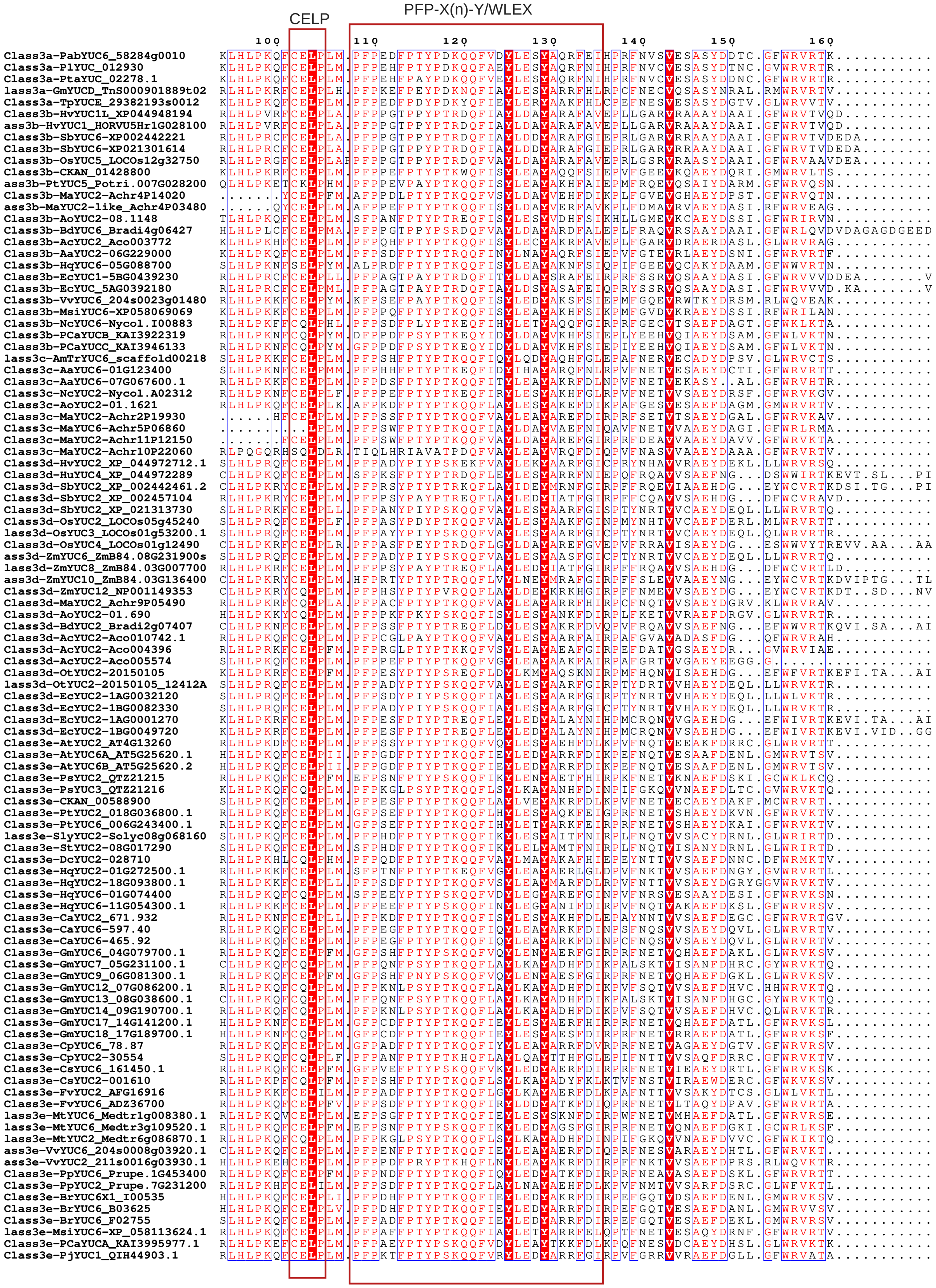
CELP
PFP-X(n)-Y/WLEX
100
110
120
130
140
150
160

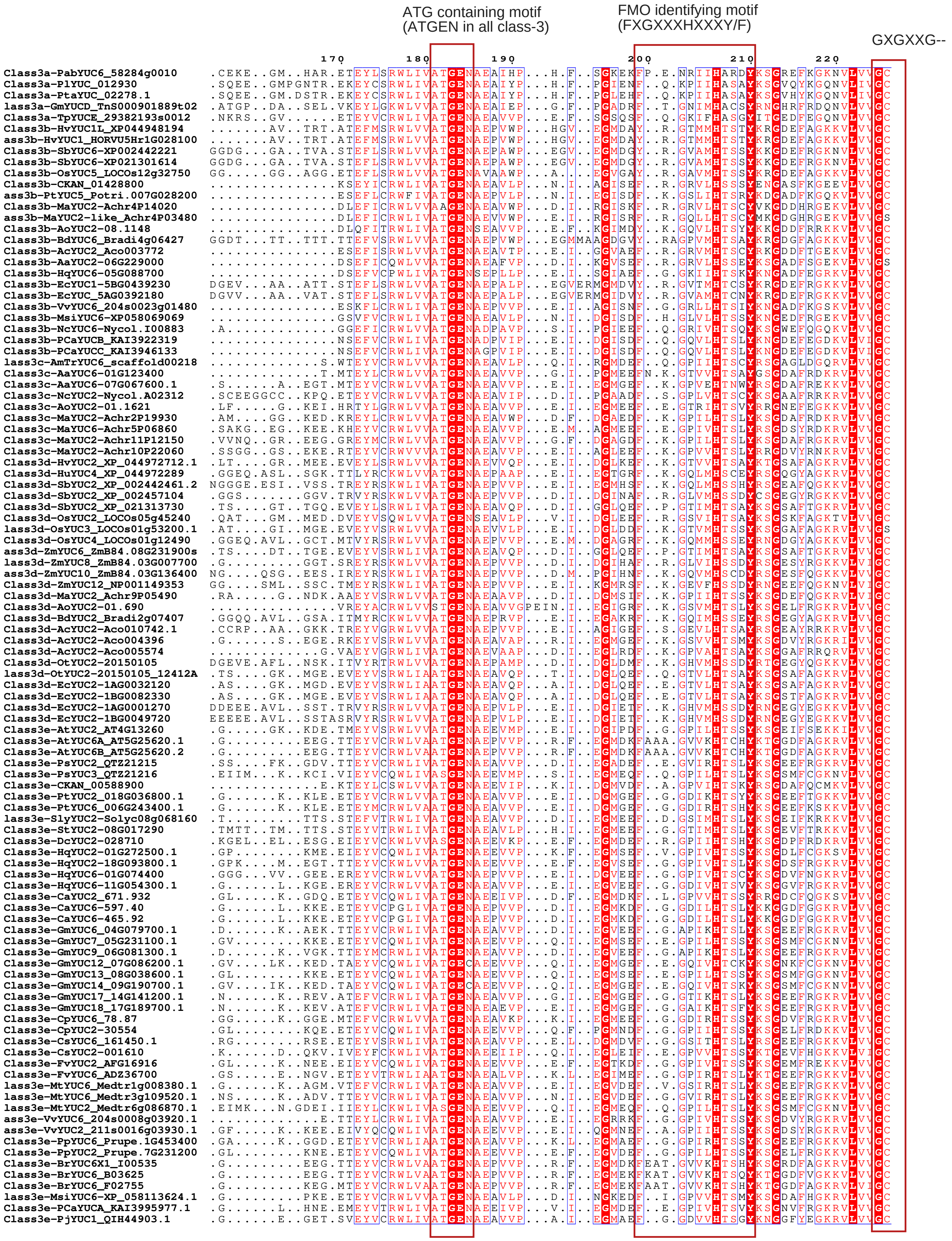
ATG containing motif
(ATGEN in all class-3)
FMO identifying motif
(FXGXXXHXXXY/F)
GXGXXG--

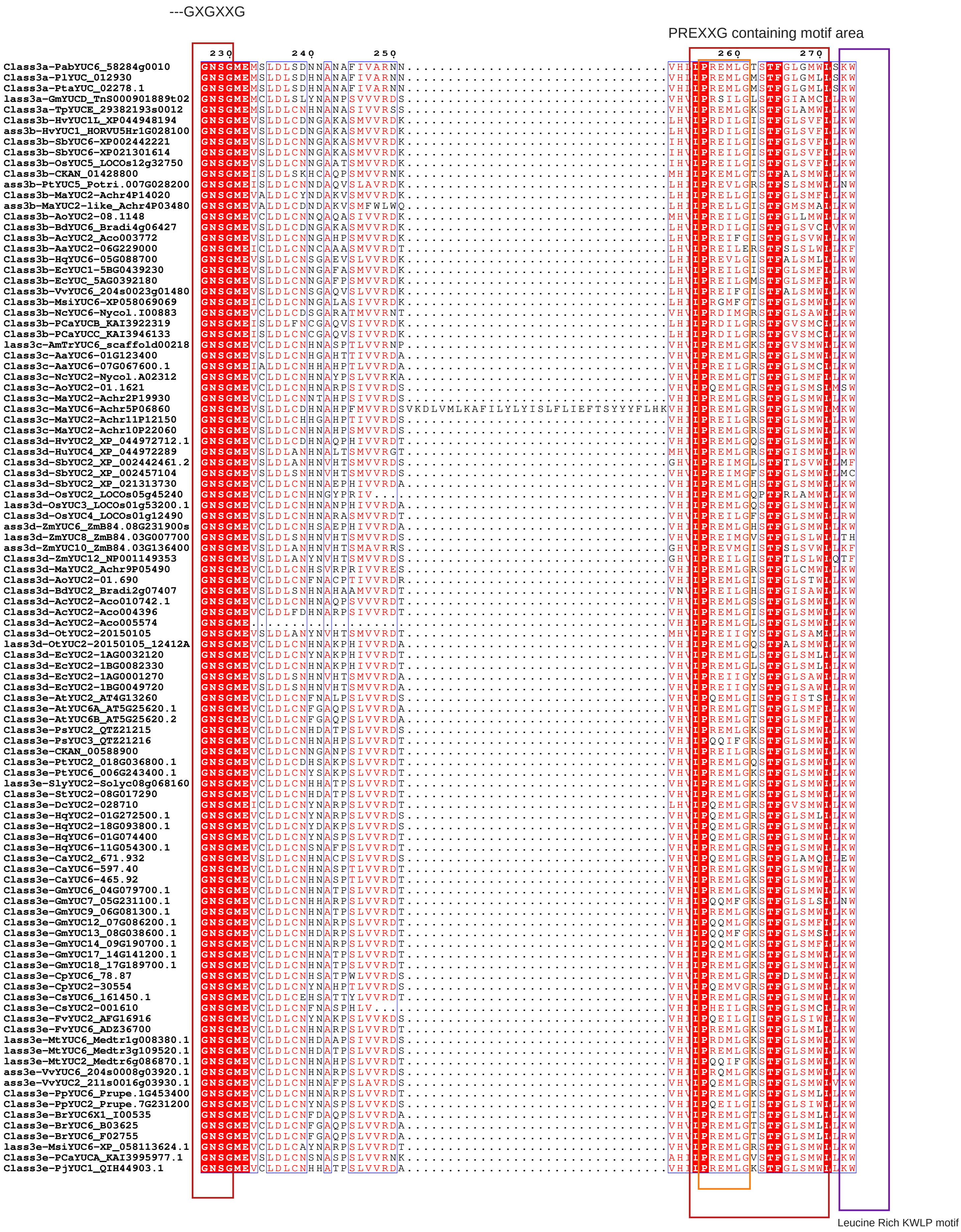


Notes:
The second GXGXXG is conserved highly along with neighbouring residues 'ME'

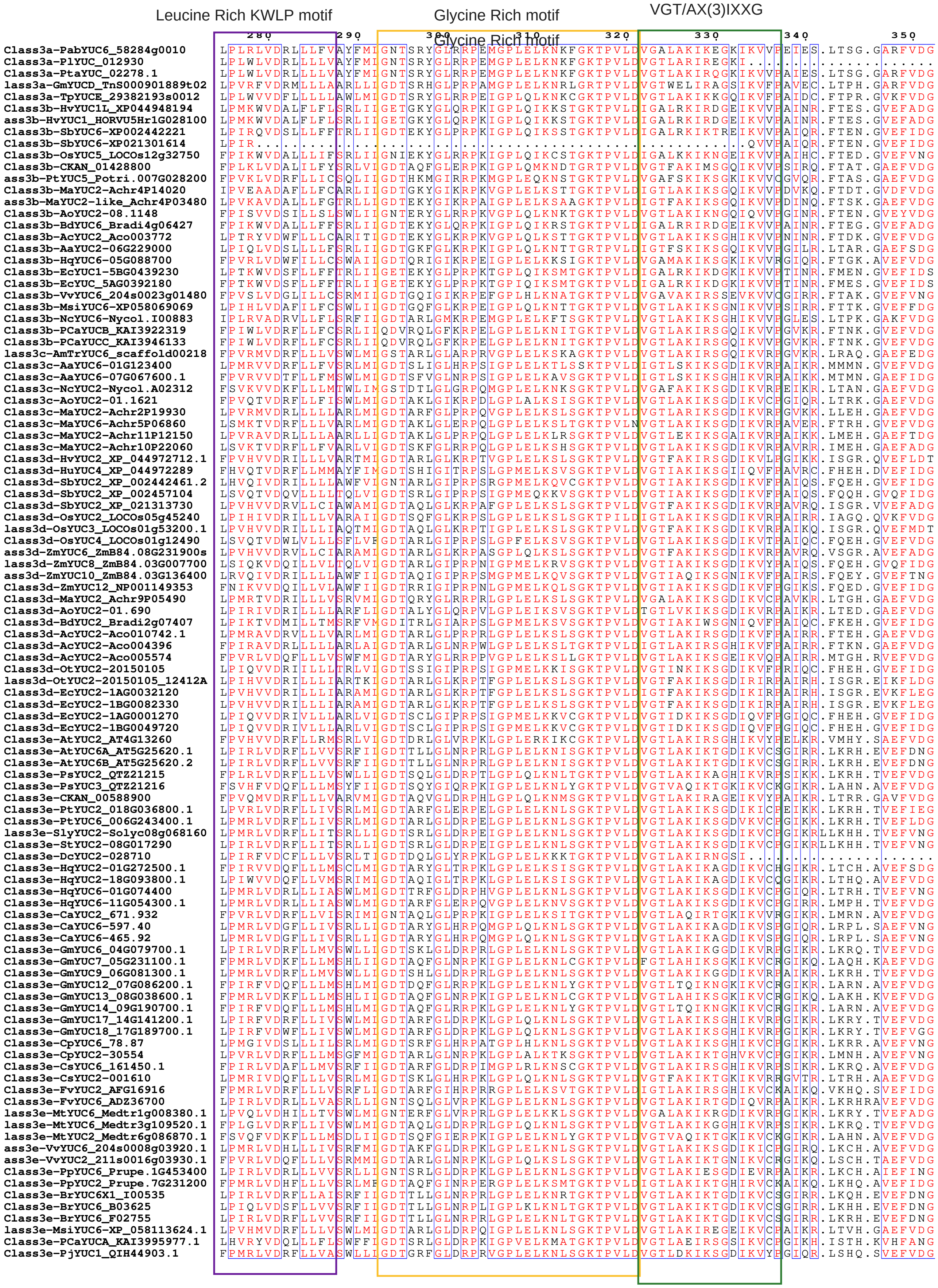
Leucine Rich KWLP motif
Glycine Rich motif
VGT/AX(3)IXXG

#### Class-3 YUCCAs

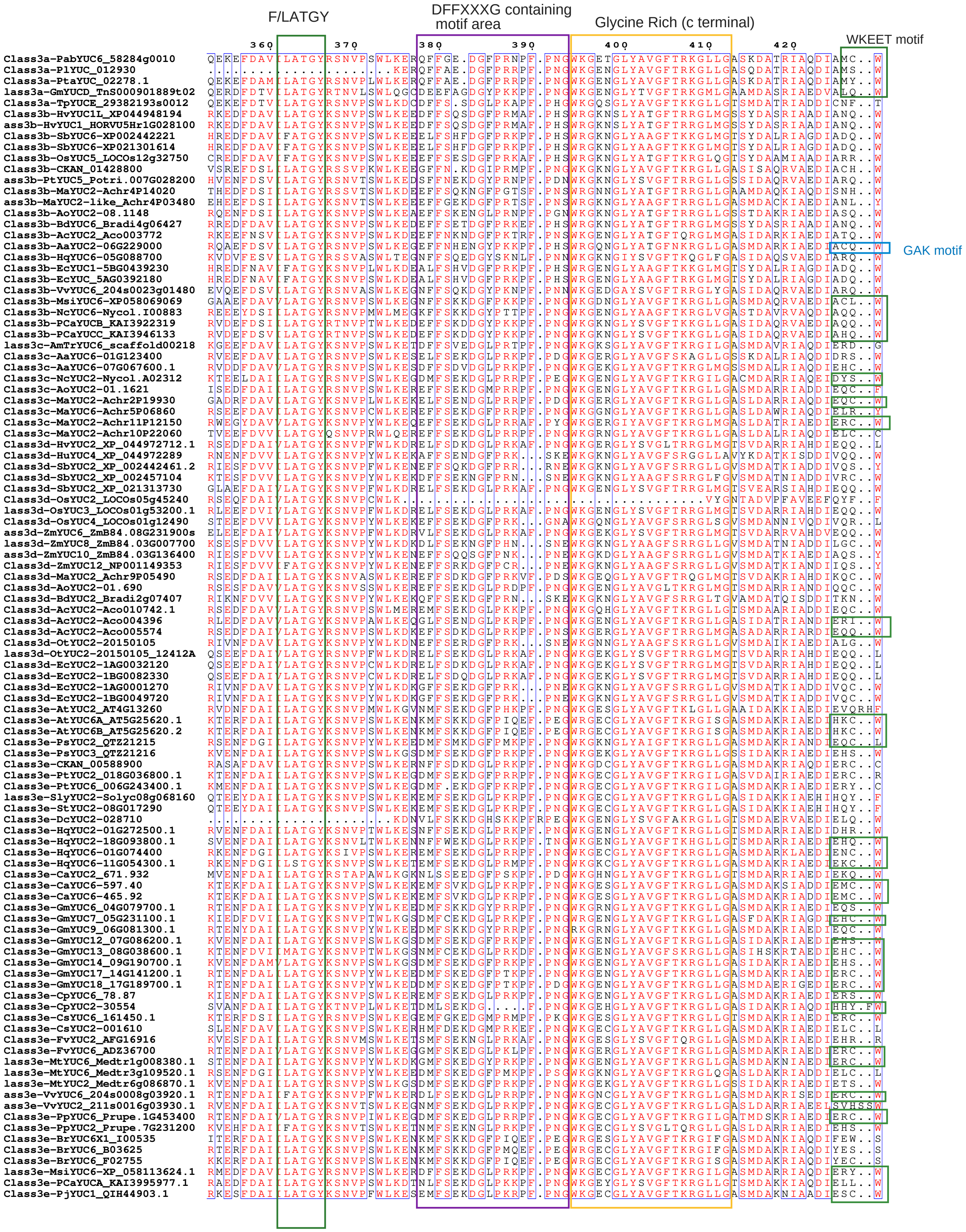

```
                                  430       440
Class3a-PabYUC6_58284g0010       .KAE.SVASKG.SQTNN................................................
Class3a-PlYUC_012930             .KAE.AVSKRA.STPAAAN..............................................
Class3a-PtaYUC_02278.1           .KAE.VVSKSA.STSAAAN..............................................
lass3a-GmYUCD_TnS000901889t02    .KAE.EARAER.SHFS.................................................
Class3a-TpYUCE_29382193s0012     .KKG.LLGTSM.DATRVAEDIFNSMDATQIAERSRWRAEAKQFIIYNKPPC..............
Class3b-HvYUC1L_XP044948194      .TEA.LARNIT.AHNNA................................................
ass3b-HvYUC1_HORVU5Hr1G028100    .TEA.LARNIT.AHNNA................................................
Class3b-SbYUC6-XP002442221       .TET.FASPTA.TNRSSDHGA............................................
Class3b-SbYUC6-XP021301614       .TET.FASPTA.TNRSSDHGA............................................
Class3b-OsYUC5_LOCOs12g32750     .TKS.LAGPTA.AADADHHETYIAN........................................
Class3b-CKAN_01428800            .DCE.TKHLCL.ELQTILN..............................................
ass3b-PtYUC5_Potri.007G028200    .NCE.TKHLRI.ES...................................................
Class3b-MaYUC2-Achr4P14020       .SCR.S...........................................................
ass3b-MaYUC2-like_Achr4P03480    .GAV.DIRHKL.NTITS................................................
Class3b-AoYUC2-08.1148           .NLK.PKHPPS.DL...................................................
Class3b-BdYUC6_Bradi4g06427      .TEA.LAASSV.AAAAADTN.............................................
Class3b-AcYUC2_Aco003772         .NSK.PKHLPC.GLL..................................................
GAK motif
Class3b-AaYUC2-06G229000         .NSG.AKHLPL.QLLFTS...............................................
Class3b-HqYUC6-05G088700         .NSE.MKHLSF.QF...................................................
Class3b-EcYUC1-5BG0439230        .TEA.LSSPTL.LLAGASTVRD.........................A.................
Class3b-EcYUC_5AG0392180         .TEA.LASPTL.LLTGASTVRD.........................A.................
Class3b-VvYUC6_204s0023g01480    .KSQ.MKHLHL.DL...................................................
Class3b-MsiYUC6-XP058069069      .KSE.TKHLRL.EM...................................................
Class3b-NcYUC6-Nycol.I00883      .NQE.AKYFTF.....................................PPSKK..........NI
Class3b-PCaYUCB_KAI3922319       .KSE.NKHFHL.RV...................................................
Class3b-PCaYUCC_KAI3946133       .KSE.TKHFHL.RV...................................................
lass3c-AmTrYUC6_scaffold00218    .RTSQGI..........................................................
Class3c-AaYUC6-01G123400         .KSI.SLLDMI.SSS..................................................
Class3c-AaYUC6-07G067600.1       .KAH.A...........................................................
Class3c-NcYUC2-Nycol.A02312      .KAE.SKPIFL.ATATPTSLPY...........................................
Class3c-AoYUC2-01.1621           .EVE.ANQFMV.FSCSPN...............................................
Class3c-MaYUC2-Achr2P19930       .KAE.LKKIMI..................................FACP................
Class3c-MaYUC6-Achr5P06860       .KAT.KK..........................................................
Class3c-MaYUC2-Achr11P12150      .KAE.GSIAGG.GNHVMTNLSVQTY....IATKSCW....SQNLCFMSNAIKQTNQTETQRS...
Class3c-MaYUC2-Achr10P22060      .KTD.ENCR........................................................
Class3d-HvYUC2_XP_044972712.1    .SAE.GKL.........................................................
Class3d-HuYUC4_XP_044972289      .RNM.CMEDVR.ESSSNQR............................TP.............CPI
Class3d-SbYUC2_XP_002442461.2    .NNI.............................................................
Class3d-SbYUC2_XP_002457104      .NDF.GYERHK.RK...................................................
Class3d-SbYUC2_XP_021313730      .KAK.GTHPDA.G....................................................
Class3d-OsYUC2_LOCOs05g45240     .............................................................S...
lass3d-OsYUC3_LOCOs01g53200.1    .KAR.GKHPGV.LL...................................................
Class3d-OsYUC4_LOCOs01g12490     .HDM.GYERSE.NN...................................................
ass3d-ZmYUC6_ZmB84.08G231900s    RKAK.GTHRDG.VPLPLVAVVYHG......................................DGL
lass3d-ZmYUC8_ZmB84.03G007700    .NDF.GYERHK.RK...................................................
ass3d-ZmYUC10_ZmB84.03G136400    .NNI.YKLQRS......................................................
Class3d-ZmYUC12_NP001149353      .NNI.IFHPMDIQRSQDD...............................................
Class3d-MaYUC2_Achr9P05490       .MAE.PKQRML.PSQT.................................................
Class3d-AoYUC2-01.690            .KND.AKRCML......................................................
Class3d-BdYUC2_Bradi2g07407      .RDM.HTRDVR.EDPSSRSGTVIFN........................................
Class3d-AcYUC2-Aco010742.1       .KVQ.KASQLV.MSFSLPVIN............................................
Class3d-AcYUC2-Aco004396         .KTQH............................................................
Class3d-AcYUC2-Aco005574         .NAE.TKHGQS.....................RS...............................
Class3d-OtYUC2-20150105          ............................................MF...................
lass3d-OtYUC2-20150105_12412A    .QSQ.GLRPDV.FL...................................................
Class3d-EcYUC2-1AG0032120        .KSQ.GLHPDV.FL...................................................
Class3d-EcYUC2-1BG0082330        .KSQ.GLHPDV.FL...................................................
Class3d-EcYUC2-1AG0001270        .HDM.GYGRQK.SK...................................................
Class3d-EcYUC2-1BG0049720        .LDM.GYGRQK.SK...................................................
Class3e-AtYUC2_AT4G13260         ...............................................LPLARPQHC.........
Class3e-AtYUC6A_AT5G25620.1      .KQDEQVK.KI......................................................
Class3e-AtYUC6B_AT5G25620.2      .KQDEQLQCKL.GKRMKRK.FS.........................ES.....DCG.....GN.
Class3e-PsYUC2_QTZ21215          .KAE.AKHGS.......................................................
Class3e-PsYUC3_QTZ21216          .KAL.KAKPLA......................................................
Class3e-CKAN_00588900            .KPR.LSNSKP.....................................PSGWHK...........
Class3e-PtYUC2_018G036800.1      .NEE.AAPCDR....SVLMKS...........................PPPPPPSFSCLE..FDL
Class3e-PtYUC6_006G243400.1      .RNEEAAPYDH.HHRSVLLLK...........................SSSSSPS........P.
lass3e-SlyYUC2-Solyc08g068160    .Q...............................................................
Class3e-StYUC2-08G017290         .Q...............................................................
Class3e-DcYUC2-028710            .YLK.DSKIIT.ESVDQVPEIERVL.....................................CNL
Class3e-HqYUC2-01G272500.1       .NSE.GNEFNA......................................................
Class3e-HqYUC2-18G093800.1       .KSE.KV..........................................................
Class3e-HqYUC6-01G074400         .RME.SKHLSR.FASSYY...............................................
Class3e-HqYUC6-11G054300.1       .KDE.LKHLTA.FAHSQ................................................
Class3e-CaYUC2_671.932           .KVR.AVQFQG..................................LC..................
Class3e-CaYUC6-597.40            .KAE.SKHFSY.FARPSSLQS............................................
Class3e-CaYUC6-465.92            .KAE.SKHFSY.FARPSSLQS............................................
Class3e-GmYUC6_04G079700.1       .KAG.ANHRTT.FARSHL..............................PQPNS............
Class3e-GmYUC7_05G231100.1       .KAA.N...........................................................
Class3e-GmYUC9_06G081300.1       .ETG.ANHRTT.LAPSHL..............................PQPNS............
Class3e-GmYUC12_07G086200.1      .KAE.ATHVLE.FPCPLA...............................................
Class3e-GmYUC13_08G038600.1      .KAA.NTTRV.......................................................
Class3e-GmYUC14_09G190700.1      .KAE.AKHVLE.FPRPLA...............................................
Class3e-GmYUC17_14G141200.1      .KAK.HSTS...FSLSLNV.............................PQSNS............
Class3e-GmYUC18_17G189700.1      .KEK.HSSTS..FSLSLNV.............................PQSNS............
Class3e-CpYUC6_78.87             .KLS.IAFGSQ..........S...........................................
Class3e-CpYUC2-30554             .KAE.ASSFMA.FPTRPP..............................P................
Class3e-CsYUC6_161450.1          .KAD.AKLCTP.TMQSPPST.............................................
Class3e-CsYUC2-001610            .EN..............................................................
Class3e-FvYUC2_AFG16916          .KAEKATHIMPRLVTCAS..............................PLAQP............
Class3e-FvYUC6_ADZ36700          .KAE.AKHSTP.FTRSHL..............................PLSSSP...........
lass3e-MtYUC6_Medtr1g008380.1    .KAE.AKHIFP.QSNS.................................................
lass3e-MtYUC6_Medtr3g109520.1    .ESE.AKYGS.......................................................
lass3e-MtYUC2_Medtr6g086870.1    .KSI.KAKPLA......................................................
ass3e-VvYUC6_204s0008g03920.1    .KAD.AKRLTV.KSHT.................................................
ass3e-VvYUC2_211s0016g03930.1    .P...FQHFRH.DNHDL................................................
Class3e-PpYUC6_Prupe.1G453400    .KAE.AKHCTP.FKGSLF..............................PLSSSSSP.........
Class3e-PpYUC2_Prupe.7G231200    .KAE.ATHFMA.FTACAL..............................PPTPPPQP.........
Class3e-BrYUC6X1_I00535          .RKSYQAHRHI.QVLCMSRRSG.........................QPYSRLLE..........
Class3e-BrYUC6_B03625            .RKSGQPHRHI.QVFMARK.SD.........................QACSRVLDG.........
Class3e-BrYUC6_F02755            .RKSDQAHRHI.QVFMSSK.PD.........................QA................
lass3e-MsiYUC6-XP_058113624.1    .KAK.TKQFMA.FSCPPTAS.............................................
Class3e-PCaYUCA_KAI3995977.1     .KAD.SKHYLS.ALFYSPSP............................PPPPSLGH.........
Class3e-PjYUC1_QIH44903.1        .RAE.AKHLSA.ISSPCSMQKH..........................ILSDQLINELIN.....
```

-WKEET motif area highlighted in green rectangle, but this is not conserved in all

-----------------------------------------------------------------------------------------------------------------

### Class-4 YUCCAs

| | |
|---|---|
| Class4A-GbYUCB_Gb32507 | ................................................................................ |
| Class4A-PlYUC_PILAhq064253 | ................................................................................ |
| Class4A-PtaYUC_PITA32685 | ................................................................................ |
| Class4A-PtaYUC_PITA21706 | ................................................................................ |
| Class4A-TpYUCD_29380002s0001 | ................................................................................ |
| Class4B-PabYUC5_218308g0010 | ................................................................................ |
| Class4B-PabYUC_5857317g0010 | ................................................................................ |
| Class4B-GbYUCA_39502 | ................................................................................ |
| Class4B-PlYUC_045322 | ................................................................................ |
| Class4B-PlYUC_038950 | ................................................................................ |
| Class4B-PtaYUC_06172 | ................................................................................ |
| Class4B-PtaYUC_03225 | ................................................................................ |
| Class4B-GmYUCCTnS000183447t02 | ................................................................................ |
| Class4c-PabYUCB_215174g0010 | ................................................................................ |
| Class4c-GbYUCC_Gb40961 | ................................................................................ |
| Class4c-PlYUC_075634 | .................................ML............................................. |
| Class4c-PtaYUC_05784 | ................................................................................ |
| Class4c-GmYUCATnS000225445t01 | ................................................................................ |
| Class4c-TpYUCB_29377126s0001 | ................................................................................ |
| Class4D-AmTrYUC_0003987 | ................................................................................ |
| Class4D-CKAN-01805100 | ................................................................................ |
| Class4D-CKAN-02673000 | ................................................................................ |
| Class4D-CKAN-02545400 | ................................................................................ |
| Class4D-CKAN-01519000 | ................................................................................ |
| Class4D-MsiYUC5-XP058095048 | ................................................................................ |
| Class4D-MsiYUC5-XP-058101305 | ................................................................................ |
| Class4D-NcYUC5_Nycol.L00992 | ................................................................................ |
| Class4D-NcYUC5_Nycol.D01636 | ................................................................................ |
| Class4E-HvYUC3_KAE8779170 | ................................................................................ |
| Class4E-HvYUC5_XP044971244 | ................................................................................ |
| Class4E-HvYUC5_XP044970914 | ................................................................................ |
| Class4E-HvYUC5-1G116980 | RALVANRIPSPHQTPT..........................................HP.................... |
| Class4E-HvYUC9_XP_044950031 | ................................................................................ |
| Class4E-HvYUC9-1G050630 | ................................................................................ |
| Class4E-SbYUC5_XP_002447449 | ................................................................................ |
| Class4E-SbYUC5_XP_021308125 | ................................................................................ |
| Class4E-SbYUC5_XP_002461905.2 | ................................................................................ |
| Class4E-SbYUC_EES15155 | ................................................................................ |
| Class4E-OsYUC6_LOCOs07g25540 | ................................................................................ |
| Class4E-OsYUC7_LOC_Os04g03980 | ................................................................................ |
| Class4E-ZmYUC2_ZmB840G121000 | ................................................................................ |
| Class4E-ZmYUC4_02G199500 | ................................................................................ |
| Class4E-ZmYUC5_07G064900.2 | ................................................................................ |
| Class4E-ZmYUC7_NP001358724 | ................................................................................ |
| Class4E-MaYUC3_P04200 | ................................................................................ |
| Class4E-MaYUC5-Achr4P25050 | ................................................................................ |
| Class4E-MaYUC5-Achr5P20550 | ................................................................................ |
| Class4E-MaYUC5-Achr8P12830 | ................................................................................ |
| Class4E-MaYUC5-Achr2P04250 | ................................................................................ |
| Class4E-MaYUC3-Achr11P07720 | ................................................................................ |
| Class4E-MaYUC5-Achr1P03360 | ................................................................................ |
| Class4E-MaYUC5-Achr11P21300 | ................................................................................ |
| Class4E-MaYUC5-Achr1P24210 | ................................................................................ |
| Class4E-AoYUC5-04.275 | ................................................................................ |
| Class4E-AoYUC3-07910 | ................................................................................ |
| Class4E-BdYUC5_Bradi5g01327 | ................................................................................ |
| Class4E-BdYUC9_Bradi1g28967 | ................................................................................ |
| Class4E-AcYUC5_Aco014634 | ................................................................................ |
| Class4E-AcYUC5_Aco008902 | ................................................................................ |
| Class4E-OtYUC5-25620A | ................................................................................ |
| Class4E-OtYUC_13179A | ................................................................................ |
| Class4E-AaYUC5-01G200900 | ................................................................................ |
| Class4E-AaYUC5-Acora0G153700 | ................................................................................ |
| Class4E-EcYUC5-7AG0564960 | ................................................................................ |
| Class4E-EcYUC5.4BG0359200 | ................................................................................ |
| Class4E-EcYUC5.4AG0327230 | ................................................................................ |
| Class4f-AtYUC3-AT1G04610 | ..MYGN.......................................................................... |
| Class4f-AtYUC5-AT5G43890 | ................................................................................ |
| Class4f-AtYUC7-AT2G33230 | ..MCNN.......................................................................... |
| Class4f-AtYUC8-AT4G28720 | ................................................................................ |
| Class4f-AtYUC9-AT1G04180 | ................................................................................ |
| Class4f-PsYUC5-QTZ21218 | ................................................................................ |
| Class4f-SlyYUC3-09g064160 | ................................................................................ |
| Class4f-SlyYUC5-06g083700 | ................................................................................ |
| Class4f-SlyYUC3-09g091090 | ................................................................................ |
| Class4f-SlyYUC8-06g008050 | ................................................................................ |
| Class4f-StYUC3-09G018700 | ................................................................................ |
| Class4f-StYUC3-09G027910 | ................................................................................ |
| Class4f-StYUC5-06G034100 | ................................................................................ |
| Class4f-StYUC3-06G002120 | ................................................................................ |
| lass4f-GmYUC1-Glyma.03G169600 | ................................................................................ |
| Class4f-GmYUC15-Glyma0G128700 | ................................................................................ |
| Class4f-GmYUC20-Glyma9G170800 | ................................................................................ |
| Class4f-GmYUC22-20G080000 | ................................................................................ |
| Class4f-CsYUC8-348750 | ................................................................................ |
| Class4f-CsYUC5-40470 | ................................................................................ |
| Class4f-CsYUC3-255980 | ................................................................................ |
| Class4f-MtYUC9-Medtr1g069275 | ................................................................................ |
| Class4f-MtYUC8-Medtr7g099330 | ................................................................................ |
| Class4f-MtYUC3-Medtr1g046230 | ................................................................................ |
| Class4f-PpYUC5-8G211000 | ................................................................................ |
| Class4f-PpYUC3-PrupeG054300 | ................................................................................ |
| Class4f-BrYUC9-I05505 | ................................................................................ |
| Class4f-BrYUC5-B02765 | ................................................................................ |
| Class4f-BrYUC8-A00847 | ................................................................................ |
| Class4f-BrYUC9-J00235 | ................................................................................ |
| Class4f-BrYUC3X2-J00274 | ..MHRN.......................................................................... |
| Class4f-BrYUC8-H01467 | ................................................................................ |
| Class4f-BrYUC3-I05483 | ..MYRN.......................................................................... |
| Class4f-BrYUC5-F03781 | ................................................................................ |
| Class4f-BrYUC7-E01069 | ..MCTN.......................................................................... |
| Class4f-PCaYUCD-KAI3983187 | ................................................................................ |
| Class4f-PCaYUCF-KAI3990778 | ................................................................................ |
| Class4f-PjYUC2-QIH44904 | ................................................................................ |
| Class4f-PjYUC3-QIH44905 | ................................................................................ |
| Class4f-PjYUC4-QIH44906 | ................................................................................ |
| Class4f-VvYUC8-207s0005g04800 | ................................................................................ |
| Class4f-VvYUC3-205s0051g00060 | ................................................................................ |
| Class4f-DcYUC3-030776 | ................................................................................ |
| Class4f-DcYUC3-027361 | ................................................................................ |
| Class4f-HqYUC3-0G114700 | ................................................................................ |
| Class4f-CaYUC3-315.833 | ................................................................................ |
| Class4f-FvYUC7-AFG16917 | ................................................................................ |
| Class4f-CpYUC3-59.54 | ................................................................................ |
| Class4f-PtYUC3-008G174600 | ................................................................................ |
| Class4f-PtYUC7-010G062400.2 | ..MAMTL...PYLTPSATSFSPTPFTSNKLHFILLSTLRPIHLSTQTSSLLVSSFGRQHPRHSFLLSLIFYFPLWRLTTF |
| Class4f-PtYUC8-002G254200 | ................................................................................ |
| Class4f-DcYUC5-000550 | ................................................................................ |
| Class4f-DcYUC5-012617 | ................................................................................ |
| Class4f-DcYUC5A-000561 | ................................................................................ |
| Class4f-DcYUC5B-000562 | ................................................................................ |
| Class4f-DcYUC5C-000563 | ................................................................................ |
| Class4f-DcYUC5D-000560 | ................................................................................ |
| Class4f-CaYUC3-2016.447 | ................................................................................ |
| Class4f-CaYUC8-635.212 | ................................................................................ |
| Class4f-CpYUC8-87.71 | ................................................................................ |
| Class4f-FvYUC3-AFG16918 | ................................................................................ |
| Class4f-HqYUC5-04G157000 | ................................................................................ |

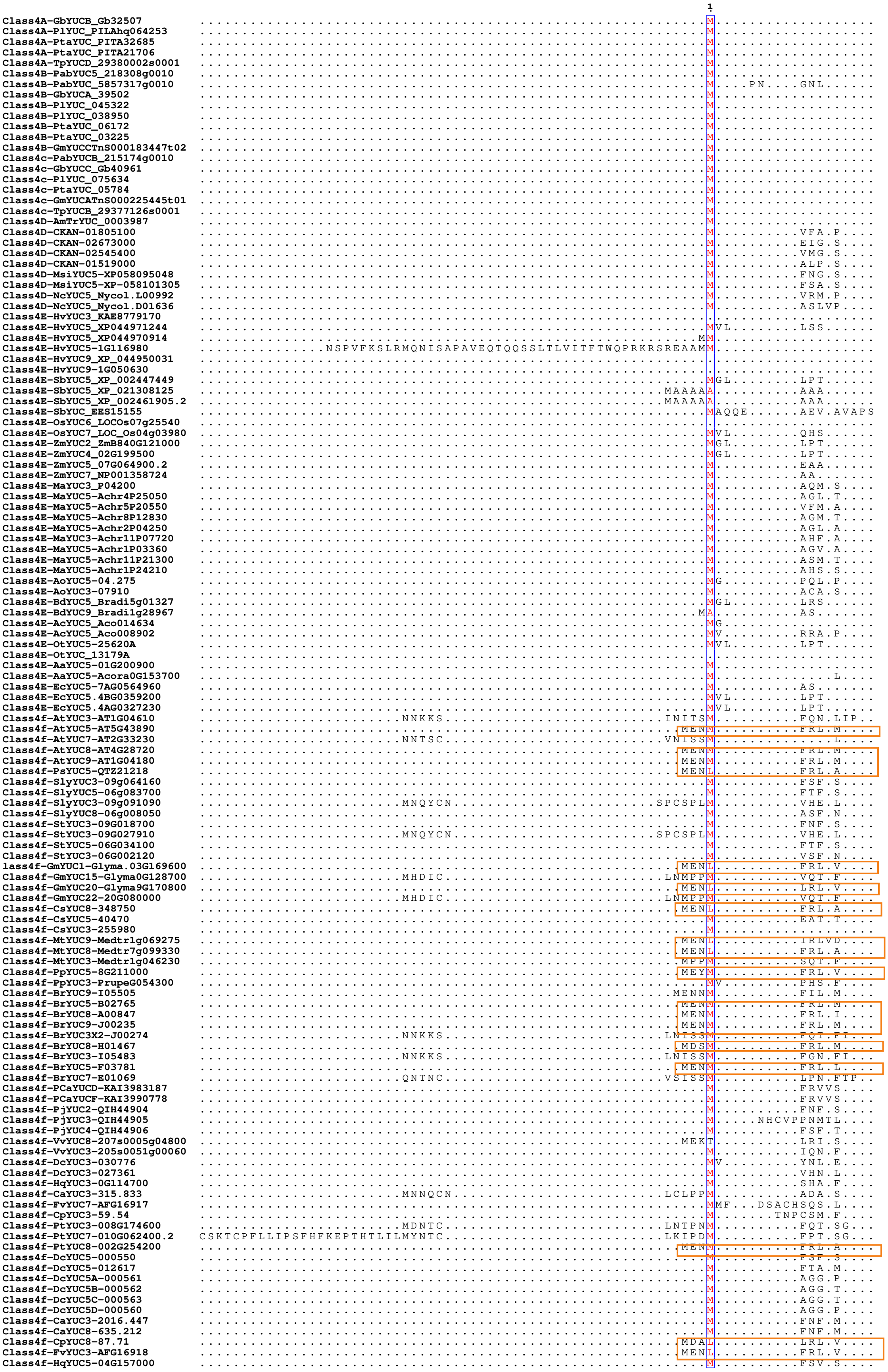

N-terminal-MENMFRLXDHED---

### Class-4 YUCCAs

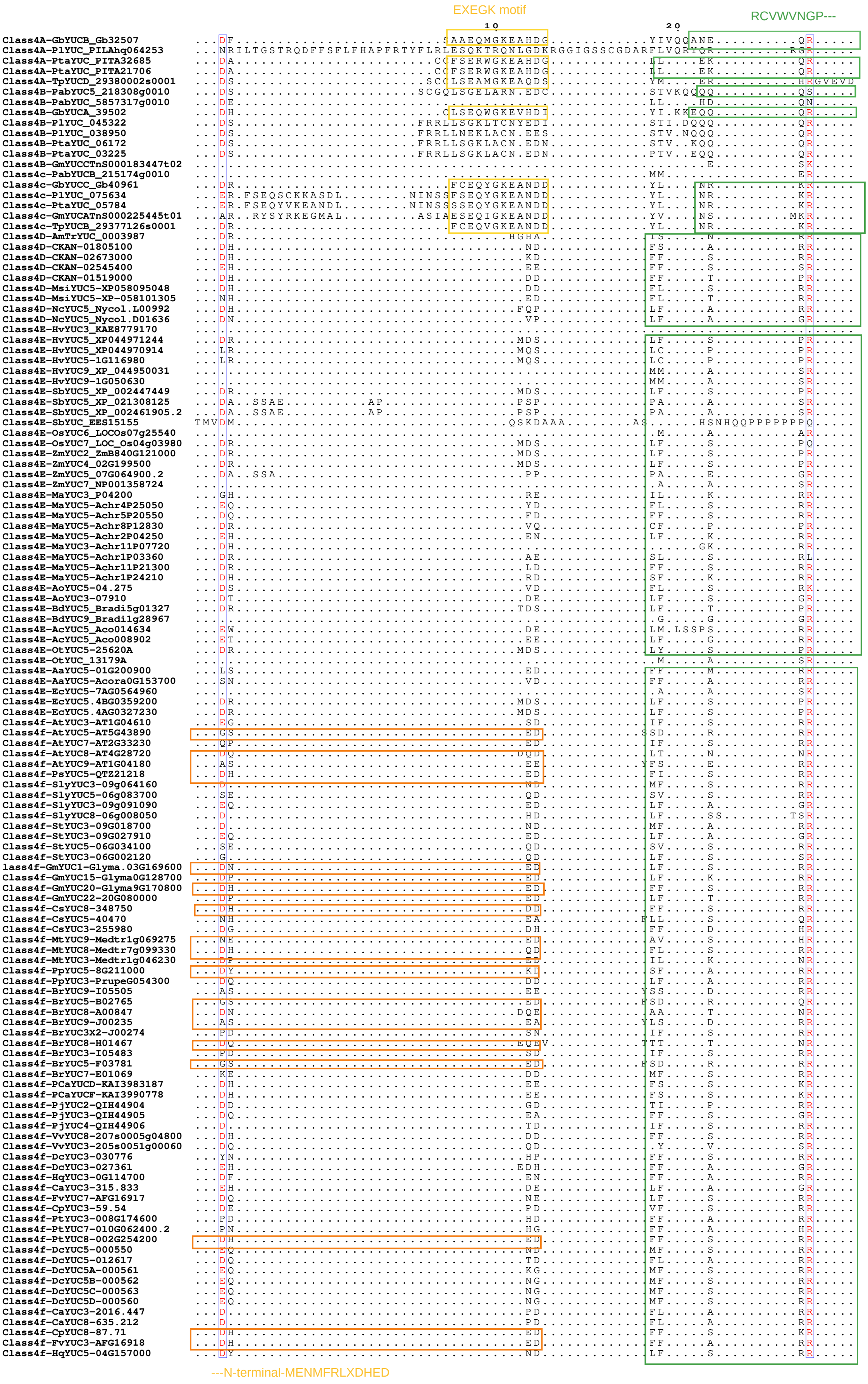

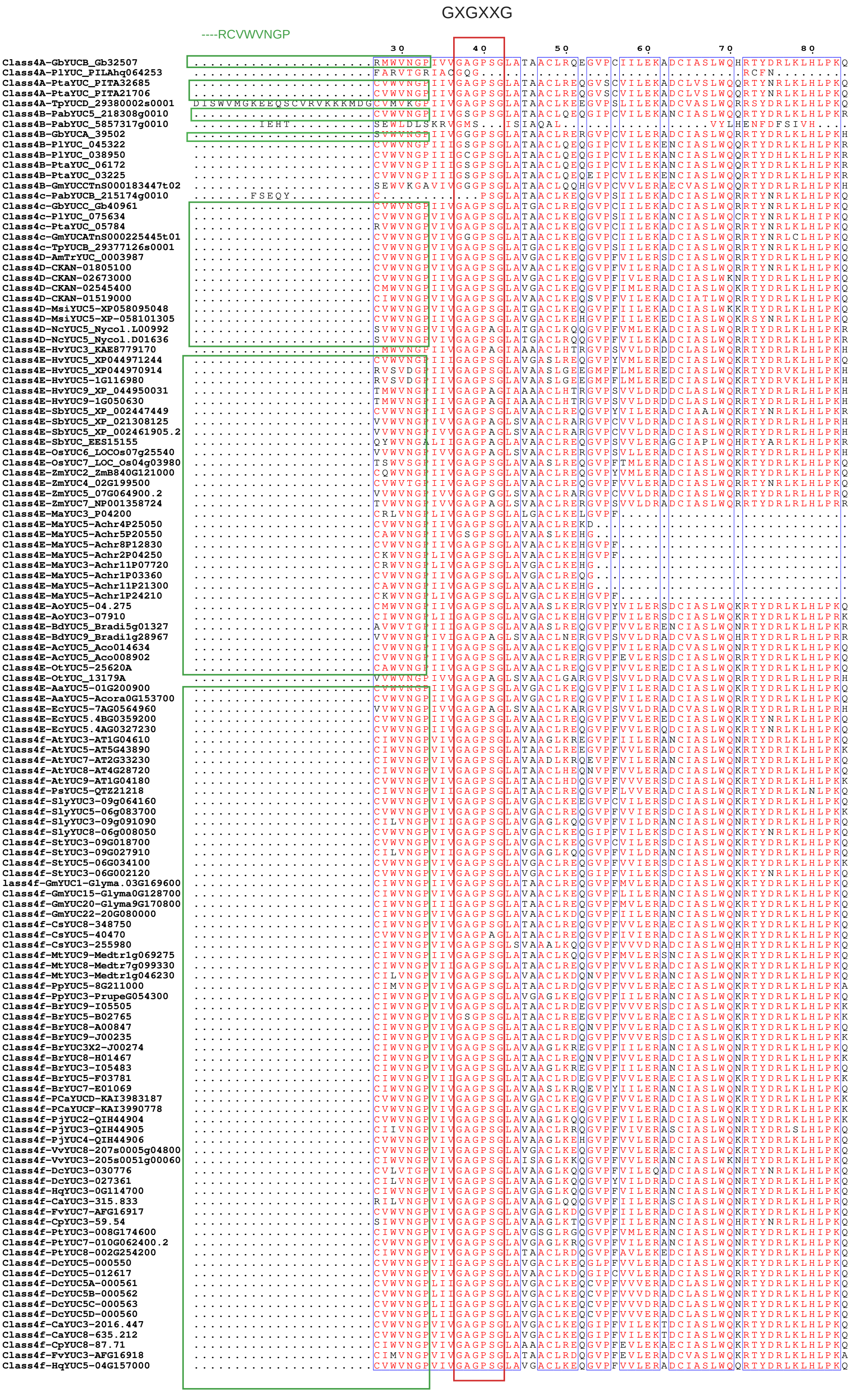
GXGXXG
----RCVWVNGP
30
40
50
60
70
80

### Class-4 YUCCAs

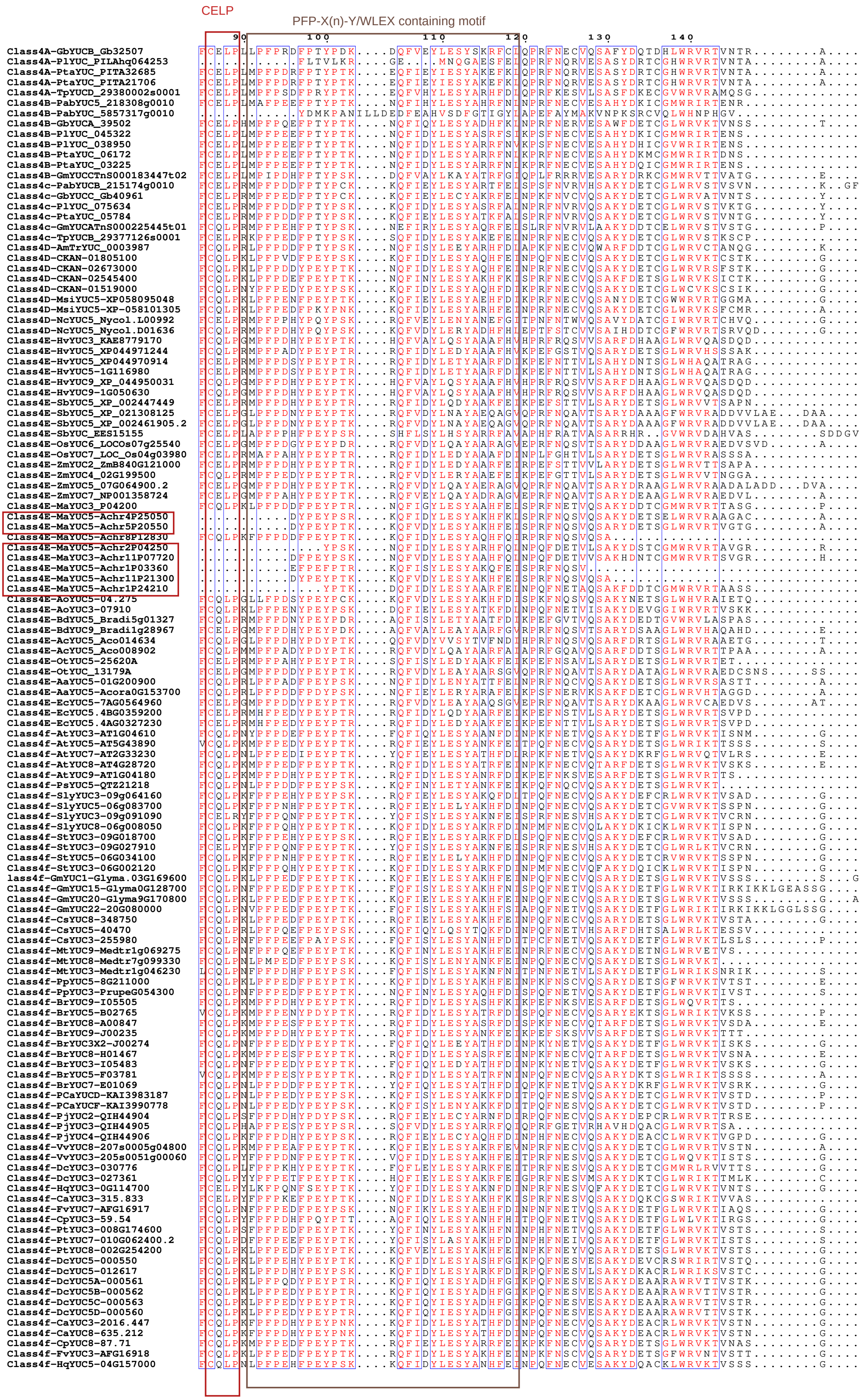

#### Class-4 YUCCAs

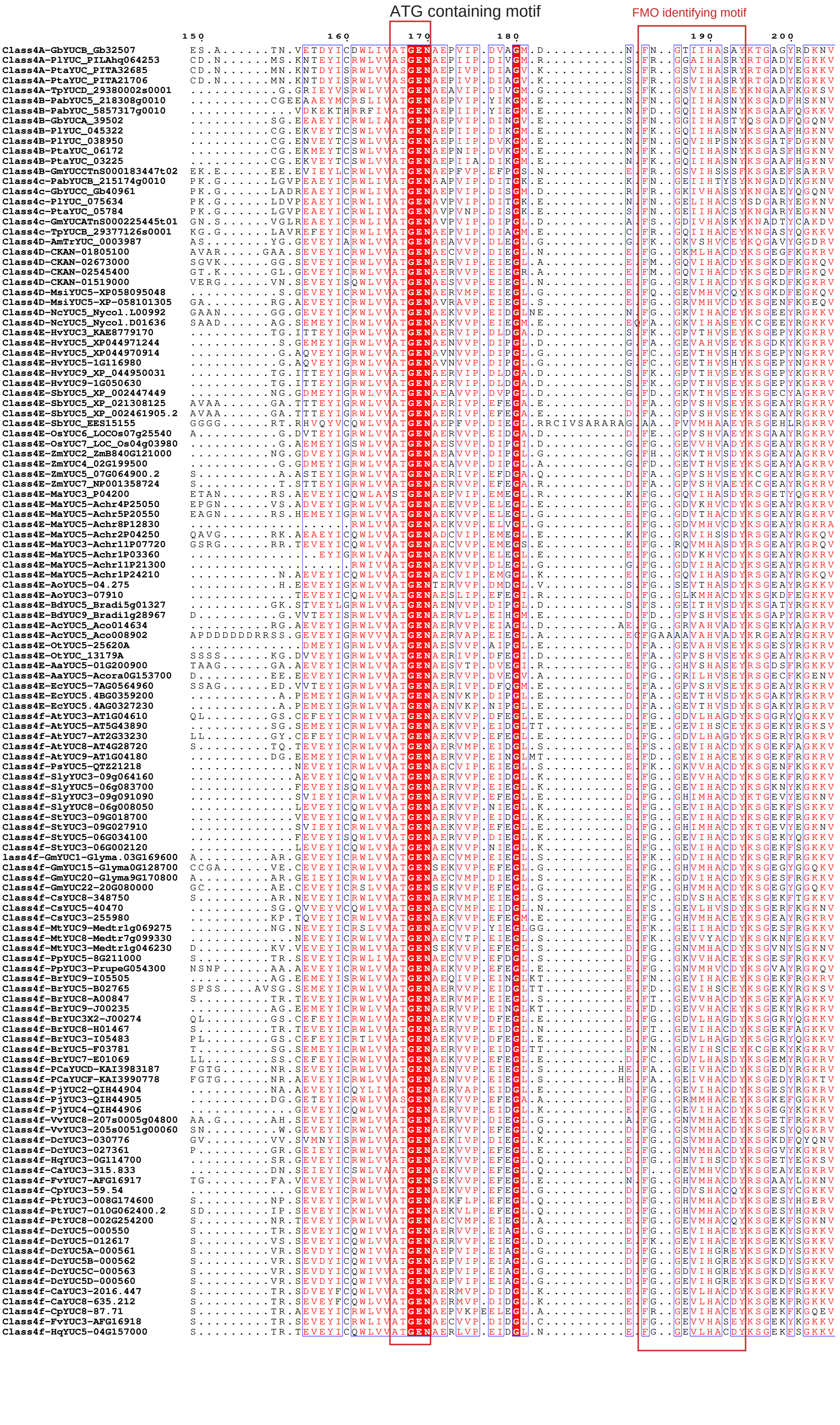

### Class-4 YUCCAs

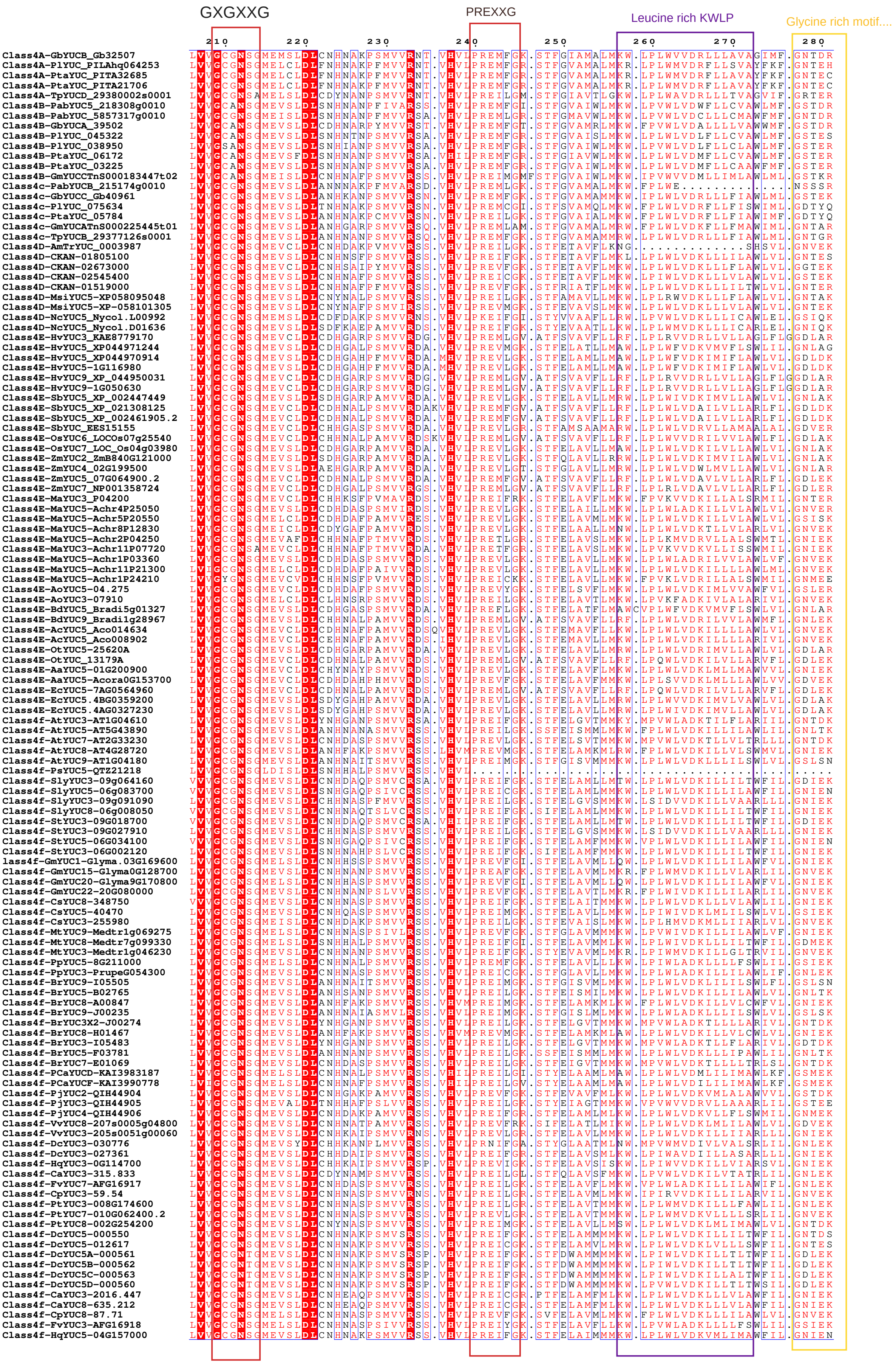

#### Class-4 YUCCAs

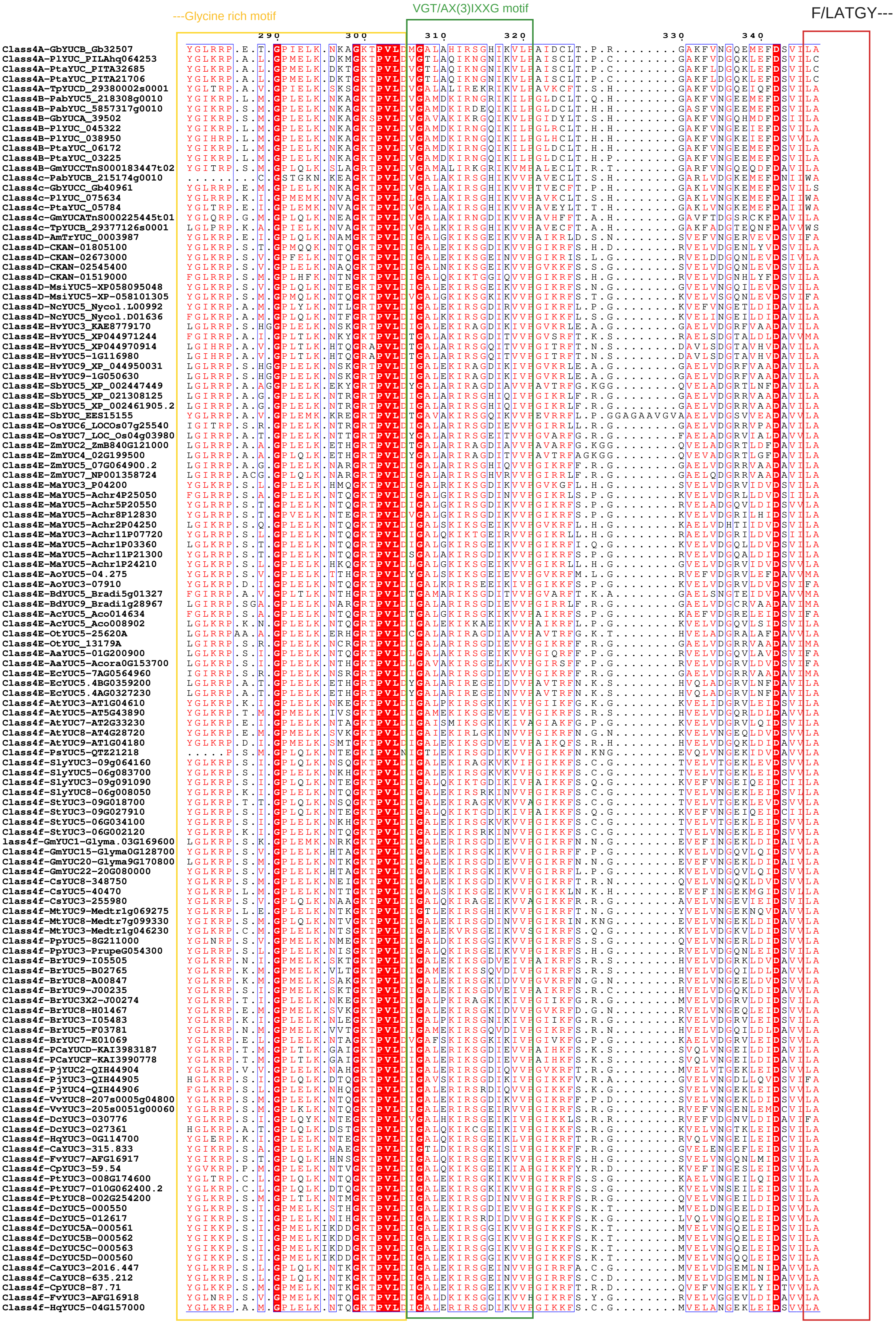

#### Class-4 YUCCAs

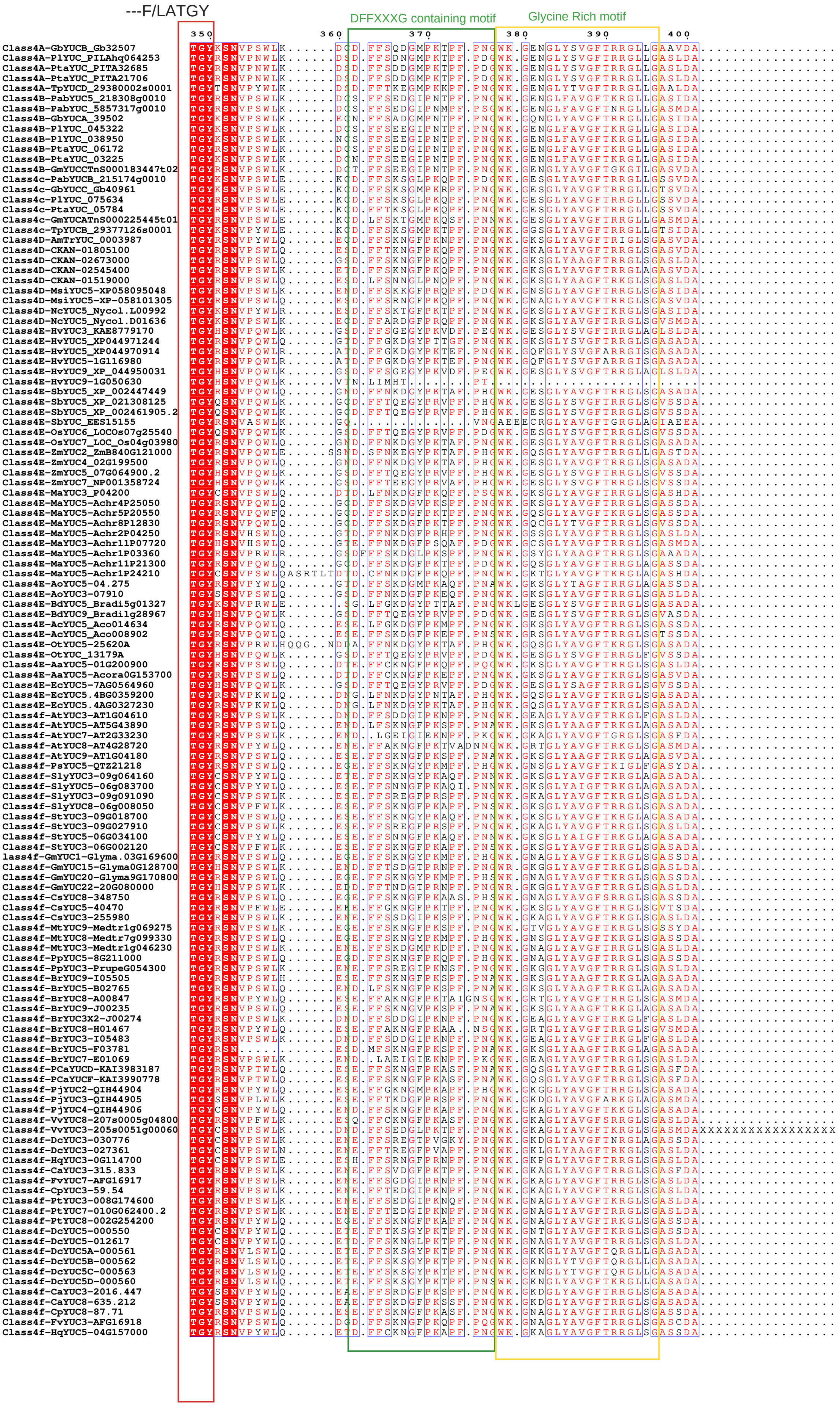

#### Class-4 YUCCAs

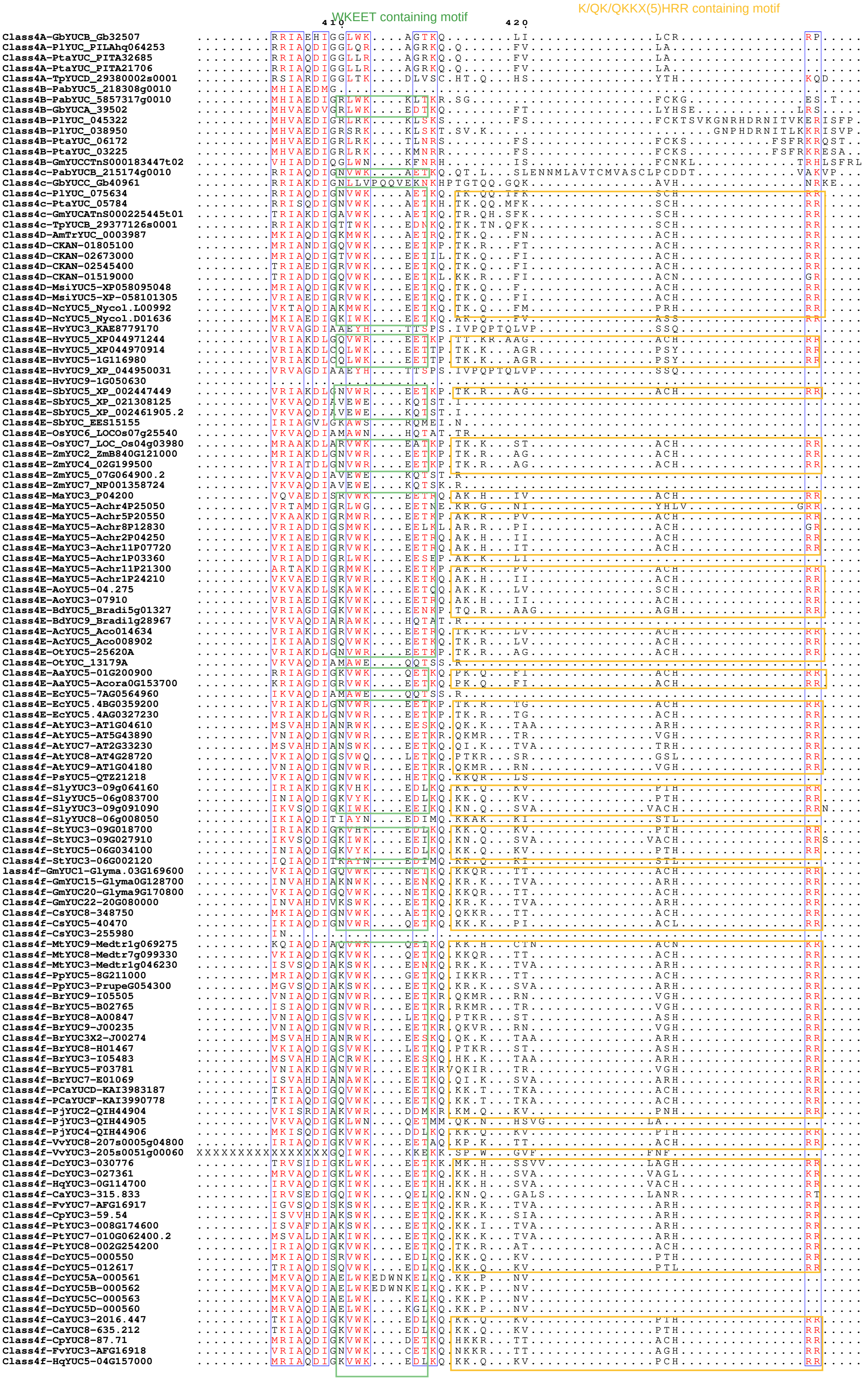

```
Class4A-GbYUCB_Gb32507              ....................................................
Class4A-PlYUC_PILAhq064253          ....................................................
Class4A-PtaYUC_PITA32685            ....................................................
Class4A-PtaYUC_PITA21706            ....................................................
Class4A-TpYUCD_29380002s0001        ....................................................
Class4B-PabYUC5_218308g0010         ....................................................
Class4B-PabYUC_5857317g0010         ....................................................
Class4B-GbYUCA_39502                .............DQNP...................................
Class4B-PlYUC_045322                .............PSNYSVLSGT.............................
Class4B-PlYUC_038950                .............PNNYFVLSRT.................QLNE........
Class4B-PtaYUC_06172                ....................................................
Class4B-PtaYUC_03225                ....................................................
Class4B-GmYUCCTnS000183447t02       IRTSSSSSLSSSTPSTPQSLSLS..PRFRSSAPRSAPSS.SLFS........
Class4c-PabYUCB_215174g0010         ..................CYKTLLGPIAGIENP..GPRNERLEHNKPLGRVA
Class4c-GbYUCC_Gb40961              .EGLKSLLLSGG.....LCSDKEFCK.........................A
Class4c-PlYUC_075634                ..................CISQF.............................
Class4c-PtaYUC_05784                ..................CSSQF.............................
Class4c-GmYUCATnS000225445t01       ..................CISQF.............................
Class4c-TpYUCB_29377126s0001        ..................CISQF.............................
Class4D-AmTrYUC_0003987             ..................CISQF.............................
Class4D-CKAN-01805100               ..................CISQF.............................
Class4D-CKAN-02673000               ..................CISQF.............................
Class4D-CKAN-02545400               ..................CTSQF.............................
Class4D-CKAN-01519000               ..................FISQF.............................
Class4D-MsiYUC5-XP058095048         ..................CISMF.............................
Class4D-MsiYUC5-XP-058101305        ..................CISQF.............................
Class4D-NcYUC5_Nycol.L00992         ..................CISQF.............................
Class4D-NcYUC5_Nycol.D01636         ..................RISQM.............................
Class4E-HvYUC3_KAE8779170           ..................SQAET.............................
Class4E-HvYUC5_XP044971244          ..................CISVIF............................
Class4E-HvYUC5_XP044970914          ..................SISVVL............................
Class4E-HvYUC5-1G116980             ..................SISVVL............................
Class4E-HvYUC9_XP_044950031         ..................SQAET.............................
Class4E-HvYUC9-1G050630             ....................................................
Class4E-SbYUC5_XP_002447449         ..................CISVVF............................
Class4E-SbYUC5_XP_021308125         ....................................................
Class4E-SbYUC5_XP_002461905.2       ....................................................
Class4E-SbYUC_EES15155              ....................................................
Class4E-OsYUC6_LOCOs07g25540        ....................................................
Class4E-OsYUC7_LOC_Os04g03980       ..................CISVIF............................
Class4E-ZmYUC2_ZmB840G121000        ..................CISVVF............................
Class4E-ZmYUC4_02G199500            ..................CISVVF............................
Class4E-ZmYUC5_07G064900.2          ....................................................
Class4E-ZmYUC7_NP001358724          ....................................................
Class4E-MaYUC3_P04200               ..................CISQI.............................
Class4E-MaYUC5-Achr4P25050          ..................T.................................
Class4E-MaYUC5-Achr5P20550          ..................CISQI.............................
Class4E-MaYUC5-Achr8P12830          ..................CISQI.............................
Class4E-MaYUC5-Achr2P04250          ..................CTSQN.............................
Class4E-MaYUC3-Achr11P07720         ..................CTSQN.............................
Class4E-MaYUC5-Achr1P03360          ..................CLP...............................
Class4E-MaYUC5-Achr11P21300         ..................CISQI.............................
Class4E-MaYUC5-Achr1P24210          ..................CISQI.............................
Class4E-AoYUC5-04.275               ..................CISQIY............................
Class4E-AoYUC3-07910                ..................CMSQM.............................
Class4E-BdYUC5_Bradi5g01327         ..................SISVIF............................
Class4E-BdYUC9_Bradi1g28967         ....................................................
Class4E-AcYUC5_Aco014634            ..................CISQI.............................
Class4E-AcYUC5_Aco008902            ..................CISQI.............................
Class4E-OtYUC5-25620A               ..................CISVVF............................
Class4E-OtYUC_13179A                ....................................................
Class4E-AaYUC5-01G200900            ..................CISQI.............................
Class4E-AaYUC5-Acora0G153700        ..................CISQI.............................
Class4E-EcYUC5-7AG0564960           ....................................................
Class4E-EcYUC5.4BG0359200           ..................CISVVF............................
Class4E-EcYUC5.4AG0327230           ..................CISVVF............................
Class4f-AtYUC3-AT1G04610            ..................CISHF.............................
Class4f-AtYUC5-AT5G43890            ..................CISVA.............................
Class4f-AtYUC7-AT2G33230            ..................CISHF.............................
Class4f-AtYUC8-AT4G28720            ..................CISQQF............................
Class4f-AtYUC9-AT1G04180            ..................CISVA.............................
Class4f-PsYUC5-QTZ21218             ....................................................
Class4f-SlyYUC3-09g064160           ..................CISTF.............................
Class4f-SlyYUC5-06g083700           ..................CISTF.............................
Class4f-SlyYUC3-09g091090           ..................KLSSF.............................
Class4f-SlyYUC8-06g008050           ....................................................
Class4f-StYUC3-09G018700            ..................CISTF.............................
Class4f-StYUC3-09G027910            ..................KLPSF.............................
Class4f-StYUC5-06G034100            ..................CISTF.............................
Class4f-StYUC3-06G002120            ....................................................
lass4f-GmYUC1-Glyma.03G169600       ..................CISQF.............................
Class4f-GmYUC15-Glyma0G128700       ..................CISHF.............................
Class4f-GmYUC20-Glyma9G170800       ..................CISQF.............................
Class4f-GmYUC22-20G080000           ..................CISHF.............................
Class4f-CsYUC8-348750               ..................CISQF.............................
Class4f-CsYUC5-40470                ..................CISHF.............................
Class4f-CsYUC3-255980               ....................................................
Class4f-MtYUC9-Medtr1g069275        ..................CISQF.............................
Class4f-MtYUC8-Medtr7g099330        ..................CISQF.............................
Class4f-MtYUC3-Medtr1g046230        ..................CISHF.............................
Class4f-PpYUC5-8G211000             ..................CISQF.............................
Class4f-PpYUC3-PrupeG054300         ..................CISHF.............................
Class4f-BrYUC9-I05505               ..................CISVA.............................
Class4f-BrYUC5-B02765               ..................CISVA.............................
Class4f-BrYUC8-A00847               ..................CISQQF............................
Class4f-BrYUC9-J00235               ..................CISVA.............................
Class4f-BrYUC3X2-J00274             ..................CISHF.............................
Class4f-BrYUC8-H01467               ..................CISQQF............................
Class4f-BrYUC3-I05483               ..................CISHF.............................
Class4f-BrYUC5-F03781               ..................CISIA.............................
Class4f-BrYUC7-E01069               ..................CISQF.............................
Class4f-PCaYUCD-KAI3983187          ..................CISQF.............................
Class4f-PCaYUCF-KAI3990778          ..................CISQF.............................
Class4f-PjYUC2-QIH44904             ..................CISTF.............................
Class4f-PjYUC3-QIH44905             ....................................................
Class4f-PjYUC4-QIH44906             ..................CISTF.............................
Class4f-VvYUC8-207s0005g04800       ..................CISQF.............................
Class4f-VvYUC3-205s0051g00060       ....................................................
Class4f-DcYUC3-030776               ..................CKSHV.............................
Class4f-DcYUC3-027361               ..................YKSQF.............................
Class4f-HqYUC3-0G114700             ..................SKSHC.............................
Class4f-CaYUC3-315.833              ..................CKS...............................
Class4f-FvYUC7-AFG16917             ..................CISHF.............................
Class4f-CpYUC3-59.54                ..................CISHF.............................
Class4f-PtYUC3-008G174600           ..................CISHF.............................
Class4f-PtYUC7-010G062400.2         ..................CISHF.............................
Class4f-PtYUC8-002G254200           ..................CISQF.............................
Class4f-DcYUC5-000550               ..................CISTF.............................
Class4f-DcYUC5-012617               ..................CISTF.............................
Class4f-DcYUC5A-000561              ....................................................
Class4f-DcYUC5B-000562              ....................................................
Class4f-DcYUC5C-000563              ....................................................
Class4f-DcYUC5D-000560              ....................................................
Class4f-CaYUC3-2016.447             ..................CISQF.............................
Class4f-CaYUC8-635.212              ..................CISQF.............................
Class4f-CpYUC8-87.71                ..................CISQQF............................
Class4f-FvYUC3-AFG16918             ..................CISQTF............................
Class4f-HqYUC5-04G157000            ..................CISQF.............................
```

### CLASS 5 YUCCAS

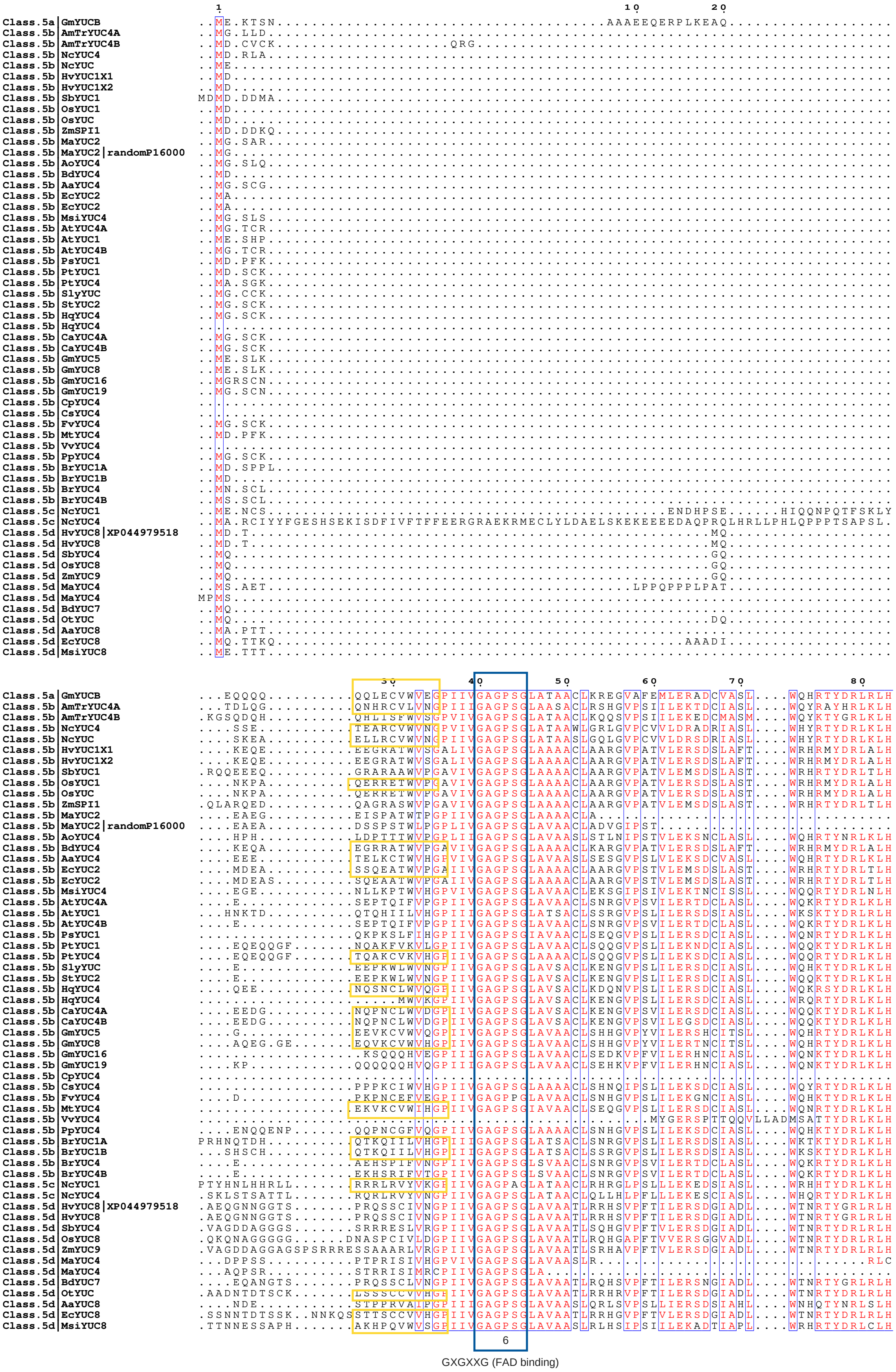

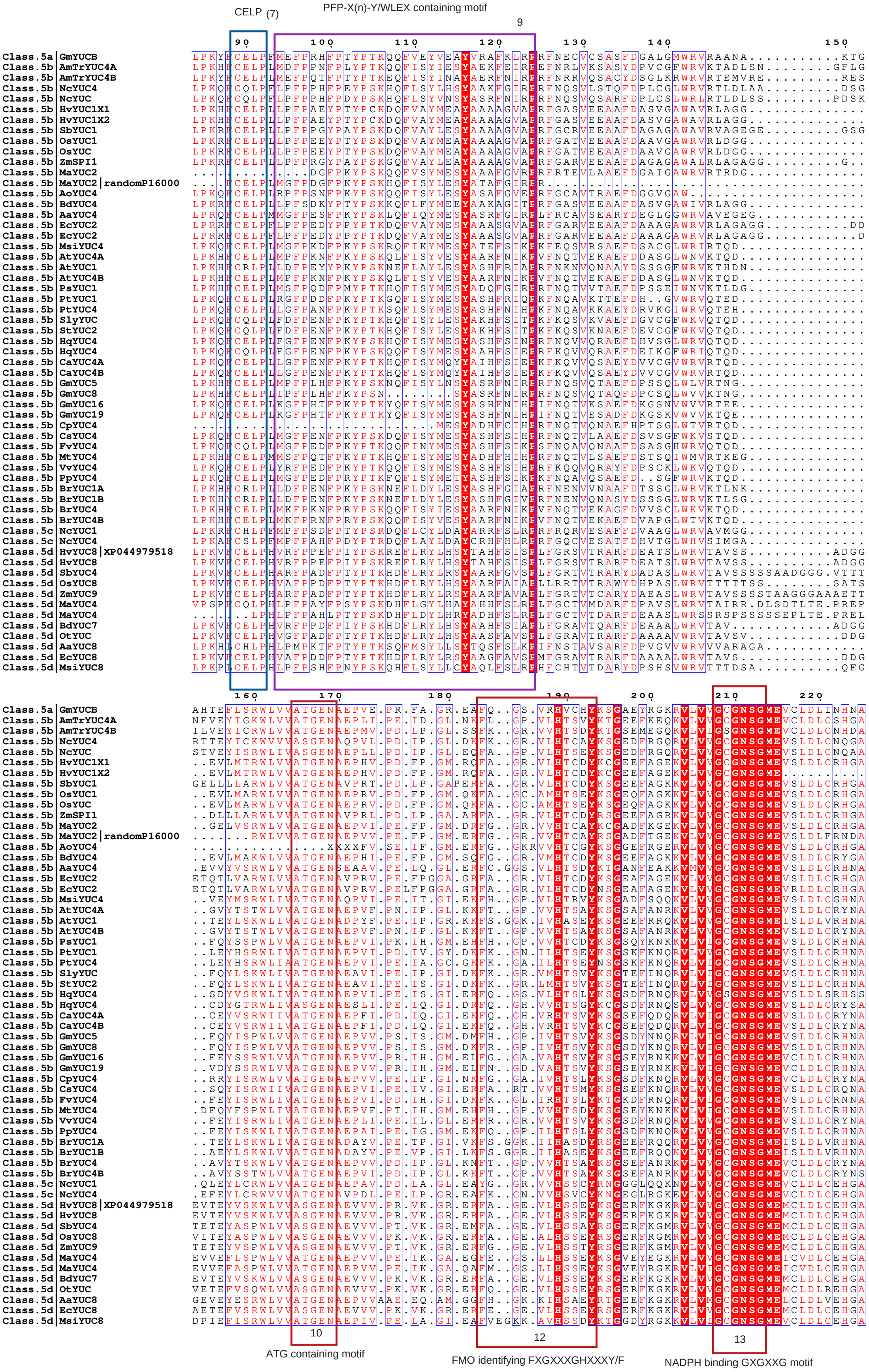
CELP (7)
PFP-X(n)-Y/WLEX containing motif
9
10
12
13
ATG containing motif
FMO identifying FXGXXXGHXXXY/F
NADPH binding GXGXXG motif

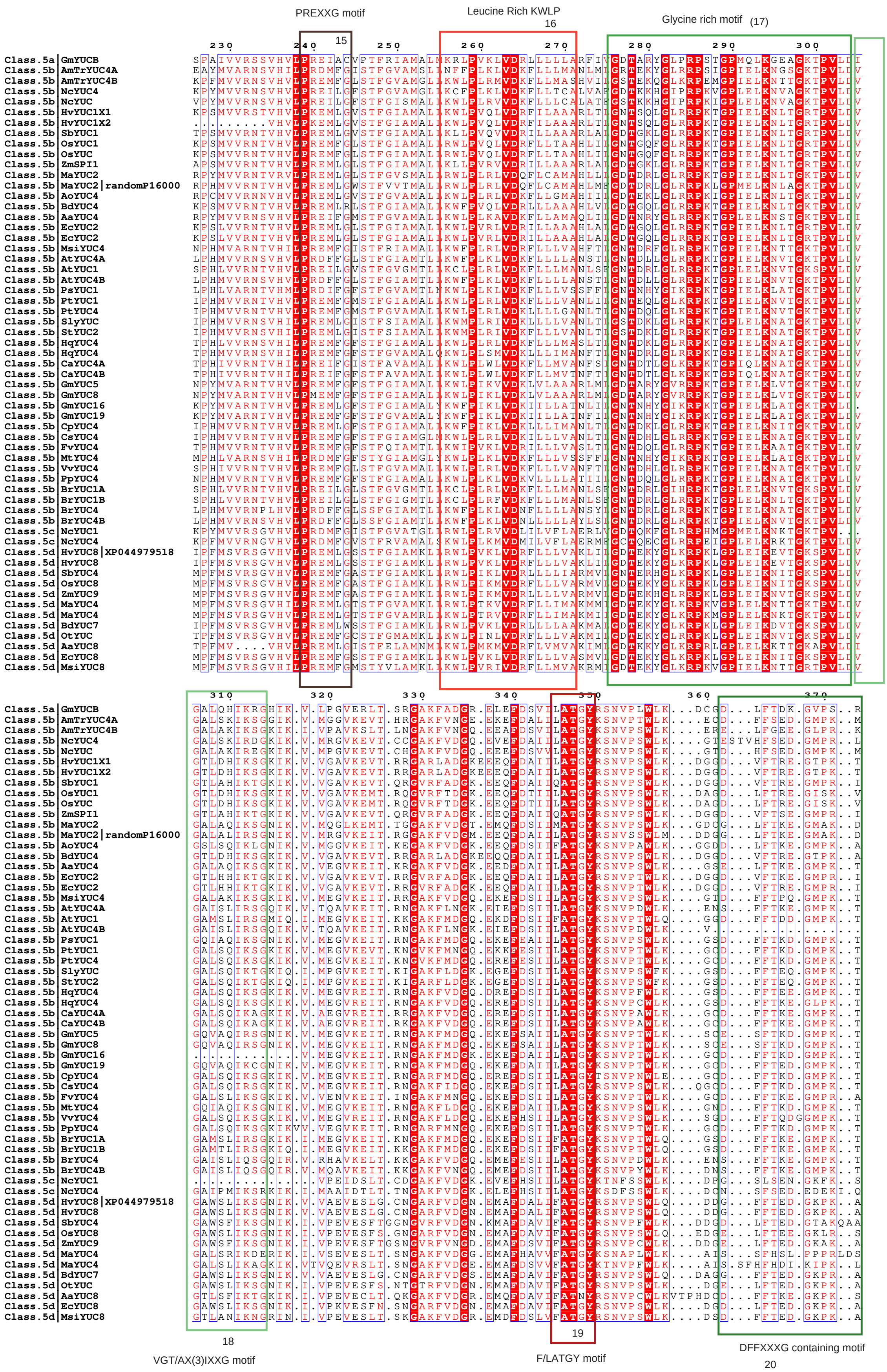
PREXXG motif
15
Leucine Rich KWLP
16
Glycine rich motif (17)
18
VGT/AX(3)IXXG motif
19
F/LATGY motif
DFFXXXG containing motif
20

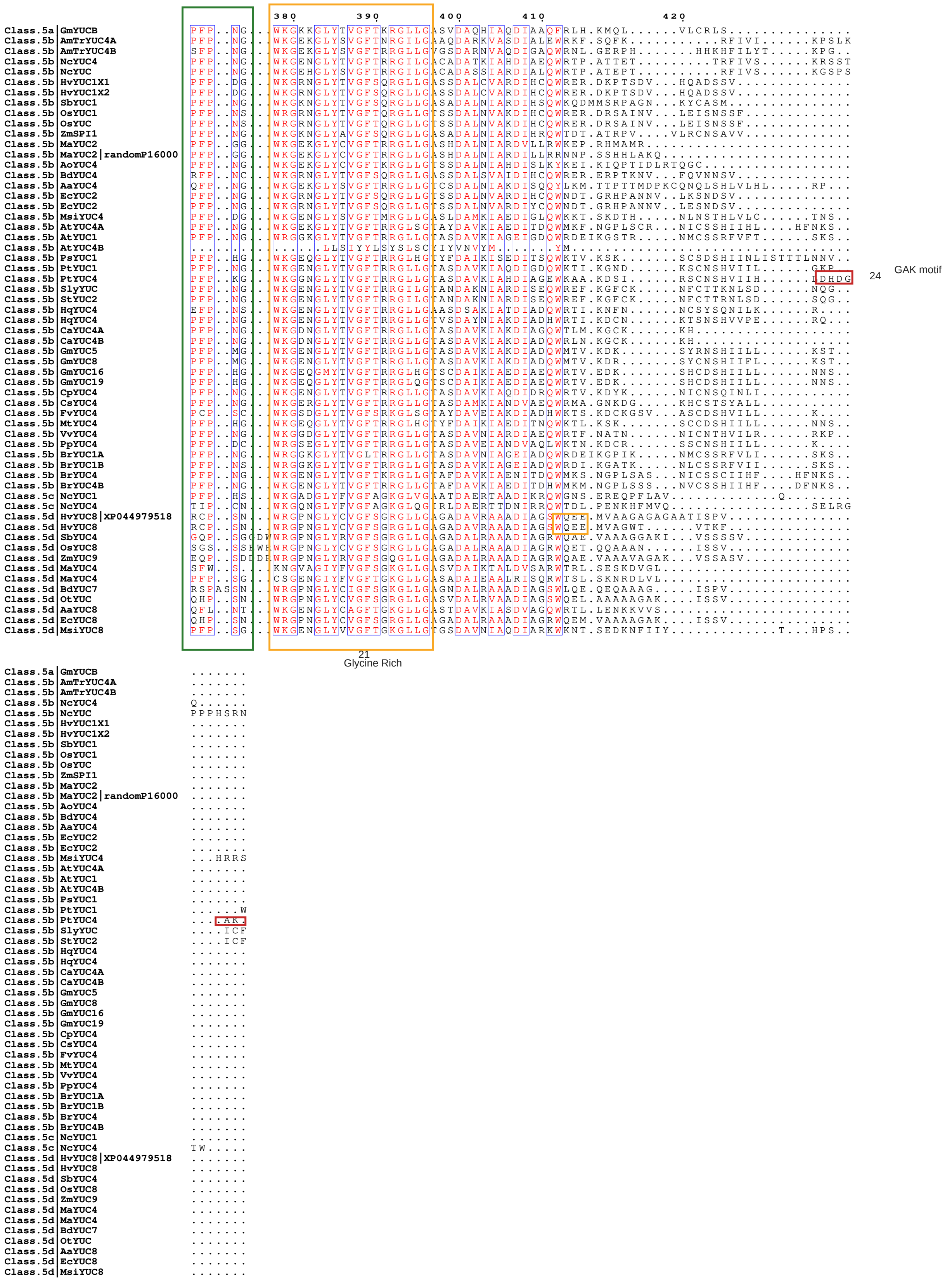
380
390
400
410
420
GAK motif
24
21
Glycine Rich

### Sequence alignment was done in MAFFT version 7 (https://mafft.cbrc.jp/alignment/server/)
#### Visualized in ESPript 3.0 (https://espript.ibcp.fr/ESPript/cgi-bin/ESPript.cgi).

(Sister YUCCAs (sYUCs) retrieved based on the paper Carrillo-Carrasco et al, 2023 (https://doi.org/10.15252/embj.2022113018).

```
                                             1                        10        20        30          40
SisterYUCCA|Mapoly0052s0062       ............MASNGTL................ENGSSDNSKIVKAWLSYLDEALQK..QDIAHLVTLF.....DE
SisterYUCCA|Mapoly0122s0026       ..................................MGSEELSAGQKITSWLSEFDTVLQD..EDVDQILELF.....DD
SisterYUCCA|Sm113792              ............MAT..................VNGSPKIDPDALVDAWLARLDSSLAS..GDASAAANLF.....EE
SisterYUCCA|Sm422043              ............MAT..................MNGSPKIDPDALVDAWLTRLDSSLAS..GDPSAAANLF.....EE
SisterYUCCA|kfl00109_0340         ....................................MATQTLTEDVRAWLERFSAALEE..SDLEKVLALF......G
4265-Phaeomegaceros_coriaceus     .................................MGKLEAEPERSRAKAWLLELERTLVA..EDIAGTLELF......G
W-AKXB-2007730-Phaeomegaceros     ..............................................................................
W-AKXB-2007731-Phaeomegaceros     ..............................................................................
W-AKXB-2067423-Phaeomegaceros     .......................................MAATEAVKAWLSSFEDALHK..QDVSAAVELF.....QE
BMJR-2002471-Adiantum_tenerum     ..............................................................................
BMJR-2059933-Adiantum_tenerum     ..............................................................................
BMJR-2062494-Adiantum_tenerum     ..............................................................................
U-2019103-Radula_lindenbergia     ..............................................................................
U-2086103-Radula_lindenbergia     ...........MLAAQPQPHDISLHLFNKTQPQVTVKENLPPREIAQQWLDKFAAALSGNGNDLSQLPSLI......H
CA|HW-DXOU-2035997-Nothoceros     ..............................................................................
FAJB-2001596-Paraphymatoceros     ..............................................................................
FAJB-2004773-Paraphymatoceros     .......................................MTATDAARAWLADFSDALQR..QDVNAVVELF.....QE
FAJB-2008542-Paraphymatoceros     ..............................................................................
CA|GA-FPCO-2029146-Interfilum     ....................................MANPSLTEAVSAWLENFGSALSE..GGIESALSLF......G
GA-FQLP-2033521-Klebsormidium     ....................................MASPTLTEDVSLWLGSFGAALQE..NDLEKALSLF......G
YUCCA|LM-GANB-2011315-Cyathea     ..............................................................................
YUCCA|LM-GANB-2011316-Cyathea     ..............................................................................
YUCCA|LM-GANB-2017253-Cyathea     ..............................................................................
ANB-2070212-Cyathea_spinulosa     .........................................MAERVNAWLQSLAFALEK..RDLIATAELF.....RE
LW-HERT-2045223-Sphaerocarpos     ............M......................GSLELSGREVVQAWLADFDDALQK..QDIPQLLGLF.....QE
UCCA|RA-IEHF-2003087-Dumontia     ..............................................................................
O-2067228-Marchantia_paleacea     ............MASNGSM................VNGSSDNTKVVKAWLLDFDEALQK..QDTAHLVTLF.....DD
O-2067575-Marchantia_paleacea     ..................................MGSEELSAGQKITSWLSEFDTALQN..EDVDQVMKLF.....GD
-2046311-Conocephalum_conicum     ..................................MG..EISNSQKITAWLCEFDAALQQEEEDLDKVMELF.....AA
-2046658-Conocephalum_conicum     ............MASNGVSNG..........ELLKTNGASDNTETVKAWLSEFDEAIQK..QDIQQLLILF.....DE
-2181954-Selaginella_wallacei     ............MPASP..............GQCTQAMAEQANAEIDAWLSRLDAALKS..RKLDEAVDLFAADDDDD
A|LP-KUXM-2009452-Selaginella     ..........................................MAEVDEWLRDFDGAMTI..RDIEGAVDLF......N
|LM-PIVW-2024618-Ceratopteris     .........................................MAERINAWLREFDSVLQN..EDIPAAVLLF.....QE
2005240-Isoetes_tegetiformans     ARFRTG.LELPAMAIDAS.............NGSFNSTADPSNAKVDNFLSDLDHYLQI..RDVPKIIDLFG....DG
2005241-Isoetes_tegetiformans     ..............................................................................
2008401-Hymenophyllum_bivalve     .........................................MAERIRAWLKSLDCALQK..GDVSAAVELF.....HG
003161-Megaceros_vincentianus     ............................................................AEDVAGTLELF......G
007840-Megaceros_vincentianus     .....................................MAAGLNGTVAAWLSRFEEALQK..EDVSQAVELF.....LE
013674-Megaceros_vincentianus     ..............................................................................
016968-Megaceros_vincentianus     .......................................MAATEAVKAWLSSFENALHK..QDVTALLEHF.....QE
019507-Megaceros_vincentianus     ..............................................................................
2000839-Marchantia_emarginata     ..............................................................................
2004520-Marchantia_emarginata     ..................................MGSEEMSASQKITAWLSEFDTALQE..EEVDKVMKLF.....LD
2069965-Marchantia_emarginata     ..............................................................................
2074864-Marchantia_emarginata     ..............................................................................
2050988-Anthoceros_agrestis-A     ........................................MMAEAVRAWLSEFSDGLQK..QDVCAVVELF.....QE
VB-2006165-Lunularia_cruciata     ..................................MSTISSKTREEVSAWLSQFNEALLE..HDVPGVLNLF.....HD
VB-2006166-Lunularia_cruciata     ............MSRLTQS..............MNSTLSTGGTEELATWLSQFKEAIVK..YDISGVLDLF.....AD
VB-2006167-Lunularia_cruciata     ..............................................................................
VB-2006168-Lunularia_cruciata     ..............................................................................
VB-2007869-Lunularia_cruciata     ............METNGSVELASEVVNENGSSQVVKSNGVSDTEVVKAWLSELQDTLEK..QDIVHLVTLF.....LE
VB-2075591-Lunularia_cruciata     .....................................MGTRAEDQITAWLATFDTALQG..GDVEHVLELF.....AP
CRN-2050303-Megaceros_tosanus     ..............................................................................
CRN-2052782-Megaceros_tosanus     .....................................MAAEASAIAAAWLSRFEEALHK..EEVSQAVELF.....VE
VITX-2002371-Blechnum_spicant     ..............................................................................
VITX-2009142-Blechnum_spicant     .......SLSLSLVTQRATE....................RMAGRISAWLQSFDSALKS..RDVSAATELF.....CP
VITX-2009143-Blechnum_spicant     S..........LSHT.......................HAHMAGRITAWLESLDSALQK..KDISSAAHLF.....YE
VITX-2016791-Blechnum_spicant     .....................................MASAEMMRVQSWLQSFDAALQH..GDVDAVVDMF.....DE
VITX-2098911-Blechnum_spicant     .........................................MAERVRVWLQSLECALNE..RDVDAAAELF.....HD
VITX-2099259-Blechnum_spicant     .........................................MAERVKVWLQSFECALQA..QDVRAAAELF.....HE
CA|HW-WCZB-2015184-Phaeoceros     ..............................................................................
CA|HW-WCZB-2016004-Phaeoceros     .......................................MAATDSVRAWLADFSDALQK..QDVNAVVELF.....QE
CA|HW-WCZB-2117013-Phaeoceros     .......MHL.LLEYCVRIHR..........LGQGISEDMAANDLVKAWLANFQDALQN..QDVRAVVELF.....RE
CA|HW-WCZB-2118293-Phaeoceros     ..............................................................................
U-2004486-Leucostegia_immersa     ..............................................................................
U-2004559-Leucostegia_immersa     ..............................................................................
JLO-2000942-Riccia_berychiana     ..............................................................................
JLO-2002497-Riccia_berychiana     ............MEAAPLLQH..............LDQQQGGGEIVQAWLSEFDEALRN..QDVAHLVTLF.....DE
JLO-2033265-Riccia_berychiana     .....................................EELNASQKISSWLFEFDAALQK..EDVDQAMELF.....VD
|LM-WQML-2003640-Cryptogramma     ............THTHKQTE....................TMANRISAWLQNLEWALKQ..RDIVATTHLF.....QE
|LM-WQML-2004411-Cryptogramma     ..............................................................................
|LM-WQML-2067992-Cryptogramma     ..............................................................................
2095928-Ophioglossum_vulgatum     ..............................................................................
2024024-Polysiphonia_japonica     ............MADETNG...............TNEANESSEKTITDWIQHFSDALIG..DDIRLVVDLF......H
-2100101-Pallavicinia_lyellii     ..............................................................................
JJY-2002336-Woodsia_scopulina     ..............................................................................
-2008616-Woodsia_scopulina.p2     ...............KERER....................EMAERVRVWLQSLECALNE..RDVGAAAELF.....HE
JJY-2076923-Woodsia_scopulina     ..............................................................................
JJY-2077796-Woodsia_scopulina     .......LFLFSLKARDQQR....................KMAGAISAWLQSFDSALKS..RDLSAATELF.....CP
A-2072483-Polypodium_amorphum     .......................................EMAHERVTVWLQSFERALHE..RDVDAAAELF.....HP
A-2073185-Polypodium_amorphum     .........................................MAGRISKWLQSFDSALKS..NDVSGATELF.....CA
A-2073213-Polypodium_amorphum     NHVFSQRLTLFLTHTQQHAQ...............PTDHPHMAGRVRAWLDSLDSALQK..KDISAAAHLF.....HA
-2003047-Cystopteris_protrusa     .........................................MAERVREWLKSLEWALNE..RDVGAAAELF.....QE
-2007507-Cystopteris_protrusa     .........................................MASRISAWLASFDSALQS..GDLSAATELF.....CA
-2012306-Cystopteris_protrusa     ...........HTHT.......................HARMAGRISAWLESLDSALQK..KDISAAAHLF.....HG
038519-Heterosiphonia_pulchra     ..............................................................................
042241-Selaginella_kraussiana     .....................................MAEAANVQVDAWLARFEASLKK..NDVQNISDLF......A
A|LP-ZZOL-2000160-Selaginella     ............MAT..................MNGSPKIDPDALVDAWLARLDSSLAS..GDASAAANLF.....EE
```

```
                                         50        60         70           80                      90
SisterYUCCA|Mapoly0052s0062     EECFWRDMLAFTWNLYTAESRDEIAAMMNSTLAT...VKPSGWEL...DGE..........V.EE......KG.G.A.
SisterYUCCA|Mapoly0122s0026     QECFWRDLLPFTWNIETFESKDEILKMLTENLAR...VKPRAFVI...DGD..........V.IDRP.....N.G.L.
SisterYUCCA|Sm113792            STSFWRDLIAFTWNIVTVEGRSQIHAMLQSTLES...VAPKGWVR...NGT..........A.SYSA....DS.G.V.
SisterYUCCA|Sm422043            STSFWRDLIAFTWNIVTVEGRSQIHSMLESTLES...VAPKGWTR...HGT..........A.NYSA....DS.G.V.
SisterYUCCA|kf100109_0340       DQCFWRDMVSFTWNVKTAEGKDEIRAMLEATLPS...AAPASWRV...EGE..........A.MEMF.......G.A.
4265-Phaeomegaceros_coriaceus   EECYWRDLVAFTWNIYTAESRDEIKAMLEATLAQ...TKPSSFQL...EET..............SE....QA.G.I.
W-AKXB-2007730-Phaeomegaceros   ..........................MLEANLAE...IKPSRWEM...DGE..........A.VEVE....ES.G.T.
W-AKXB-2007731-Phaeomegaceros   ..............................................................................
W-AKXB-2067423-Phaeomegaceros   EECFWRDLVAFTWNIYTAETKDEIEDMLEARLAE...VKPHRWEV...DGE..........A.VE......VG.G.T.
BMJR-2002471-Adiantum_tenerum   ..........................MLEVTLGH...TKPTNWVI...DGE..........A.DEAD....DG.A.L.
BMJR-2059933-Adiantum_tenerum   ..............................................................................
BMJR-2062494-Adiantum_tenerum   ..........................MLEATLPH...ITPSLWTL...DGP..........V.EEAE....DG.A.V.
U-2019103-Radula_lindenbergia   ..............................................................................
U-2086103-Radula_lindenbergia   AESWWRDHLALEWDFHTLHTLPTITSFLQQRLPG...TQFSKFKLAE.SGPFAPS......ASTPIE.......G.L.
CA|HW-DXOU-2035997-Nothoceros   ..............................................................................
FAJB-2001596-Paraphymatoceros   ..............................................................................
FAJB-2004773-Paraphymatoceros   EECFWRDLLAFTWNIYTAESRDEVAEMLKATLSE...VKPELWEL...DGE..........A.LE......VG.G.T.
FAJB-2008542-Paraphymatoceros   ..............................................................................
CA|GA-FPCO-2029146-Interfilum   EDCFWRDMVAFTWNIRTAEGREEIRTMLDATLKD...TSPKNWKI...EGE..........A.VEMF.......G.A.
GA-FQLP-2033521-Klebsormidium   DQCFWRDMVSFTWNIKTAEGKDAIRAMLEATLPS...TAPSNWRI...EGE..........A.TDMF.......G.A.
YUCCA|LM-GANB-2011315-Cyathea   ..........................MLEATLDD...TKPFGWEI...EGE..........A.EEGQ....DG..V.
YUCCA|LM-GANB-2011316-Cyathea   ..............................................................................
YUCCA|LM-GANB-2017253-Cyathea   ..........................MLEATLEH...TKPSLWAI...DGE..........A.EVAE....GGAA.V.
ANB-2070212-Cyathea_spinulosa   DECFWRDLVAFTWNLYTAESREEILAMLEATLKE...TMPSRWAI...EGE..........V.EEGE....DG.Q.L.
LW-HERT-2045223-Sphaerocarpos   EECFWRDLLSFTWNLYTAESREEIAQMMKATLEE...AKPSRWEL...DGE..........V.VE......NG.G.A.
UCCA|RA-IEHF-2003087-Dumontia   ..............MPGLPLTPPMPVLMPTCVCQMPACVPPPCSL.............................S.C.
O-2067228-Marchantia_paleacea   EECFWRDLLAFTWNLYTAESRDEIAAMMKSALAT...VKPSGWEV...DGE..........V.ED......KG.G.V.
O-2067575-Marchantia_paleacea   QECFWKDLLAFTWNFDTFESRDEILQMLEENLAT...VKPRGWEI...NGH..........V.FEKP.....G.G.L.
-2046311-Conocephalum_conicum   PECFWRDFVAFSWNIQTFELRDEIVPMLKENLAR...VKPRGFEI...NGP..........T.FEKP.......M.M.
-2046658-Conocephalum_conicum   EGCFWRDLLAFTWNFYTAESRDQIGTMLKSTLAE...TKPSGWKV...DGE..........V.QE......NN.H.V.
-2181954-Selaginella_wallacei   GGCLWRDLVSFTWNITTLEGRSQVLQMLQACLDQ...ANPSNWKR...QGN..........A.SPDD....PS.G.IG
A|LP-KUXM-2009452-Selaginella   VDGYWRDLVAFTWNIITVQGLSEIASLIKGTVKH...IDPVGWTR...EGN..........V.VKED.....S.D.T.
|LM-PIVW-2024618-Ceratopteris   EECFWRDLLSFTWNIYTAESRQEIRTMLEETLTN...TKPSNWSI...QGE..........P.EELED..ADG.P.Y.
2005240-Isoetes_tegetiformans   GDAYWRDLLSFTWNIITVEGREEIGRMLESTLEH...TNPKGFIR...DSE..........A.TAYR....DD.G...
2005241-Isoetes_tegetiformans   ..............................................................................
2008401-Hymenophyllum_bivalve   DECFWRDLVAFTWNIYTAESRKEIQAMLEATLEE...SKPCGWEL...DGE..........V.EEDA....TQ.G.V.
003161-Megaceros_vincentianus   EECYWRDLLAFTWN................................................................
007840-Megaceros_vincentianus   DECFWRDLVAFTWNVYTAESRDEIGAMLDANLAG...IKPQRWEV...DGE..........A.VQVE....ES.G.T.
013674-Megaceros_vincentianus   ..............................................................................
016968-Megaceros_vincentianus   EECFWRDLLAFTWNIYTAENRDEIKDMMEARLAE...VKPDRWAV...DGE..........A.VE......VG.G.I.
019507-Megaceros_vincentianus   ..............................................................................
2000839-Marchantia_emarginata   ..........................MLEQTLET...TKPSGWEI...DGP..........A.LPMD.......G.G.
2004520-Marchantia_emarginata   QECLWRDLLAFTWNIQTFESRDEIVSMLKENLAR...VKPRGWQI...DGL..........V.YEKP.....G.G.L.
2069965-Marchantia_emarginata   ..............................................................................
2074864-Marchantia_emarginata   ..............................................................................
2050988-Anthoceros_agrestis-A   EECFWRDLLAFTWNIYTAESKQEIAAMLEATLDE...VKPDLWEL...DGE..........V.TE......AG.G.V.
VB-2006165-Lunularia_cruciata   EECFWRDFVAFTWNIHTAESKEEIAALLEATLDA...TKPSGWEI...DATVEQTDEPSDQA.EEVN.......A.L.
VB-2006166-Lunularia_cruciata   DECFWRDFVAFTWNIQTLESKDAIAAMLEATLEA...TSASGWEL...SGD..........A.REAN.......G.L.
VB-2006167-Lunularia_cruciata   ..............................................................................
VB-2006168-Lunularia_cruciata   ..............................................................................
VB-2007869-Lunularia_cruciata   EDCYWRDLVAFTWNFYTAESRDEIAAMMKSTLNE...VKSVRWEL...DGN..........V.ES......KD.G.L.
VB-2075591-Lunularia_cruciata   EECFWRDFVAFTWNIYTADTKDQIAALLRDNLTR...VKPTGWEI...DDA..........A.FERP.....G.G.V.
CRN-2050303-Megaceros_tosanus   ..............................................................................
CRN-2052782-Megaceros_tosanus   GECFWRDLVAFTWNVYTAESRDEIRAMLDANLSG...IKPSRWQV...DGA..........A.VEVE....ES.G.T.
VITX-2002371-Blechnum_spicant   ..............................................................................
VITX-2009142-Blechnum_spicant   EECFWRDLVAFTWNLYTAESRDDIRAMLEATLRH...ANPSNWAI...DGN..........A.EEAE....GG.A.V.
VITX-2009143-Blechnum_spicant   EECFWRDFVTFTWNLYTAESREDICAMLEATLEH...TNPSVWAI...DGE..........A.EESE....GG.A.L.
VITX-2016791-Blechnum_spicant   EECFWRDLVSFTWNIETAESRDEIRAMLDATLHE...TKPCNWEI...VGD..........V.EILP....DD.K.V.
VITX-2098911-Blechnum_spicant   HECFWRDLVAFTWNFYTAESREEIRAMLQATLED...AKPSAWQL...EGD..........V.EEAE....DG.T.L.
VITX-2099259-Blechnum_spicant   EECFWRDMVAFTWNYYTAESRENIIAMLDATLEH...TKPSGWAI...QGE..........V.EEIC....KG.G.I.
CA|HW-WCZB-2015184-Phaeoceros   ..............................................................................
CA|HW-WCZB-2016004-Phaeoceros   EECFWRDLLAFTWNIYTAESKDEVAEMLKATLSE...VKPDLWEV...DGE..........A.VE......VG.G.T.
CA|HW-WCZB-2117013-Phaeoceros   EECFWRDLVAFTWNLYTAESRDEIAAMLDATLAD...VRPVHWEL...DGE..........A.KET.....DA.G.L.
CA|HW-WCZB-2118293-Phaeoceros   ..............................................................................
U-2004486-Leucostegia_immersa   ..............................................................................
U-2004559-Leucostegia_immersa   ..........................MLEATLPH...TRPSNWAV...DGH..........G.EEAE....GG.A.V.
JLO-2000942-Riccia_berychiana   ..............................................................................
JLO-2002497-Riccia_berychiana   EECFWRDLLAFTWNLYTAESRDEIAAMMKSTLAE...TKPSGWQV...DGK..........V.EE......KD.N.V.
JLO-2033265-Riccia_berychiana   SECFWRDFLTFTWNIETFESKEEIIAMLKETLAR...VKPRGWQI...SGS..........A.HEKP.......M.M.
|LM-WQML-2003640-Cryptogramma   EECFWRDFVSFTWNLYTAESREEICAMLEATLDH...TKPSNWSI...DGP..........A.DEAE....DG.A.L.
|LM-WQML-2004411-Cryptogramma   ..........................MLDATLKA...TKPSNWSV...VGD..........V.EAVEVLGPDGVD.VI
|LM-WQML-2067992-Cryptogramma   ..............................................................................
2095928-Ophioglossum_vulgatum   ..........................MLEATLDL...TKPWGWTV...EKE..........L..PAE....DG.G.I.
2024024-Polysiphonia_japonica   DDCYWRDLVSLTWNIVTLEGKPAIRQMLETTHSS.KDTCSSDWKL...ESM..........Y.HSSS.......E.T.
-2100101-Pallavicinia_lyellii   ..............................................................................
JJY-2002336-Woodsia_scopulina   ..............................................................................
-2008616-Woodsia_scopulina.p2   HECFWRDLVTFTWNFYTAESREEIRAMLHATLQH...ANPSAWQL...EGD..........V.EEAE....DG.N.L.
JJY-2076923-Woodsia_scopulina   ..............................................................................
JJY-2077796-Woodsia_scopulina   GECFWRDLVAFTWNLFTAESRSDIRAMLEATLHH...TNPSNWAI...DGN..........A.DEAE....GG.I.I.
A-2072483-Polypodium_amorphum   HECFWRDLVAFTWNFYTAESREEIQAMLHATLDY...TNPSDWQI...QGD..........V.EEGE....DG.S.L.
A-2073185-Polypodium_amorphum   EECFWRDLIAFTWNLYTAESREDIHSMLGATLHH...THPSNWAL...DGT..........A.EEVE....SG.A.V.
A-2073213-Polypodium_amorphum   EECFWRDFVSFTWNLYTAESREDICAMLDATLEL...TQPSAWAI...DGE..........VEEEAEG..GAG.A.L.
-2003047-Cystopteris_protrusa   HECFWRDLVAFTWNFYTAESRQEIRAMLHATLED...VKPSAWQI...DGD..........V.EEGE....DG.SQL.
-2007507-Cystopteris_protrusa   EECLWRDLVAFTW....XESRHDIRAMLHATLPH...TCPSAWAI...DGE..........V.QELE....AG.A.V.
-2012306-Cystopteris_protrusa   DECFWRDFVTFTWNLYTAESRKDICAMLEATLEH...TNPSVWAI...DGE..........A.EEAE....GG.A.L.
038519-Heterosiphonia_pulchra   ..........................MLQETLSD...TRPNEWTVCEEDGVHEQH........EEDR....DV.K.V.
042241-Selaginella_kraussiana   EECFWRDLVAFTWNILTLEGRTQVTSMLESCLQQ...ASPCEWKR...EGD..........A.SLEN....DG.N.G.
A|LP-ZZOL-2000160-Selaginella   STSFWRDLIAFTWNIVTVEGRSQIHAMLQYTLES...VAPKGWVR...NGT..........A.SYSA....DS.G.V.
```

```
                                       100           110                 120                130          140          150
SisterYUCCA|Mapoly0052s0062       ..LQVFLKFETSIARGRGHLRLKGG.....KCWTL....LTAMTELKGYEEHLGSARPMGVKNENVPGR.KT..WLEE
SisterYUCCA|Mapoly0122s0026       ..MGSAFKFETSLAWCRGFVWLKGG.....KCWTI....TTSMWDLKGYEEKISRSRPFGPTLGQIKGR.KT..AYQE
SisterYUCCA|Sm113792              ..IEAWIKFETRDAVCTGHLRLMATTS...LCRTL....LTAMEGLRDFPENKGRTRPNGVTHGVIRHR.AS..WLDG
SisterYUCCA|Sm422043              ..IEAWIKFETRVAVCTGHLRLVATTS...LCRTL....LTAMEGLRDFPEKKGRTRPNGVTHGVIRHR.AS..WLDG
SisterYUCCA|kfl00109_0340         ..VTAFVAFETATGRGKGHLRLVDG.....KCFTL....LTTLYELKGFEEKIGASRLPGRNLAPEPNR.QS..WAAK
4265-Phaeomegaceros_coriaceus     ..IQAFFTFETGVARGRGIARLQGE.....KCWIL....TTSIHELKGYEEAIGARRPF..................
W-AKXB-2007730-Phaeomegaceros     ..VEACIRFETAVARGRGHVRLKGG.....KCWTL....LTAIGELKGHEEKAGKLRVNGVTHGAVSGR.TT..WLQE
W-AKXB-2007731-Phaeomegaceros     ..............................................................................
W-AKXB-2067423-Phaeomegaceros     ..VDAMLRFETAVGKGRGHVRLKGG.....KCWTM....FTALRELKGHEEKVGEARPPGVTYGAVPGR.KT..WREE
BMJR-2002471-Adiantum_tenerum     ..VQAFFTFETERSRGRGLVKLKGE.....KCWCL....FTSMVELKGYEEKAGLRREMGVQHGSFKGR.KT..WLEG
BMJR-2059933-Adiantum_tenerum     ..............................................................................
BMJR-2062494-Adiantum_tenerum     ..LQAFITFETQRARGRGYVKLRGD.....KCWCL....LTSMLELKGFEEKVGIRREPGVRHGAIKGR.KT..WLDS
U-2019103-Radula_lindenbergia     ..............................................................................
U-2086103-Radula_lindenbergia     EWIESMFSFETKVGRGKGMIRLVQGEDGVYKGYML....HTSLQELKGFEESSGARRPHGGNNSLSGGV.KSGNWLER
CA|HW-DXOU-2035997-Nothoceros     ..............................................................................
FAJB-2001596-Paraphymatoceros     ..............................................................................
FAJB-2004773-Paraphymatoceros     ..VDATLKFETAVGRGRGHVRLKGN.....KCWTM....FTALRELKGHEEKIGKARPTGLLYGAVPGR.KT..WREE
FAJB-2008542-Paraphymatoceros     .............................................................................K
CA|GA-FPCO-2029146-Interfilum     ..VNAFFTFETKVGRGRGHLRLVDN.....RCFTL....LTTLYELKGFEEKVGRNRPSGRELFSQSKK.ET.....Q
GA-FQLP-2033521-Klebsormidium     ..VNAFVAFDTATGRGKGHLRLVDG.....KCFTL....LTTLCELKGFEGKTGASRSGRNLTPEPNR.RS..WGAK
YUCCA|LM-GANB-2011315-Cyathea     ..LQAFFRFETQTARGRGVVRLRGG.....KCWCL....ITSMLELKGHEENAGFRRQLGVQHGAVKGR.KT..WLEG
YUCCA|LM-GANB-2011316-Cyathea     ..............................................................................
YUCCA|LM-GANB-2017253-Cyathea     ..IQAFITFETQRARGRGHVKLRGD.....KCWCL....FTSIVELKGYEE....RREFGVQHGAIKGR.KS..WLER
ANB-2070212-Cyathea_spinulosa     ..LQAFISFETQRARGRGHVKIRGD.....KCWCL....LTAMQELKGHEEKAGLRRELGVEHGAVEGR.KT..WLEG
LW-HERT-2045223-Sphaerocarpos     ..LCAFIKFETSLSRGRGHLRLRGG.....RCWTL....LTAMTELKGHEEPLSTLRPLGTGKAG.....KT..WVDG
UCCA|RA-IEHF-2003087-Dumontia     .................VSLRAM.....KSWSVPLARLACITPSPVKPSQYLPARPVCA.........LS..WVSS
O-2067228-Marchantia_paleacea     ..LQVFLKFETSVARGRGHLRLKNG.....KCWTL....LTAMTELKGHEEHLGPARPMGVKNENVPGR.KT..WLEE
O-2067575-Marchantia_paleacea     ..IGSAFEFETSLAWGRGFVWLRGG.....KCWTI....MTSMWDLKGHEEKIARSRPFGATIGQVKGR.KT..WYQQ
-2046311-Conocephalum_conicum     ..TGAGIKFETSVGWGRGFLWLRGS.....KCWTI....MTSMWELKGHEEGIARFRPFGATPGSARDR.KT..WAQE
-2046658-Conocephalum_conicum     ..LSVFLIFETAIGRGRGHLRLKGG.....KCWTL....LTALTELKGHEQPVGMARPRGNQDGNVPGR.KP..WHEE
-2181954-Selaginella_wallacei     RGPTAWLCFETSVGDCTGHVRLQSGGGGGGRCTTL....LTTMAGLKAHPEAIGKNRVKGVEHGASKQR.KT..WLEK
A|LP-KUXM-2009452-Selaginella     ..IAAWFSFQTSVGACKGHLRLKNG.....KCWTL....FTSITELTGHPQKIGRNRPK...............YNQ
 |LM-PIVW-2024618-Ceratopteris    ..IQAFVRFETEKARCRGLVKLRGD.....KCWCL....FTTMVELKGFEEPAGRRREIGVQHGSVKGR.KS..WLQK
2005240-Isoetes_tegetiformans     ..LQAWFRFETDVGNCKGHVRIKGDG....KCRTL....FTAIAELKGYPEQKGRNRPYGAEHGVIQGR.KT..WLDR
2005241-Isoetes_tegetiformans     ..............................................................................
2008401-Hymenophyllum_bivalve     ..VQAFISFETARARGRGHVRIKGDN....KCWTL....LTAMTELKGHEEKARRTRPKGVEHGVVRGR.KT..WLEE
003161-Megaceros_vincentianus     .......................................ITSIHELKGYEEAIGQRRPF..................
007840-Megaceros_vincentianus     ..VEACIRFETAAARGRGHVRLKGG.....KCWTL....LTAMGELKGHEEKAGKLRANGLTHSVVSGR.KT..WLQE
013674-Megaceros_vincentianus     ..............................................................................
016968-Megaceros_vincentianus     ..VDALLKFETAVGKGRGHVRLKGG.....KCWTM....FTALRELKGHEEKVGDARPQGVIYGAVPGR.KT..WREE
019507-Megaceros_vincentianus     ..............................................................................
2000839-Marchantia_emarginata     ..VEACITFETSVARGRGYVKLKEG.....KCWVL....MTAMWELKGHEEKAGLFRPQGVPYGTIRGK.KT..WLEK
2004520-Marchantia_emarginata     ..IGSGVTFETSFAWGRGFLWLKGG.....KCWTI....MTTMWDLKGHEEKIARSRSLGPTIGQISGR.KS..WLQA
2069965-Marchantia_emarginata     ..............................................................................
2074864-Marchantia_emarginata     ..............................................................................
2050988-Anthoceros_agrestis-A     ..VEAMLKFETAAGRGRGVVRLKDG.....KCWTM....QTAVRELKGYEEKVGQARPHGVVFGAVPGR.KT..WRAK
VB-2006165-Lunularia_cruciata     ..VEVPIKFETSVARCRGNVKLKRG.....KCWIL....FTAAWELKGHEEPAGRFRPIGKPYGVVRNR.KT..WLEK
VB-2006166-Lunularia_cruciata     ..VEGNLTFETSVARCRGYVKLKGG.....KCWVL....FTAAWELKGHEEEASKFRPIGLPYGSVRNR.KT..WLQK
VB-2006167-Lunularia_cruciata     ..............................................................................
VB-2006168-Lunularia_cruciata     ..............................................................................
VB-2007869-Lunularia_cruciata     ..QSALIKFETSVGRGRGHLRLRGG.....KCWTL....LTALTELKGYEEQLNGARPLGTLHGAIGGG.KT..WLEE
VB-2075591-Lunularia_cruciata     ..VGAKIKFETSVGWARGFVWLKGE.....KCWIM....MTAIWDLKGHEEKIGKRRPTGPTFGQVCGR.KS..WIEQ
CRN-2050303-Megaceros_tosanus     ..............................................................................
CRN-2052782-Megaceros_tosanus     ..VEAWVSFETAAGRGRGHVRLKGR.....KCWTL....LTAMVELKGHEELAGKLRVSGVTHGAVSGR.KT..WLEA
VITX-2002371-Blechnum_spicant     ..............................................................................
VITX-2009142-Blechnum_spicant     ..LQAFITFETESARCRGYVKLRGV.....KCWCL....LTSMLELKGHEENAGLRRELGVQHGAVKGR.KT..WLDS
VITX-2009143-Blechnum_spicant     ..IQAFITFETQRARGRGHVKLRGD.....KCWCL....FTSIVELKGYEEKAGFRREFGVQHGAIKGR.KT..WLES
VITX-2016791-Blechnum_spicant     ..VECFVRFETERGRGKGIVKLKGD.....KCWCL....LTTLQELKGFEEETGPTREVGAVHGAIKGR.KT..WLEG
VITX-2098911-Blechnum_spicant     ..VQAFIKFETERARGKGHVKLIGD.....KCWCL....LTTMQELKGLEERAGLKREVGVEHGAAKGK.RT..WLEC
VITX-2099259-Blechnum_spicant     ELVEAFISFETESARGKGHLSLRGD.....KCWCL....LTAMQELKGHEEIAP.......................VER
CA|HW-WCZB-2015184-Phaeoceros     ..............................................................................
CA|HW-WCZB-2016004-Phaeoceros     ..VDAMLKFETAVGRGRGHVRLKGN.....KCWTM....FTALRELKGHEEKVGKARPTGLVYGAVPGR.KT..WREE
CA|HW-WCZB-2117013-Phaeoceros     ..VDACLKFETRVARGRGHVRLKGG.....KCWTL....LTAIAELKGHEEKAGSLRDRGLQYGAVRGR.KS..WLEE
CA|HW-WCZB-2118293-Phaeoceros     ..............................................................................
U-2004486-Leucostegia_immersa     ............................................................................EG
U-2004559-Leucostegia_immersa     ..LQAFITFETKSARCRGYVKLRGD.....KCWCL....LTSMLELKGYEEKAGVRRALGVHHGAIKGR.KT..WLDR
JLO-2000942-Riccia_berychiana     ..............................................................................
JLO-2002497-Riccia_berychiana     ..LSVFLKFESAIGRGRGHLRLKGG.....KCWTL....LTAMTELKGHEEPLGMARPSGVKDGNVPGR.KS..WLEG
JLO-2033265-Riccia_berychiana     ..VGAGIKFETSFAWGRGFLWLKGG.....KCWTM....MTTMWELKGHEERIARSRPFGVTIGSVSGR.ET..WVQQ
 |LM-WQML-2003640-Cryptogramma    ..IQAFITFETERGRGRGHVKLRGE.....KCWCL....FTSMVELKGYEEKAGIRRDMGVQHGAIKGR.KT..WLEG
 |LM-WQML-2004411-Cryptogramma    HYDQCFATFETERGRCKAMVKLHGD.....KCWCL....LTALQELKGYEERA....DVGVEHGAIRGR.KT..WLER
 |LM-WQML-2067992-Cryptogramma    ...................................................QHSAAKGR.KT..WLEL
2095928-Ophioglossum_vulgatum     ..VNAFISFETERAHCRGHVRLRGD.....KCWTL....LTSMEGLRGYEEKAGAIREKGATHGAIKGR.KT..WLDM
2024024-Polysiphonia_japonica     ..VEAWVKFNTSYIRGLAHVRLTKNEKDEMLCLTM....LTTAQEIIGHEEPHGLTRPLGTDHEVYKGR.ES..WIEK
-2100101-Pallavicinia_lyellii     ..............................................................................
JJY-2002336-Woodsia_scopulina     ..............................................................................
-2008616-Woodsia_scopulina.p2     ..VQAFIRFETERARGKGHVKLLGD.....KCWCL....LTTMQELKGLEERAGLKREVGVQHGAAKGK.RT..WLEC
JJY-2076923-Woodsia_scopulina     ..............................................................................
JJY-2077796-Woodsia_scopulina     ..LQAFITFETERARCRGYVKLKGD.....KCWCL....LTTMLELKGHEEKAGLRRELGIQHGAIKGR.KT..WLDS
A-2072483-Polypodium_amorphum     ..VQAFISFETERAKGSGHVKLIGD.....KCWCL....LTTMQELKGFEERAGPKREVGVQHGATKGR.KT..WAES
A-2073185-Polypodium_amorphum     ..LQAFITFETERARCRGYVKLRGD.....KCWCL....LTSMLELKGHEENAGVRRELGVQHGAFKGR.KT..WLDS
A-2073213-Polypodium_amorphum     ..TQAFITFETQRARGRGYVKLHGD.....KCWCL....FTSMVELKGFEEKAGVRRESGVQHGAIKGR.KT..WLES
-2003047-Cystopteris_protrusa     ..VQAFIKFETERARGKGHVKLIGD.....KCWCL....LTTMQELKGLEESAGLKREVGIQHGAAKGK.RT..WLEC
-2007507-Cystopteris_protrusa     ..LQAFFTFETQAALCRGILKLRDN.....KCWCL....LTSMLELKGHEEKAGLGRELGVQHVATKGR.KT..WLDG
-2012306-Cystopteris_protrusa     ..IQAFISFETQRARGRGHVKLRGD.....KCWCL....FTSMVELKGYEEKAGFRREFGVQHGTIKGR.KS..WLES
038519-Heterosiphonia_pulchra     ..VSFWINFQTTHCHCLGRVVLHNS.....KCHVL....LTTAQALRRFPEQCGKSRELGKRRGDNPGAEFS..WGSR
042241-Selaginella_kraussiana     ..TSAWVRFETEVGLCSGHIRLASDGG...TATTL....FTTLVSFKQYPETTGDKRPIGAEHGVHRPPGKT..WLQT
A|LP-ZZOL-2000160-Selaginella     ..IEAWIKFETRDAVCTGHLRLMATTS...LCRTL....LTAMEGLKDFPEKKGHTRPKGVTHGVIRHR.AS..WLDG
```

GXGXXG

```
                                      160       170       180       190       200       210
SisterYUCCA|Mapoly0052s0062         RMQE..RQELGYKS.QPYCVIVGGGQAGIALAAR...LRMLNVP.....ALIIEKN.....ARPGDSWRNR.......
SisterYUCCA|Mapoly0122s0026         KQQE..QHDMGRSK.QPYCLVVGAGQGGVMLGAR...LRMLGVP.....AIIVEKN.....ERLGDNWRNR.......
SisterYUCCA|Sm113792                RKEE..ERTLGSTV.QPYCVIVGGGQAGIGLAAR...LRQLGVP.....CIVVEKN.....PRPGDSWRNR.......
SisterYUCCA|Sm422043                RKED..ERTLGSTV.QPYCVIVGGGQAGIGLAAR...LRQLGVP.....CIVVEKN.....PRPGDSWRNR.......
SisterYUCCA|kf100109_0340           RELE..RGELGYGK.QPFVVIVGAGHSGVMLAAR...LRALNVP.....TIVVDKG......SPGDSWRNR.......
4265-Phaeomegaceros_coriaceus       RAAS..STARTSDA.EPYCVIVGGGQAGISLAAR...LKHLDVR.....AIVVDKN.....HLPGDSWRNR.......
W-AKXB-2007730-Phaeomegaceros       RLSE..EAELGYSK.QPYCVVVGGGQGGIVLGAR...LRRLGVP.....TIIVEKN.....PRAGDSWRNR.......
W-AKXB-2007731-Phaeomegaceros       ..............................................................................
W-AKXB-2067423-Phaeomegaceros       RVEE..EKKLGYSQ.QPYCVVVGGGQAGIALGAR...LRRLGVP.....TIIVEKN.....GRPGDSWRNR.......
BMJR-2002471-Adiantum_tenerum       RIAE..QEELGKSK.QPYCLIVGGGQGGIALGAR...LRRLGVP.....TIIVEKN.....KRPGDSWRNR.......
BMJR-2059933-Adiantum_tenerum       ........................GQGGITLGAR...LRRLGVP.....SIIVEKN.....ARAGDSWRQR.......
BMJR-2062494-Adiantum_tenerum       RVEE..EQELGRSR.QPHCLIVGGGQAGIALGAR...LRRLCVP.....TIIIEKN.....KRPGDSWRNR.......
U-2019103-Radula_lindenbergia       ..............................................................................
U-2086103-Radula_lindenbergia       RQ....RAVEFQEE.QPQVLIIGAGQAGLNMAAR...LQSLNMS.....SLLIDKN.....ARIGDNWRSR.......
CA|HW-DXOU-2035997-Nothoceros       ..............................................................................
FAJB-2001596-Paraphymatoceros       ..............................................................................
FAJB-2004773-Paraphymatoceros       RQEE..EGKLGYSE.QPYCVVVGGGQSGIALGAR...LRRLGVP.....TIIVEKN.....KRPGDSWRNR.......
FAJB-2008542-Paraphymatoceros       KRQE..EAELGYSK.QPYCVVVGGGQGGIALGAR...LRRLGVP.....SIIVEKN.....ARAGDSWRNR.......
CA|GA-FPCO-2029146-Interfilum       LSNE..QVELDLSK.QPYVVIVGAGHSGVMLAAR...LKILKVP.....TILVEKG......APGDSWRKR.......
GA-FQLP-2033521-Klebsormidium       RELE..RGELGYDK.QPFVVIVGGGHSGMMLAAR...LRALNVP.....TIEVDKG......SPGDSWRNR.......
YUCCA|LM-GANB-2011315-Cyathea       RLEE..IEELGRTR.QPYCLIVGGGQGGIALGAR...LRRLGVP.....TIIVEQN.....KRPGDSWRNR.......
YUCCA|LM-GANB-2011316-Cyathea       ..............................................................................
YUCCA|LM-GANB-2017253-Cyathea       RIEE..QEELGRTK.QPYCLIVGGGQGGIALGAR...LRQLQVP.....TIIVEKH.....QRPGDSWRNR.......
ANB-2070212-Cyathea_spinulosa       RVEE..LEELGRSR.QPYCVIVGGGQGGIALGAR...LRRLGVP.....TIIIEKN.....ERAGDSWRNR.......
LW-HERT-2045223-Sphaerocarpos       RMEE..KQQLGYES.QPYCVIVGGGQAGIILGAR...LRMLKVP.....TIIIEKN.....ERPGDSWRKR.......
UCCA|RA-IEHF-2003087-Dumontia       RTLS......................SSLSVVARVVFLWLLASPAW..MSLPLSSNVTGALGTPGERDTARCACTIQC
O-2067228-Marchantia_paleacea       RLQE..RQELGYKS.QPYCVIVGGGQAGITLGAR...LRMLNVP.....SIIIEKN.....ARPGDSWRNR.......
O-2067575-Marchantia_paleacea       KQQE..QQDVGRSK.QPYCLIVGAGHAGAMLGAR...LRMLGVP.....TIIVERN.....ERAGDNWRNR.......
-2046311-Conocephalum_conicum       QEQE..RKELGHSK.QPYCLIVGGGHSGVMLGAR...LKMLGVP.....TLIVERN.....ARLGDNWRNR.......
-2046658-Conocephalum_conicum       RQEE..KKELGYKT.QPYVVIAGGGQAGIILAAR...LRMLSVP.....TIIVEKN.....ARPGDSWRNR.......
-2181954-Selaginella_wallacei       RAEE..ESKLGYEQ.QPYVVIVGGGQAGIVLGAR...LRQLGVP.....AIIVEKN.....ARAGDSWRNR.......
A|LP-KUXM-2009452-Selaginella       LQET..ENSLGYQI.QPYCVIIGGGQAGITLGAR...LKLLGVP.....SIIVEKN.....ATPGDSWRLR.......
|LM-PIVW-2024618-Ceratopteris       RIEE..EEALGKTK.QPYVLIVGGGQGGIMLGAR...LRRLQVP.....TIIVDKR.....KRPGDSWRSR.......
2005240-Isoetes_tegetiformans       RKEE..EQTLGYSI.QPYCVIVGGGQAGIGLGAR...LKQLQVP.....TLIVEKN.....GRAGDSWRNR.......
2005241-Isoetes_tegetiformans       ..............................................................................
2008401-Hymenophyllum_bivalve       RQEE..RQLGTGKQ.QPYCVVVGGGQGGIALGAR...LRRLGVP.....TIIVEKN.....ARPGDSWRNR.......
003161-Megaceros_vincentianus       REAS..STARTNNA.EPYCVIVGGGQAGISLAAR...LKHLDVP.....AIVVDKN.....RRPGDSWRNR.......
007840-Megaceros_vincentianus       RLSE..EAELGYSK.QPYCVVVGGGQAGIVLGAR...LRRLGVP.....TIIVEKN.....PRAGDSWRNR.......
013674-Megaceros_vincentianus       ..............................................................................
016968-Megaceros_vincentianus       RAEE..EKKLGYSQ.QPYCVVVGGGQAGIALGAR...LRRLGVP.....SIIVEKN.....ARPGDSWRNR.......
019507-Megaceros_vincentianus       ..............................................................................
2000839-Marchantia_emarginata       RVEE..EQELGHSK.QPYCVVVGGGHSGIALGAR...LRALEVP.....TIILEEN.....ERAGDNWRNR.......
2004520-Marchantia_emarginata       KQQE..QQVLGHSK.QPYCLIVGGGQAGVMLGAR...LRVLGVP.....TIIVEKN.....ERAGDNWRNR.......
2069965-Marchantia_emarginata       ..............................................................................
2074864-Marchantia_emarginata       .......................GGQAGICLGAR...LRMLNVP.....TLIIEKN.....ARPGDSWRNR.......
2050988-Anthoceros_agrestis-A       REEE..EGKLGHSV.QPYCVVVGGGQAGIALGAR...LRRLEVP.....TIVVEKN.....NKPGDSWRNR.......
VB-2006165-Lunularia_cruciata       KEAE..KEELESTN.RPYCVIVGGGQAGIILGAR...LKVLGVP.....TIILEKN.....DKAGDSWRNR.......
VB-2006166-Lunularia_cruciata       KEEE..EKELECNK.RPYCVIVGGGQAGIMLGAR...LKVLGVP.....TIILEKN.....DKAGDSWRNR.......
VB-2006167-Lunularia_cruciata       .............................MLGAR...LKVLGVP.....TIILEKN.....DKAGDSWRNR.......
VB-2006168-Lunularia_cruciata       ........................MQAGIILGAR...LKVLGVP.....TIILEKN.....DKAGDSWRNRHN.....
VB-2007869-Lunularia_cruciata       SLVE..KQKLGYTT.QPFCLIVGGGQAGISLGAR...LRMLKVP.....TIIVEKN.....AKAGDSWRNR.......
VB-2075591-Lunularia_cruciata       RKEE..EQELGHSK.QPYCVIVGGGQTGLAVGAR...LKVLGVP.....TIIVEKN.....ETAGDTWRNR.......
CRN-2050303-Megaceros_tosanus       ..............................................................................
CRN-2052782-Megaceros_tosanus       KVAE..EAELGYSK.QPYCVVVGGGQGGIALGAR...LRRLGVP.....TIVVEKN.....ARAGDSWRKR.......
VITX-2002371-Blechnum_spicant       ..............................................................................
VITX-2009142-Blechnum_spicant       RVEE..LEGLGRTK.QPYCLIVGGGQGGIALGAR...LRRLGVP.....TIIIEKN.....RRPGDSWRNR.......
VITX-2009143-Blechnum_spicant       RIEE..QEELGTTK.QPYCLIVGGGQGGIALGAR...LRRLQVP.....TIIVEKN.....KRPGDSWRNR.......
VITX-2016791-Blechnum_spicant       RIED..MQELGKSK.QPYCLIVGGGQGGIALGAR...LRRLGVPTIIVXTIIVEKN.....KRAGDSWRHR.......
VITX-2098911-Blechnum_spicant       RTEE..LQELGRSK.QPYCIIVGGGQAGVALGAR...LRRLGVP.....TIIVEKN.....KRPGDTWRQR.......
VITX-2099259-Blechnum_spicant       RKEELHVQELGRSK.QPYCLIVGGGHSGIMLAAR...LVRLGVP.....TIVVEKN.....KRAGDSWRKR.......
CA|HW-WCZB-2015184-Phaeoceros       ..............................................................................
CA|HW-WCZB-2016004-Phaeoceros       RQEE..EGKLGYSE.QPYCVVVGGGQSGIALGAR...LRKLGVP.....TIIVEKN.....ERPGDSWRKR.......
CA|HW-WCZB-2117013-Phaeoceros       KVQE..AAELGFSK.QPYCVVVGGGQGGIALGAR...LRRLGVP.....SIIVEKN.....PRAGDSWRNR.......
CA|HW-WCZB-2118293-Phaeoceros       ...................VVVGGGQGGIALGAR...LRRLGVP.....TIIVEKN.....PRAGDSWRNR.......
U-2004486-Leucostegia_immersa       RMEE..MQELGKSK.QPYCLIVGGGQAGIALGAR...LRRLGVP.....TIIVEKN.....ERAGDSWRHR.......
U-2004559-Leucostegia_immersa       RSEE..LEGLGTSK.QPYCLVVGGGQGGIALGAR...LRRLEVP.....TIIIDKN.....KHPGDSWRNR.......
JLO-2000942-Riccia_berychiana       ..............................................................................
JLO-2002497-Riccia_berychiana       RLEE..KENLGYNT.QPYVVIIGGGQAGIILAAR...LRMLNVP.....TIIVEKN.....ARPGDSWRNR.......
JLO-2033265-Riccia_berychiana       KERE..QQELGHEK.QPYCLIVGAGQAGLMLGAR...LRMLGVP.....TIIVEKN.....ARPGDNWRNR.......
|LM-WQML-2003640-Cryptogramma       RIEE..QEQLGKSK.QPYCLIVGGGQGGIALGAR...LRRLHVP.....TIIVEKN.....ERPGDSWRNR.......
|LM-WQML-2004411-Cryptogramma       RTEE..LRELGSSK.QPYCLIVGGGHSGISLGAR...LRRLGVP.....NIVVERN.....PRPGDSWRNR.......
|LM-WQML-2067992-Cryptogramma       RNEE..VQQLESGRLQPYCVIVGGGQAGVALAAR...LRRLGVA.....AIIVEKR.....KRAGDTWRNR.......
2095928-Ophioglossum_vulgatum       KMQE..QQELGVSK.QPYCVIVGGGQGGIMLGAR...LQRLGVP.....TIIVEKN.....ARPGDSWRNR.......
2024024-Polysiphonia_japonica       RLEE..IQQ..NRQ.QPYVVIIGGGQAGITLGSR...LSRLNVP.....YVILERN.....VRAGDSWRKR.......
-2100101-Pallavicinia_lyellii       ..............................................................................
JJY-2002336-Woodsia_scopulina       ..............................................................................
-2008616-Woodsia_scopulina.p2       RTQE..LEELGRSR.QPYCIIVGGGQAGVALGAR...LRRLGVP.....TIIVEKN.....KRPGDTWRHR.......
JJY-2076923-Woodsia_scopulina       ................PYCLIVGGGQCGIALSAY...LDRLGVP.....TIVVDKK.....KHPGDAWRER.......
JJY-2077796-Woodsia_scopulina       RLEE..LEGLGRAK.QPYCLIVGGGQGGIALGAR...LRRLGVP.....TIIIEKN.....KHPGDSWRNR.......
A-2072483-Polypodium_amorphum       RTEE..LEGLERCK.KTYCIIVGGGQAGVALGAR...LRRLGVP.....TIIVERN.....KKAGDTWRQR.......
A-2073185-Polypodium_amorphum       RVEE..LQGLGRTQ.QPYCLIVGGGQCGIALGAR...LRQLRVP.....TIIVEKN.....KRPGDSWRNR.......
A-2073213-Polypodium_amorphum       RIEE..QEELGRSK.QPYCLIVGGGQGGIALGAR...LRRLGVP.....TIIVEKN.....KHPGDSWRNR.......
-2003047-Cystopteris_protrusa       RTEE..LQKLEMRK.QPYCIIVGGGQAGVALGAR...LHRLGVP.....TIIVEKN.....KRPGDTWRHR.......
-2007507-Cystopteris_protrusa       RVNE..LESLGRTR.QPYVLIVGGGQGGISLGAR...LRRLEVP.....TIIVEKN.....KRPGDSWRNR.......
-2012306-Cystopteris_protrusa       RIEE..QEKLGRTE.QPYCLIVGGGQGGIALGAR...LRRLEVP.....TIIVEKN.....KHPGDSWRNR.......
038519-Heterosiphonia_pulchra       RAAR..RQTLGMEE.QPYVVIVGGGQAGVMLGAR...LKRQNVP.....TIILESH.....GRAGDSWRTR.......
042241-Selaginella_kraussiana       RTEE..ESTLGYTV.QPYVVVVGGGQAGIVLGAR...LRYLGVP.....TIVLERN.....ARAGDSWRRR.......
A|LP-ZZOL-2000160-Selaginella       RKEE..ERTLGSTV.QPYCVIVGGGQAGIGLAAR...LRQLGVP.....CIVVEKN.....PRPGDSWRSR.......
```

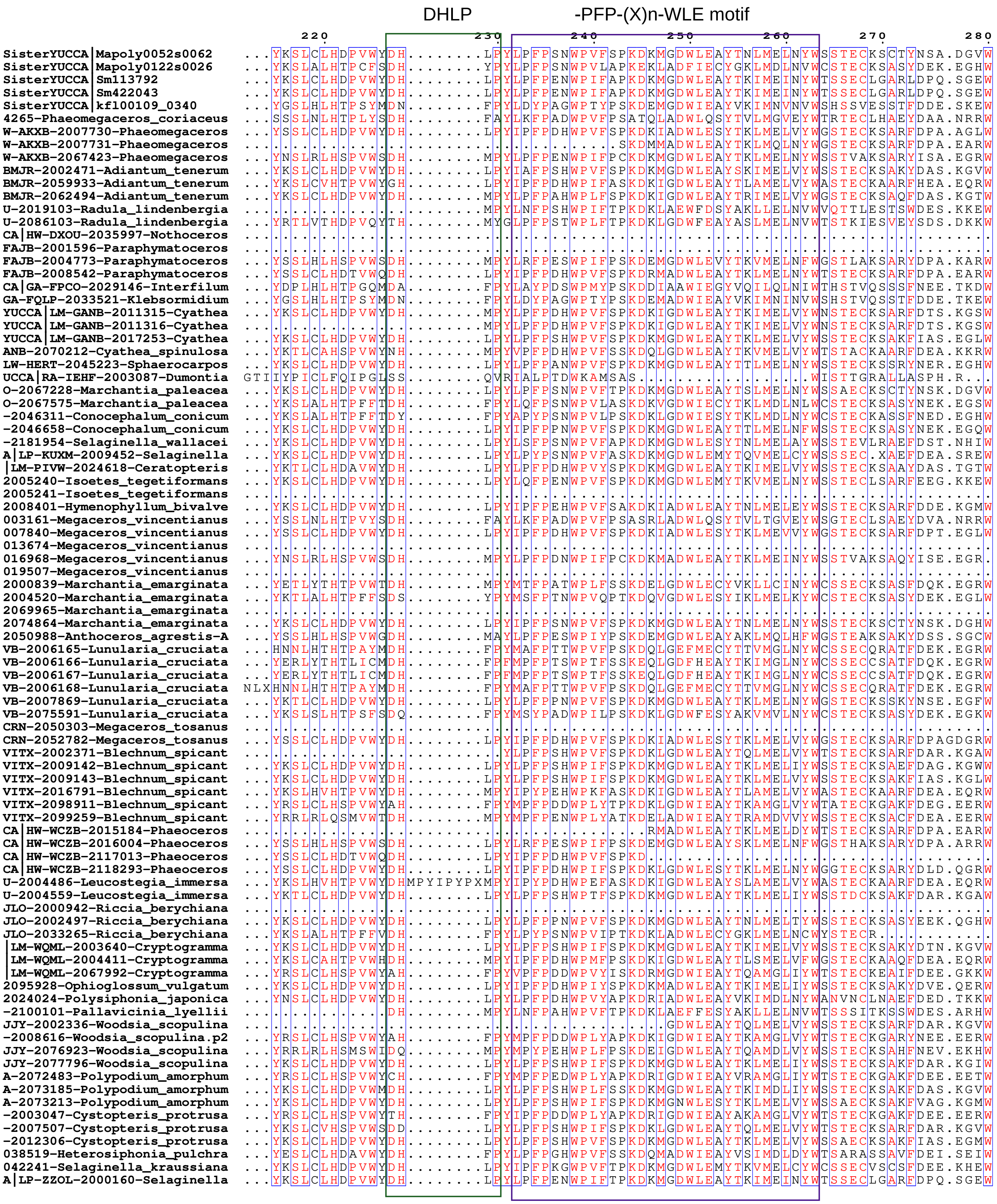
DHLP
-PFP-(X)n-WLE motif
220
230
240
250
260
270
280
SisterYUCCA|Mapoly0052s0062
SisterYUCCA|Mapoly0122s0026
SisterYUCCA|Sm113792
SisterYUCCA|Sm422043
SisterYUCCA|kfl00109_0340
4265-Phaeomegaceros_coriaceus
W-AKXB-2007730-Phaeomegaceros
W-AKXB-2007731-Phaeomegaceros
W-AKXB-2067423-Phaeomegaceros
BMJR-2002471-Adiantum_tenerum
BMJR-2059933-Adiantum_tenerum
BMJR-2062494-Adiantum_tenerum
U-2019103-Radula_lindenbergia
U-2086103-Radula_lindenbergia
CA|HW-DXOU-2035997-Nothoceros
FAJB-2001596-Paraphymatoceros
FAJB-2004773-Paraphymatoceros
FAJB-2008542-Paraphymatoceros
CA|GA-FPCO-2029146-Interfilum
GA-FQLP-2033521-Klebsormidium
YUCCA|LM-GANB-2011315-Cyathea
YUCCA|LM-GANB-2011316-Cyathea
YUCCA|LM-GANB-2017253-Cyathea
ANB-2070212-Cyathea_spinulosa
LW-HERT-2045223-Sphaerocarpos
UCCA|RA-IEHF-2003087-Dumontia
O-2067228-Marchantia_paleacea
O-2067575-Marchantia_paleacea
-2046311-Conocephalum_conicum
-2046658-Conocephalum_conicum
-2181954-Selaginella_wallacei
A|LP-KUXM-2009452-Selaginella
|LM-PIVW-2024618-Ceratopteris
2005240-Isoetes_tegetiformans
2005241-Isoetes_tegetiformans
2008401-Hymenophyllum_bivalve
003161-Megaceros_vincentianus
007840-Megaceros_vincentianus
013674-Megaceros_vincentianus
016968-Megaceros_vincentianus
019507-Megaceros_vincentianus
2000839-Marchantia_emarginata
2004520-Marchantia_emarginata
2069965-Marchantia_emarginata
2074864-Marchantia_emarginata
2050988-Anthoceros_agrestis-A
VB-2006165-Lunularia_cruciata
VB-2006166-Lunularia_cruciata
VB-2006167-Lunularia_cruciata
VB-2006168-Lunularia_cruciata
VB-2007869-Lunularia_cruciata
VB-2075591-Lunularia_cruciata
CRN-2050303-Megaceros_tosanus
CRN-2052782-Megaceros_tosanus
VITX-2002371-Blechnum_spicant
VITX-2009142-Blechnum_spicant
VITX-2009143-Blechnum_spicant
VITX-2016791-Blechnum_spicant
VITX-2098911-Blechnum_spicant
VITX-2099259-Blechnum_spicant
CA|HW-WCZB-2015184-Phaeoceros
CA|HW-WCZB-2016004-Phaeoceros
CA|HW-WCZB-2117013-Phaeoceros
CA|HW-WCZB-2118293-Phaeoceros
U-2004486-Leucostegia_immersa
U-2004559-Leucostegia_immersa
JLO-2000942-Riccia_berychiana
JLO-2002497-Riccia_berychiana
JLO-2033265-Riccia_berychiana
|LM-WQML-2003640-Cryptogramma
|LM-WQML-2004411-Cryptogramma
|LM-WQML-2067992-Cryptogramma
2095928-Ophioglossum_vulgatum
2024024-Polysiphonia_japonica
-2100101-Pallavicinia_lyellii
JJY-2002336-Woodsia_scopulina
-2008616-Woodsia_scopulina.p2
JJY-2076923-Woodsia_scopulina
JJY-2077796-Woodsia_scopulina
A-2072483-Polypodium_amorphum
A-2073185-Polypodium_amorphum
A-2073213-Polypodium_amorphum
-2003047-Cystopteris_protrusa
-2007507-Cystopteris_protrusa
-2012306-Cystopteris_protrusa
038519-Heterosiphonia_pulchra
042241-Selaginella_kraussiana
A|LP-ZZOL-2000160-Selaginella

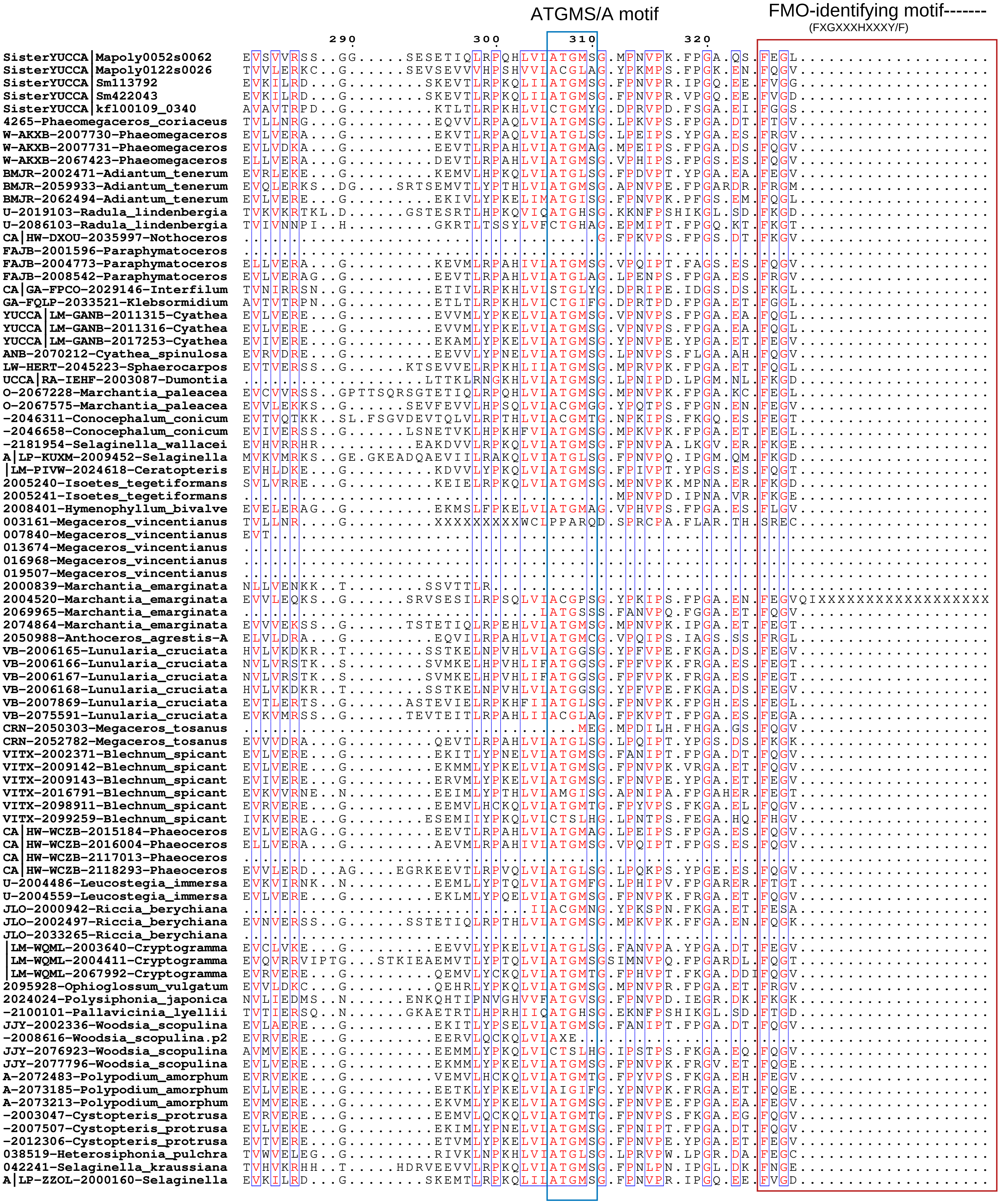
ATGMS/A motif
FMO-identifying motif-------
(FXGXXXHXXXY/F)
290
300
310
320
SisterYUCCA|Mapoly0052s0062
SisterYUCCA|Mapoly0122s0026
SisterYUCCA|Sm113792
SisterYUCCA|Sm422043
SisterYUCCA|kfl00109_0340
4265-Phaeomegaceros_coriaceus
W-AKXB-2007730-Phaeomegaceros
W-AKXB-2007731-Phaeomegaceros
W-AKXB-2067423-Phaeomegaceros
BMJR-2002471-Adiantum_tenerum
BMJR-2059933-Adiantum_tenerum
BMJR-2062494-Adiantum_tenerum
U-2019103-Radula_lindenbergia
U-2086103-Radula_lindenbergia
CA|HW-DXOU-2035997-Nothoceros
FAJB-2001596-Paraphymatoceros
FAJB-2004773-Paraphymatoceros
FAJB-2008542-Paraphymatoceros
CA|GA-FPCO-2029146-Interfilum
GA-FQLP-2033521-Klebsormidium
YUCCA|LM-GANB-2011315-Cyathea
YUCCA|LM-GANB-2011316-Cyathea
YUCCA|LM-GANB-2017253-Cyathea
ANB-2070212-Cyathea_spinulosa
LW-HERT-2045223-Sphaerocarpos
UCCA|RA-IEHF-2003087-Dumontia
O-2067228-Marchantia_paleacea
O-2067575-Marchantia_paleacea
-2046311-Conocephalum_conicum
-2046658-Conocephalum_conicum
-2181954-Selaginella_wallacei
A|LP-KUXM-2009452-Selaginella
|LM-PIVW-2024618-Ceratopteris
2005240-Isoetes_tegetiformans
2005241-Isoetes_tegetiformans
2008401-Hymenophyllum_bivalve
003161-Megaceros_vincentianus
007840-Megaceros_vincentianus
013674-Megaceros_vincentianus
016968-Megaceros_vincentianus
019507-Megaceros_vincentianus
2000839-Marchantia_emarginata
2004520-Marchantia_emarginata
2069965-Marchantia_emarginata
2074864-Marchantia_emarginata
2050988-Anthoceros_agrestis-A
VB-2006165-Lunularia_cruciata
VB-2006166-Lunularia_cruciata
VB-2006167-Lunularia_cruciata
VB-2006168-Lunularia_cruciata
VB-2007869-Lunularia_cruciata
VB-2075591-Lunularia_cruciata
CRN-2050303-Megaceros_tosanus
CRN-2052782-Megaceros_tosanus
VITX-2002371-Blechnum_spicant
VITX-2009142-Blechnum_spicant
VITX-2009143-Blechnum_spicant
VITX-2016791-Blechnum_spicant
VITX-2098911-Blechnum_spicant
VITX-2099259-Blechnum_spicant
CA|HW-WCZB-2015184-Phaeoceros
CA|HW-WCZB-2016004-Phaeoceros
CA|HW-WCZB-2117013-Phaeoceros
CA|HW-WCZB-2118293-Phaeoceros
U-2004486-Leucostegia_immersa
U-2004559-Leucostegia_immersa
JLO-2000942-Riccia_berychiana
JLO-2002497-Riccia_berychiana
JLO-2033265-Riccia_berychiana
|LM-WQML-2003640-Cryptogramma
|LM-WQML-2004411-Cryptogramma
|LM-WQML-2067992-Cryptogramma
2095928-Ophioglossum_vulgatum
2024024-Polysiphonia_japonica
-2100101-Pallavicinia_lyellii
JJY-2002336-Woodsia_scopulina
-2008616-Woodsia_scopulina.p2
JJY-2076923-Woodsia_scopulina
JJY-2077796-Woodsia_scopulina
A-2072483-Polypodium_amorphum
A-2073185-Polypodium_amorphum
A-2073213-Polypodium_amorphum
-2003047-Cystopteris_protrusa
-2007507-Cystopteris_protrusa
-2012306-Cystopteris_protrusa
038519-Heterosiphonia_pulchra
042241-Selaginella_kraussiana
A|LP-ZZOL-2000160-Selaginella

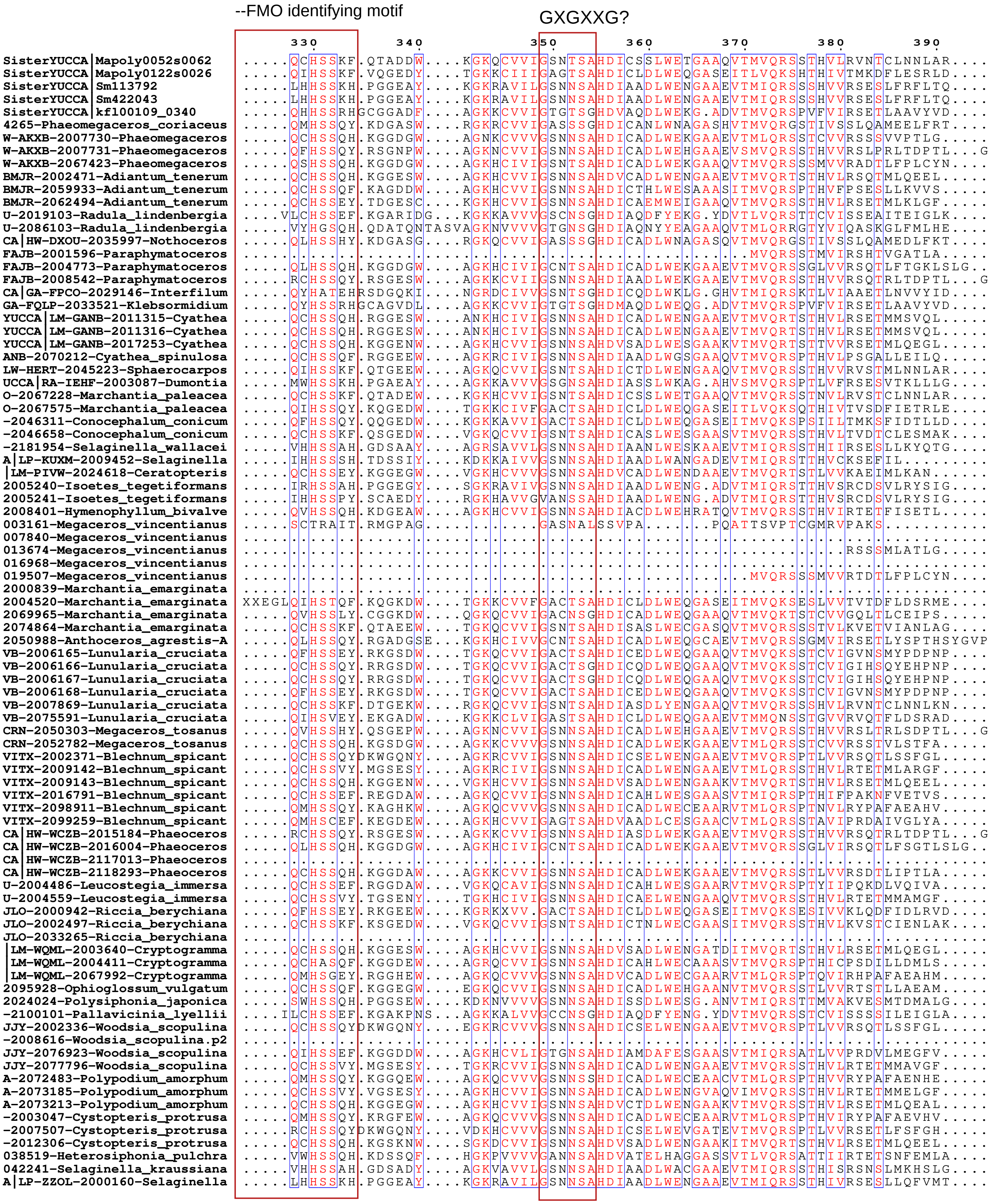
--FMO identifying motif
GXGXXG?
330
340
350
360
370
380
390

```
                                   400       410       420       430       440       450       460
SisterYUCCA|Mapoly0052s0062     DGYGEG....GPPVDTVDLAFASVPYKIF.HSFAIPAMQRIAKAEAPFYEGLSKAGFVLDWGADGSGLFTKYLRRGGG
SisterYUCCA|Mapoly0122s0026     PQLEKAELDTNFDTDAFDLKSASLPYKVL.PRLNVPIYARIAQKEAELYDKLHKAGFKTFMGEDESGLILFLLRRAAG
SisterYUCCA|Sm113792            EIYSESAVESGITTDKADMIFASLPYKIM.GDAQRCNSDAIRAHDKDFYAALTNTGFMLDFGDDDSGLFMKYLRRGSG
SisterYUCCA|Sm422043            EIYSESAVESGITTDKADMIFASLPYKIM.GDAQRCNSDAIRAHDKDFYAALTKTGFMLDFGDDDSGLFMKYLRRGSG
SisterYUCCA|kfl00109_0340       PLFHDSS...PMSQQHADLLMVSTPYRLV.APFQAPMVEEIKRRDGDLHERLRERGFRVHFAEDGSGLAGLLNRTATG
4265-Phaeomegaceros_coriaceus   IGFSEENIDTPSGSTKCDLVAVSAPYKLM.RDMHLPLVRMMQKRDSMLVESLRKIGFQCDYCEDEAGLFANYARELSG
W-AKXB-2007730-Phaeomegaceros   SLYAEGL...PFSTDTADLLNASLPYKLF.PRVHIPVYQRVAELDADLLHRLRKVGFLLDFGDDGSGMFVKYLRRGSG
W-AKXB-2007731-Phaeomegaceros   GLYKEDS...LFSTDTCDLLIASLPYKLL.PRLHIPVYQRIAQQDADLYDRLRKAGFLLDFGEDGSGSFVQYLRRGSG
W-AKXB-2067423-Phaeomegaceros   GLYTGKP...CPPPDTADLILASIPHKLF.ARLQIPFTDTIREMDAAFYSKLSKSGFQLDFGEDGTGIQVKYLRRGSG
BMJR-2002471-Adiantum_tenerum   HPYREGS...KYSTDDADILFASIPYKIF.HKAQIPVYQKIADIDAEFYARLKESGFMLDFGEDGSGLFTKYLRRGSG
BMJR-2059933-Adiantum_tenerum   AFFGG.....KYSTAEGDLLAASTPYRLL.PQVHIPVARKAAADHAELYKELEKVGFMHTFGDDGSGMMPLYLRRASG
BMJR-2062494-Adiantum_tenerum   GEYSENP...RHSIDDCDLLYASVPYKLF.HRDLIPACKRLQEIDAEFYRRIKDCGFLLDFGDDESGVLLKFLRRFSG
U-2019103-Radula_lindenbergia   GVYSE....DGPPVDDADMWLHSLPSEVL.KALQVNITAAQAEFDKKTLDGLTKAGFKLDRGPEDAGLFMKYFQRGGG
U-2086103-Radula_lindenbergia   GLYDE....TGPPIEDADVYGQSLPIAVQ.FALNVGLTERIKAAEKENIEGLEKAGFKLDFGHDGSGIYRKYITRGGG
CA|HW-DXOU-2035997-Nothoceros   IGFSEENSESNSGSTKCDLDAVSAPYKLM.REMHLPLVRVMQKRDSMLIEGLRKIGFQCDYCEDEAGLFANYARNFSG
FAJB-2001596-Paraphymatoceros   GLYAEDL...PFSTETADLIVASVPYKIF.PRLQIPVYQRLAEVDADLLDRLRKVGFMLDYGDDGSGLFAKFLRRGSG
FAJB-2004773-Paraphymatoceros   GLFVENA...RFSPDTADLILASIPHKVL.TGLQVQFTQQIREADAPFYDRLQKSGFQLDFGEDGSGLLLKYLRRGSG
FAJB-2008542-Paraphymatoceros   GLYREDS...PFSTDTADLLVASIPYKLL.PRRHIPVYQRIAQQDADLYDRLTKAGFLLNFGEDGSGGFVMYLRRGSG
CA|GA-FPCO-2029146-Interfilum   PLFLEES...PFTREQADVLMVSTPYRLL.APFQMPMVQEAAKQDADLHERLRKVGFELSFAEDGSGVGGLLNRTATG
GA-FQLP-2033521-Klebsormidium   PLFHDSS...PLSQQHADLPMVSTPYRLI.APFQAPIVEEVKRRDGDLHERLREPGFRFHFAEDGSGLAGLLNRTATG
YUCCA|LM-GANB-2011315-Cyathea   SLYREGS...KXMPT...FLLPLHPTKSY.TKLRYLCTKGLQTLMLISMGALRKL.......................
YUCCA|LM-GANB-2011316-Cyathea   SLYREGS...KYSTDDADLLVASTPYKIL.HKAQIPVYKRIADIDADFYGRLKKAGFMLDFGEDGSGLLLKYLRRGSG
YUCCA|LM-GANB-2017253-Cyathea   VFYREGS...KYSTSDADILLASRPYRIF.HKFEAAVYRKIAEIDAEFYTRLEERGFLLDFGEDRSGHFTKYLRRGSG
ANB-2070212-Cyathea_spinulosa   PFYRERS...KHSTADNDLLLAACPYKLF.PQLQIPFLNKIAQTYGEFYHRLLNAGFMHTFGEDGSGAFSMYMRRGSG
LW-HERT-2045223-Sphaerocarpos   DGYGEG....GPPVDTVDVGFASVPYKLF.PSFSIPGVQKIAKDESTFYEGLSKAGFELSWGEDGSGLFTMYLRRGGG
UCCA|RA-IEHF-2003087-Dumontia   SLYSEEAVANGITTEVADLIFASIPYQML.GELHKTVCKQVKQRDAEFYEKLREREFRLSFGPDESGLFLMYLRRGSG
O-2067228-Marchantia_paleacea   DGYGEG....GPPVDTVDLTFASLPYKIF.HRFMIPVMQQIAKAESSFYEGLTKAGFVLDWGDDGSGLFPKYLRRGGG
O-2067575-Marchantia_paleacea   PHLEGAEYDTNFDSDAFDLKTASLPFKVL.PRVQIPVYAGIAKKEADLYDRLLKAGFKTFLGEDESGIPLMLLRRGGG
-2046311-Conocephalum_conicum   PDLQGKEFDHDFDMDSYDIKGASLPFKVM.PRVLVPVYNSIAEKEAELHSKLRKMGFDLHMGEDGSGLPMALVRRAGG
-2046658-Conocephalum_conicum   DGYGEG....GPPVDDTDLKMASVPYKVF.PHLVIPLFQQLAKEQAPFYERLSKAGFVLDWGDDGSGLFAKYLRRGGG
-2181954-Selaginella_wallacei   PLYSEEAVAQGITTDVADMMFASVPMRLV.HLMFKTVVEGIKKEDADFYSALKASGFMLDFGVDETGLFPKYLRRGSG
A|LP-KUXM-2009452-Selaginella   ..............................................................................
|LM-PIVW-2024618-Ceratopteris   ELYQEGS...KYSTEDADILFASVPYKIF.HQPSILLYKKIKEWDAEFYSRLRESGFMLDFGEDESGLFPKYLRRGSG
2005240-Isoetes_tegetiformans   ELYSESAVESGITTDLADMIFASIPYKIM.TEFQRPIIDKIRAQDADFYKDLQKSGFMLDFGDDETGLFMKYLRRGSG
2005241-Isoetes_tegetiformans   ELYSESAVESGITTDLADMIFASIPYKIM.TEFQRPIIDKIRAQDADFYKDLQKSGFMLDFGDDETGLFMKYLRRGSG
2008401-Hymenophyllum_bivalve   KHYSEDA...VYSTESADLFFASFPNKLV.PRFQIPANSKIAEMDANFYERLKRAGFLLDFGEDGSGLFGKYLRRGSG
003161-Megaceros_vincentianus   ..........QWSSAAPRLFPACKP.....................................................
007840-Megaceros_vincentianus   ..............................................................................
013674-Megaceros_vincentianus   SLYAEGL...PLSTDTADILMASLPYKVF.PRIQVTVHERVAELDADLWDRLRRVGFLLDLGDDGSGMFVKYLRRGSG
016968-Megaceros_vincentianus   ..............................................................................
019507-Megaceros_vincentianus   GLYTGKP...CHTPDTADLIMASIPQKLF.ARLQIPFTNSLREMDASFYADLTKSGFQVDFGEDGSGIQVKYLRRGSG
2000839-Marchantia_emarginata   ..............................................................................
2004520-Marchantia_emarginata   PDLKGGEYDYTFDTDAFDIKTASLPFKVL.PRVQIPVYAKISEKEAELHQRLNKAGFKTWMGEDKSGIPLMLLRRAGG
2069965-Marchantia_emarginata   GMENEDN.SSTSAVDRMDLRMICVPYKLR.ARLDIPFYEQLAKTHADFYNRLTKAGFKIDFGEDGSGLLMKLFRRGGG
2074864-Marchantia_emarginata   DGYGEG....GPPVDTVDLTFASLPYKIF.HRFMIPVMQRIAKEQASFYEGLAKAGFALDWGEDGSGLFPKYLRKGGG
2050988-Anthoceros_agrestis-A   GLFVHDS...PWTLDTADLMLASFPHKVL.PRLQTLFSQAIQEQDAAFYARLKKSGFQLDFGEDGSGLLYKYYRRGSG
VB-2006165-Lunularia_cruciata   AAFVDAP...LLAAENFDLLSSSNPYKVL.TQQCVPMVKAIAEKDAKFYEDLNKAGFKTCFGEDNTGLPMLFHRRLGG
VB-2006166-Lunularia_cruciata   EVMQDAP...LATLERFEPMLFATPYKVM.HRITIPITKKIAENDAEFYSDLAKVGFRTDFGEDGSGLLMKLLRRQGG
VB-2006167-Lunularia_cruciata   EVMQDAP...LATLERFEPMLFATPYKVM.HRITIPITKKIAENDAEFYSDLAKVGFRTDFGEDGSGLLMKLLRRQGG
VB-2006168-Lunularia_cruciata   AAFVDAP...LLAAENFDLLSSSNPYKVL.TQQCVPMVKAIAEKDAKFYEDLNKAGFKTCFGEDNTGLPMLFHRRLGG
VB-2007869-Lunularia_cruciata   EGYCEG....GPPVDTVDLTYASIPYKVF.PEFAKVGMKRIAQEEAPFYERLSKSGYVLDFGDDGSGLWLKYLRRGGG
VB-2075591-Lunularia_cruciata   ADLPGTENDRAFDIDAFDLKSASLPFKVL.TRMMIGTYTKIAERDAKFLEDLKKVGFQWTMGEDGSGLSLMLIRRGGG
CRN-2050303-Megaceros_tosanus   GLYKEDS...LFSTETCDLLLFSIPYKLL.PRQHIPVFQRIALRDADLYNRLRNAGFLLDFGEDGSGAFVKYLRRGSG
CRN-2052782-Megaceros_tosanus   SLYSESA...PFPTDTADLLMASFPYKLF.SQQQIPVYQRVAELDADLLDRLKKVGFLLDFGEDGSGMFVKYLRRGSG
VITX-2002371-Blechnum_spicant   GPFCENS...KYSIEDADLLFASTPYKIF.HRDHILATKMLQIVDSEFYARLKDTGFLLNFGEDKSGLLPMYLRRGSG
VITX-2009142-Blechnum_spicant   GEYREDP...KYSIDDVDLLFASTPYKIF.PRESMPACKQLQETDTEFYGRLKQCGFMLDFGEDESGLFIKYLRRGSG
VITX-2009143-Blechnum_spicant   VFYREGS...KYSTSDADILFASIPYKIF.HKFQIPVYRKIGEIDAKFYARLEERGFLLDFGEDGSGLFTKYLRRGSG
VITX-2016791-Blechnum_spicant   TFFIG.....ERRTSEGDLLLASAPYKLL.PRIHTQVAKKALKENGELYKGLEEVGFMHTFGEDGSGMVPLYLRRGSG
VITX-2098911-Blechnum_spicant   PFYCADS...KYSTSEADLVLASFPHKLF.PRLQAPLYTKFAETDKAFYQSLADVGFMHSFGEDNSGIFSMYYRRGSG
VITX-2099259-Blechnum_spicant   PIYNS.....KYSTAEGDLLVVAYPHKLL.ARLQGPAYKQIGEKYAEFYRGLQEAGFKHSFGEDGAGLYAMYLRGESG
CA|HW-WCZB-2015184-Phaeoceros   GLYREDS...PFSTDTADLLVASIPYKLL.PRHHIPVHQRIAQRDADLYGRLTKAGFLLNFGEDGSGGFVMYLRRGSG
CA|HW-WCZB-2016004-Phaeoceros   GLYAEDS...RFSPDTADLIMSSIPHKVL.TKLQIQFTQQIRDLDAPFYDRLQKSGFQVDFGEDGSGILLKYLRRGSG
CA|HW-WCZB-2117013-Phaeoceros   ..............................................................................
CA|HW-WCZB-2118293-Phaeoceros   GLYAEGL...PFSTDTADLISASLPYKLF.PRVHIPVYQKVAEVDADLLDRLRKVGFMLDYGDDGSGLFIKYLRRGSG
U-2004486-Leucostegia_immersa   PVFKG.....ERTTSEADLLLASSPYKLIGAGIHKEGSERAAKENEELYKGLEEVGFMHTFGDDGSGLIPLYVRRASG
U-2004559-Leucostegia_immersa   GEFREDP...KYSIDDADLLFASTPYKIF.HRILMPACQRLQEIDAEFYGRLKQCGFMLDFGDDESGLLPKYLRRGSG
JLO-2000942-Riccia_berychiana   PDFQGTEIDGGFDIDSFDIKSASLPYKVL.PRVLVPVYNRIAEKEAEFYDRLRKVGFRLHMGEDGTGISLGLVRRGGG
JLO-2002497-Riccia_berychiana   DGYGEG....GPPVDDSDLRFASVPYKIF.HRFAIPAMQQIAKQHAQFYERLSNAGFVLDWGDDGSGLFTKYLRRGGG
JLO-2033265-Riccia_berychiana   ..............................................................................
|LM-WQML-2003640-Cryptogramma   AFYREGS...KYCTSDADILFASIPYKLF.HRMEIPAYQKIAEIDAEFYARLKKSGFMLDFGEDGSGLFTKYLRRGSG
|LM-WQML-2004411-Cryptogramma   PFFAGK....HYITADADLLLASFPYKLL.HRVHTPVAKKAAERHAELYRDLEKVGFMHTFGDDGSGMLPLYLRRGSG
|LM-WQML-2067992-Cryptogramma   PFYCADS...KYSTVEADLLLASYPYKLF.PLIQIPLYKKFAESDKSFYEGLSEVGFMHTFGEDHSGLFSMYYRRGSG
2095928-Ophioglossum_vulgatum   GNYVEGS...PYSTTDADFLFASVPYKIF.HLFHIPVYKRIAKLDAAFYESLSKAGFLLDFGEDGSGVLTKYMRRGSG
2024024-Polysiphonia_japonica   GLYSEAAVKRGDVADVSDLIFASIPYKIL.PQGQKQLYEKMRVKDGEFYKQLECAGFMLDFGVDDTGLFMKYLRRGSG
-2100101-Pallavicinia_lyellii   GLYDE....TGPPVEDADVFFHSIPNSVL.KSIHVDVTAA......................................
JJY-2002336-Woodsia_scopulina   GPFCENS...KYSIEDADLLFASTPYKIF.HRDHILATKMLQIVDSQFYARLKDTGFLLDFGEDKSGLFPKYLRRGSG
-2008616-Woodsia_scopulina.p2   ..............................................................................
JJY-2076923-Woodsia_scopulina   PIYKS.....KYSTAEADLLVASVPHKVF.PRQQRQMYKMIGEMCADFYKGLREAGFEHSFGEDGAGFFAMFLRRASG
JJY-2077796-Woodsia_scopulina   GEYREDP...KYSIDDADLLFASTPYKIF.HRNLMPACQRLQEMDAEFYNRLKQCGFMLDFGEDESGLFLKYLRRGSG
A-2072483-Polypodium_amorphum   PYYCADA...KYSTTEADLLEASYPYKIY.PQTQIPQYKKFAERDKTFYEALAQVGFMHTFGEDHSGLFSMYYRRGSG
A-2073185-Polypodium_amorphum   GGYCEDP...RYSIDDADLLFASTPYKIF.HRFLMPVCKLVQEIDAEFYGRLKQCGFMLDFGDDESGVLPKYLRRGSG
A-2073213-Polypodium_amorphum   VLYREGS...KLSTSDADIIFASIPCKVF.HRAEIPVYKKIAEIDAEFYARLEESGFMLDFGEDGSGLFAKYLRRGSG
-2003047-Cystopteris_protrusa   PFYCADS...KYSTSEADLISASYPHKIF.PRTHVPFYIKFAETDKAFYESLAHRGFMHTFGEDHSGIFPMYYRRGSG
-2007507-Cystopteris_protrusa   GPYGENS...RYSVEDADLLLASTPFRIF.HRDHIPITKMLQIIDSDFYMRLKHSGFLLDFGEDESGAFPKYLRRGSG
-2012306-Cystopteris_protrusa   AFYREGS...KYSTSDADILFASIPYKLF.HKFQIPLYRKIAEIDAKFYARLEECGFLLDFGEDGSGLFTKYLRRGSG
038519-Heterosiphonia_pulchra   ELYSEEAVKKGVTVEKADLLFATTPYALL.SGRHQVLYERIRKKDKGLYDGLREVGFKVDFGEDESGHFLKYLRRGSG
042241-Selaginella_kraussiana   ALYSEAAAANGITTEKADMMFASVPYKQM.PEKYRLQMAAIKKDDSEFYKKLVAAGFLLDYGADDTGLFLKYLRRGSG
A|LP-ZZOL-2000160-Selaginella   NTYSESAVESGLTTDKADMIFASVPYKIM.GDVNRPTYDAIRAHDKDFYAALTKTGFMLDFGDDDSGLFMKYLRRGSG
```

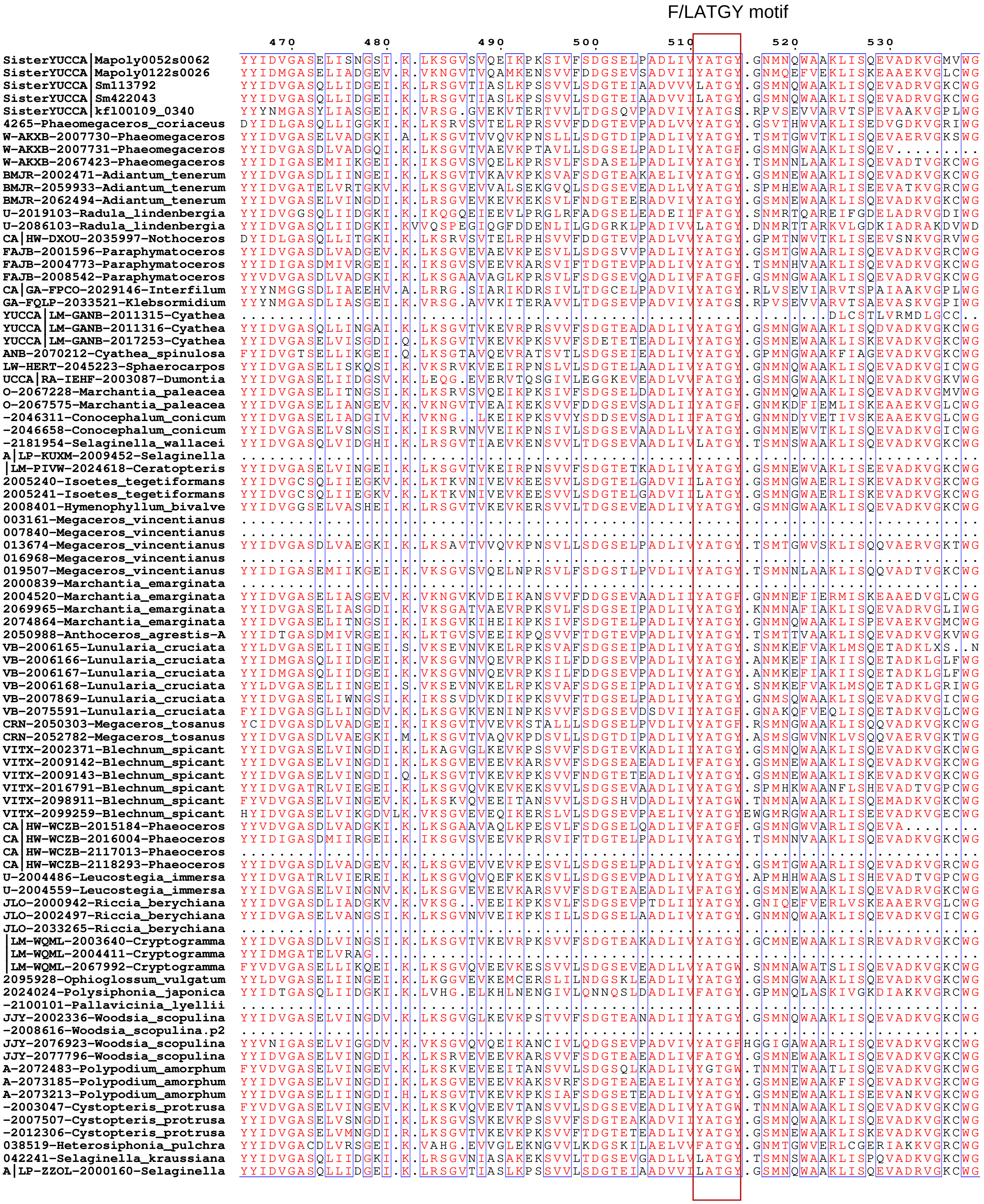
F/LATGY motif
470
480
490
500
510
520
530
SisterYUCCA|Mapoly0052s0062
SisterYUCCA|Mapoly0122s0026
SisterYUCCA|Sm113792
SisterYUCCA|Sm422043
SisterYUCCA|kfl00109_0340
4265-Phaeomegaceros_coriaceus
W-AKXB-2007730-Phaeomegaceros
W-AKXB-2007731-Phaeomegaceros
W-AKXB-2067423-Phaeomegaceros
BMJR-2002471-Adiantum_tenerum
BMJR-2059933-Adiantum_tenerum
BMJR-2062494-Adiantum_tenerum
U-2019103-Radula_lindenbergia
U-2086103-Radula_lindenbergia
CA|HW-DXOU-2035997-Nothoceros
FAJB-2001596-Paraphymatoceros
FAJB-2004773-Paraphymatoceros
FAJB-2008542-Paraphymatoceros
CA|GA-FPCO-2029146-Interfilum
GA-FQLP-2033521-Klebsormidium
YUCCA|LM-GANB-2011315-Cyathea
YUCCA|LM-GANB-2011316-Cyathea
YUCCA|LM-GANB-2017253-Cyathea
ANB-2070212-Cyathea_spinulosa
LW-HERT-2045223-Sphaerocarpos
UCCA|RA-IEHF-2003087-Dumontia
O-2067228-Marchantia_paleacea
O-2067575-Marchantia_paleacea
-2046311-Conocephalum_conicum
-2046658-Conocephalum_conicum
-2181954-Selaginella_wallacei
A|LP-KUXM-2009452-Selaginella
|LM-PIVW-2024618-Ceratopteris
2005240-Isoetes_tegetiformans
2005241-Isoetes_tegetiformans
2008401-Hymenophyllum_bivalve
003161-Megaceros_vincentianus
007840-Megaceros_vincentianus
013674-Megaceros_vincentianus
016968-Megaceros_vincentianus
019507-Megaceros_vincentianus
2000839-Marchantia_emarginata
2004520-Marchantia_emarginata
2069965-Marchantia_emarginata
2074864-Marchantia_emarginata
2050988-Anthoceros_agrestis-A
VB-2006165-Lunularia_cruciata
VB-2006166-Lunularia_cruciata
VB-2006167-Lunularia_cruciata
VB-2006168-Lunularia_cruciata
VB-2007869-Lunularia_cruciata
VB-2075591-Lunularia_cruciata
CRN-2050303-Megaceros_tosanus
CRN-2052782-Megaceros_tosanus
VITX-2002371-Blechnum_spicant
VITX-2009142-Blechnum_spicant
VITX-2009143-Blechnum_spicant
VITX-2016791-Blechnum_spicant
VITX-2098911-Blechnum_spicant
VITX-2099259-Blechnum_spicant
CA|HW-WCZB-2015184-Phaeoceros
CA|HW-WCZB-2016004-Phaeoceros
CA|HW-WCZB-2117013-Phaeoceros
CA|HW-WCZB-2118293-Phaeoceros
U-2004486-Leucostegia_immersa
U-2004559-Leucostegia_immersa
JLO-2000942-Riccia_berychiana
JLO-2002497-Riccia_berychiana
JLO-2033265-Riccia_berychiana
|LM-WQML-2003640-Cryptogramma
|LM-WQML-2004411-Cryptogramma
|LM-WQML-2067992-Cryptogramma
2095928-Ophioglossum_vulgatum
2024024-Polysiphonia_japonica
-2100101-Pallavicinia_lyellii
JJY-2002336-Woodsia_scopulina
-2008616-Woodsia_scopulina.p2
JJY-2076923-Woodsia_scopulina
JJY-2077796-Woodsia_scopulina
A-2072483-Polypodium_amorphum
A-2073185-Polypodium_amorphum
A-2073213-Polypodium_amorphum
-2003047-Cystopteris_protrusa
-2007507-Cystopteris_protrusa
-2012306-Cystopteris_protrusa
038519-Heterosiphonia_pulchra
042241-Selaginella_kraussiana
A|LP-ZZOL-2000160-Selaginella

```
540 550 560 570 580
SisterYUCCA|Mapoly0052s0062 LGSNTTNDPGPW.............EKELRNMWKPTQQPGL...........W..FMG.GNLHQC.....RHYSLVVA
SisterYUCCA|Mapoly0122s0026 LGSGFKGDPGPW.............VGELKNMYRPLPVPAL...........W..ILS.GGLSHC.....RFYSRHVA
SisterYUCCA|Sm113792 LGSNTRKDPGPW.............EGELRNMWKPTAQEGL...........W..FHG.GNLHQA.....RHYSLYLA
SisterYUCCA|Sm422043 LGSNTRKDPGPW.............EGELRNMWKPTAQEGL...........W..FHG.GNLHQA.....RHYSLYLA
SisterYUCCA|kf100109_0340 YGSGLDGDPGPW.............EGELRNIYKPTAQEGL...........W..FHI.GGLVQT.....RFGSLLVA
4265-Phaeomegaceros_coriaceus LGSGFGKDVGPW.............EGEIRNMWKPTHQPGL...........W..FHG.GGLNHT.....RIFSAYLA
W-AKXB-2007730-Phaeomegaceros LGSGTTKDPGPW.............ERELRNMWKPTQQPAL...........W..FHA.GNLHQA.....RHYSLVLA
W-AKXB-2007731-Phaeomegaceros ..........................................................................
W-AKXB-2067423-Phaeomegaceros LGSGTARDPGPW.............ERELRNMWKPTQQPGL...........W..FHG.GTVQEN.....RHFSLVLA
BMJR-2002471-Adiantum_tenerum LGSGTRKDPGPW.............ERELRNMWKPTKQEGL...........W..FHG.GNLHQS.....RHYSLMLA
BMJR-2059933-Adiantum_tenerum LGSNTANDPGPW.............ERELRNMWKPTPQPNL...........W..FHG.GGFQEA.....RHFSLYLA
BMJR-2062494-Adiantum_tenerum LGSGTTKDPGPW.............ERELRNMWKPTQQENL...........W..FHG.GNLQQA.....RHYSLLLA
U-2019103-Radula_lindenbergia YDA......................EGEMRTIWRKSGHPGF...........W..FMG.GNLALC.....RYYSKVLA
U-2086103-Radula_lindenbergia LDE......................EGEVNAMWRPCGHPRL...........W..FMG.GSLALC.....RTYSRFVA
CA|HW-DXOU-2035997-Nothoceros LGSGFGKDTGPW.............EGEIRNMWKPTHQPGL...........W..FHG.GGLNHT.....RIFSAYLA
FAJB-2001596-Paraphymatoceros LGSSTTKDPGPW.............ERELRNMWKPTQQAGL...........W..FHG.GNLQQS.....RHYSLMLA
FAJB-2004773-Paraphymatoceros LGSGTARDPGPW.............ERELRNMWKPTHQEGL...........W..FHG.GTVQEN.....RHFSLVLA
FAJB-2008542-Paraphymatoceros LGSATTKDPGPW.............ERELRNMWKPTQQPGL...........W..FQG.GNLLQA.....RYYSLMLA
CA|GA-FPCO-2029146-Interfilum YGSGLDGDKGPW.............EGELRNVFKPTAQNGL...........W..FFT.GGLVQT.....RFFSLMLA
GA-FQLP-2033521-Klebsormidium YGSGLDGDPGPW.............EGELRNIYKPTSQEGL...........W..FHI.GGLVQT.....RFGSLIVA
YUCCA|LM-GANB-2011315-Cyathea .SSTCVEDQGTI.............L.....MW...............................................
YUCCA|LM-GANB-2011316-Cyathea LGSDTRKDPGPW.............ERELRNMWKPTQQCGL...........W..FHG.GNLHQS.....RHYSLPLA
YUCCA|LM-GANB-2017253-Cyathea LGSGTRKDPGPW.............ERELRNMWKPTQQEGL...........W..FHG.GNLHQS.....RHYSLMLA
ANB-2070212-Cyathea_spinulosa LGSDTAKDPGPW.............ERELRNMWKPTHQPSL...........W..FHG.GSFQES.....RAYSLYLA
LW-HERT-2045223-Sphaerocarpos LGSSTTNDPGPW.............ERELRNMWKPTQQEGL...........W..FMG.GNLHQC.....RYFSLPLA
UCCA|RA-IEHF-2003087-Dumontia LGSNTVKDPGPW.............EGELRNMWKPTQQEGL...........W..FQG.GNLHQA.....RHYSRFLA
O-2067228-Marchantia_paleacea LGSNTTNDPGPW.............EKELRNMWKPTQQPGL...........W..FMG.GNLHQC.....RHYSLVVA
O-2067575-Marchantia_paleacea LGSDTKGDPGPW.............VGELKNMYKPLAVPAL...........W..VAS.GGFSQC.....RFYSRHVA
-2046311-Conocephalum_conicum LGSNTNKDPGPW.............EGELKNLWKPLPVPNL...........W..MTS.GGFSQS.....RFHSRHLA
-2046658-Conocephalum_conicum LGSNTTGDPGPW.............EGELRNMWKPTQQPAL...........W..FMG.GNLHQC.....RHFSLPLA
-2181954-Selaginella_wallacei YGSDTPKDPGPW.............VGELRNMWKPTKQEAL...........W..FHG.GNLHQV.....RHYSLYLA
A|LP-KUXM-2009452-Selaginella ..............................................................................
|LM-PIVW-2024618-Ceratopteris LGSDTRKDPGPW.............ERELRNMWKPTRQEAL...........W..FQG.GNLHQS.....RHYSLMLA
2005240-Isoetes_tegetiformans LGSDTTKDPGPW.............EGELRNMWKPTRQEAL...........W..FHG.GNLHQA.....RHYSLYVA
2005241-Isoetes_tegetiformans LGSDTTKDPGPW.............EGELRNMWKPTRQEAL...........W..FHG.GNLHQA.....RHYSLYVA
2008401-Hymenophyllum_bivalve LGSNTRKDPGPW.............ERELRNMWKPTKQPCL...........W..FHG.GNFHQA.....RHYSLLLA
003161-Megaceros_vincentianus ....................................................W.................RTFSR...
007840-Megaceros_vincentianus ..............................................................................
013674-Megaceros_vincentianus LGSGTTKDPGPW.............ERELRNMWKPTQQPGL...........W..FHG.GNLHQA.....RHYSLILG
016968-Megaceros_vincentianus ..............................................................................
019507-Megaceros_vincentianus LGSGTFRDPGPW.............ERELRNMWKPTHQPGL...........W..FHG.GTVQEN.....RHFSLVLA
2000839-Marchantia_emarginata ..............................................................................
2004520-Marchantia_emarginata LGSDTKGDPGPW.............VGELKNMWKPLPVPAL...........W..IMSGGGFSHC.....RFYSRHLA
2069965-Marchantia_emarginata LGSGTTNDPG....................................................................
2074864-Marchantia_emarginata LGSNTSNDPGPW.............EKELRNMWKPTQQPGL...........W..FMG.GNLHQC.....RHYSLVLA
2050988-Anthoceros_agrestis-A LGSGTARDPGPW.............ERELRNMWKPTHQEGL...........W..FHG.GAVQES.....RSFSLVLA
VB-2006165-Lunularia_cruciata LGSRFKHRP.............................................................RSRALG
VB-2006166-Lunularia_cruciata LGSNTTNDPGPW.............VGEHRNIFKPLSQPNL...........W..MTG.GSFSPT.....RMMSRYLS
VB-2006167-Lunularia_cruciata LGSNTTNDPGPW.............VGEHRNIFKPLSQPNL...........W..MTG.GSFSPT.....RMMSRYLS
VB-2006168-Lunularia_cruciata LGSNTAHDPGPW.............VGEYRNVWKPTAQANF...........W..TTG.AAINQA.....RQMSRYLA
VB-2007869-Lunularia_cruciata LGSNTTNDPGPW.............EKELRNMWKPTQQPGL...........W..FMG.GNLHQC.....RHFSACLA
VB-2075591-Lunularia_cruciata LGSNTKGDPGPW.............EGELRNMYKPTAQPGL...........W..IAS.GGFSHA.....RFYSRYLA
CRN-2050303-Megaceros_tosanus LGSATAYDPGPLGTGASKHVEAHPSAGIVVPRWQPSPSQVL...........LAPFST.PDQSKVR....RHSDASLR
CRN-2052782-Megaceros_tosanus LGSGTAKDPGPW.............EGELRNMWKPTQQPGL...........W..FHG.GNLHQS.....RHYSLILA
VITX-2002371-Blechnum_spicant LGSDTTKDPGPW.............EGELRNMWKPTQQPHL...........W..FHG.GNLQQS.....RHFSLSLG
VITX-2009142-Blechnum_spicant LGSGTRKDPGPW.............ESELRNMWKPTQQDGL...........W..FHG.GNLHQS.....RHYSLMLA
VITX-2009143-Blechnum_spicant LGSGTRKDPGPW.............ERELRNMWKPTQQEGL...........W..FHG.GNLHQS.....RHYSLVLA
VITX-2016791-Blechnum_spicant LGSNTTKDPGPW.............ERELRNMWKPTLQQNL...........W..FHG.GGFHEA.....RHYSLYLA
VITX-2098911-Blechnum_spicant LGSGTTKDPGPW.............EREIRNMWKPTRHPSL...........W..FHG.GGFQET.....RHFSLYLA
VITX-2099259-Blechnum_spicant LGSGTIHDPGPW.............EGELRNMYKPTRQQNL...........W..FHA.GGLFDS.....RIHSQYLA
CA|HW-WCZB-2015184-Phaeoceros .........................................................A.G..................
CA|HW-WCZB-2016004-Phaeoceros LGSGTTRDPGPW.............ERELRNMWKPTHQEGL...........W..FHG.GTVQEN.....RHFSLVLA
CA|HW-WCZB-2117013-Phaeoceros ..............................................................................
CA|HW-WCZB-2118293-Phaeoceros LGSSTTKDPGPW.............ERELRNMWKPTQQPGL...........W..FHG.GNLHQS.....RHYSLMLA
U-2004486-Leucostegia_immersa LGSNTTKDPGPW.............QRELRNMWKPTLQQNL...........W..FHG.GGFQEA.....RHFSLYLA
U-2004559-Leucostegia_immersa LGSGTSKDLGPW.............ERELRNMWKPTQQEGL...........W..FHG.GNLQQS.....RHYSLSLA
JLO-2000942-Riccia_berychiana VGSNTKLDPGPW.............EGELRNMWKPLAVPXXXXXXXWRGLQPWQILQS.SCCSSAESQNGRHSNSCLQ
JLO-2002497-Riccia_berychiana LGSGTTNDPGPW.............ERELRNMWKPTQQPGL...........W..FMG.GNLHQC.....RHFSLPLA
JLO-2033265-Riccia_berychiana ..............................................................................
|LM-WQML-2003640-Cryptogramma LGSGTRKDPGPW.............ERELRNMWKPTKQEGL...........W..FHG.GNLHQS.....RHYSLMLA
|LM-WQML-2004411-Cryptogramma ..............................................................................
|LM-WQML-2067992-Cryptogramma LGSDTAKDPGPW.............EREIRNMWKPTHVPNL...........W..FHG.GGFQEA.....RHYSLYLA
2095928-Ophioglossum_vulgatum LGSATTKDPGPW.............ERELRNMWKPTQQPSL...........W..FHG.GNLHQS.....RHYSLYLA
2024024-Polysiphonia_japonica LGSNTPKDPGPW.............EGELRNMYKPTRQEGM...........W..YFG.GNLHQC.....RHYSLCLA
-2100101-Pallavicinia_lyellii ..............................................................................
JJY-2002336-Woodsia_scopulina LGSDTAKDPGPW.............EGELRNMWKPTQQPQL...........W..FQG.GNLQQS.....RYFSLLLA
-2008616-Woodsia_scopulina.p2 ..............................................................................
JJY-2076923-Woodsia_scopulina LGSNTKGDPGPW.............ERELRNMWKPTQQEAL...........W..FHA.GGFIDA.....RFHSRFLA
JJY-2077796-Woodsia_scopulina LGSGTRKDPGPW.............ERELRNMWKPTQQDGL...........W..FHG.GNLHQS.....RHYSLSLA
A-2072483-Polypodium_amorphum LGSDTTKDPGPW.............EREIRNMWKPTHQQGL...........W..FHG.GSFQET.....RHVSLYLA
A-2073185-Polypodium_amorphum LGSGTRKDPGPW.............EIELRNMWKPTQQEGL...........W..FHG.GNLQQS.....RHYSLLLA
A-2073213-Polypodium_amorphum LGSGTRKDPGPW.............ERELRNMWKPTRQEGL...........W..FHG.GNLHQS.....RHYSLLLA
-2003047-Cystopteris_protrusa LGSNTTKDPGPW.............EGEIRNMWKPTRQQSL...........W..FHG.GGFEET.....RHFSLYLA
-2007507-Cystopteris_protrusa LGSDTAKDPGPW.............EGELRNMWKPTQQPHL...........W..FHG.GKLQQS.....RHFSLLLG
-2012306-Cystopteris_protrusa LGSGTRKDPGPW.............ERELRNMWKPTQQEGL...........W..FHG.GNLHQS.....RHYSLLLA
038519-Heterosiphonia_pulchra LGSGTRDDPGPW.............QGELRNMWKPLRQKGL...........W..MFG.GNLHQC.....RHYSLVVA
042241-Selaginella_kraussiana LGSETPKDPGPW.............EGELRNMWKPTQQEAL...........W..FHG.GNLHQA.....RHYSMYLA
A|LP-ZZOL-2000160-Selaginella LGSNTRKDPGPW.............EGELRNMWKPTAQEGL...........W..FHG.GNLHQA.....RHYSLYLA
```

```
                                            590          600
                                              .            .
SisterYUCCA|Mapoly0052s0062       L......QLK..ARLEGIPTPVYALPKV.NHA...............................
SisterYUCCA|Mapoly0122s0026       L......QLK..ARMEGIPTPVYKLVRH.KEPDSPVLASKM...............SSIQMQD
SisterYUCCA|Sm113792              L......QLK..ARMEGIATPVYGMPKV.HHLS..............................
SisterYUCCA|Sm422043              L......QLK..ARMEGIATPVYGVPKV.HHLS..............................
SisterYUCCA|kf100109_0340         L......QLK..ARFEGLPTPVYR.......................................
4265-Phaeomegaceros_coriaceus     I......QIK..ARQLGISMQVYGMSKV.HHLR..............................
W-AKXB-2007730-Phaeomegaceros     L......QIK..ARLEGIPTPVYGMPPV.HHLT..............................
W-AKXB-2007731-Phaeomegaceros     ...............................................................
W-AKXB-2067423-Phaeomegaceros     L......QIK..ARLEGIATNAYGLQPV.HHLS..............................
BMJR-2002471-Adiantum_tenerum     L......QLK..ARMEGIPVHVYGLPEV.HHSK..............................
BMJR-2059933-Adiantum_tenerum     L......QLK..AR...IPTPVFRFPPI.GSSSL.............................
BMJR-2062494-Adiantum_tenerum     L......QLK..ARMEGLPTPVYGLAKV.HHTK..............................
U-2019103-Radula_lindenbergia     L......QIK..GLLEGLYKY..........................................
U-2086103-Radula_lindenbergia     L......QIK..AIEEGLASG..........................................
CA|HW-DXOU-2035997-Nothoceros     V......QIK..VRQLGISIQVYGMGKV.HHLC..............................
FAJB-2001596-Paraphymatoceros     L......QIK..ARLEGIPTPVYGLPQV.HHLS..............................
FAJB-2004773-Paraphymatoceros     L......QMK..ARYEGISTPVYGMQPV.FHLS..............................
FAJB-2008542-Paraphymatoceros     L......QIK..ARLEGIPTPLYGLQQV.HHLA..............................
CA|GA-FPCO-2029146-Interfilum     L......QLK..ARFEGLPTPVFG.......................................
GA-FQLP-2033521-Klebsormidium     L......QLK..ARFEGLLTPVYS.......................................
YUCCA|LM-GANB-2011315-Cyathea     ...........................V.HHSC..............................
YUCCA|LM-GANB-2011316-Cyathea     L......QLK..ARMEGIPTPVYGLPKV.HHSN..............................
YUCCA|LM-GANB-2017253-Cyathea     L......QLK..ARMEGIPTPVYGMPEV.HHIK..............................
ANB-2070212-Cyathea_spinulosa     L......QIK..ARMEGIPTPVYGLPEV.HHTM..............................
LW-HERT-2045223-Sphaerocarpos     L......QLK..ARMEAIPTPTYGVPQV.HHVA..............................
UCCA|RA-IEHF-2003087-Dumontia     L......QLK..ARHVGIETPVYEVQEA.WHQC..............................
O-2067228-Marchantia_paleacea     L......QLK..ARMEGIPTPIYALPQV.NHVRV.............................
O-2067575-Marchantia_paleacea     I......QLK..ARMEGIPTPVYTRSK....................................
-2046311-Conocephalum_conicum     L......QLK..ARMEGIPTPVFGKSLP.KSVK..............................
-2046658-Conocephalum_conicum     L......QLK..ARMEGIPTPTYGVPPV.NHVKL.............................
-2181954-Selaginella_wallacei     L......QLK..ARMEGIPTPVYGIPEV.HHLA..............................
A|LP-KUXM-2009452-Selaginella     ...............................................................
|LM-PIVW-2024618-Ceratopteris     L......QLK..ARMEGIPTPVFGLAEV.HHTR..............................
2005240-Isoetes_tegetiformans     L......QLK..ARMEGIPTPVYGLPSTTYHLS..............................
2005241-Isoetes_tegetiformans     L......QLK..ARMEGIPTPVYGLPSTTYHLS..............................
2008401-Hymenophyllum_bivalve     L......QLK..ARMEGIPTPVYGQPQV.HHTS..............................
003161-Megaceros_vincentianus     .................................P.............................
007840-Megaceros_vincentianus     ...............................................................
013674-Megaceros_vincentianus     L......QIK..ARLEGIPTPVYGIPPV.HHVT..............................
016968-Megaceros_vincentianus     ...............................................................
019507-Megaceros_vincentianus     L......QIK..ARFEGIAANAYGMQPV.HHLS..............................
2000839-Marchantia_emarginata     ...............................................................
2004520-Marchantia_emarginata     L......QLK..ARMEGIPTPVWTLPK....................................
2069965-Marchantia_emarginata     ...............................................................
2074864-Marchantia_emarginata     L......QLK..ARMEGIATPAYGVPVV.NHPVA.............................
2050988-Anthoceros_agrestis-A     L......QIT..ARYQGISTPVYGMQPV.FHLS..............................
VB-2006165-Lunularia_cruciata     R......GVQ..ERVETYSTS..........................................
VB-2006166-Lunularia_cruciata     I......QLK..ARFEGIQTPVYNDPAS.HRSS..............................
VB-2006167-Lunularia_cruciata     I......QLK..ARFEGIQTPVYNDPAS.HRSS..............................
VB-2006168-Lunularia_cruciata     I......QLK..ARMEGIQVSVYNDPAS.PQST..............................
VB-2007869-Lunularia_cruciata     L......QLK..ARLEGIPTPVYGLPPV.HHVL..............................
VB-2075591-Lunularia_cruciata     V......QLK..ARMEGIPTPVYTPHPS.IALK..............................
CRN-2050303-Megaceros_tosanus     LAACTTSQVN..CRLTS.....YRM......................................
CRN-2052782-Megaceros_tosanus     L......QIK..ARLEGIPTPVYGMPPV.RHLV..............................
VITX-2002371-Blechnum_spicant     L......QLK..ARMEGIPTPVYGLPKV.QHTQ..............................
VITX-2009142-Blechnum_spicant     L......QLK..ARMEGIPTPVYGLPKV.HHTK..............................
VITX-2009143-Blechnum_spicant     L......QLK.....................................................
VITX-2016791-Blechnum_spicant     L......QLK..ARLEHIPTPVFRFSTL.ANTGPSGKVPT...GNGVHA....PNGNGMLMM.
VITX-2098911-Blechnum_spicant     L......QIK..ARMESIHTPVYGLPQV.HHSS..............................
VITX-2099259-Blechnum_spicant     L......QLK..AGMEAIPTPVYGLPHV.HHTS..............................
CA|HW-WCZB-2015184-Phaeoceros     ...............................................................
CA|HW-WCZB-2016004-Phaeoceros     L......QLK..ARFEGISTPVYGMQPV.FHLS..............................
CA|HW-WCZB-2117013-Phaeoceros     ...............................................................
CA|HW-WCZB-2118293-Phaeoceros     L......QIK..ARLEGIPTPVYGLPQV.HHLS..............................
U-2004486-Leucostegia_immersa     L......QLK..ARMERITTPIFRFPPL.TSNSELDITPSIWPSNSKRAWSAFPPPSFASLP.
U-2004559-Leucostegia_immersa     L......QLK..ARMEGIPTPVYGLPKV.HHIN..............................
JLO-2000942-Riccia_berychiana     P.....SQVSQYARLEN...........YIVI...............................
JLO-2002497-Riccia_berychiana     L......QLK..ARMEGIPTPAYGIPSV.NHVS..............................
JLO-2033265-Riccia_berychiana     ...............................................................
 |LM-WQML-2003640-Cryptogramma    L......QLK..ARMEGIPTPVYGIPEV.HHIQ..............................
 |LM-WQML-2004411-Cryptogramma    ...............................................................
 |LM-WQML-2067992-Cryptogramma    L......QIK..ARLEAIPTPVYGLPLV.HHLA..............................
2095928-Ophioglossum_vulgatum     L......QLK..ARMEGIPTPVYALAQV.HHTS..............................
2024024-Polysiphonia_japonica     L......QLK..ARHVGIPTPVYNLATV.YHMW..............................
-2100101-Pallavicinia_lyellii     ...............................................................
JJY-2002336-Woodsia_scopulina     L......QLK..ARMEG..............................................
-2008616-Woodsia_scopulina.p2     ...............................................................
JJY-2076923-Woodsia_scopulina     L......QLK..ARMEAIPTPVYGLPRV.HHTS..............................
JJY-2077796-Woodsia_scopulina     L......QLK..ARMEGIPTPVYGLPKV.HHTN..............................
A-2072483-Polypodium_amorphum     L......Q.......................................................
A-2073185-Polypodium_amorphum     L......QLK..ARMEGIPTPVYGLSKV.YHTK..............................
A-2073213-Polypodium_amorphum     L......QLK..ARMEGIPTPVYGMPEV.HHIK..............................
-2003047-Cystopteris_protrusa     L......QIK..ARLEGIPTPVYGLPQV.HHSS..............................
-2007507-Cystopteris_protrusa     L......QLK..ARMEGIPTPVYGLPKV.HHTQ..............................
-2012306-Cystopteris_protrusa     L......QIK..ARMEGIPTPVYGMPEV.HHIK..............................
038519-Heterosiphonia_pulchra     L......QLK..ARLEGLPISVYGIPVA.TPLD..............................
042241-Selaginella_kraussiana     L......QLK..ARMEGISTPVYGIPKV.HHLA..............................
A|LP-ZZOL-2000160-Selaginella     L......QLK..ARMEGITTPVYGIPKV.RHLS..............................
```
