## Supplementary Data (Data S1-S10) for "The auxin gatekeepers: Evolution and diversification of the YUCCA family": Data S5.pdf

### Supplementary Data S5

**MEME (Multiple Em for Motif Elicitation) analysis for YUCs with other class B FMOs (S-OX, N-OX and sYUCs)**

(https://meme-suite.org/; accessed on 08/07/2024)

(Reference: Timothy L. Bailey and Charles Elkan, "Fitting a mixture model by expectation maximization to discover motifs in biopolymers", Proceedings of the Second International Conference on Intelligent Systems for Molecular Biology, pp. 28-36, AAAI Press, Menlo Park, California, 1994)

#### MEME Running parameters used: -

| | |
|---|---|
| Selection for site distribution | ZOOPS: Zero or one site per sequence |
| Motif Count | Searched for 30 motifs |
| Motif Width | Between 5 wide and 50 wide (inclusive). |
| Motif E-value Threshold | No limit |
| MEME Version used | MEME version 5.5.5 |

#### Identified Motifs

| | Logo | E-value | Sites | Width |
|---|---|---|---|---|
| 1. | 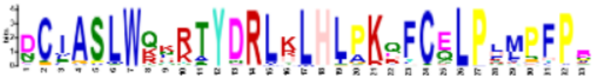 | 4.1e-5870 | 442 | 33 |
| 2. | 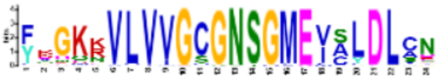 | 2.5e-4355 | 473 | 24 |
| 3. | 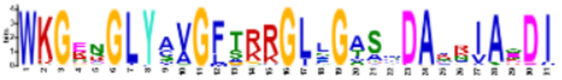 | 1.4e-4955 | 461 | 31 |
| 4. | 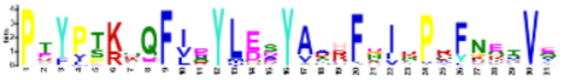 | 7.1e-4430 | 460 | 31 |
| 5. | 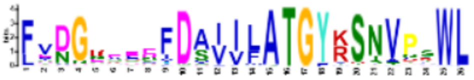 | 1.1e-3873 | 448 | 26 |
| 6. | 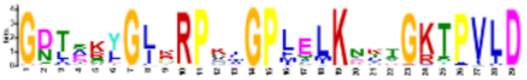 | 2.0e-4245 | 453 | 29 |
| 7. | 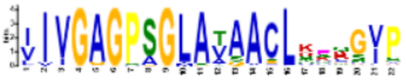 | 3.9e-3455 | 456 | 22 |
| 8. | 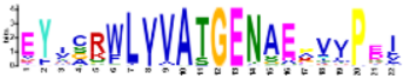 | 7.5e-3350 | 460 | 22 |
| 9. | 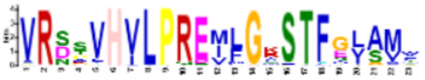 | 7.0e-2785 | 350 | 23 |
| 10. | 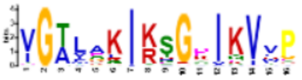 | 1.1e-1910 | 463 | 16 |
| 11. | 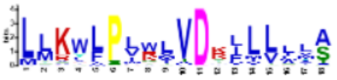 | 1.6e-1697 | 452 | 18 |
| 12. | 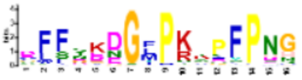 | 2.7e-1457 | 405 | 16 |
| 13. | 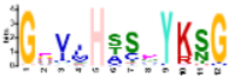 | 2.9e-1347 | 467 | 12 |
| 14. | 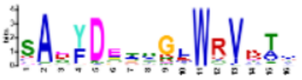 | 2.6e-1294 | 427 | 16 |

| | | E-value | Sites | Width |
|---|---|---|---|---|
| 15. | 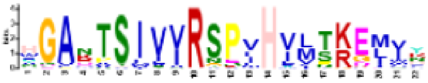 | 7.3e-554 | 107 | 22 |
| 16. | 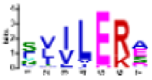 | 3.9e-444 | 444 | 7 |
| 17. | 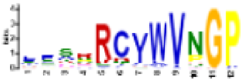 | 3.0e-431 | 191 | 12 |
| 18. | 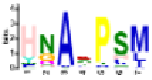 | 1.8e-354 | 298 | 7 |
| 19. | 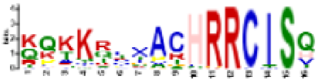 | 2.6e-293 | 78 | 16 |
| 20. | 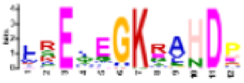 | 9.5e-199 | 91 | 12 |
| 21. | 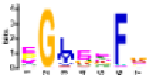 | 4.7e-147 | 446 | 7 |
| 22. | 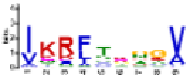 | 2.2e-109 | 321 | 9 |
| 23. | 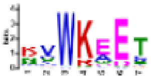 | 3.7e-083 | 133 | 7 |
| 24. | 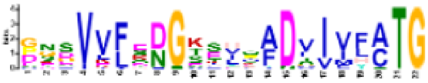 | 1.5e-060 | 29 | 22 |
| 25. | 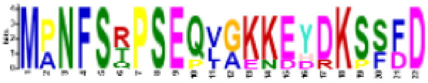 | 5.6e-031 | 6 | 22 |
| 26. | 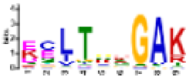 | 9.9e-029 | 67 | 9 |
| 27. | 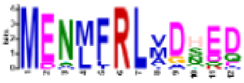 | 5.7e-026 | 19 | 12 |
| 28. | 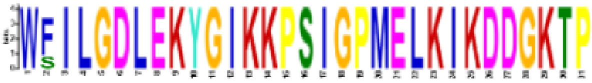 | 3.9e-025 | 4 | 31 |
| 29. | 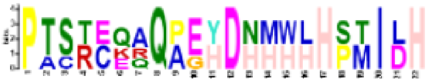 | 6.0e-023 | 4 | 22 |
| 30. | 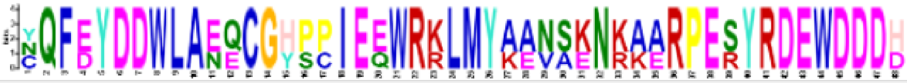 | 8.9e-022 | 3 | 48 |

**More motif details are given below**

| Name | *p*-value | Motif Locations |
|---|---|---|
| AsFMO_GS-OX_BAS32646_Allium_sativum | 4.28e-72 | |
| AtFMO_GS-OX_NP001321123.1_A.thaliana | 9.31e-46 | |
| OsFMO_GS-OX_XP015612837_O.sativa | 8.10e-94 | |
| HvFMO_GS-OX_XP044971712.1_H.vulgare | 2.31e-98 | |
| SmFMO_GS-OX_XP_002973683.1_S.moellendorffii | 2.92e-19 | |
| MpFMO_GS-OX_PTQ49826.1__Marchantia | 1.50e-34 | |
| PcFMO_GS-OX_scaffold2006551_P.carolinianus | 5.04e-20 | |
| AtFMO1_N-OX_AT1G19250.1_A.thaliana | 2.19e-29 | |
| OsFMO1_N-OX_LOCOs03g08410.1_O.sativa | 1.41e-32 | |
| HvFMO1_N-OX_HORVU4Hr1G077170_H.vulgare | 6.13e-25 | |
| SmFMO1_N-OX_XP002967925.1_S.moellendorffii | 2.91e-19 | |
| PcFMO1_N-OX_scaffold2088977_P.carolinianus | 1.12e-14 | |
| SisterYUCCA\|HW-WCZB-2015184-P.carolinianus | 5.57e-40 | |
| SisterYUCCA\|HW-WCZB-2016004-P.carolinianus | 3.11e-49 | |
| SisterYUCCA\|HW-WCZB-2118293-P.carolinianus | 7.46e-44 | |
| SisterYUCCA\|Mapoly0052s0062_Marchantia | 4.14e-45 | |
| SisterYUCCA\|Mapoly0122s0026_Marchantia | 2.21e-43 | |
| SisterYUCCA\|Sm113792_S.moellendorffii | 5.55e-48 | |
| SisterYUCCA\|Sm422043_S.moellendorffii | 3.10e-48 | |
| SisterYUCCA\|EFJ16321.1_S.moellendorffii | 5.55e-47 | |
| ED_a_CpYUCE_CepurGG1.9G038900.1.p_C.purpureus | 1.18e-232 | |
| ED_a_PpYUCB_Pp3c11_11790V3.1.p_P.patens | 7.20e-231 | |
| ED_b_CpYUCA_CepurGG1.4G246000.1.p_C.purpureus | 1.98e-222 | |
| ED_b_PpYUCC_Pp3c1_11500V3.1.p_P.patens | 8.92e-221 | |
| ED_b_PpYUCD_Pp3c2_27740V3.1.p_P.patens | 4.90e-213 | |
| ED_b_PpYUCE_Pp3c13_21970V3.1.p_P.patens | 1.44e-215 | |
| ED_b_PpYUCF_Pp3c3_20490V3.1.p_P.patens | 1.77e-216 | |
| ED_c_MpYUC1_Mapoly0127s0005.1.p_M.polymorpha | 1.13e-219 | |
| ED_c_MpYUC2_Mapoly0063s0040.1.p_M.polymorpha | 5.47e-248 | |
| ED_d_AaYUC_scaffold_2048594_Anthoceros_agrestis | 1.03e-150 | |
| ED_d_PcYUCA_scaffold_2017088_P.carolinianus | 9.40e-246 | |
| ED_d_PcYUCB_scaffold_2088798_P.carolinianus | 4.30e-247 | |
| ED_d_PcYUCC_scaffold_2018152_P.carolinianus | 3.30e-175 | |

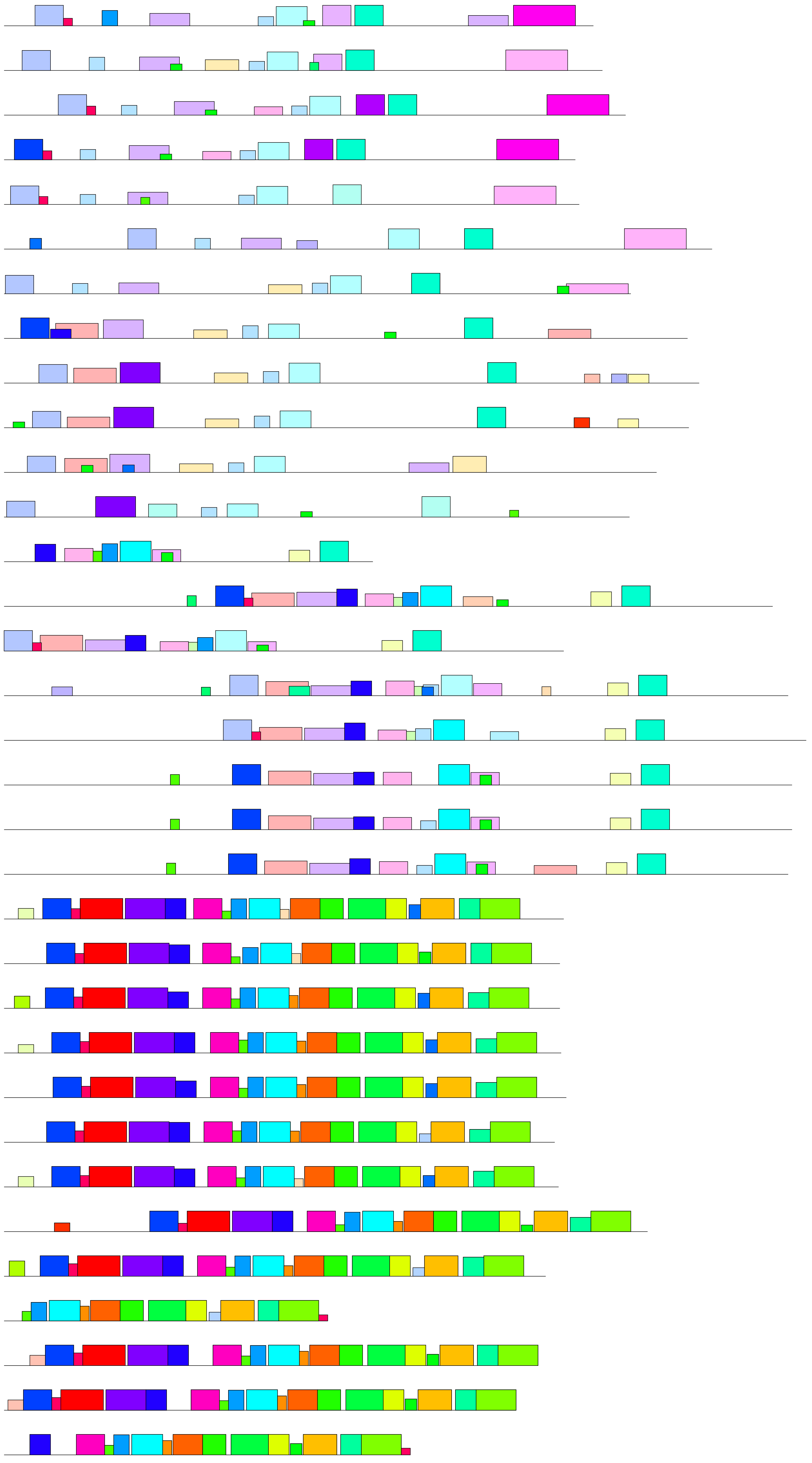

| Motif | Symbol | Motif Consensus |
|---|---|---|
| 1. | | DCIASLWQKRTYDRLKLHLPKQFCZLPLMPFPE |
| 2. | | FRGKKVLVVGCGNSGMEVSLDLCN |
| 3. | | WKGENGLYAVGFTRRGLLGASMDAVRIAZDI |
| 4. | | PTYPTKQQFIDYLESYAAHFDIRPRFNETVZ |
| 5. | | FVDGREEEFDAIILATGYKSNVPSWL |
| 6. | | GBTSKYGJKRPKIGPLELKNKTGKTPVLD |
| 7. | | IIVGAGPSGLATAACLKEQGVP |
| 8. | | EYICRWLVVATGENAEPVVPEI |
| 9. | | VRSSVHVLPREMLGKSTFGLAMW |
| 10. | | VGTLAKIKSGDIKVVP |
| 11. | | LLKWLPLWLVDKLLLLLA |
| 12. | | DFFSKDGFPKTPFPNG |
| 13. | | GEVIHSSDYKSG |
| 14. | | SAEYDETSGLWRVKTV |
| 15. | | HGANTSIVVRSPVHVLTKEMVY |
| 16. | | SVILERA |
| 17. | | LFSRRCVWVNGP |
| 18. | | HNAKPSM |
| 19. | | KQKKRFVACHRRCISQ |
| 20. | | LRELEGKRAHDP |
| 21. | | EGLEEFG |
| 22. | | IKRFTRGGV |
| 23. | | KVWKEET |
| 24. | | GNSVVFEBGKSHQADVIVFATG |
| 25. | | MPNFSRPSEQVGKKEYDKSSFD |
| 26. | | ECLTPHGAK |
| 27. | | MENMFRLVDHED |
| 29. | | PTSRCQAQPEHDNHWHHPTIDH |
| 30. | | NQFEYDDWLAEQCGHPPIEEWRKLMYAANSKNKAARPESYRDEWDDDH |

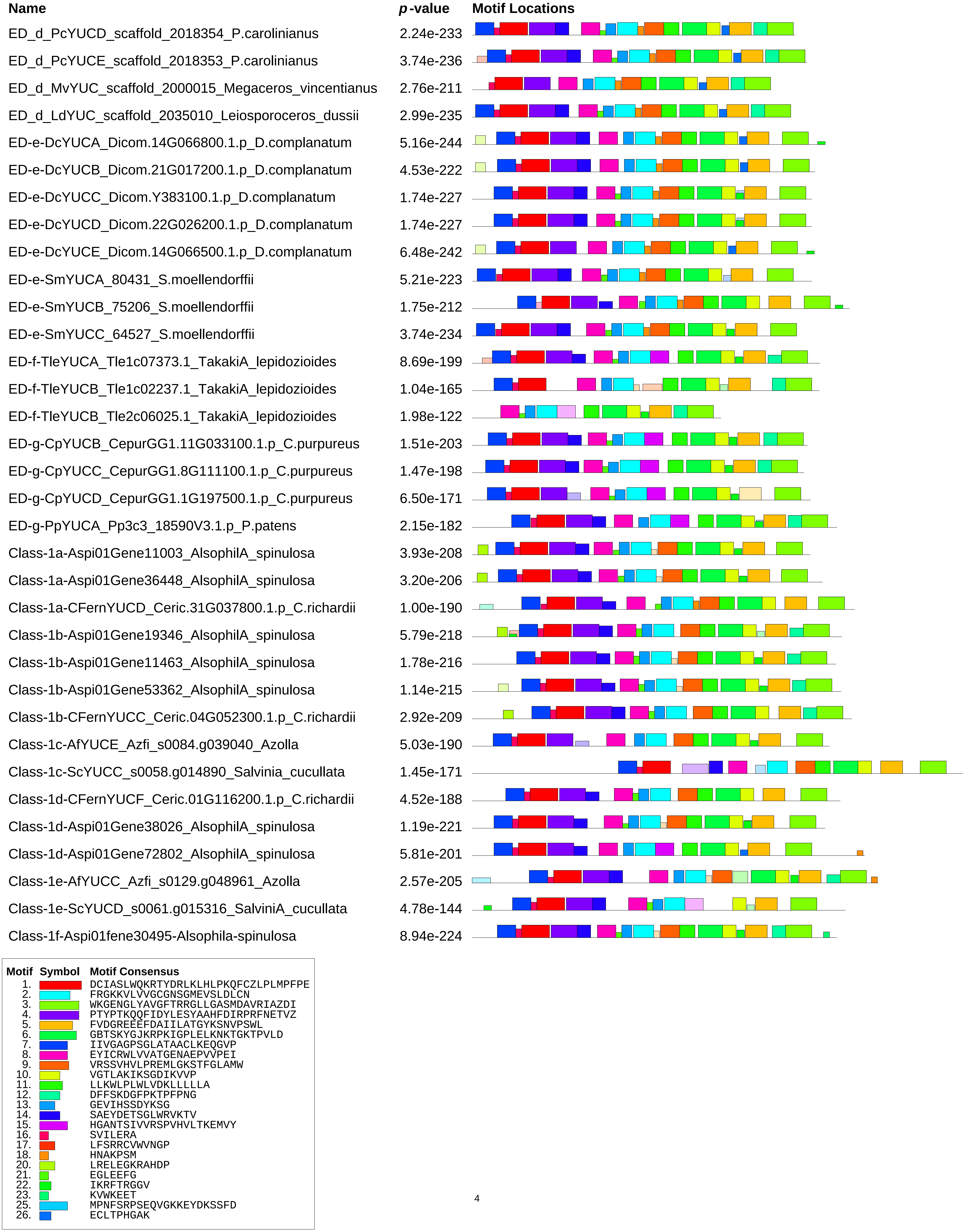
Name
p-value
Motif Locations
ED_d_PcYUCD_scaffold_2018354_P.carolinianus 2.24e-233
ED_d_PcYUCE_scaffold_2018353_P.carolinianus 3.74e-236
ED_d_MvYUC_scaffold_2000015_Megaceros_vincentianus 2.76e-211
ED_d_LdYUC_scaffold_2035010_Leiosporoceros_dussii 2.99e-235
ED-e-DcYUCA_Dicom.14G066800.1.p_D.complanatum 5.16e-244
ED-e-DcYUCB_Dicom.21G017200.1.p_D.complanatum 4.53e-222
ED-e-DcYUCC_Dicom.Y383100.1.p_D.complanatum 1.74e-227
ED-e-DcYUCD_Dicom.22G026200.1.p_D.complanatum 1.74e-227
ED-e-DcYUCE_Dicom.14G066500.1.p_D.complanatum 6.48e-242
ED-e-SmYUCA_80431_S.moellendorffii 5.21e-223
ED-e-SmYUCB_75206_S.moellendorffii 1.75e-212
ED-e-SmYUCC_64527_S.moellendorffii 3.74e-234
ED-f-TleYUCA_Tle1c07373.1_TakakiA_lepidozioides 8.69e-199
ED-f-TleYUCB_Tle1c02237.1_TakakiA_lepidozioides 1.04e-165
ED-f-TleYUCB_Tle2c06025.1_TakakiA_lepidozioides 1.98e-122
ED-g-CpYUCB_CepurGG1.11G033100.1.p_C.purpureus 1.51e-203
ED-g-CpYUCC_CepurGG1.8G111100.1.p_C.purpureus 1.47e-198
ED-g-CpYUCD_CepurGG1.1G197500.1.p_C.purpureus 6.50e-171
ED-g-PpYUCA_Pp3c3_18590V3.1.p_P.patens 2.15e-182
Class-1a-Aspi01Gene11003_AlsophilA_spinulosa 3.93e-208
Class-1a-Aspi01Gene36448_AlsophilA_spinulosa 3.20e-206
Class-1a-CFernYUCD_Ceric.31G037800.1.p_C.richardii 1.00e-190
Class-1b-Aspi01Gene19346_AlsophilA_spinulosa 5.79e-218
Class-1b-Aspi01Gene11463_AlsophilA_spinulosa 1.78e-216
Class-1b-Aspi01Gene53362_AlsophilA_spinulosa 1.14e-215
Class-1b-CFernYUCC_Ceric.04G052300.1.p_C.richardii 2.92e-209
Class-1c-AfYUCE_Azfi_s0084.g039040_Azolla 5.03e-190
Class-1c-ScYUCC_s0058.g014890_Salvinia_cucullata 1.45e-171
Class-1d-CFernYUCF_Ceric.01G116200.1.p_C.richardii 4.52e-188
Class-1d-Aspi01Gene38026_AlsophilA_spinulosa 1.19e-221
Class-1d-Aspi01Gene72802_AlsophilA_spinulosa 5.81e-201
Class-1e-AfYUCC_Azfi_s0129.g048961_Azolla 2.57e-205
Class-1e-ScYUCD_s0061.g015316_SalviniA_cucullata 4.78e-144
Class-1f-Aspi01fene30495-Alsophila-spinulosa 8.94e-224
Motif Symbol Motif Consensus
1. DCIASLWQKRTYDRLKLHLPKQFCZLPLMPFPE
2. FRGKKVLVVGCGNSGMEVSLDLCN
3. WKGENGLYAVGFTRRGLLGASMDAVRIAZDI
4. PTYPTKQQFIDYLESYAAHFDIRPRFNETVZ
5. FVDGREEEFDAIILATGYKSNVPSWL
6. GBTSKYGJKRPKIGPLELKNKTGKTPVLD
7. IIVGAGPSGLATAACLKEQGVP
8. EYICRWLVVATGENAEPVVPEI
9. VRSSVHVLPREMLGKSTFGLAMW
10. VGTLAKIKSGDIKVVP
11. LLKWLPLWLVDKLLLLLA
12. DFFSKDGFPKTPFPNG
13. GEVIHSSDYKSG
14. SAEYDETSGLWRVKTV
15. HGANTSIVVRSPVHVLTKEMVY
16. SVILERA
17. LFSRRCVWVNGP
18. HNAKPSM
20. LRELEGKRAHDP
21. EGLEEFG
22. IKRFTRGGV
23. KVWKEET
25. MPNFSRPSEQVGKKEYDKSSFD
26. ECLTPHGAK

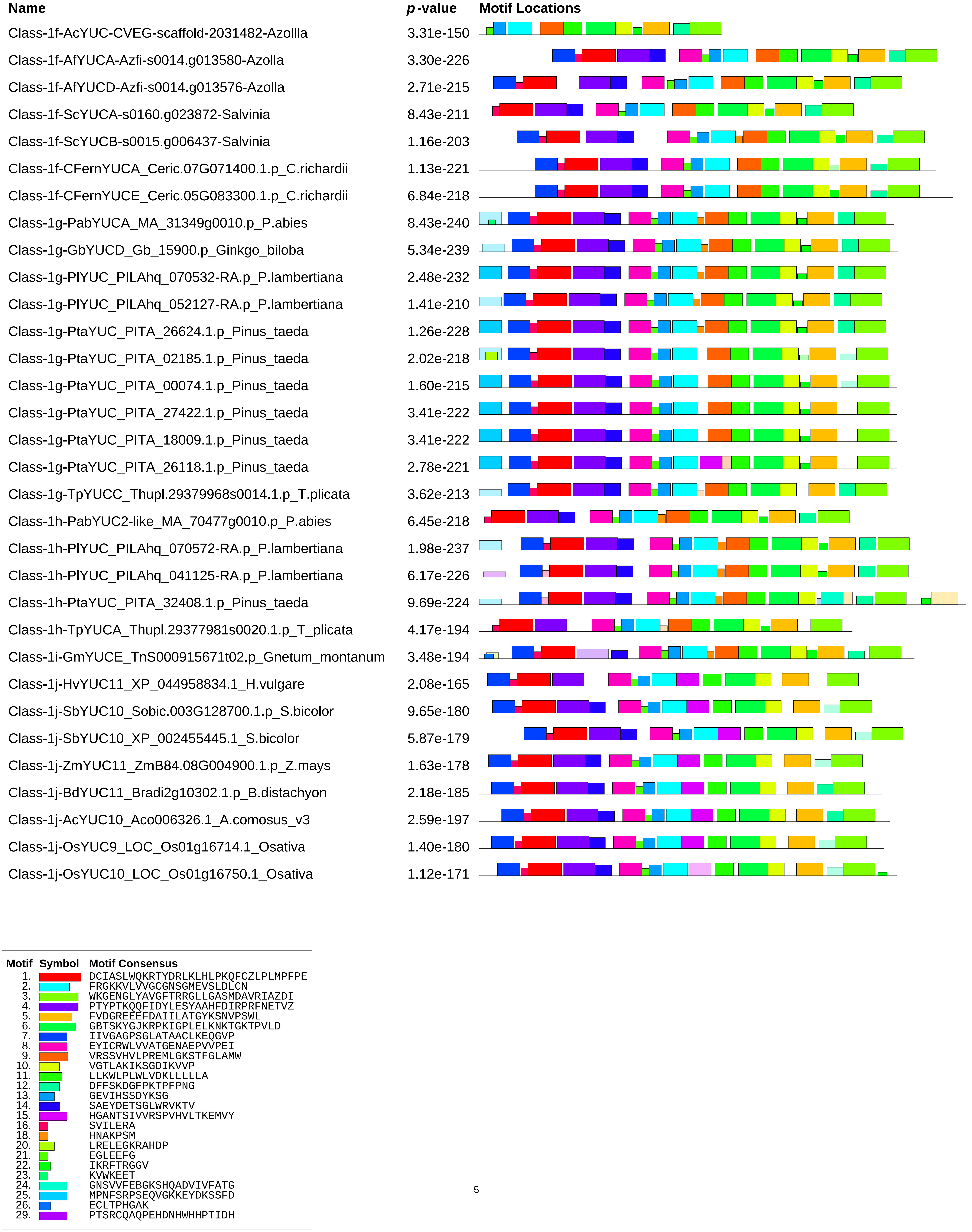
Name
p-value
Motif Locations
Class-1f-AcYUC-CVEG-scaffold-2031482-Azollla 3.31e-150
Class-1f-AfYUCA-Azfi-s0014.g013580-Azolla 3.30e-226
Class-1f-AfYUCD-Azfi-s0014.g013576-Azolla 2.71e-215
Class-1f-ScYUCA-s0160.g023872-Salvinia 8.43e-211
Class-1f-ScYUCB-s0015.g006437-Salvinia 1.16e-203
Class-1f-CFernYUCA_Ceric.07G071400.1.p_C.richardii 1.13e-221
Class-1f-CFernYUCE_Ceric.05G083300.1.p_C.richardii 6.84e-218
Class-1g-PabYUCA_MA_31349g0010.p_P.abies 8.43e-240
Class-1g-GbYUCD_Gb_15900.p_Ginkgo_biloba 5.34e-239
Class-1g-PlYUC_PILAhq_070532-RA.p_P.lambertiana 2.48e-232
Class-1g-PlYUC_PILAhq_052127-RA.p_P.lambertiana 1.41e-210
Class-1g-PtaYUC_PITA_26624.1.p_Pinus_taeda 1.26e-228
Class-1g-PtaYUC_PITA_02185.1.p_Pinus_taeda 2.02e-218
Class-1g-PtaYUC_PITA_00074.1.p_Pinus_taeda 1.60e-215
Class-1g-PtaYUC_PITA_27422.1.p_Pinus_taeda 3.41e-222
Class-1g-PtaYUC_PITA_18009.1.p_Pinus_taeda 3.41e-222
Class-1g-PtaYUC_PITA_26118.1.p_Pinus_taeda 2.78e-221
Class-1g-TpYUCC_Thupl.29379968s0014.1.p_T.plicata 3.62e-213
Class-1h-PabYUC2-like_MA_70477g0010.p_P.abies 6.45e-218
Class-1h-PlYUC_PILAhq_070572-RA.p_P.lambertiana 1.98e-237
Class-1h-PlYUC_PILAhq_041125-RA.p_P.lambertiana 6.17e-226
Class-1h-PtaYUC_PITA_32408.1.p_Pinus_taeda 9.69e-224
Class-1h-TpYUCA_Thupl.29377981s0020.1.p_T_plicata 4.17e-194
Class-1i-GmYUCE_TnS000915671t02.p_Gnetum_montanum 3.48e-194
Class-1j-HvYUC11_XP_044958834.1_H.vulgare 2.08e-165
Class-1j-SbYUC10_Sobic.003G128700.1.p_S.bicolor 9.65e-180
Class-1j-SbYUC10_XP_002455445.1_S.bicolor 5.87e-179
Class-1j-ZmYUC11_ZmB84.08G004900.1.p_Z.mays 1.63e-178
Class-1j-BdYUC11_Bradi2g10302.1.p_B.distachyon 2.18e-185
Class-1j-AcYUC10_Aco006326.1_A.comosus_v3 2.59e-197
Class-1j-OsYUC9_LOC_Os01g16714.1_Osativa 1.40e-180
Class-1j-OsYUC10_LOC_Os01g16750.1_Osativa 1.12e-171
Motif Symbol Motif Consensus
1. DCIASLWQKRTYDRLKLHLPKQFCZLPLMPFPE
2. FRGKKVLVVGCGNSGMEVSLDLCN
3. WKGENGLYAVGFTRRGLLGASMDAVRIAZDI
4. PTYPTKQQFIDYLESYAAHFDIRPRFNETVZ
5. FVDGREEEFDAIILATGYKSNVPSWL
6. GBTSKYGJKRPKIGPLELKNKTGKTPVLD
7. IIVGAGPSGLATAACLKEQGVP
8. EYICRWLVVATGENAEPVVPEI
9. VRSSVHVLPREMLGKSTFGLAMW
10. VGTLAKIKSGDIKVVP
11. LLKWLPLWLVDKLLLLLA
12. DFFSKDGFPKTPFPNG
13. GEVIHSSDYKSG
14. SAEYDETSGLWRVKTV
15. HGANTSIVVRSPVHVLTKEMVY
16. SVILERA
18. HNAKPSM
20. LRELEGKRAHDP
21. EGLEEFG
22. IKRFTRGGV
23. KVWKEET
24. GNSVVFEBGKSHQADVIVFATG
25. MPNFSRPSEQVGKKEYDKSSFD
26. ECLTPHGAK
29. PTSRCQAQPEHDNHWHHPTIDH

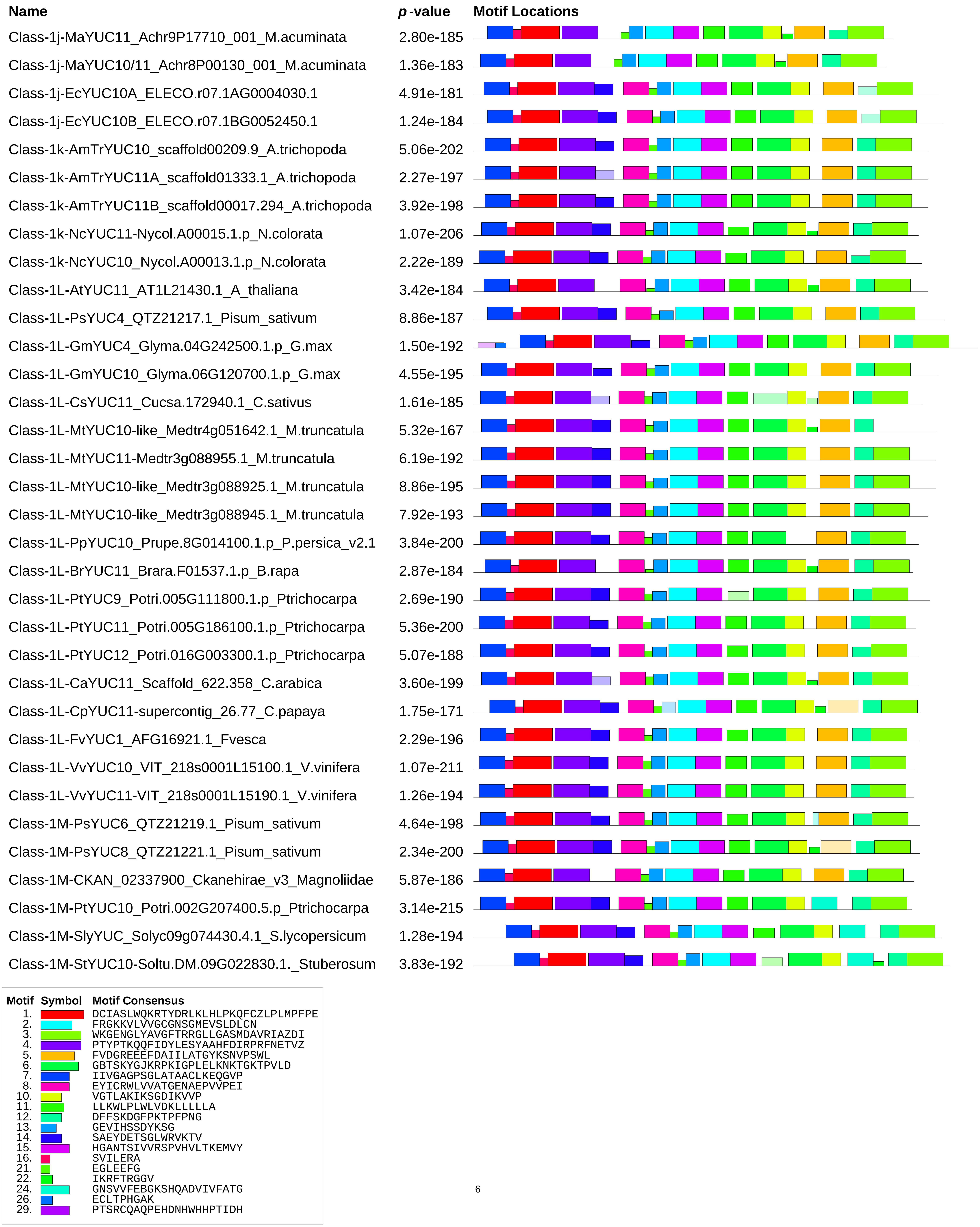
Name
p-value
Motif Locations
Class-1j-MaYUC11_Achr9P17710_001_M.acuminata 2.80e-185
Class-1j-MaYUC10/11_Achr8P00130_001_M.acuminata 1.36e-183
Class-1j-EcYUC10A_ELECO.r07.1AG0004030.1 4.91e-181
Class-1j-EcYUC10B_ELECO.r07.1BG0052450.1 1.24e-184
Class-1k-AmTrYUC10_scaffold00209.9_A.trichopoda 5.06e-202
Class-1k-AmTrYUC11A_scaffold01333.1_A.trichopoda 2.27e-197
Class-1k-AmTrYUC11B_scaffold00017.294_A.trichopoda 3.92e-198
Class-1k-NcYUC11-Nycol.A00015.1.p_N.colorata 1.07e-206
Class-1k-NcYUC10_Nycol.A00013.1.p_N.colorata 2.22e-189
Class-1L-AtYUC11_AT1L21430.1_A_thaliana 3.42e-184
Class-1L-PsYUC4_QTZ21217.1_Pisum_sativum 8.86e-187
Class-1L-GmYUC4_Glyma.04G242500.1.p_G.max 1.50e-192
Class-1L-GmYUC10_Glyma.06G120700.1.p_G.max 4.55e-195
Class-1L-CsYUC11_Cucsa.172940.1_C.sativus 1.61e-185
Class-1L-MtYUC10-like_Medtr4g051642.1_M.truncatula 5.32e-167
Class-1L-MtYUC11-Medtr3g088955.1_M.truncatula 6.19e-192
Class-1L-MtYUC10-like_Medtr3g088925.1_M.truncatula 8.86e-195
Class-1L-MtYUC10-like_Medtr3g088945.1_M.truncatula 7.92e-193
Class-1L-PpYUC10_Prupe.8G014100.1.p_P.persica_v2.1 3.84e-200
Class-1L-BrYUC11_Brara.F01537.1.p_B.rapa 2.87e-184
Class-1L-PtYUC9_Potri.005G111800.1.p_Ptrichocarpa 2.69e-190
Class-1L-PtYUC11_Potri.005G186100.1.p_Ptrichocarpa 5.36e-200
Class-1L-PtYUC12_Potri.016G003300.1.p_Ptrichocarpa 5.07e-188
Class-1L-CaYUC11_Scaffold_622.358_C.arabica 3.60e-199
Class-1L-CpYUC11-supercontig_26.77_C.papaya 1.75e-171
Class-1L-FvYUC1_AFG16921.1_Fvesca 2.29e-196
Class-1L-VvYUC10_VIT_218s0001L15100.1_V.vinifera 1.07e-211
Class-1L-VvYUC11-VIT_218s0001L15190.1_V.vinifera 1.26e-194
Class-1M-PsYUC6_QTZ21219.1_Pisum_sativum 4.64e-198
Class-1M-PsYUC8_QTZ21221.1_Pisum_sativum 2.34e-200
Class-1M-CKAN_02337900_Ckanehirae_v3_Magnoliidae 5.87e-186
Class-1M-PtYUC10_Potri.002G207400.5.p_Ptrichocarpa 3.14e-215
Class-1M-SlyYUC_Solyc09g074430.4.1_S.lycopersicum 1.28e-194
Class-1M-StYUC10-Soltu.DM.09G022830.1._Stuberosum 3.83e-192
Motif Symbol Motif Consensus
1. DCIASLWQKRTYDRLKLHLPKQFCZLPLMPFPE
2. FRGKKVLVVGCGNSGMEVSLDLCN
3. WKGENGLYAVGFTRRGLLGASMDAVRIAZDI
4. PTYPTKQQFIDYLESYAAHFDIRPRFNETVZ
5. FVDGREEEFDAIILATGYKSNVPSWL
6. GBTSKYGJKRPKIGPLELKNKTGKTPVLD
7. IIVGAGPSGLATAACLKEQGVP
8. EYICRWLVVATGENAEPVVPEI
10. VGTLAKIKSGDIKVVP
11. LLKWLPLWLVDKLLLLLA
12. DFFSKDGFPKTPFPNG
13. GEVIHSSDYKSG
14. SAEYDETSGLWRVKTV
15. HGANTSIVVRSPVHVLTKEMVY
16. SVILERA
21. EGLEEFG
22. IKRFTRGGV
24. GNSVVFEBGKSHQADVIVFATG
26. ECLTPHGAK
29. PTSRCQAQPEHDNHWHHPTIDH

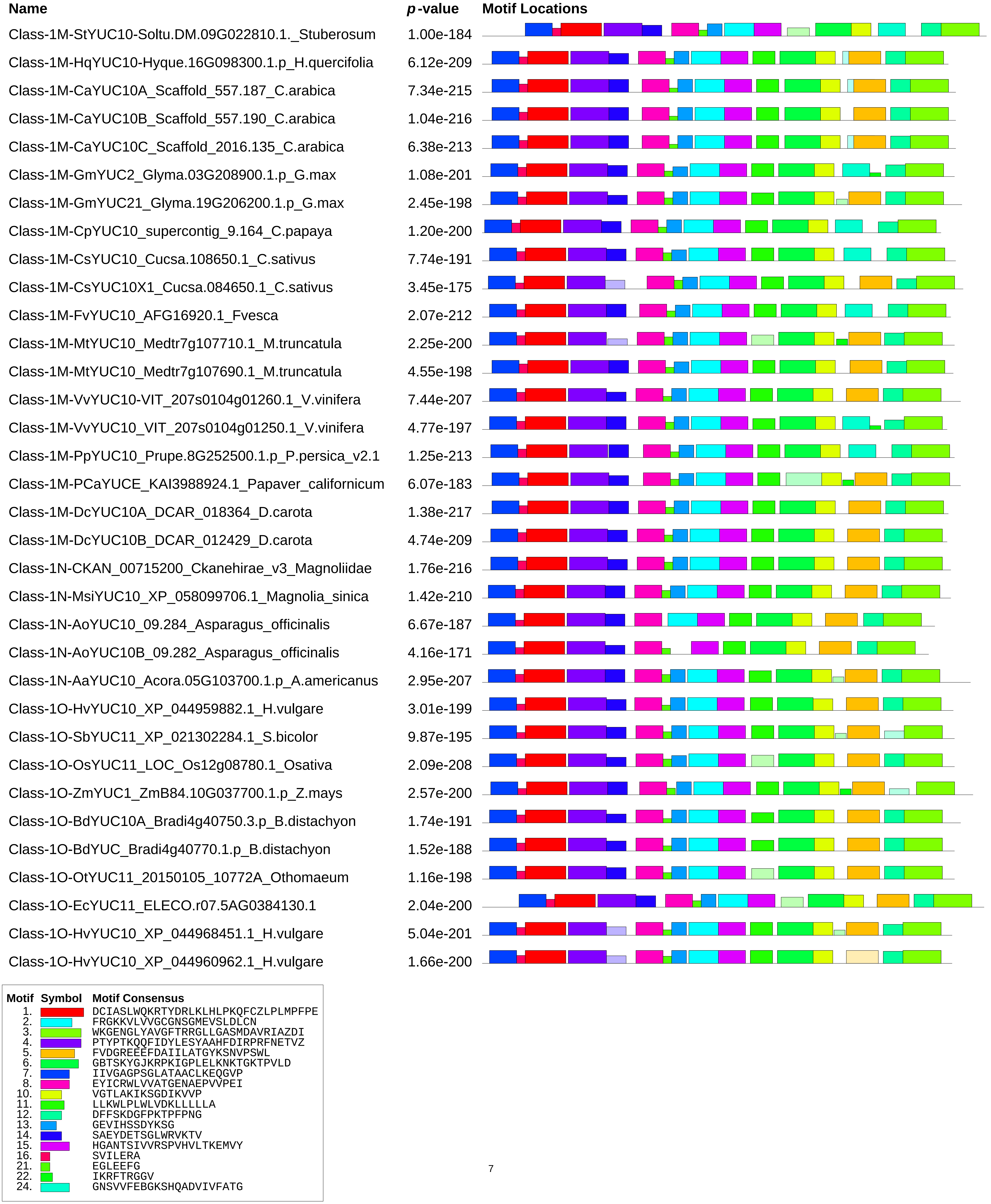
Name
p-value
Motif Locations
Class-1M-StYUC10-Soltu.DM.09G022810.1._Stuberosum 1.00e-184
Class-1M-HqYUC10-Hyque.16G098300.1.p_H.quercifolia 6.12e-209
Class-1M-CaYUC10A_Scaffold_557.187_C.arabica 7.34e-215
Class-1M-CaYUC10B_Scaffold_557.190_C.arabica 1.04e-216
Class-1M-CaYUC10C_Scaffold_2016.135_C.arabica 6.38e-213
Class-1M-GmYUC2_Glyma.03G208900.1.p_G.max 1.08e-201
Class-1M-GmYUC21_Glyma.19G206200.1.p_G.max 2.45e-198
Class-1M-CpYUC10_supercontig_9.164_C.papaya 1.20e-200
Class-1M-CsYUC10_Cucsa.108650.1_C.sativus 7.74e-191
Class-1M-CsYUC10X1_Cucsa.084650.1_C.sativus 3.45e-175
Class-1M-FvYUC10_AFG16920.1_Fvesca 2.07e-212
Class-1M-MtYUC10_Medtr7g107710.1_M.truncatula 2.25e-200
Class-1M-MtYUC10_Medtr7g107690.1_M.truncatula 4.55e-198
Class-1M-VvYUC10-VIT_207s0104g01260.1_V.vinifera 7.44e-207
Class-1M-VvYUC10_VIT_207s0104g01250.1_V.vinifera 4.77e-197
Class-1M-PpYUC10_Prupe.8G252500.1.p_P.persica_v2.1 1.25e-213
Class-1M-PCaYUCE_KAI3988924.1_Papaver_californicum 6.07e-183
Class-1M-DcYUC10A_DCAR_018364_D.carota 1.38e-217
Class-1M-DcYUC10B_DCAR_012429_D.carota 4.74e-209
Class-1N-CKAN_00715200_Ckanehirae_v3_Magnoliidae 1.76e-216
Class-1N-MsiYUC10_XP_058099706.1_Magnolia_sinica 1.42e-210
Class-1N-AoYUC10_09.284_Asparagus_officinalis 6.67e-187
Class-1N-AoYUC10B_09.282_Asparagus_officinalis 4.16e-171
Class-1N-AaYUC10_Acora.05G103700.1.p_A.americanus 2.95e-207
Class-1O-HvYUC10_XP_044959882.1_H.vulgare 3.01e-199
Class-1O-SbYUC11_XP_021302284.1_S.bicolor 9.87e-195
Class-1O-OsYUC11_LOC_Os12g08780.1_Osativa 2.09e-208
Class-1O-ZmYUC1_ZmB84.10G037700.1.p_Z.mays 2.57e-200
Class-1O-BdYUC10A_Bradi4g40750.3.p_B.distachyon 1.74e-191
Class-1O-BdYUC_Bradi4g40770.1.p_B.distachyon 1.52e-188
Class-1O-OtYUC11_20150105_10772A_Othomaeum 1.16e-198
Class-1O-EcYUC11_ELECO.r07.5AG0384130.1 2.04e-200
Class-1O-HvYUC10_XP_044968451.1_H.vulgare 5.04e-201
Class-1O-HvYUC10_XP_044960962.1_H.vulgare 1.66e-200
Motif Symbol Motif Consensus
1. DCIASLWQKRTYDRLKLHLPKQFCZLPLMPFPE
2. FRGKKVLVVGCGNSGMEVSLDLCN
3. WKGENGLYAVGFTRRGLLGASMDAVRIAZDI
4. PTYPTKQQFIDYLESYAAHFDIRPRFNETVZ
5. FVDGREEEFDAIILATGYKSNVPSWL
6. GBTSKYGJKRPKIGPLELKNKTGKTPVLD
7. IIVGAGPSGLATAACLKEQGVP
8. EYICRWLVVATGENAEPVVPEI
10. VGTLAKIKSGDIKVVP
11. LLKWLPLWLVDKLLLLLA
12. DFFSKDGFPKTPFPNG
13. GEVIHSSDYKSG
14. SAEYDETSGLWRVKTV
15. HGANTSIVVRSPVHVLTKEMVY
16. SVILERA
21. EGLEEFG
22. IKRFTRGGV
24. GNSVVFEBGKSHQADVIVFATG

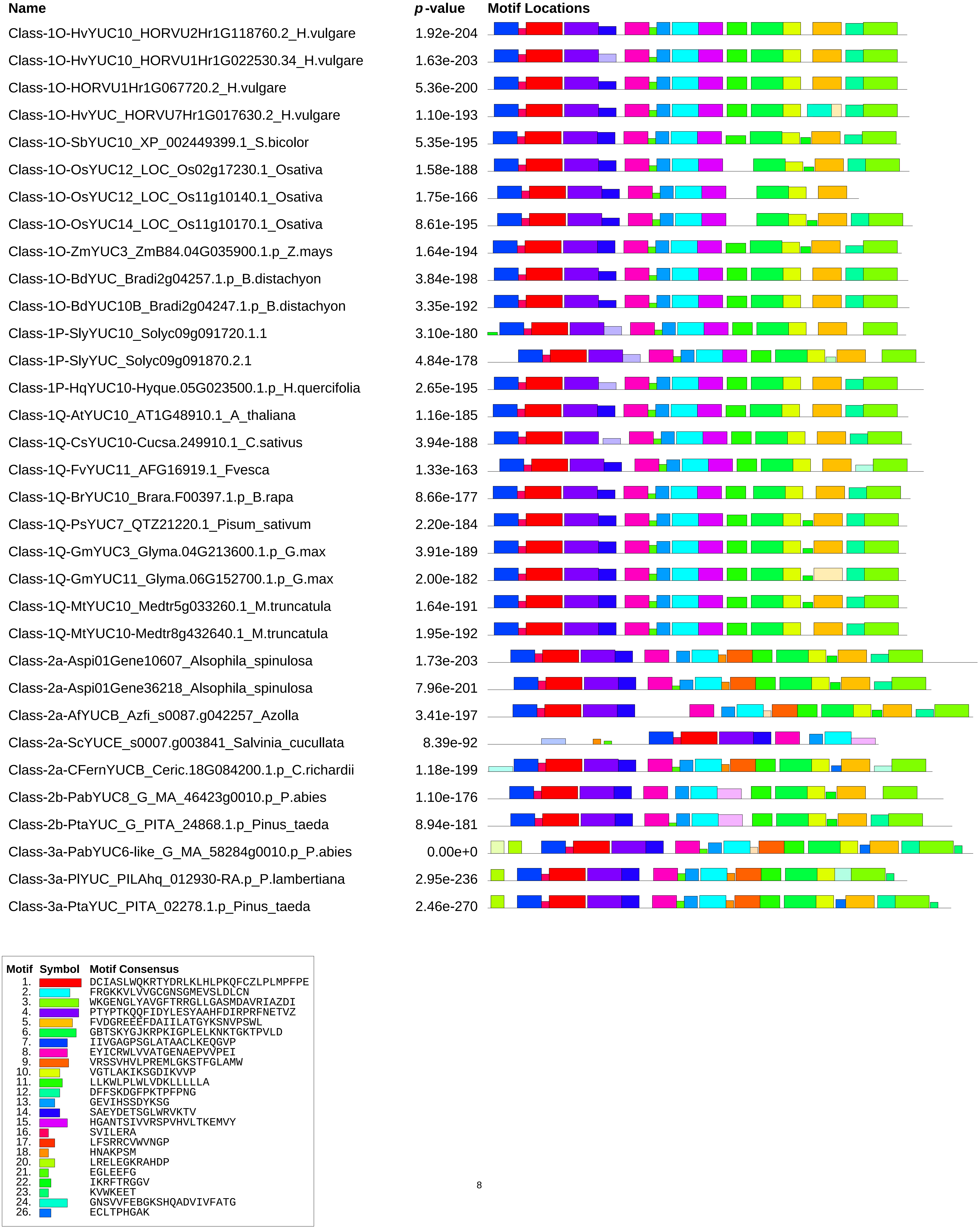
Name
p-value
Motif Locations
Class-1O-HvYUC10_HORVU2Hr1G118760.2_H.vulgare 1.92e-204
Class-1O-HvYUC10_HORVU1Hr1G022530.34_H.vulgare 1.63e-203
Class-1O-HORVU1Hr1G067720.2_H.vulgare 5.36e-200
Class-1O-HvYUC_HORVU7Hr1G017630.2_H.vulgare 1.10e-193
Class-1O-SbYUC10_XP_002449399.1_S.bicolor 5.35e-195
Class-1O-OsYUC12_LOC_Os02g17230.1_Osativa 1.58e-188
Class-1O-OsYUC12_LOC_Os11g10140.1_Osativa 1.75e-166
Class-1O-OsYUC14_LOC_Os11g10170.1_Osativa 8.61e-195
Class-1O-ZmYUC3_ZmB84.04G035900.1.p_Z.mays 1.64e-194
Class-1O-BdYUC_Bradi2g04257.1.p_B.distachyon 3.84e-198
Class-1O-BdYUC10B_Bradi2g04247.1.p_B.distachyon 3.35e-192
Class-1P-SlyYUC10_Solyc09g091720.1.1 3.10e-180
Class-1P-SlyYUC_Solyc09g091870.2.1 4.84e-178
Class-1P-HqYUC10-Hyque.05G023500.1.p_H.quercifolia 2.65e-195
Class-1Q-AtYUC10_AT1G48910.1_A_thaliana 1.16e-185
Class-1Q-CsYUC10-Cucsa.249910.1_C.sativus 3.94e-188
Class-1Q-FvYUC11_AFG16919.1_Fvesca 1.33e-163
Class-1Q-BrYUC10_Brara.F00397.1.p_B.rapa 8.66e-177
Class-1Q-PsYUC7_QTZ21220.1_Pisum_sativum 2.20e-184
Class-1Q-GmYUC3_Glyma.04G213600.1.p_G.max 3.91e-189
Class-1Q-GmYUC11_Glyma.06G152700.1.p_G.max 2.00e-182
Class-1Q-MtYUC10_Medtr5g033260.1_M.truncatula 1.64e-191
Class-1Q-MtYUC10-Medtr8g432640.1_M.truncatula 1.95e-192
Class-2a-Aspi01Gene10607_Alsophila_spinulosa 1.73e-203
Class-2a-Aspi01Gene36218_Alsophila_spinulosa 7.96e-201
Class-2a-AfYUCB_Azfi_s0087.g042257_Azolla 3.41e-197
Class-2a-ScYUCE_s0007.g003841_Salvinia_cucullata 8.39e-92
Class-2a-CFernYUCB_Ceric.18G084200.1.p_C.richardii 1.18e-199
Class-2b-PabYUC8_G_MA_46423g0010.p_P.abies 1.10e-176
Class-2b-PtaYUC_G_PITA_24868.1.p_Pinus_taeda 8.94e-181
Class-3a-PabYUC6-like_G_MA_58284g0010.p_P.abies 0.00e+0
Class-3a-PlYUC_PILAhq_012930-RA.p_P.lambertiana 2.95e-236
Class-3a-PtaYUC_PITA_02278.1.p_Pinus_taeda 2.46e-270
Motif Symbol Motif Consensus
1. DCIASLWQKRTYDRLKLHLPKQFCZLPLMPFPE
2. FRGKKVLVVGCGNSGMEVSLDLCN
3. WKGENGLYAVGFTRRGLLGASMDAVRIAZDI
4. PTYPTKQQFIDYLESYAAHFDIRPRFNETVZ
5. FVDGREEEFDAIILATGYKSNVPSWL
6. GBTSKYGJKRPKIGPLELKNKTGKTPVLD
7. IIVGAGPSGLATAACLKEQGVP
8. EYICRWLVVATGENAEPVVPEI
9. VRSSVHVLPREMLGKSTFGLAMW
10. VGTLAKIKSGDIKVVP
11. LLKWLPLWLVDKLLLLLA
12. DFFSKDGFPKTPFPNG
13. GEVIHSSDYKSG
14. SAEYDETSGLWRVKTV
15. HGANTSIVVRSPVHVLTKEMVY
16. SVILERA
17. LFSRRCVWVNGP
18. HNAKPSM
20. LRELEGKRAHDP
21. EGLEEFG
22. IKRFTRGGV
23. KVWKEET
24. GNSVVFEBGKSHQADVIVFATG
26. ECLTPHGAK

| Name | *p*-value | Motif Locations |
|---|---|---|
| Class-3a-GmYUCD_TnS000901889t02.p_Gnetum_montanum | 4.58e-247 | |
| Class-3a-TpYUCE_Thupl.29382193s0012.1.p_T.plicata | 0.00e+0 | |
| Class-3b-HvYUC1L_XP_044948194.1_H.vulgare | 4.92e-230 | |
| Class-3b-HvYUC1_HORVU5Hr1G028100.1_H.vulgare | 4.15e-188 | |
| Class-3b-SbYUC6-XP002442221.1_S.bicolor | 2.16e-268 | |
| Class-3b-SbYUC6-XP021301614.1_S.bicolor | 3.91e-224 | |
| Class-3b-OsYUC5_LOCOs12g32750.1_Osativa | 3.46e-230 | |
| Class-3b-CKAN_01428800_Ckanehirae_v3_Magnoliidae | 7.88e-237 | |
| Class-3b-PtYUC5_Potri.007G028200.1.p_Ptrichocarpa | 2.27e-241 | |
| Class-3b-MaYUC2-Achr4P14020_001_M.acuminata | 3.00e-212 | |
| Class-3b-MaYUC2-like_Achr4P03480_001_M.acuminata | 6.58e-200 | |
| Class-3b-AoYUC2-08.1148_Asparagus_officinalis | 4.84e-252 | |
| Class-3b-BdYUC6_Bradi4g06427.1.p_B.distachyon | 6.52e-234 | |
| Class-3b-AcYUC2_Aco003772.1_A.comosus_v3 | 3.64e-255 | |
| Class-3b-AaYUC2-Acora.06G229000.1.p_A.americanus | 5.05e-247 | |
| Class-3b-HqYUC6-Hyque.05G088700.1.p_H.quercifolia | 5.61e-232 | |
| Class-3b-EcYUC1-ELECO.r07.5BG0439230.1 | 3.90e-232 | |
| Class-3b-EcYUC_ELECO.r07.5AG0392180.1 | 1.87e-237 | |
| Class-3b-VvYUC6_VIT_204s0023g01480.1_V.vinifera | 6.36e-232 | |
| Class-3b-MsiYUC6-XP_058069069.1_Magnolia_sinica | 3.32e-259 | |
| Class-3b-NcYUC6-like_Nycol.I00883.1.p_N.colorata | 2.90e-261 | |
| Class-3b-PCaYUCB_KAI3922319.1_Papaver_californicum | 1.21e-244 | |
| Class-3b-PCaYUCC_KAI3946133.1_Papaver_californicum | 2.39e-245 | |
| Class-3c-AmTrYUC6_scaffold00218.8_A.trichopoda | 0.00e+0 | |
| Class-3c-AaYUC6-Acora.01G123400.1.p_A.americanus | 0.00e+0 | |
| Class-3c-AaYUC6-Acora.07G067600.1.p_A.americanus | 2.90e-264 | |
| Class-3c-NcYUC2-Nycol.A02312.1.p_N.colorata | 0.00e+0 | |
| Class-3c-AoYUC2-01.1621_Asparagus_officinalis | 0.00e+0 | |
| Class-3c-MaYUC2-Achr2P19930_M.acuminata | 9.69e-243 | |
| Class-3c-MaYUC6-Achr5P06860_M.acuminata | 2.71e-222 | |
| Class-3c-MaYUC2-Achr11P12150_M.acuminata | 3.88e-233 | |
| Class-3c-MaYUC2-Achr10P22060_M.acuminata | 1.54e-211 | |
| Class-3d-HvYUC2_XP_044972712.1_H.vulgare | 5.66e-265 | |

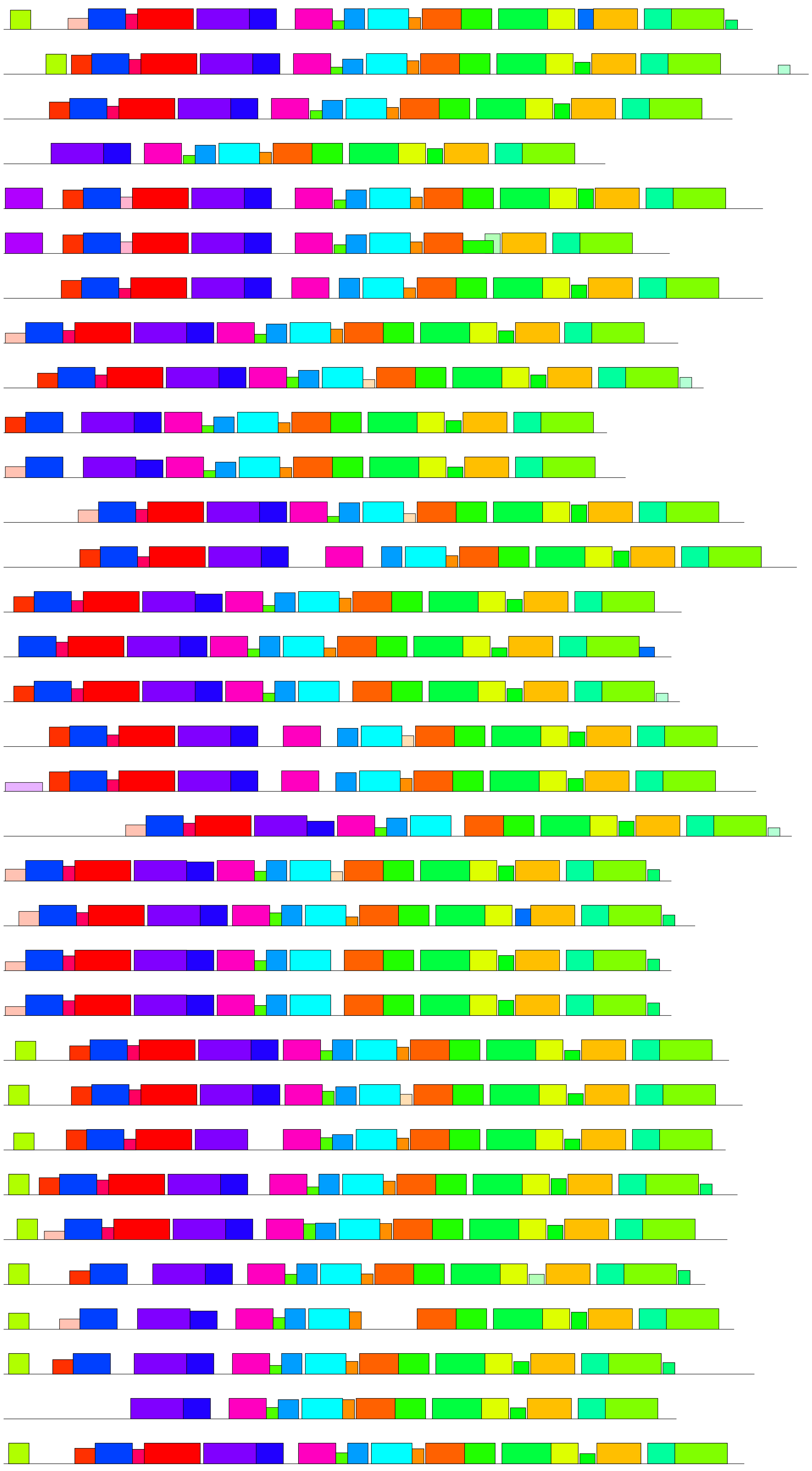

| Motif | Symbol | Motif Consensus |
|---|---|---|
| 1. | | DCIASLWQKRTYDRLKLHLPKQFCZLPLMPFPE |
| 2. | | FRGKKVLVVGCGNSGMEVSLDLCN |
| 3. | | WKGENGLYAVGFTRRGLLGASMDAVRIAZDI |
| 4. | | PTYPTKQQFIDYLESYAAHFDIRPRFNETVZ |
| 5. | | FVDGREEEFDAIILATGYKSNVPSWL |
| 6. | | GBTSKYGJKRPKIGPLELKNKTGKTPVLD |
| 7. | | IIVGAGPSGLATAACLKEQGVP |
| 8. | | EYICRWLVVATGENAEPVVPEI |
| 9. | | VRSSVHVLPREMLGKSTFGLAMW |
| 10. | | VGTLAKIKSGDIKVVP |
| 11. | | LLKWLPLWLVDKLLLLLA |
| 12. | | DFFSKDGFPKTPFPNG |
| 13. | | GEVIHSSDYKSG |
| 14. | | SAEYDETSGLWRVKTV |
| 16. | | SVILERA |
| 17. | | LFSRRCVWVNGP |
| 18. | | HNAKPSM |
| 20. | | LRELEGKRAHDP |
| 21. | | EGLEEFG |
| 22. | | IKRFTRGGV |
| 23. | | KVWKEET |
| 26. | | ECLTPHGAK |
| 29. | | PTSRCQAQPEHDNHWHHPTIDH |

| Name | *p*-value | Motif Locations |
|---|---|---|
| Class-3d-HuYUC4_XP_044972289.1_H.vulgare | 1.29e-225 | |
| Class-3d-SbYUC2_XP_002442461.2_S.bicolor | 8.43e-227 | |
| Class-3d-SbYUC2_XP_002457104.1_S.bicolor | 7.01e-236 | |
| Class-3d-SbYUC2_XP_021313730.1_S.bicolor | 4.05e-265 | |
| Class-3d-OsYUC2_LOC_Os05g45240.1_Osativa | 4.20e-204 | |
| Class-3d-OsYUC3_LOC_Os01g53200.1_Osativa | 0.00e+0 | |
| Class-3d-OsYUC4_LOC_Os01g12490.1_Osativa | 1.18e-236 | |
| Class-3d-ZmYUC6_ZmB84.08G231900.1.p_Z.mays | 1.73e-264 | |
| Class-3d-ZmYUC8_ZmB84.03G007700.1.p_Z.mays | 1.95e-238 | |
| Class-3d-ZmYUC10_ZmB84.03G136400.1.p_Z.mays | 2.39e-221 | |
| Class-3d-ZmYUC12_NP_001149353.1_Zea_mays | 1.17e-219 | |
| Class-3d-MaYUC2_Achr9P05490_001_M.acuminata | 9.51e-268 | |
| Class-3d-AoYUC2-01.690_Asparagus_officinalis | 6.80e-258 | |
| Class-3d-BdYUC2_Bradi2g07407.3.p_B.distachyon | 1.37e-227 | |
| Class-3d-AcYUC2-like_Aco010742.1_A.comosus_v3 | 4.60e-270 | |
| Class-3d-AcYUC2-like_Aco004396.1_A.comosus_v3 | 1.06e-251 | |
| Class-3d-AcYUC2-like_Aco005574.1_A.comosus_v3 | 1.34e-244 | |
| Class-3d-OtYUC2-20150105_17835A_Othomaeum | 2.44e-231 | |
| Class-3d-OtYUC2-like_20150105_12412A_Othomaeum | 9.22e-271 | |
| Class-3d-EcYUC2-ELECO.r07.1AG0032120.1 | 3.36e-265 | |
| Class-3d-EcYUC2-ELECO.r07.1BG0082330.1 | 8.42e-270 | |
| Class-3d-EcYUC2-ELECO.r07.1AG0001270.1 | 5.54e-245 | |
| Class-3d-EcYUC2-like_ELECO.r07.1BG0049720.1 | 1.55e-245 | |
| Class-3e-AtYUC2_AT4G13260.1_A_thaliana | 1.24e-255 | |
| Class-3e-AtYUC6A_AT5G25620.1_A_thaliana | 0.00e+0 | |
| Class-3e-AtYUC6B_AT5G25620.2_A_thaliana | 0.00e+0 | |
| Class-3e-PsYUC2_QTZ21215.1_Pisum_sativum | 0.00e+0 | |
| Class-3e-PsYUC3_QTZ21216.1_Pisum_sativum | 0.00e+0 | |
| Class-3e-CKAN_00588900_Ckanehirae | 9.46e-264 | |
| Class-3e-PtYUC2_Potri.018G036800.1.p_Ptrichocarpa | 0.00e+0 | |
| Class-3e-PtYUC6_Potri.006G243400.1.p_Ptrichocarpa | 0.00e+0 | |
| Class-3e-SlyYUC2-Solyc08g068160.2.1_S.lycopersicum | 0.00e+0 | |
| Class-3e-StYUC2-Soltu.DM.08G017290.1_S.tuberosum | 0.00e+0 | |
| Class-3e-DcYUC2-DCAR_028710_D.carota | 1.85e-219 | |

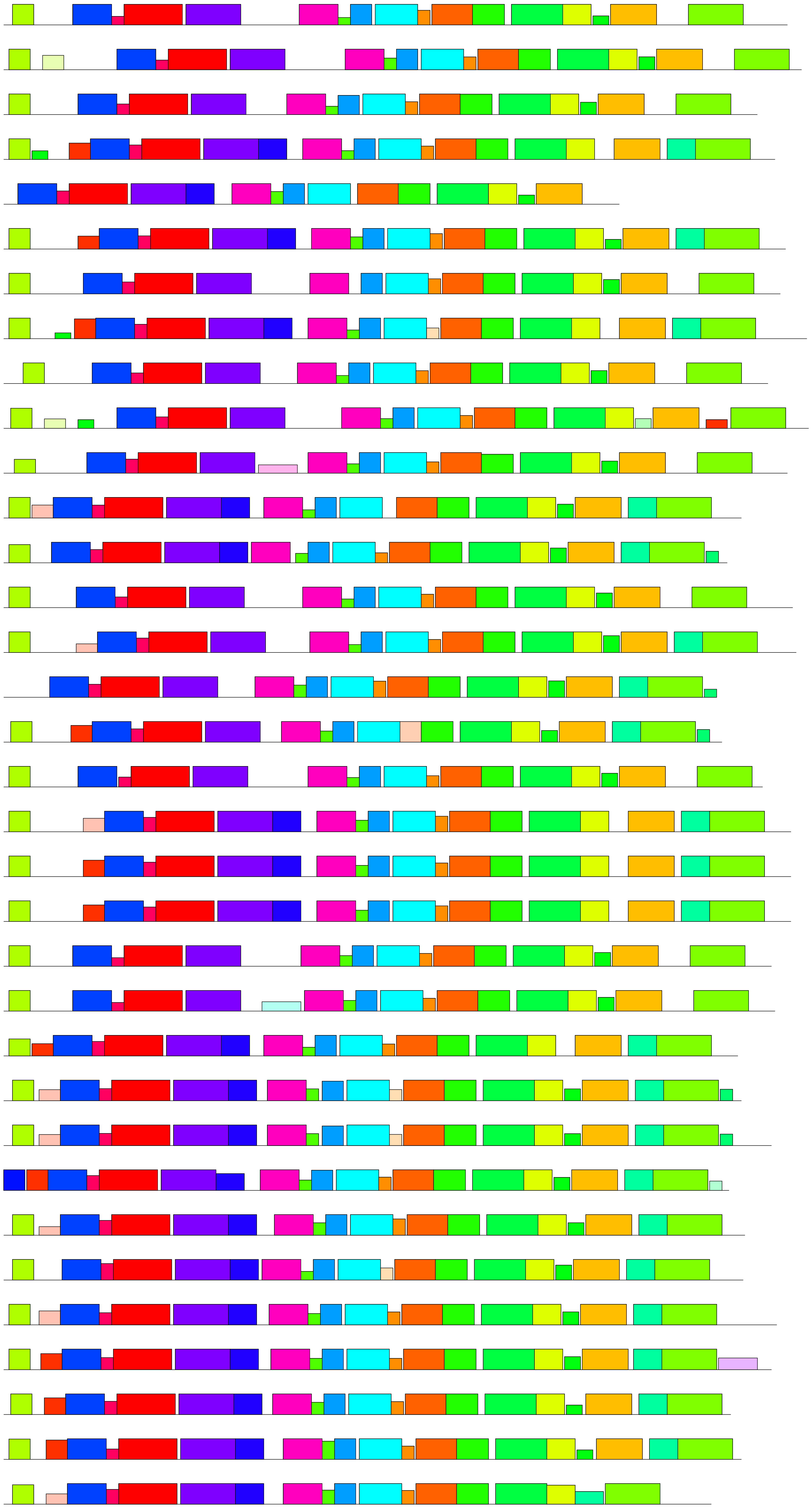

| Motif | Symbol | Motif Consensus |
|---|---|---|
| 1. | | DCIASLWQKRTYDRLKLHLPKQFCZLPLMPFPE |
| 2. | | FRGKKVLVVGCGNSGMEVSLDLCN |
| 3. | | WKGENGLYAVGFTRRGLLGASMDAVRIAZDI |
| 4. | | PTYPTKQQFIDYLESYAAHFDIRPRFNETVZ |
| 5. | | FVDGREEEFDAIILATGYKSNVPSWL |
| 6. | | GBTSKYGJKRPKIGPLELKNKTGKTPVLD |
| 7. | | IIVGAGPSGLATAACLKEQGVP |
| 8. | | EYICRWLVVATGENAEPVVPEI |
| 9. | | VRSSVHVLPREMLGKSTFGLAMW |
| 10. | | VGTLAKIKSGDIKVVP |
| 11. | | LLKWLPLWLVDKLLLLLA |
| 12. | | DFFSKDGFPKTPFPNG |
| 13. | | GEVIHSSDYKSG |
| 14. | | SAEYDETSGLWRVKTV |
| 16. | | SVILERA |
| 17. | | LFSRRCVWVNGP |
| 18. | | HNAKPSM |
| 20. | | LRELEGKRAHDP |
| 21. | | EGLEEFG |
| 22. | | IKRFTRGGV |
| 23. | | KVWKEET |
| 24. | | GNSVVFEBGKSHQADVIVFATG |
| 27. | | MENMFRLVDHED |
| 29. | | PTSRCQAQPEHDNHWHHPTIDH |

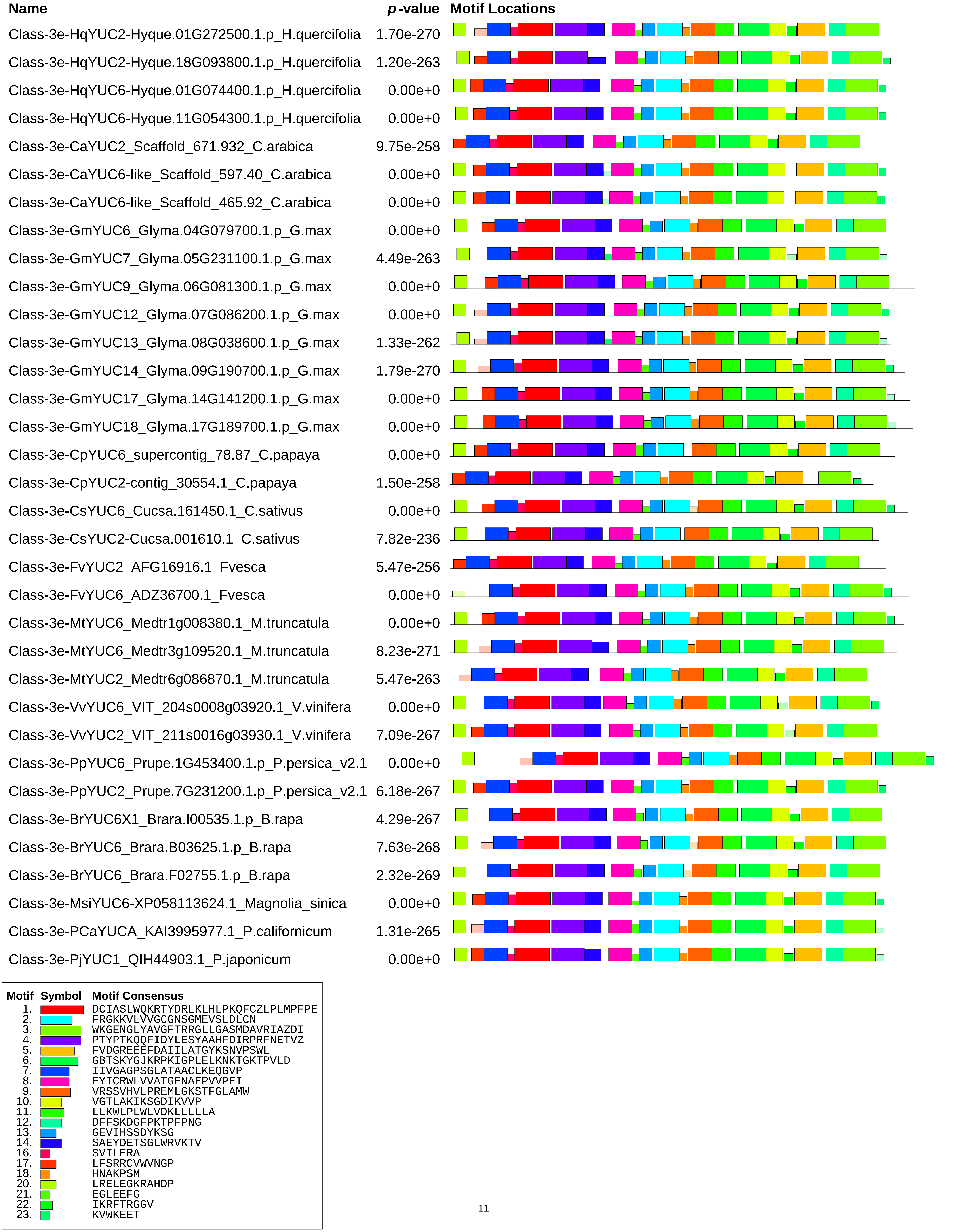

Name
p-value
Motif Locations
Class-3e-HqYUC2-Hyque.01G272500.1.p_H.quercifolia 1.70e-270
Class-3e-HqYUC2-Hyque.18G093800.1.p_H.quercifolia 1.20e-263
Class-3e-HqYUC6-Hyque.01G074400.1.p_H.quercifolia 0.00e+0
Class-3e-HqYUC6-Hyque.11G054300.1.p_H.quercifolia 0.00e+0
Class-3e-CaYUC2_Scaffold_671.932_C.arabica 9.75e-258
Class-3e-CaYUC6-like_Scaffold_597.40_C.arabica 0.00e+0
Class-3e-CaYUC6-like_Scaffold_465.92_C.arabica 0.00e+0
Class-3e-GmYUC6_Glyma.04G079700.1.p_G.max 0.00e+0
Class-3e-GmYUC7_Glyma.05G231100.1.p_G.max 4.49e-263
Class-3e-GmYUC9_Glyma.06G081300.1.p_G.max 0.00e+0
Class-3e-GmYUC12_Glyma.07G086200.1.p_G.max 0.00e+0
Class-3e-GmYUC13_Glyma.08G038600.1.p_G.max 1.33e-262
Class-3e-GmYUC14_Glyma.09G190700.1.p_G.max 1.79e-270
Class-3e-GmYUC17_Glyma.14G141200.1.p_G.max 0.00e+0
Class-3e-GmYUC18_Glyma.17G189700.1.p_G.max 0.00e+0
Class-3e-CpYUC6_supercontig_78.87_C.papaya 0.00e+0
Class-3e-CpYUC2-contig_30554.1_C.papaya 1.50e-258
Class-3e-CsYUC6_Cucsa.161450.1_C.sativus 0.00e+0
Class-3e-CsYUC2-Cucsa.001610.1_C.sativus 7.82e-236
Class-3e-FvYUC2_AFG16916.1_Fvesca 5.47e-256
Class-3e-FvYUC6_ADZ36700.1_Fvesca 0.00e+0
Class-3e-MtYUC6_Medtr1g008380.1_M.truncatula 0.00e+0
Class-3e-MtYUC6_Medtr3g109520.1_M.truncatula 8.23e-271
Class-3e-MtYUC2_Medtr6g086870.1_M.truncatula 5.47e-263
Class-3e-VvYUC6_VIT_204s0008g03920.1_V.vinifera 0.00e+0
Class-3e-VvYUC2_VIT_211s0016g03930.1_V.vinifera 7.09e-267
Class-3e-PpYUC6_Prupe.1G453400.1.p_P.persica_v2.1 0.00e+0
Class-3e-PpYUC2_Prupe.7G231200.1.p_P.persica_v2.1 6.18e-267
Class-3e-BrYUC6X1_Brara.I00535.1.p_B.rapa 4.29e-267
Class-3e-BrYUC6_Brara.B03625.1.p_B.rapa 7.63e-268
Class-3e-BrYUC6_Brara.F02755.1.p_B.rapa 2.32e-269
Class-3e-MsiYUC6-XP058113624.1_Magnolia_sinica 0.00e+0
Class-3e-PCaYUCA_KAI3995977.1_P.californicum 1.31e-265
Class-3e-PjYUC1_QIH44903.1_P.japonicum 0.00e+0
Motif Symbol Motif Consensus
1. DCIASLWQKRTYDRLKLHLPKQFCZLPLMPFPE
2. FRGKKVLVVGCGNSGMEVSLDLCN
3. WKGENGLYAVGFTRRGLLGASMDAVRIAZDI
4. PTYPTKQQFIDYLESYAAHFDIRPRFNETVZ
5. FVDGREEEFDAIILATGYKSNVPSWL
6. GBTSKYGJKRPKIGPLELKNKTGKTPVLD
7. IIVGAGPSGLATAACLKEQGVP
8. EYICRWLVVATGENAEPVVPEI
9. VRSSVHVLPREMLGKSTFGLAMW
10. VGTLAKIKSGDIKVVP
11. LLKWLPLWLVDKLLLLLA
12. DFFSKDGFPKTPFPNG
13. GEVIHSSDYKSG
14. SAEYDETSGLWRVKTV
16. SVILERA
17. LFSRRCVWVNGP
18. HNAKPSM
20. LRELEGKRAHDP
21. EGLEEFG
22. IKRFTRGGV
23. KVWKEET

| Name | *p*-value |
|---|---|
| Class-4A-GbYUCB_Gb32507.p_G.biloba | 0.00e+0 |
| Class-4A-PlYUC_PILAhq064253-RA.p_P.lambertiana | 6.77e-204 |
| Class-4A-PtaYUC_PITA32685.1.p_P.taeda | 0.00e+0 |
| Class-4A-PtaYUC_PITA21706.1.p_P.taeda | 0.00e+0 |
| Class-4A-TpYUCD_Thupl.29380002s0001.1.p_T.plicata | 3.21e-271 |
| Class-4B-PabYUC5_MA_218308g0010.p_P.abies | 0.00e+0 |
| Class-4B-PabYUC_MA_5857317g0010.p_P.abies | 1.81e-186 |
| Class-4B-GbYUCA_Gb_39502.p_Ginkgo_biloba | 0.00e+0 |
| Class-4B-PlYUC_PILAhq_045322-RA.p_P.lambertiana | 0.00e+0 |
| Class-4B-PlYUC_PILAhq_038950-RA.p_P.lambertiana | 0.00e+0 |
| Class-4B-PtaYUC_PITA_06172.1.p_Pinus_taeda | 0.00e+0 |
| Class-4B-PtaYUC_PITA_03225.1.p_Pinus_taeda | 0.00e+0 |
| Class-4B-GmYUCC_TnS000183447t02.p_Gnetum_montanum | 2.24e-246 |
| Class-4c-PabYUCB/YUC5-G-MA-215174g0010.p-P.abies | 2.83e-225 |
| Class-4c-GbYUCC-G-Gb-40961.p-Ginkgo-biloba | 0.00e+0 |
| Class-4c-PlYUC-G-PILAhq-075634-RA.p-P.lambertiana | 0.00e+0 |
| Class-4c-PtaYUC-G-PITA-05784.1.p-P.taeda | 0.00e+0 |
| Class-4c-GmYUCA-G-TnS000225445t01.p-G.montanum | 0.00e+0 |
| Class-4c-TpYUCB-Thupl.29377126s0001.1.p-T.plicata | 0.00e+0 |
| Class-4D-AmTrYUC_scaffold00039.187_A.trichopoda | 0.00e+0 |
| Class-4D-CKAN-01805100-Ckanehirae-v3-Magnoliidae | 0.00e+0 |
| Class-4D-CKAN-02673000-Ckanehirae-v3-Magnoliidae | 0.00e+0 |
| Class-4D-CKAN-02545400-Ckanehirae-v3-Magnoliidae | 0.00e+0 |
| Class-4D-CKAN-01519000-Ckanehirae-v3-Magnoliidae | 0.00e+0 |
| Class-4D-MsiYUC5-XP-058095048.1-MagnoliA-sinica | 0.00e+0 |
| Class-4D-MsiYUC5-XP-058101305.1-MagnoliA-sinica | 0.00e+0 |
| Class-4D-NcYUC5_Nycol.L00992.1.p_N.colorata | 0.00e+0 |
| Class-4D-NcYUC5_Nycol.D01636.1.p_N.colorata | 0.00e+0 |
| Class-4E-HvYUC3_KAE8779170.1_Hordeum_vulgare | 5.19e-228 |
| Class-4E-HvYUC5_XP_044971244.1_H.vulgare | 0.00e+0 |
| Class-4E-HvYUC5_XP_044970914.1_H.vulgare | 1.69e-253 |
| Class-4E-HvYUC5_HORVU2Hr1G116980.1_H.vulgare | 8.96e-252 |
| Class-4E-HvYUC9_XP_044950031.1_H.vulgare | 1.99e-234 |

**Motif Locations**

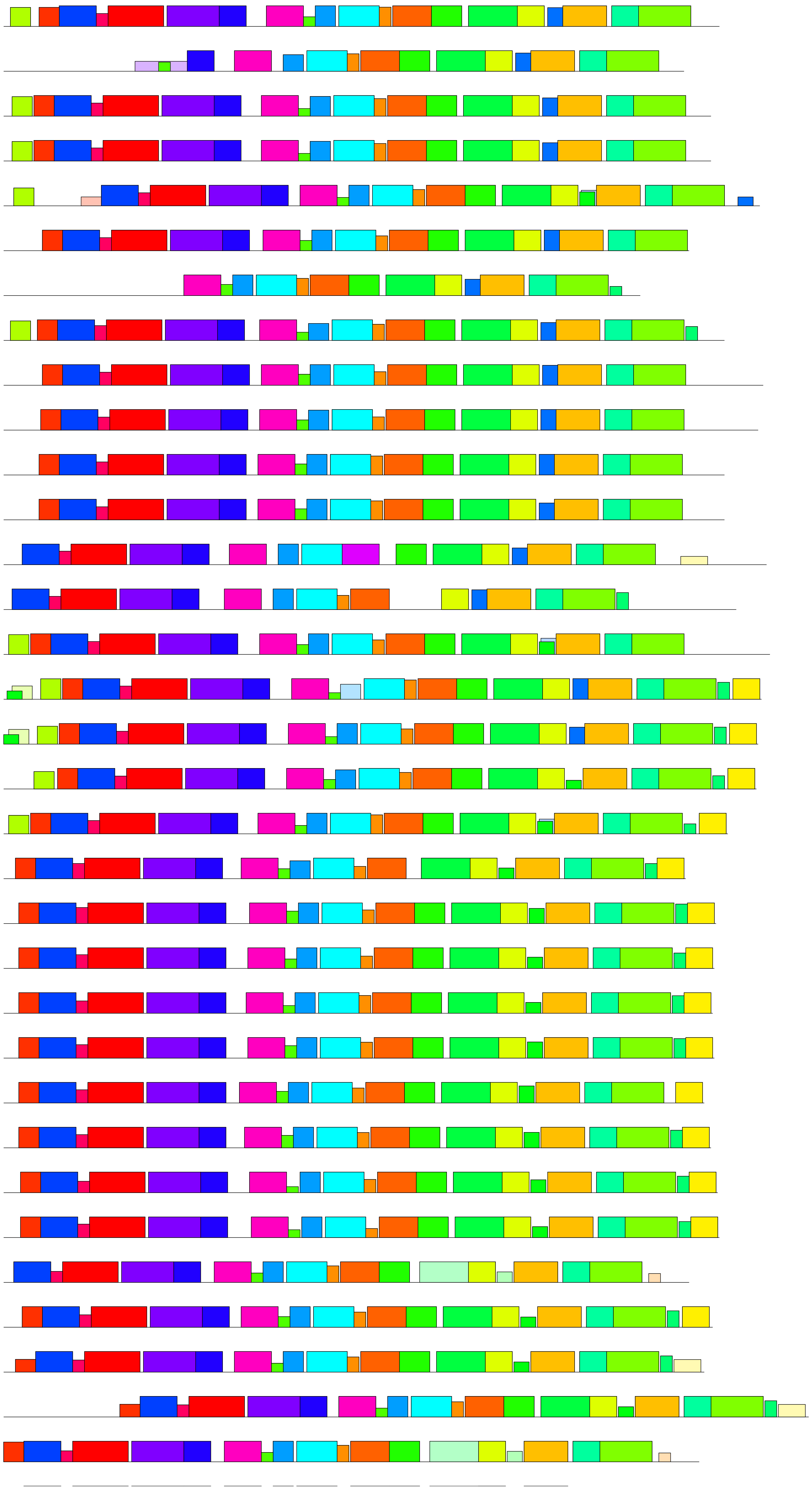

| Motif | Symbol | Motif Consensus |
|---|---|---|
| 1. | | DCIASLWQKRTYDRLKLHLPKQFCZLPLMPFPE |
| 2. | | FRGKKVLVVGCGNSGMEVSLDLCN |
| 3. | | WKGENGLYAVGFTRRGLLGASMDAVRIAZDI |
| 4. | | PTYPTKQQFIDYLESYAAHFDIRPRFNETVZ |
| 5. | | FVDGREEEFDAIILATGYKSNVPSWL |
| 6. | | GBTSKYGJKRPKIGPLELKNKTGKTPVLD |
| 7. | | IIVGAGPSGLATAACLKEQGVP |
| 8. | | EYICRWLVVATGENAEPVVPEI |
| 9. | | VRSSVHVLPREMLGKSTFGLAMW |
| 10. | | VGTLAKIKSGDIKVVP |
| 11. | | LLKWLPLWLVDKLLLLLA |
| 12. | | DFFSKDGFPKTPFPNG |
| 13. | | GEVIHSSDYKSG |
| 14. | | SAEYDETSGLWRVKTV |
| 15. | | HGANTSIVVRSPVHVLTKEMVY |
| 16. | | SVILERA |
| 17. | | LFSRRCVWVNGP |
| 18. | | HNAKPSM |
| 19. | | KQKKRFVACHRRCISQ |
| 20. | | LRELEGKRAHDP |
| 21. | | EGLEEFG |
| 22. | | IKRFTRGGV |
| 23. | | KVWKEET |
| 26. | | ECLTPHGAK |

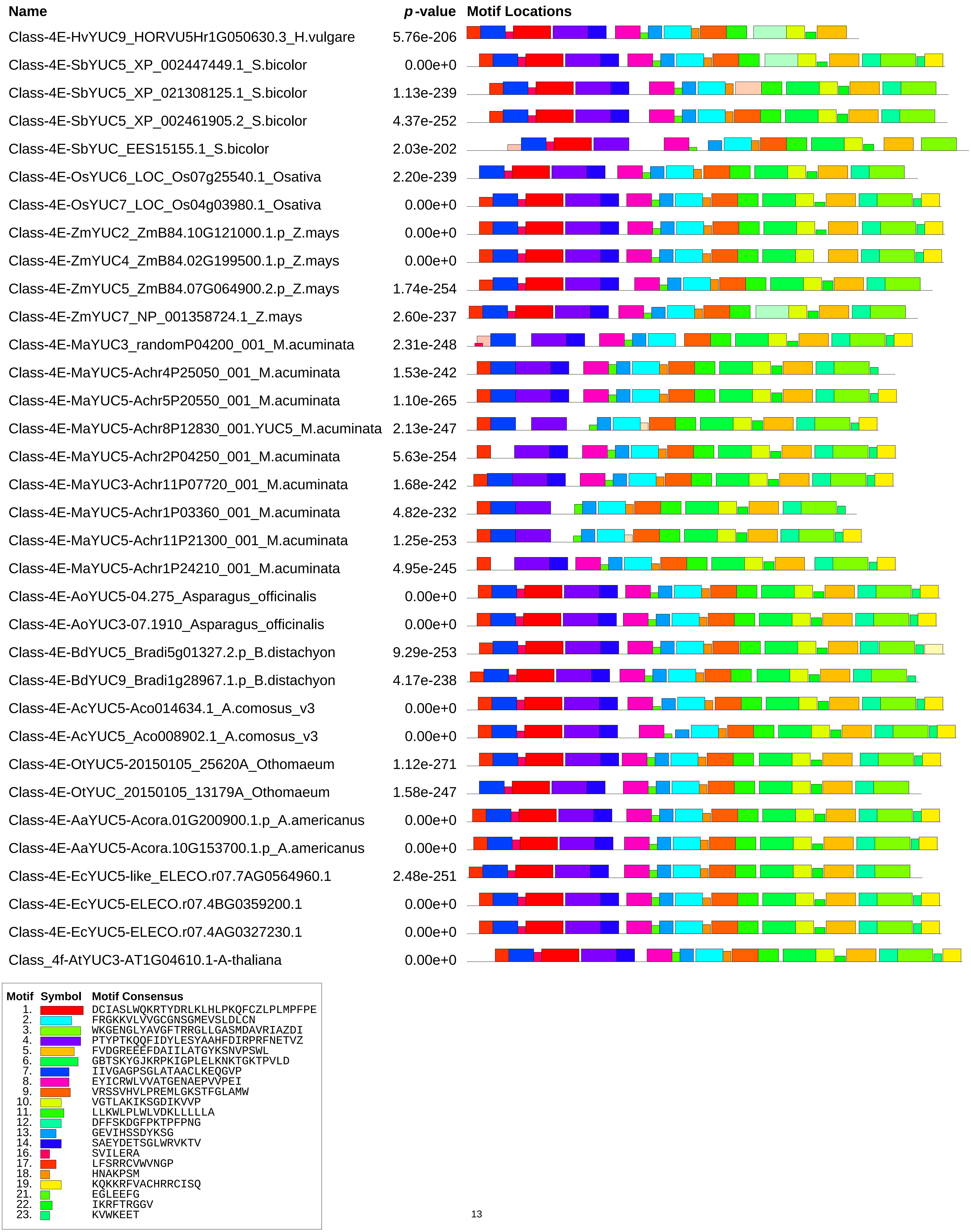
Name
p-value
Motif Locations
Class-4E-HvYUC9_HORVU5Hr1G050630.3_H.vulgare 5.76e-206
Class-4E-SbYUC5_XP_002447449.1_S.bicolor 0.00e+0
Class-4E-SbYUC5_XP_021308125.1_S.bicolor 1.13e-239
Class-4E-SbYUC5_XP_002461905.2_S.bicolor 4.37e-252
Class-4E-SbYUC_EES15155.1_S.bicolor 2.03e-202
Class-4E-OsYUC6_LOC_Os07g25540.1_Osativa 2.20e-239
Class-4E-OsYUC7_LOC_Os04g03980.1_Osativa 0.00e+0
Class-4E-ZmYUC2_ZmB84.10G121000.1.p_Z.mays 0.00e+0
Class-4E-ZmYUC4_ZmB84.02G199500.1.p_Z.mays 0.00e+0
Class-4E-ZmYUC5_ZmB84.07G064900.2.p_Z.mays 1.74e-254
Class-4E-ZmYUC7_NP_001358724.1_Z.mays 2.60e-237
Class-4E-MaYUC3_randomP04200_001_M.acuminata 2.31e-248
Class-4E-MaYUC5-Achr4P25050_001_M.acuminata 1.53e-242
Class-4E-MaYUC5-Achr5P20550_001_M.acuminata 1.10e-265
Class-4E-MaYUC5-Achr8P12830_001.YUC5_M.acuminata 2.13e-247
Class-4E-MaYUC5-Achr2P04250_001_M.acuminata 5.63e-254
Class-4E-MaYUC3-Achr11P07720_001_M.acuminata 1.68e-242
Class-4E-MaYUC5-Achr1P03360_001_M.acuminata 4.82e-232
Class-4E-MaYUC5-Achr11P21300_001_M.acuminata 1.25e-253
Class-4E-MaYUC5-Achr1P24210_001_M.acuminata 4.95e-245
Class-4E-AoYUC5-04.275_Asparagus_officinalis 0.00e+0
Class-4E-AoYUC3-07.1910_Asparagus_officinalis 0.00e+0
Class-4E-BdYUC5_Bradi5g01327.2.p_B.distachyon 9.29e-253
Class-4E-BdYUC9_Bradi1g28967.1.p_B.distachyon 4.17e-238
Class-4E-AcYUC5-Aco014634.1_A.comosus_v3 0.00e+0
Class-4E-AcYUC5_Aco008902.1_A.comosus_v3 0.00e+0
Class-4E-OtYUC5-20150105_25620A_Othomaeum 1.12e-271
Class-4E-OtYUC_20150105_13179A_Othomaeum 1.58e-247
Class-4E-AaYUC5-Acora.01G200900.1.p_A.americanus 0.00e+0
Class-4E-AaYUC5-Acora.10G153700.1.p_A.americanus 0.00e+0
Class-4E-EcYUC5-like_ELECO.r07.7AG0564960.1 2.48e-251
Class-4E-EcYUC5-ELECO.r07.4BG0359200.1 0.00e+0
Class-4E-EcYUC5-ELECO.r07.4AG0327230.1 0.00e+0
Class_4f-AtYUC3-AT1G04610.1-A-thaliana 0.00e+0
Motif Symbol Motif Consensus
1. DCIASLWQKRTYDRLKLHLPKQFCZLPLMPFPE
2. FRGKKVLVVGCGNSGMEVSLDLCN
3. WKGENGLYAVGFTRRGLLGASMDAVRIAZDI
4. PTYPTKQQFIDYLESYAAHFDIRPRFNETVZ
5. FVDGREEEFDAIILATGYKSNVPSWL
6. GBTSKYGJKRPKIGPLELKNKTGKTPVLD
7. IIVGAGPSGLATAACLKEQGVP
8. EYICRWLVVATGENAEPVVPEI
9. VRSSVHVLPREMLGKSTFGLAMW
10. VGTLAKIKSGDIKVVP
11. LLKWLPLWLVDKLLLLLA
12. DFFSKDGFPKTPFPNG
13. GEVIHSSDYKSG
14. SAEYDETSGLWRVKTV
16. SVILERA
17. LFSRRCVWVNGP
18. HNAKPSM
19. KQKKRFVACHRRCISQ
21. EGLEEFG
22. IKRFTRGGV
23. KVWKEET

| Name | *p*-value | Motif Locations |
|---|---|---|
| Class_4f-AtYUC5-AT5G43890.1-A-thaliana | 0.00e+0 | |
| Class_4f-AtYUC7-AT2G33230.1-A-thaliana | 0.00e+0 | |
| Class_4f-AtYUC8-AT4G28720.1-A-thaliana | 0.00e+0 | |
| Class_4f-AtYUC9-AT1G04180.1-A-thaliana | 0.00e+0 | |
| Class_4f-PsYUC5-QTZ21218.1-Pisum-sativum | 4.15e-241 | |
| Class_4f-SlyYUC3-Solyc09g064160.3.1-S.lycopersicum | 0.00e+0 | |
| Class_4f-SlyYUC5-Solyc06g083700.4.1-S.lycopersicum | 0.00e+0 | |
| Class_4f-SlyYUC3-Solyc09g091090.2.1-S.lycopersicum | 0.00e+0 | |
| Class_4f-SlyYUC8-Solyc06g008050.4.1-S.lycopersicum | 0.00e+0 | |
| Class_4f-StYUC3-Soltu.DM.09G018700.1-S.tuberosum | 0.00e+0 | |
| Class_4f-StYUC3-Soltu.DM.09G027910.1-S.tuberosum | 0.00e+0 | |
| Class_4f-StYUC5-Soltu.DM.06G034100.1-S.tuberosum | 0.00e+0 | |
| Class_4f-StYUC3-Soltu.DM.06G002120.1-S.tuberosum | 0.00e+0 | |
| Class_4f-GmYUC1-Glyma.03G169600.1.p-G.max | 0.00e+0 | |
| Class_4f-GmYUC15-Glyma.10G128700.1.p-G.max | 0.00e+0 | |
| Class_4f-GmYUC20-Glyma.19G170800.1.p-G.max | 0.00e+0 | |
| Class_4f-GmYUC22-Glyma.20G080000.1.p-G.max | 0.00e+0 | |
| Class_4f-CsYUC8-Cucsa.348750.1-C.sativus-v1.0 | 0.00e+0 | |
| Class_4f-CsYUC5-Cucsa.140470.1-C.sativus-v1.0 | 0.00e+0 | |
| Class_4f-CsYUC3-Cucsa.255980.1-C.sativus-v1.0 | 2.88e-258 | |
| Class_4f-MtYUC9-Medtr1g069275.1-M.truncatula | 0.00e+0 | |
| Class_4f-MtYUC8-Medtr7g099330.1-M.truncatula | 0.00e+0 | |
| Class_4f-MtYUC3-Medtr1g046230.1-M.truncatula | 0.00e+0 | |
| Class_4f-PpYUC5-Prupe.8G211000.1.p-P.persica | 0.00e+0 | |
| Class_4f-PpYUC3-Prupe.1G054300.1.p-P.persica | 0.00e+0 | |
| Class_4f-BrYUC9-Brara.I05505.1.p-B.rapa | 0.00e+0 | |
| Class_4f-BrYUC5-Brara.B02765.1.p-B.rapa | 0.00e+0 | |
| Class_4f-BrYUC8-Brara.A00847.1.p-B.rapa | 0.00e+0 | |
| Class_4f-BrYUC9-Brara.J00235.1.p-B.rapa | 0.00e+0 | |
| Class_4f-BrYUC3X2-Brara.J00274.1.p-B.rapa | 0.00e+0 | |
| Class_4f-BrYUC8-Brara.H01467.1.p-B.rapa | 0.00e+0 | |
| Class_4f-BrYUC3-Brara.I05483.1.p-B.rapa | 0.00e+0 | |
| Class_4f-BrYUC5-Brara.F03781.1.p-B.rapa | 0.00e+0 | |
| Class_4f-BrYUC7-Brara.E01069.1.p-B.rapa | 0.00e+0 | |

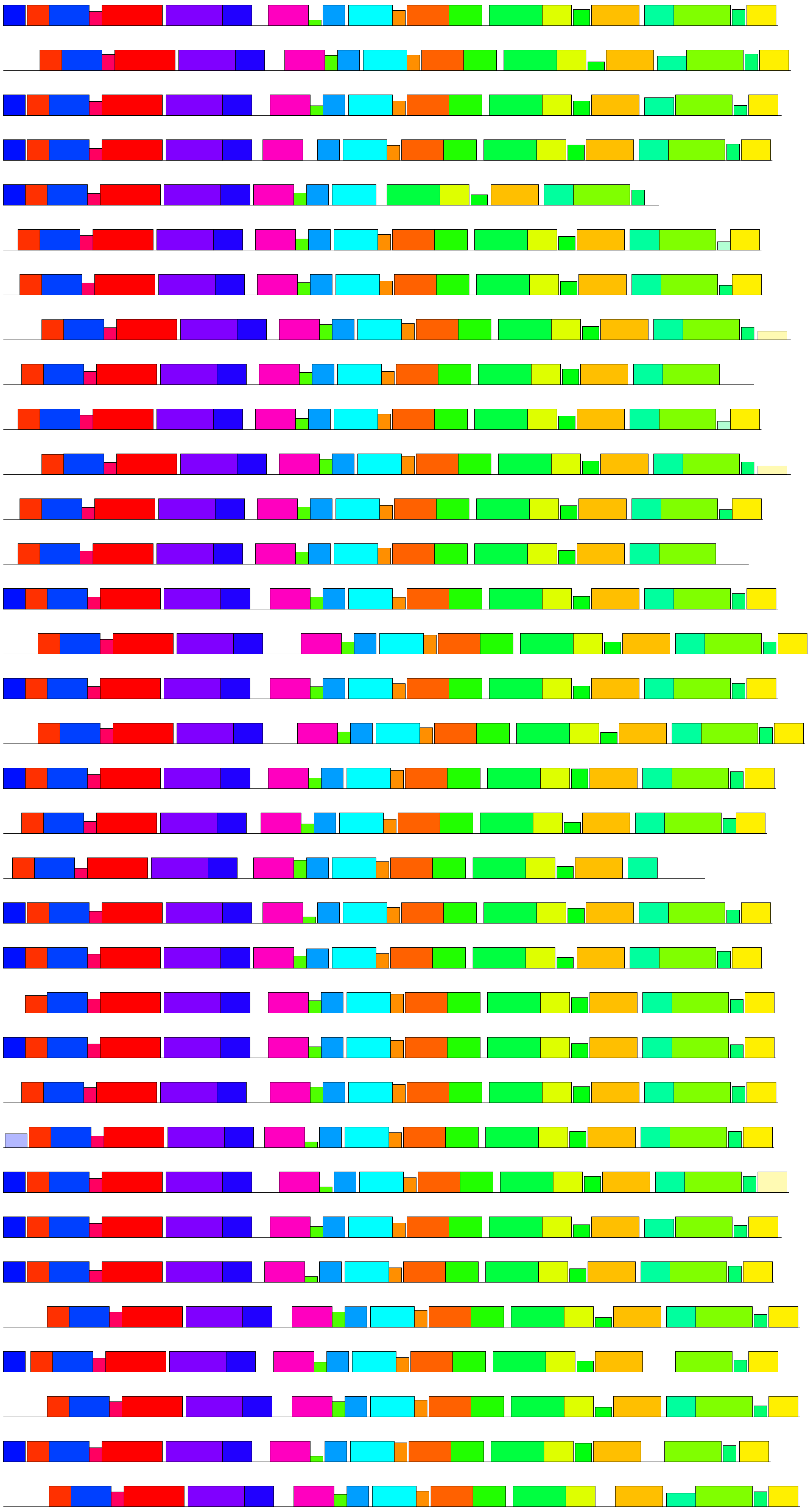

| Motif | Symbol | Motif Consensus |
|---|---|---|
| 1. | | DCIASLWQKRTYDRLKLHLPKQFCZLPLMPFPE |
| 2. | | FRGKKVLVVGCGNSGMEVSLDLCN |
| 3. | | WKGENGLYAVGFTRRGLLGASMDAVRIAZDI |
| 4. | | PTYPTKQQFIDYLESYAAHFDIRPRFNETVZ |
| 5. | | FVDGREEEFDAIILATGYKSNVPSWL |
| 6. | | GBTSKYGJKRPKIGPLELKNKTGKTPVLD |
| 7. | | IIVGAGPSGLATAACLKEQGVP |
| 8. | | EYICRWLVVATGENAEPVVPEI |
| 9. | | VRSSVHVLPREMLGKSTFGLAMW |
| 10. | | VGTLAKIKSGDIKVVP |
| 11. | | LLKWLPLWLVDKLLLLLA |
| 12. | | DFFSKDGFPKTPFPNG |
| 13. | | GEVIHSSDYKSG |
| 14. | | SAEYDETSGLWRVKTV |
| 16. | | SVILERA |
| 17. | | LFSRRCVWVNGP |
| 18. | | HNAKPSM |
| 19. | | KQKKRFVACHRRCISQ |
| 21. | | EGLEEFG |
| 22. | | IKRFTRGGV |
| 23. | | KVWKEET |
| 27. | | MENMFRLVDHED |

| Name | *p*-value |
|---|---|
| Class_4f-PCaYUCF-KAI3990778.1-Papaver-californicum | 0.00e+0 |
| Class_4f-PjYUC2-QIH44904.1-Phtheirospermum | 0.00e+0 |
| Class_4f-PjYUC3-QIH44905.1-Phtheirospermum | 7.27e-260 |
| Class_4f-PjYUC4-QIH44906.1-Phtheirospermum | 0.00e+0 |
| Class_4f-VvYUC8-VIT-207s0005g04800.1-V.vinifera | 0.00e+0 |
| Class_4f-VvYUC3-VIT-205s0051g00060.1-V.vinifera | 3.21e-271 |
| Class_4f-DcYUC3-DCAR-030776-D.carota | 2.08e-260 |
| Class_4f-DcYUC3-DCAR-027361-D.carota | 0.00e+0 |
| Class_4f-HqYUC3-Hyque.10G114700.1.p-H.quercifolia | 0.00e+0 |
| Class_4f-CaYUC3-Scaffold-315.833-C.arabica | 0.00e+0 |
| Class_4f-FvYUC7-AFG16917.1-Fragaria-vesca | 0.00e+0 |
| Class_4f-CpYUC3-supercontig-59.54-C.papaya | 0.00e+0 |
| Class_4f-PtYUC3-Potri.008G174600.1.p-Ptrichocarpa | 0.00e+0 |
| Class_4f-PtYUC7-Potri.010G062400.2.p-Ptrichocarpa | 0.00e+0 |
| Class_4f-PtYUC8-Potri.002G254200.1.p-Ptrichocarpa | 0.00e+0 |
| Class_4f-DcYUC5-DCAR-000550-D.carota | 0.00e+0 |
| Class_4f-DcYUC5-DCAR-012617-D.carota | 0.00e+0 |
| Class_4f-DcYUC5A-DCAR-000561-D.carota | 0.00e+0 |
| Class_4f-DcYUC5B-DCAR-000562-D.carota | 0.00e+0 |
| Class_4f-DcYUC5C-DCAR-000563-D.carota | 0.00e+0 |
| Class_4f-DcYUC5D-DCAR-000560-D.carota | 0.00e+0 |
| Class_4f-CaYUC3-Scaffold-2016.447-C.arabica | 0.00e+0 |
| Class_4f-CaYUC8-Scaffold-635.212-C.arabica | 0.00e+0 |
| Class_4f-CpYUC8-supercontig-87.71-C.papaya | 0.00e+0 |
| Class_4f-FvYUC3-AFG16918.1-Fragaria-vesca | 0.00e+0 |
| Class_4f-HqYUC5-Hyque.04G157000.1.p | 0.00e+0 |
| Class-5a-GmYUCB-TnS000936361t02.p-G.montanum | 7.43e-243 |
| Class-5B-AmTrYUC4A--scaffold00122.41-A.trichopoda | 8.88e-254 |
| Class-5B-AmTrYUC4B--scaffold00002.564-A.trichopoda | 1.48e-263 |
| Class-5B-NcYUC4-Nycol.F00334.1.p-N.colorata-v1.2 | 1.04e-254 |
| Class-5B-NcYUC-Nycol.A02388.1.p-N.colorata-v1.2 | 5.47e-259 |
| Class-5B-HvYUC1X1_XP_044972564.1_H.vulgare | 1.12e-228 |
| Class-5B-HvYUC1X2_XP_044972565.1_H.vulgare | 6.27e-199 |

**Motif Locations**

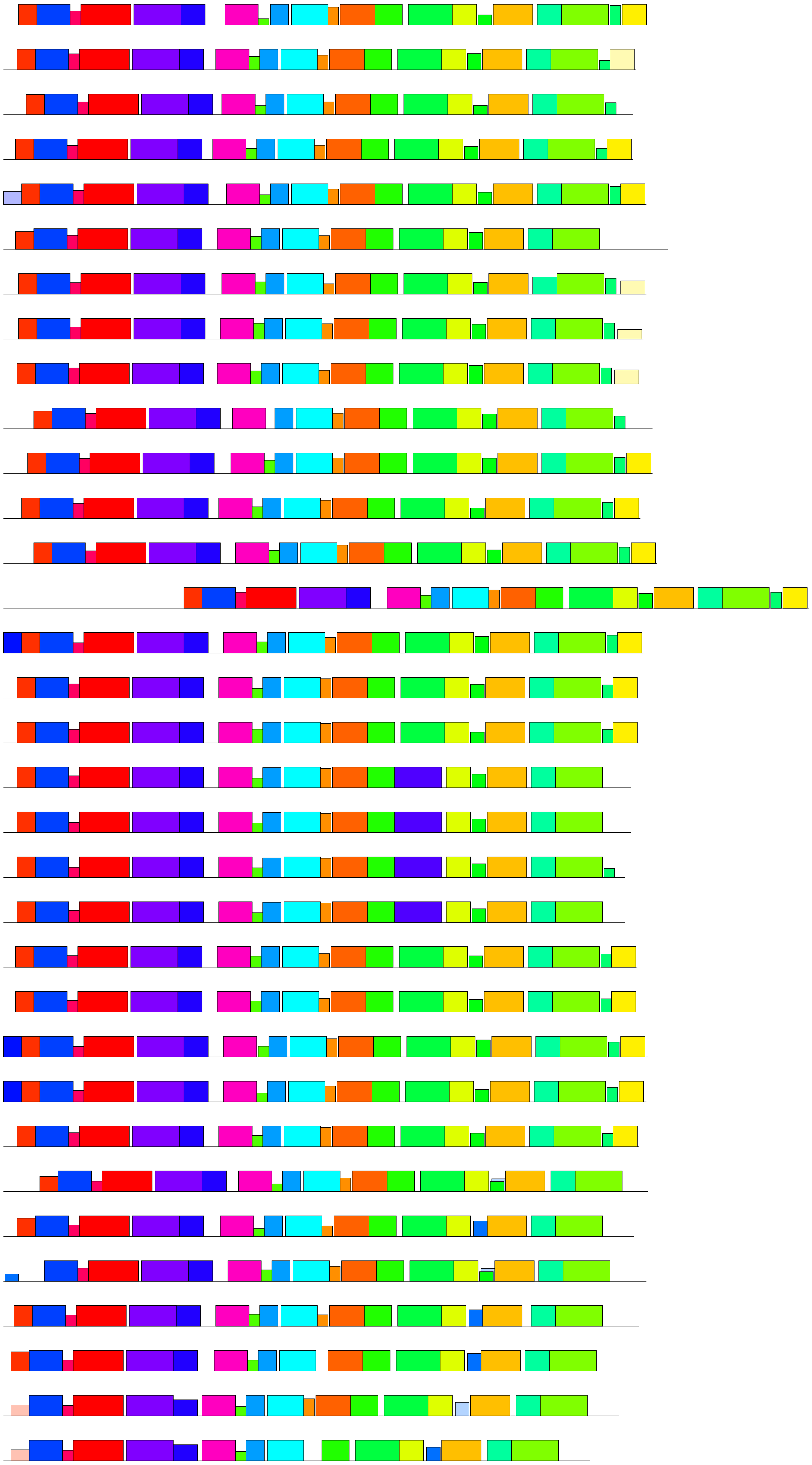

| Motif | Symbol | Motif Consensus |
|---|---|---|
| 1. | | DCIASLWQKRTYDRLKLHLPKQFCZLPLMPFPE |
| 2. | | FRGKKVLVVGCGNSGMEVSLDLCN |
| 3. | | WKGENGLYAVGFTRRGLLGASMDAVRIAZDI |
| 4. | | PTYPTKQQFIDYLESYAAHFDIRPRFNETVZ |
| 5. | | FVDGREEEFDAIILATGYKSNVPSWL |
| 6. | | GBTSKYGJKRPKIGPLELKNKTGKTPVLD |
| 7. | | IIVGAGPSGLATAACLKEQGVP |
| 8. | | EYICRWLVVATGENAEPVVPEI |
| 9. | | VRSSVHVLPREMLGKSTFGLAMW |
| 10. | | VGTLAKIKSGDIKVVP |
| 11. | | LLKWLPLWLVDKLLLLLA |
| 12. | | DFFSKDGFPKTPFPNG |
| 13. | | GEVIHSSDYKSG |
| 14. | | SAEYDETSGLWRVKTV |
| 16. | | SVILERA |
| 17. | | LFSRRCVWVNGP |
| 18. | | HNAKPSM |
| 19. | | KQKKRFVACHRRCISQ |
| 21. | | EGLEEFG |
| 22. | | IKRFTRGGV |
| 23. | | KVWKEET |
| 26. | | ECLTPHGAK |
| 27. | | MENMFRLVDHED |
| 28. | | WFILGDLEKYGIKKPSIGPMELKIKDDGKTP |

| Name | *p*-value |
|---|---|
| Class-5B-SbYUC1_XP_002456044.2_S.bicolor | 1.98e-242 |
| Class-5B-OsYUC1_LOC_Os01g45760_Osativa | 9.75e-247 |
| Class-5B-ZmSPI1-YUCCA1-ZmB84.03G318800.2.p_Z.mays | 1.37e-235 |
| Class-5B-MaYUC2-Achr1P05290_001_M.acuminata | 2.11e-205 |
| Class-5B-AoYUC4_08.665_Asparagus_officinalis | 1.42e-236 |
| Class-5B-BdYUC4_Bradi2g45105.2.p_B.distachyon | 3.70e-230 |
| Class-5B-AaYUC4_Acora.12G097100.1.p_A.americanus | 1.31e-262 |
| Class-5B-EcYUC2-ELECO.r07.1AG0026300.1 | 1.61e-245 |
| Class-5B-EcYUC2-ELECO.r07.1BG0076760.1 | 1.42e-239 |
| Class-5B-MsiYUC4_XP_058112037.1_Magnolia_sinica | 1.17e-264 |
| Class-5b_AtYUC1_AT4G32540.1_A_thaliana | 5.42e-248 |
| Class-5B-AtYUC4A_AT5G11320.1_A_thaliana | 1.51e-260 |
| Class-5B-AtYUC4B_AT5G11320.2_A_thaliana | 8.88e-224 |
| Class-5B-PsYUC1_QTZ21214.1_Pisum_sativum | 8.12e-258 |
| Class-5B-PtYUC1_Potri.006G248200.1.p_Ptrichocarpa | 2.66e-268 |
| Class-5B-PtYUC4_Potri.018G033200.1.p_Ptrichocarpa | 4.56e-267 |

**Motif Locations**

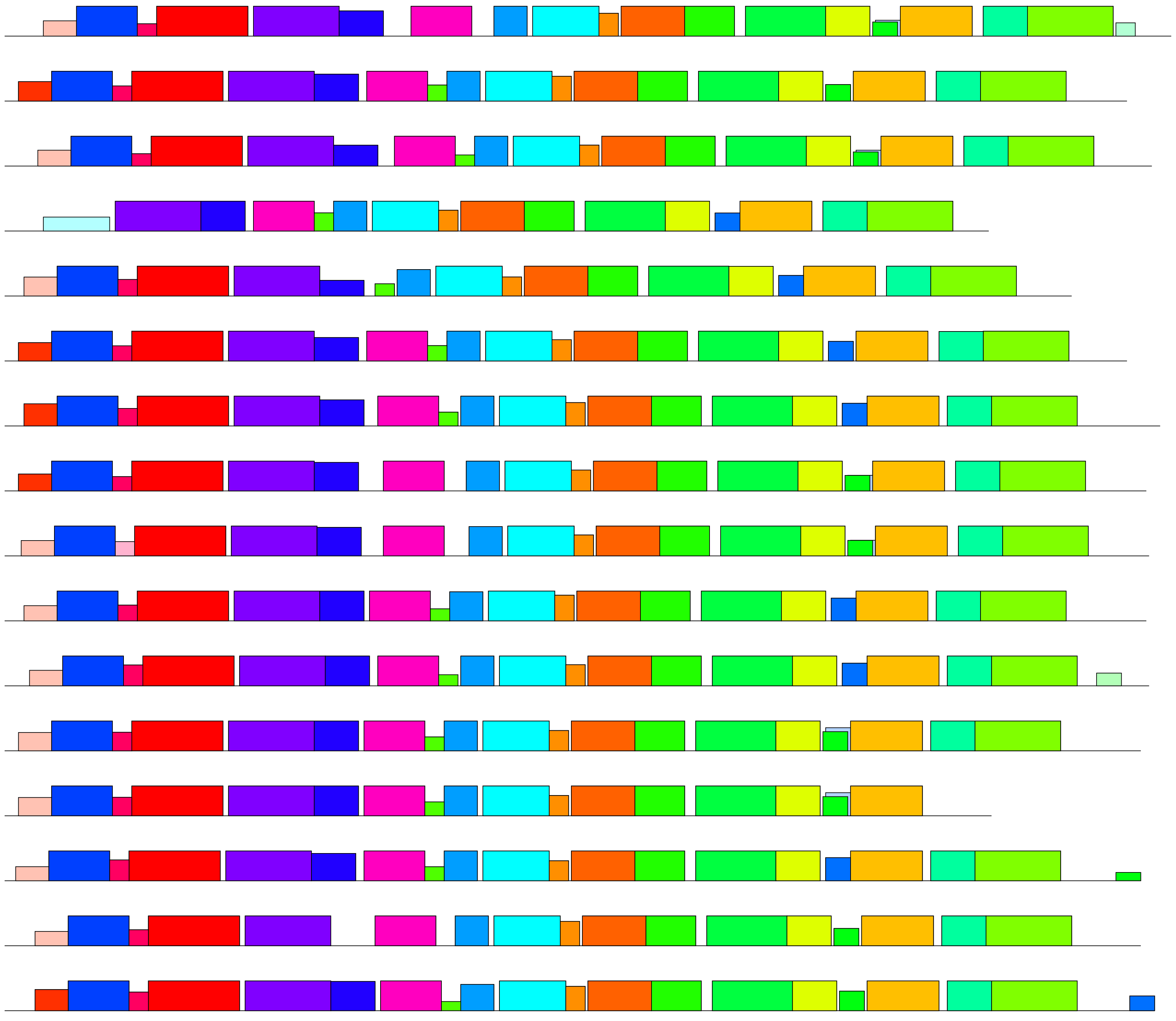

| Motif | Motif Consensus |
|---|---|
| 1. | DCIASLWQKRTYDRLKLHLPKQFCZLPLMPFPE |
| 2. | FRGKKVLVVGCGNSGMEVSLDLCN |
| 3. | WKGENGLYAVGFTRRGLLGASMDAVRIAZDI |
| 4. | PTYPTKQQFIDYLESYAAHFDIRPRFNETVZ |
| 5. | FVDGREEEFDAIILATGYKSNVPSWL |
| 6. | GBTSKYGJKRPKIGPLELKNKTGKTPVLD |
| 7. | IIVGAGPSGLATAACLKEQGVP |
| 8. | EYICRWLVVATGENAEPVVPEI |
| 9. | VRSSVHVLPREMLGKSTFGLAMW |
| 10. | VGTLAKIKSGDIKVVP |
| 11. | LLKWLPLWLVDKLLLLLA |
| 12. | DFFSKDGFPKTPFPNG |
| 13. | GEVIHSSDYKSG |
| 14. | SAEYDETSGLWRVKTV |
| 16. | SVILERA |
| 17. | LFSRRCVWVNGP |
| 18. | HNAKPSM |
| 21. | EGLEEFG |
| 22. | IKRFTRGGV |
| 23. | KVWKEET |
| 26. | ECLTPHGAK |

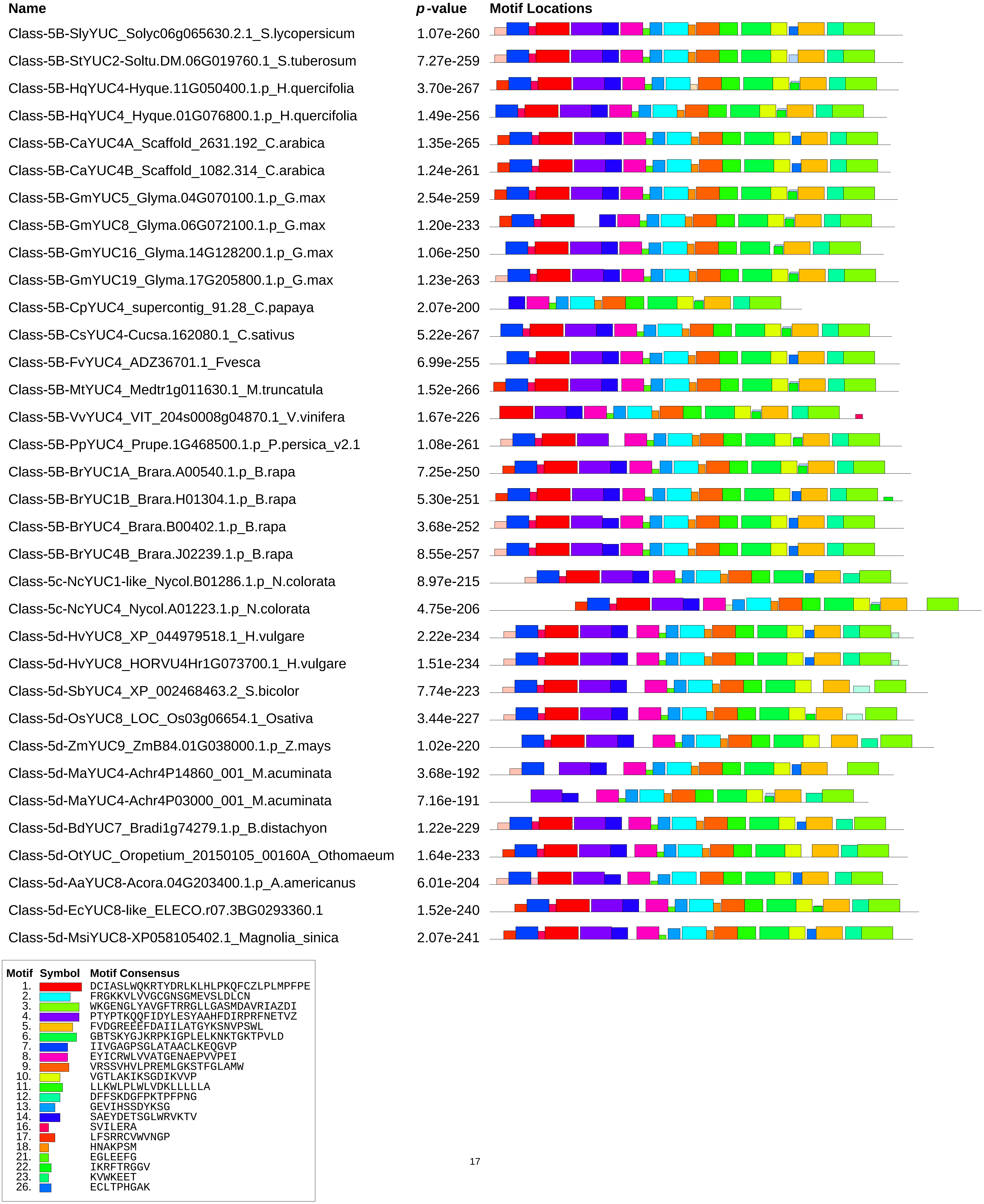
Name
p-value
Motif Locations
Class-5B-SlyYUC_Solyc06g065630.2.1_S.lycopersicum 1.07e-260
Class-5B-StYUC2-Soltu.DM.06G019760.1_S.tuberosum 7.27e-259
Class-5B-HqYUC4-Hyque.11G050400.1.p_H.quercifolia 3.70e-267
Class-5B-HqYUC4_Hyque.01G076800.1.p_H.quercifolia 1.49e-256
Class-5B-CaYUC4A_Scaffold_2631.192_C.arabica 1.35e-265
Class-5B-CaYUC4B_Scaffold_1082.314_C.arabica 1.24e-261
Class-5B-GmYUC5_Glyma.04G070100.1.p_G.max 2.54e-259
Class-5B-GmYUC8_Glyma.06G072100.1.p_G.max 1.20e-233
Class-5B-GmYUC16_Glyma.14G128200.1.p_G.max 1.06e-250
Class-5B-GmYUC19_Glyma.17G205800.1.p_G.max 1.23e-263
Class-5B-CpYUC4_supercontig_91.28_C.papaya 2.07e-200
Class-5B-CsYUC4-Cucsa.162080.1_C.sativus 5.22e-267
Class-5B-FvYUC4_ADZ36701.1_Fvesca 6.99e-255
Class-5B-MtYUC4_Medtr1g011630.1_M.truncatula 1.52e-266
Class-5B-VvYUC4_VIT_204s0008g04870.1_V.vinifera 1.67e-226
Class-5B-PpYUC4_Prupe.1G468500.1.p_P.persica_v2.1 1.08e-261
Class-5B-BrYUC1A_Brara.A00540.1.p_B.rapa 7.25e-250
Class-5B-BrYUC1B_Brara.H01304.1.p_B.rapa 5.30e-251
Class-5B-BrYUC4_Brara.B00402.1.p_B.rapa 3.68e-252
Class-5B-BrYUC4B_Brara.J02239.1.p_B.rapa 8.55e-257
Class-5c-NcYUC1-like_Nycol.B01286.1.p_N.colorata 8.97e-215
Class-5c-NcYUC4_Nycol.A01223.1.p_N.colorata 4.75e-206
Class-5d-HvYUC8_XP_044979518.1_H.vulgare 2.22e-234
Class-5d-HvYUC8_HORVU4Hr1G073700.1_H.vulgare 1.51e-234
Class-5d-SbYUC4_XP_002468463.2_S.bicolor 7.74e-223
Class-5d-OsYUC8_LOC_Os03g06654.1_Osativa 3.44e-227
Class-5d-ZmYUC9_ZmB84.01G038000.1.p_Z.mays 1.02e-220
Class-5d-MaYUC4-Achr4P14860_001_M.acuminata 3.68e-192
Class-5d-MaYUC4-Achr4P03000_001_M.acuminata 7.16e-191
Class-5d-BdYUC7_Bradi1g74279.1.p_B.distachyon 1.22e-229
Class-5d-OtYUC_Oropetium_20150105_00160A_Othomaeum 1.64e-233
Class-5d-AaYUC8-Acora.04G203400.1.p_A.americanus 6.01e-204
Class-5d-EcYUC8-like_ELECO.r07.3BG0293360.1 1.52e-240
Class-5d-MsiYUC8-XP058105402.1_Magnolia_sinica 2.07e-241
Motif Symbol Motif Consensus
1. DCIASLWQKRTYDRLKLHLPKQFCZLPLMPFPE
2. FRGKKVLVVGCGNSGMEVSLDLCN
3. WKGENGLYAVGFTRRGLLGASMDAVRIAZDI
4. PTYPTKQQFIDYLESYAAHFDIRPRFNETVZ
5. FVDGREEEFDAIILATGYKSNVPSWL
6. GBTSKYGJKRPKIGPLELKNKTGKTPVLD
7. IIVGAGPSGLATAACLKEQGVP
8. EYICRWLVVATGENAEPVVPEI
9. VRSSVHVLPREMLGKSTFGLAMW
10. VGTLAKIKSGDIKVVP
11. LLKWLPLWLVDKLLLLLA
12. DFFSKDGFPKTPFPNG
13. GEVIHSSDYKSG
14. SAEYDETSGLWRVKTV
16. SVILERA
17. LFSRRCVWVNGP
18. HNAKPSM
21. EGLEEFG
22. IKRFTRGGV
23. KVWKEET
26. ECLTPHGAK
