## Supplementary Data (Data S1-S10) for "The auxin gatekeepers: Evolution and diversification of the YUCCA family": Data S7.pdf

AtYUCs amino acid alignment (* - sites under negative selection, * - sites under positive selection)

Analysis done in Datamonkey MEME. (URL: https://www.datamonkey.org/meme)

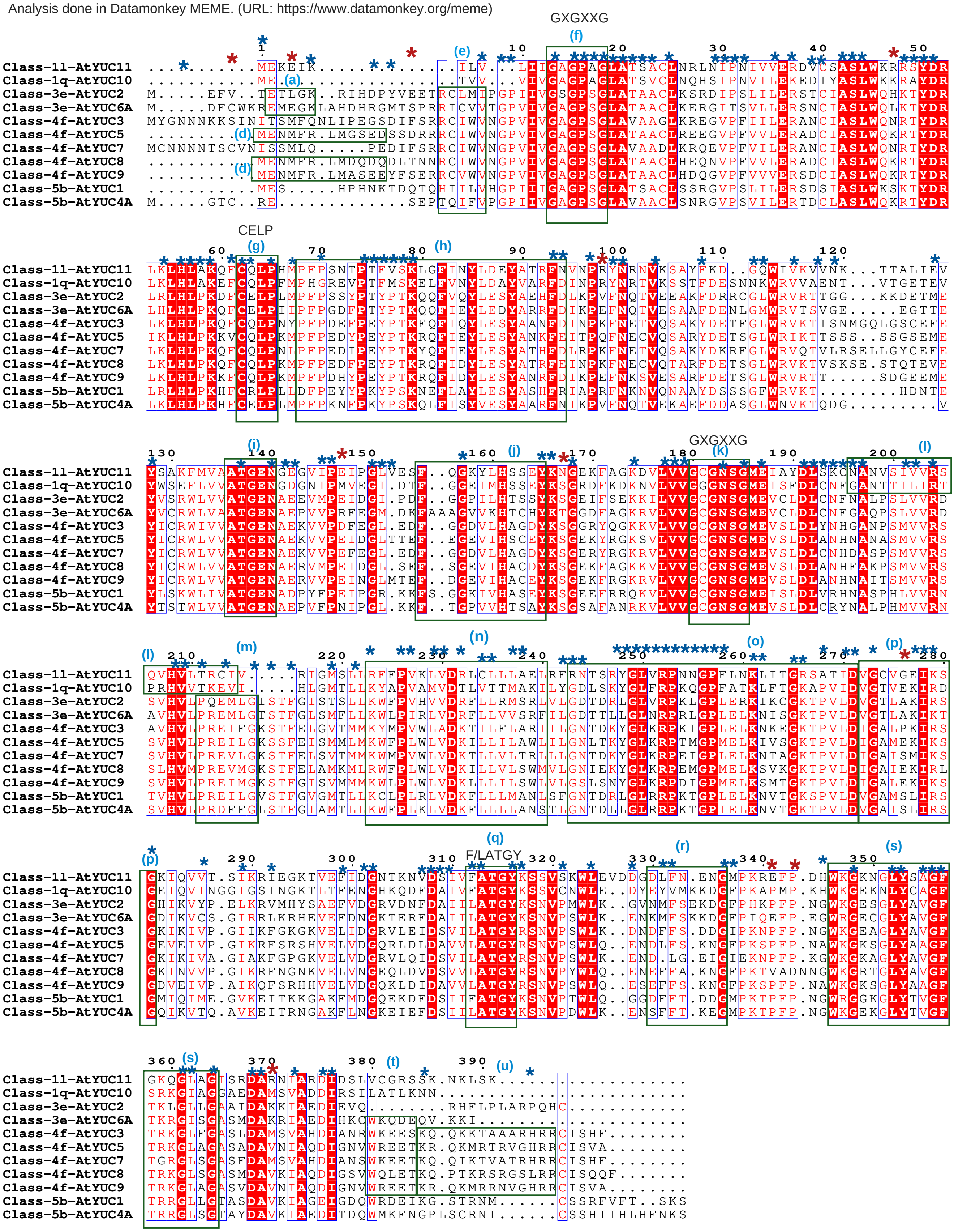


- Amino acid residues specific to codons were obtained from the tree generated in MEME and marked in the alignment.
- None of the diversifying sites are predicted in the signature motifs, which are common to all class B FMOs (motif f,h, j, k, q).

**(All the motifs in the alignment are marked based on the motif key)**

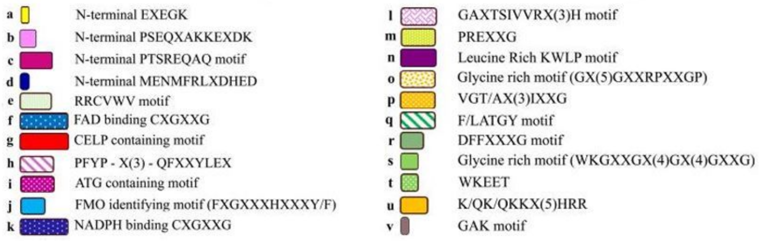


Note: Motifs are predicted in MEME suite (https://meme-suite.org/meme/) using all 469 studied sequence, and common motifs are marked

### AtYUCs codon alignment (with amino acids)

(PAL2NAL program: https://www.bork.embl.de/pal2nal/)

| | 1 | 2 | 3 | 4 | 5 | 6 | 7 | 8 | 9 | 10 | 11 | 12 | 13 | 14 | 15 | 16 | 17 | 18 | 19 | 20 |
|---|---|---|---|---|---|---|---|---|---|---|---|---|---|---|---|---|---|---|---|---|
| | | | | * | | | | | * | | | * | | | * | | * | | | |
| | - | - | - | - | - | - | - | - | - | - | - | M | E | K | E | I | K | - | - | - |
| Class-1l-AtYUC11 | --- | --- | --- | --- | --- | --- | --- | --- | --- | --- | --- | ATG | GAG | AAA | GAA | ATC | AAG | --- | --- | --- |
| | - | - | - | - | - | - | - | - | - | - | - | M | E | - | - | - | - | - | - | - |
| Class-1q-AtYUC10 | --- | --- | --- | --- | --- | --- | --- | --- | --- | --- | --- | ATG | GAG | --- | --- (a) | --- | --- | --- | --- | --- |
| | M | - | - | - | - | - | E | F | V | - | - | T | E | T | L | G | K | - | - | R |
| Class-3e-AtYUC2 | ATG | --- | --- | --- | --- | --- | GAG | TTT | GTT | --- | --- | ACA | GAA | ACG | TTA | GGC | AAG | --- | --- | AGA |
| | M | - | - | - | - | - | D | F | C | W | K | R | E | M | E | G | K | L | A | H |
| Class-3e-AtYUC6A | ATG | --- | --- | --- | --- | --- | GAT | TTC | TGT | TGG | AAG | AGA | GAG | ATG | GAA | GGT | AAA | CTA | GCA | CAT |
| | M | Y | G | N | N | N | K | K | S | I | N | I | T | S | M | F | Q | N | L | I |
| Class-4f-AtYUC3 | ATG | TAT | GGA | AAT | AAT | AAC | AAG | AAG | TCT | ATC | AAC | ATC | ACG | AGT | ATG | TTT | CAA | AAC | CTC | ATC |
| | - | - | - | - | - | - | - | - | - | - | - | M | E | N | M | F | R | - | L | M |
| Class-4f-AtYUC5 (d) | --- | --- | --- | --- | --- | --- | --- | --- | --- | --- | --- | ATG | GAG | AAC | ATG | TTT | AGG | --- | CTC | ATG |
| | M | C | N | N | N | N | T | S | C | V | N | I | S | S | M | L | Q | - | - | - |
| Class-4f-AtYUC7 | ATG | TGT | AAT | AAC | AAT | AAC | ACA | AGT | TGT | GTC | AAC | ATC | TCA | AGT | ATG | CTT | CAG | --- | --- | --- |
| | - | - | - | - | - | - | - | - | - | - | - | M | E | N | M | F | R | - | L | M |
| Class-4f-AtYUC8 | --- | --- | --- | --- | --- | --- | --- | --- | --- | --- | --- | ATG | GAG | AAT | ATG | TTT | CGT | --- | TTG | ATG |
| | - | - | - | - | - | - | - | - | - | - | - | M | E | N | M | F | R | - | L | M |
| Class-4f-AtYUC9 (d) | --- | --- | --- | --- | --- | --- | --- | --- | --- | --- | --- | ATG | GAG | AAT | ATG | TTT | AGA | --- | CTC | ATG |
| | - | - | - | - | - | - | - | - | - | - | - | M | E | S | - | - | - | - | - | H |
| Class-5b-AtYUC1 | --- | --- | --- | --- | --- | --- | --- | --- | --- | --- | --- | ATG | GAG | TCT | --- | --- | --- | --- | --- | CAT |
| | M | - | - | - | - | - | G | T | C | - | - | R | E | - | - | - | - | - | - | - |
| Class-5b-AtYUC4A | ATG | --- | --- | --- | --- | --- | GGC | ACT | TGT | --- | --- | AGA | GAA | --- | --- | --- | --- | --- | --- | --- |

| | 21 | 22 | 23 | 24 | 25 | 26 | 27 | 28 | 29 | 30 | 31 | 32 | 33 | 34 | 35 | 36 | 37 | 38 | 39 | 40 |
|---|---|---|---|---|---|---|---|---|---|---|---|---|---|---|---|---|---|---|---|---|
| | | | * | | * | | * | | | | (e) | | | * | | * | * | | | |
| | - | - | - | - | - | - | - | - | - | - | - | I | L | V | - | - | - | L | I | I |
| Class-1l-AtYUC11 | --- | --- | --- | --- | --- | --- | --- | --- | --- | --- | --- | ATT | CTG | GTT | --- | --- | --- | TTG | ATT | ATC |
| | - | - | - | - | - | - | - | - | - | - | - | T | V | V | - | - | - | V | I | V |
| Class-1q-AtYUC10 | --- | --- | --- | --- | --- | --- | --- | --- | --- | --- | --- | ACC | GTA | GTG | --- | --- | --- | GTG | ATT | GTG |
| | I | H | D | P | Y | V | E | E | T | R | C | L | M | I | P | G | P | I | I | V |
| Class-3e-AtYUC2 | ATC | CAT | GAT | CCG | TAC | GTG | GAG | GAA | ACT | AGG | TGC | TTA | ATG | ATT | CCC | GGA | CCA | ATC | ATT | GTC |
| | D | H | R | G | M | T | S | P | R | R | I | C | V | V | T | G | P | V | I | V |
| Class-3e-AtYUC6A | GAC | CAC | CGC | GGC | ATG | ACG | TCA | CCG | CGT | CGT | ATC | TGC | GTC | GTC | ACC | GGT | CCG | GTG | ATC | GTA |
| | P | E | G | S | D | I | F | S | R | R | C | I | W | V | N | G | P | V | I | V |
| Class-4f-AtYUC3 | CCC | GAA | GGC | AGC | GAC | ATT | TTC | TCC | CGG | CGT | TGC | ATC | TGG | GTC | AAT | GGA | CCC | GTC | ATC | GTC |
| | G | S | E | D | S | S | D | R | R | R | C | I | W | V | N | G | P | V | I | V |
| Class-4f-AtYUC5 (d) | GGA | AGT | GAA | GAT | TCC | TCC | GAC | AGA | AGA | CGG | TGC | ATT | TGG | GTT | AAC | GGT | CCT | GTA | ATC | GTC |
| | - | - | P | E | D | I | F | S | R | R | C | I | W | V | N | G | P | V | I | V |
| Class-4f-AtYUC7 | --- | --- | CCT | GAG | GAC | ATT | TTC | TCC | CGT | CGT | TGC | ATC | TGG | GTG | AAT | GGG | CCT | GTC | ATC | GTC |
| | D | Q | D | Q | D | L | T | N | N | R | C | I | W | V | N | G | P | V | I | V |
| Class-4f-AtYUC8 (d) | GAT | CAA | GAT | CAG | GAT | TTA | ACT | AAT | AAC | CGG | TGC | ATT | TGG | GTC | AAC | GGA | CCG | GTC | ATC | GTC |
| | A | S | E | E | Y | F | S | E | R | R | C | V | W | V | N | G | P | V | I | V |
| Class-4f-AtYUC9 | GCA | AGT | GAA | GAA | TAT | TTC | TCA | GAG | CGG | CGA | TGT | GTT | TGG | GTC | AAC | GGT | CCG | GTT | ATC | GTA |
| | P | H | N | K | T | D | Q | T | Q | H | I | I | L | V | H | G | P | I | I | I |
| Class-5b-AtYUC1 | CCT | CAC | AAC | AAA | ACT | GAC | CAG | ACC | CAG | CAT | ATC | ATC | CTC | GTA | CAC | GGT | CCC | ATC | ATC | ATC |
| | - | - | - | - | - | - | S | E | P | T | Q | I | F | V | P | G | P | I | I | V |
| Class-5b-AtYUC4A | --- | --- | --- | --- | --- | --- | TCA | GAA | CCT | ACT | CAA | ATC | TTC | GTT | CCT | GGT | CCG | ATC | ATC | GTC |

| | 41 | 42 | 43 | 44 | 45 | 46 | 47 | 48 | 49 | 50 | 51 | 52 | 53 | 54 | 55 | 56 | 57 | 58 | 59 | 60 |
|---|---|---|---|---|---|---|---|---|---|---|---|---|---|---|---|---|---|---|---|---|
| | * | | * | (f) * | * | * | * | * | * | * | * | | * | | | | * | * | * | * |
| | G | A | G | P | A | G | L | A | T | S | A | C | L | N | R | L | N | I | P | N |
| Class-1l-AtYUC11 | GGT | GCA | GGA | CCA | GCC | GGT | TTA | GCA | ACA | TCA | GCT | TGT | CTT | AAC | CGG | TTG | AAC | ATA | CCA | AAC |
| | G | A | G | P | A | G | L | A | T | S | V | C | L | N | Q | H | S | I | P | N |
| Class-1q-AtYUC10 | GGA | GCT | GGA | CCG | GCC | GGT | TTA | GCA | ACA | TCG | GTT | TGT | CTA | AAC | CAA | CAC | TCA | ATC | CCA | AAC |
| | G | S | G | P | S | G | L | A | T | A | A | C | L | K | S | R | D | I | P | S |
| Class-3e-AtYUC2 | GGT | TCC | GGG | CCG | TCG | GGA | CTG | GCC | ACA | GCG | GCA | TGT | TTA | AAG | TCG | AGA | GAC | ATC | CCT | AGT |
| | G | A | G | P | S | G | L | A | T | A | A | C | L | K | E | R | G | I | T | S |
| Class-3e-AtYUC6A | GGC | GCC | GGA | CCG | TCG | GGA | CTA | GCC | ACG | GCA | GCA | TGT | TTA | AAA | GAG | AGA | GGT | ATC | ACG | TCC |
| | G | A | G | P | S | G | L | A | V | A | A | G | L | K | R | E | G | V | P | F |
| Class-4f-AtYUC3 | GGA | GCT | GGC | CCA | TCA | GGT | CTA | GCC | GTT | GCT | GCG | GGC | TTG | AAA | CGT | GAA | GGG | GTA | CCA | TTC |
| | G | A | G | P | S | G | L | A | T | A | A | C | L | R | E | E | G | V | P | F |
| Class-4f-AtYUC5 | GGA | GCC | GGG | CCA | TCG | GGG | CTA | GCC | ACG | GCA | GCC | TGC | CTC | CGC | GAG | GAA | GGT | GTT | CCC | TTT |
| | G | A | G | P | S | G | L | A | V | A | A | D | L | K | R | Q | E | V | P | F |
| Class-4f-AtYUC7 | GGT | GCT | GGC | CCA | TCA | GGC | CTA | GCT | GTC | GCC | GCC | GAC | CTG | AAA | CGC | CAA | GAA | GTT | CCG | TTT |
| | G | A | G | P | S | G | L | A | T | A | A | C | L | H | E | Q | N | V | P | F |
| Class-4f-AtYUC8 | GGA | GCT | GGA | CCG | TCG | GGG | TTA | GCG | ACG | GCG | GCT | TGT | CTC | CAT | GAA | CAA | AAC | GTT | CCT | TTC |
| | G | A | G | P | S | G | L | A | T | A | A | C | L | H | D | Q | G | V | P | F |
| Class-4f-AtYUC9 | GGA | GCT | GGT | CCA | TCC | GGT | TTA | GCT | ACA | GCA | GCT | TGT | CTA | CAT | GAT | CAA | GGA | GTC | CCA | TTC |
| | G | A | G | P | S | G | L | A | T | S | A | C | L | S | S | R | G | V | P | S |
| Class-5b-AtYUC1 | GGA | GCT | GGC | CCT | TCT | GGT | CTT | GCC | ACT | TCA | GCA | TGT | CTC | TCG | AGC | CGT | GGA | GTC | CCT | TCT |
| | G | A | G | P | S | G | L | A | V | A | A | C | L | S | N | R | G | V | P | S |
| Class-5b-AtYUC4A | GGA | GCC | GGG | CCT | TCC | GGT | CTA | GCC | GTA | GCG | GCT | TGT | TTG | TCA | AAC | CGA | GGC | GTA | CCA | TCG |

Modified PAL2NAL output. Each codon is numbered, and all motifs are designated according to the motif keys.

| | 61 | 62 | 63 | 64 | 65 | 66 | 67 | 68 | 69 | 70 | 71 | 72 | 73 | 74 | 75 | 76 | 77 | 78 | 79 | 80 |
|---|---|---|---|---|---|---|---|---|---|---|---|---|---|---|---|---|---|---|---|---|
| | | * | | * | | | * | * | | * | * | * | | | * | * | * | * | * | |
| | I | V | V | E | R | D | V | C | S | A | S | L | W | K | R | R | S | Y | D | R |
| Class-1l-AtYUC11 | ATA | GTA | GTG | GAA | AGA | GAT | GTT | TGC | AGT | GCC | TCT | CTG | TGG | AAA | AGA | AGG | TCT | TAC | GAT | CGT |
| | V | I | L | E | K | E | D | I | Y | A | S | L | W | K | K | R | A | Y | D | R |
| Class-1q-AtYUC10 | GTG | ATC | CTC | GAG | AAA | GAA | GAC | ATC | TAT | GCA | TCC | CTT | TGG | AAG | AAA | CGA | GCG | TAC | GAT | CGT |
| | L | I | L | E | R | S | T | C | I | A | S | L | W | Q | H | K | T | Y | D | R |
| Class-3e-AtYUC2 | TTG | ATT | CTA | GAA | CGT | TCC | ACT | TGC | ATA | GCG | TCA | CTA | TGG | CAG | CAC | AAA | ACA | TAT | GAT | CGT |
| | V | L | L | E | R | S | N | C | I | A | S | L | W | Q | L | K | T | Y | D | R |
| Class-3e-AtYUC6A | GTA | CTA | CTA | GAG | AGA | TCA | AAC | TGT | ATA | GCA | TCA | CTA | TGG | CAG | CTC | AAG | ACT | TAT | GAC | CGT |
| | I | I | L | E | R | A | N | C | I | A | S | L | W | Q | N | R | T | Y | D | R |
| Class-4f-AtYUC3 | ATA | ATC | CTC | GAG | CGA | GCT | AAC | TGC | ATT | GCT | TCA | TTA | TGG | CAA | AAC | AGA | ACC | TAC | GAT | CGT |
| | V | V | L | E | R | A | D | C | I | A | S | L | W | Q | K | R | T | Y | D | R |
| Class-4f-AtYUC5 | GTG | GTA | TTG | GAG | AGA | GCA | GAT | TGC | ATA | GCT | TCA | CTA | TGG | CAA | AAA | CGA | ACT | TAC | GAT | AGA |
| | V | I | L | E | R | A | N | C | I | A | S | L | W | Q | N | R | T | Y | D | R |
| Class-4f-AtYUC7 | GTA | ATC | CTC | GAG | CGA | GCT | AAC | TGC | ATA | GCT | TCT | CTG | TGG | CAA | AAC | AGA | ACC | TAT | GAT | CGT |
| | V | V | L | E | R | A | D | C | I | A | S | L | W | Q | K | R | T | Y | D | R |
| Class-4f-AtYUC8 | GTT | GTT | CTC | GAG | AGA | GCA | GAT | TGT | ATT | GCT | TCT | CTA | TGG | CAA | AAA | CGT | ACT | TAC | GAT | CGT |
| | V | V | V | E | R | S | D | C | I | A | S | L | W | Q | K | R | T | Y | D | R |
| Class-4f-AtYUC9 | GTT | GTG | GTC | GAG | AGA | TCA | GAT | TGC | ATA | GCG | TCA | CTG | TGG | CAA | AAA | CGA | ACC | TAC | GAT | AGG |
| | L | I | L | E | R | S | D | S | I | A | S | L | W | K | S | K | T | Y | D | R |
| Class-5b-AtYUC1 | TTG | ATC | CTA | GAA | CGG | TCG | GAT | TCA | ATA | GCA | TCT | CTA | TGG | AAA | TCT | AAA | ACC | TAC | GAC | CGA |
| | V | I | L | E | R | T | D | C | L | A | S | L | W | Q | K | R | T | Y | D | R |
| Class-5b-AtYUC4A | GTT | ATC | CTC | GAG | AGA | ACC | GAT | TGT | TTA | GCT | TCT | TTA | TGG | CAA | AAA | AGA | ACC | TAC | GAC | CGT |

(g) CELP (codons 90–93)

(h) (codons 96–100)

| | 81 | 82 | 83 | 84 | 85 | 86 | 87 | 88 | 89 | 90 | 91 | 92 | 93 | 94 | 95 | 96 | 97 | 98 | 99 | 100 |
|---|---|---|---|---|---|---|---|---|---|---|---|---|---|---|---|---|---|---|---|---|
| | | * | | * | * | * | * | | * | * | * | | * | | * | | | * | | |
| | L | K | L | H | L | A | K | Q | F | C | Q | L | P | H | M | P | F | P | S | N |
| Class-1l-AtYUC11 | CTC | AAG | CTC | CAC | CTC | GCA | AAG | CAA | TTC | TGT | CAA | CTC | CCT | CAC | ATG | CCA | TTC | CCC | TCA | AAC |
| | L | K | L | H | L | A | K | E | F | C | Q | L | P | F | M | P | H | G | R | E |
| Class-1q-AtYUC10 | CTC | AAA | CTT | CAC | TTA | GCC | AAA | GAA | TTT | TGC | CAG | CTT | CCT | TTC | ATG | CCA | CAC | GGT | CGT | GAG |
| | L | R | L | H | L | P | K | D | F | C | E | L | P | L | M | P | F | P | S | S |
| Class-3e-AtYUC2 | CTT | CGA | CTT | CAT | CTC | CCT | AAA | GAT | TTC | TGT | GAG | CTT | CCA | TTG | ATG | CCT | TTT | CCT | TCA | AGC |
| | L | H | L | H | L | P | K | Q | F | C | E | L | P | I | I | P | F | P | G | D |
| Class-3e-AtYUC6A | CTT | CAT | CTT | CAC | CTT | CCT | AAA | CAA | TTC | TGT | GAA | CTT | CCG | ATT | ATA | CCC | TTC | CCC | GGA | GAT |
| | L | K | L | H | L | P | K | Q | F | C | Q | L | P | N | Y | P | F | P | D | E |
| Class-4f-AtYUC3 | CTC | AAA | CTC | CAT | CTA | CCT | AAA | CAG | TTT | TGC | CAA | TTA | CCC | AAT | TAT | CCT | TTC | CCG | GAC | GAG |
| | I | K | L | H | L | P | K | K | V | C | Q | L | P | K | M | P | F | P | E | D |
| Class-4f-AtYUC5 | ATC | AAA | CTT | CAT | TTA | CCA | AAG | AAA | GTT | TGT | CAA | TTA | CCC | AAA | ATG | CCC | TTC | CCG | GAA | GAC |
| | L | K | L | H | L | P | K | Q | F | C | Q | L | P | N | L | P | F | P | E | D |
| Class-4f-AtYUC7 | CTC | AAG | CTT | CAT | CTC | CCA | AAG | CAA | TTC | TGT | CAA | TTA | CCC | AAT | TTA | CCC | TTC | CCC | GAG | GAT |
| | L | K | L | H | L | P | K | Q | F | C | Q | L | P | K | M | P | F | P | E | D |
| Class-4f-AtYUC8 | CTC | AAG | CTT | CAC | CTT | CCT | AAG | CAG | TTT | TGT | CAA | TTA | CCG | AAA | ATG | CCC | TTC | CCT | GAG | GAC |
| | L | K | L | H | L | P | K | K | F | C | Q | L | P | K | M | P | F | P | D | H |
| Class-4f-AtYUC9 | CTC | AAG | CTT | CAC | TTG | CCC | AAG | AAA | TTC | TGC | CAA | TTA | CCG | AAA | ATG | CCC | TTC | CCG | GAT | CAC |
| | L | R | L | H | L | P | K | H | F | C | R | L | P | L | L | D | F | P | E | Y |
| Class-5b-AtYUC1 | CTC | AGA | CTC | CAT | CTC | CCA | AAA | CAC | TTT | TGC | CGG | TTA | CCC | CTC | CTG | GAC | TTC | CCT | GAA | TAT |
| | L | K | L | H | L | P | K | H | F | C | E | L | P | L | M | P | F | P | K | N |
| Class-5b-AtYUC4A | CTC | AAA | CTC | CAT | CTC | CCT | AAA | CAC | TTT | TGC | GAG | CTT | CCT | CTT | ATG | CCT | TTC | CCT | AAA | AAC |

(h) (codons 101–120)

| | 101 | 102 | 103 | 104 | 105 | 106 | 107 | 108 | 109 | 110 | 111 | 112 | 113 | 114 | 115 | 116 | 117 | 118 | 119 | 120 |
|---|---|---|---|---|---|---|---|---|---|---|---|---|---|---|---|---|---|---|---|---|
| | | * | * | * | * | * | * | | | | | | | | | | | | | |
| | T | P | T | F | V | S | K | L | G | F | I | N | Y | L | D | E | Y | A | T | R |
| Class-1l-AtYUC11 | ACT | CCT | ACC | TTC | GTC | TCC | AAG | CTA | GGT | TTC | ATC | AAT | TAC | CTT | GAC | GAA | TAC | GCC | ACA | CGT |
| | V | P | T | F | M | S | K | E | L | F | V | N | Y | L | D | A | Y | V | A | R |
| Class-1q-AtYUC10 | GTT | CCA | ACC | TTT | ATG | TCC | AAA | GAG | CTT | TTC | GTC | AAC | TAC | CTT | GAC | GCT | TAC | GTT | GCC | CGT |
| | Y | P | T | Y | P | T | K | Q | Q | F | V | Q | Y | L | E | S | Y | A | E | H |
| Class-3e-AtYUC2 | TAC | CCT | ACT | TAC | CCT | ACA | AAG | CAA | CAG | TTC | GTC | CAA | TAC | CTT | GAG | TCT | TAC | GCC | GAA | CAT |
| | F | P | T | Y | P | T | K | Q | Q | F | I | E | Y | L | E | D | Y | A | R | R |
| Class-3e-AtYUC6A | TTC | CCT | ACC | TAC | CCG | ACG | AAG | CAA | CAG | TTC | ATC | GAG | TAC | CTT | GAG | GAC | TAC | GCT | CGG | AGG |
| | F | P | E | Y | P | T | K | F | Q | F | I | Q | Y | L | E | S | Y | A | A | N |
| Class-4f-AtYUC3 | TTC | CCT | GAG | TAC | CCT | ACC | AAG | TTT | CAG | TTT | ATT | CAG | TAC | CTT | GAG | TCC | TAC | GCA | GCC | AAC |
| | Y | P | E | Y | P | T | K | R | Q | F | I | E | Y | L | E | S | Y | A | N | K |
| Class-4f-AtYUC5 | TAC | CCG | GAA | TAT | CCA | ACA | AAA | CGA | CAG | TTC | ATC | GAG | TAC | CTT | GAG | TCC | TAC | GCA | AAC | AAA |
| | I | P | E | Y | P | T | K | Y | Q | F | I | E | Y | L | E | S | Y | A | T | H |
| Class-4f-AtYUC7 | ATC | CCG | GAG | TAT | CCA | ACG | AAG | TAC | CAG | TTT | ATT | GAG | TAC | CTT | GAG | TCC | TAC | GCT | ACC | CAC |
| | F | P | E | Y | P | T | K | R | Q | F | I | D | Y | L | E | S | Y | A | T | R |
| Class-4f-AtYUC8 | TTC | CCG | GAG | TAC | CCG | ACG | AAG | CGT | CAG | TTC | ATC | GAC | TAC | CTT | GAG | TCT | TAC | GCT | ACC | CGG |
| | Y | P | E | Y | P | T | K | R | Q | F | I | D | Y | L | E | S | Y | A | N | R |
| Class-4f-AtYUC9 | TAC | CCT | GAA | TAC | CCA | ACG | AAA | CGA | CAG | TTC | ATC | GAT | TAC | CTT | GAG | TCA | TAC | GCA | AAC | CGG |
| | Y | P | K | Y | P | S | K | N | E | F | L | A | Y | L | E | S | Y | A | S | H |
| Class-5b-AtYUC1 | TAC | CCA | AAA | TAC | CCT | TCC | AAA | AAC | GAG | TTC | TTG | GCC | TAC | CTT | GAG | TCC | TAC | GCT | TCC | CAC |
| | F | P | K | Y | P | S | K | Q | L | F | I | S | Y | V | E | S | Y | A | A | R |
| Class-5b-AtYUC4A | TTC | CCC | AAA | TAC | CCA | TCC | AAG | CAA | CTA | TTC | ATC | TCC | TAC | GTT | GAG | TCC | TAC | GCT | GCC | CGG |

Modified PAL2NAL output. Each codon is numbered, and all motifs are designated according to the motif keys.

(h)

| | 121 | 122 | 123 | 124 | 125 | 126 | 127 | 128 | 129 | 130 | 131 | 132 | 133 | 134 | 135 | 136 | 137 | 138 | 139 | 140 |
|---|---|---|---|---|---|---|---|---|---|---|---|---|---|---|---|---|---|---|---|---|
| | * | * | | | * | * | * | * | | | * | | | | | * | | | | |
| | F | N | V | N | P | R | Y | N | R | N | V | K | S | A | Y | F | K | D | - | - |
| Class-1l-AtYUC11 | TTC | AAC | GTA | AAC | CCT | AGG | TAC | AAC | CGG | AAC | GTC | AAA | TCC | GCG | TAC | TTC | AAA | GAT | --- | --- |
| | F | D | I | N | P | R | Y | N | R | T | V | K | S | S | T | F | D | E | S | N |
| Class-1q-AtYUC10 | TTC | GAC | ATC | AAC | CCT | AGG | TAC | AAC | CGT | ACC | GTG | AAG | TCC | TCA | ACG | TTC | GAT | GAG | TCC | AAT |
| | F | D | L | K | P | V | F | N | Q | T | V | E | E | A | K | F | D | R | R | C |
| Class-3e-AtYUC2 | TTT | GAC | CTA | AAG | CCC | GTT | TTT | AAC | CAG | ACC | GTG | GAG | GAA | GCC | AAG | TTC | GAT | AGG | CGG | TGT |
| | F | D | I | K | P | E | F | N | Q | T | V | E | S | A | A | F | D | E | N | L |
| Class-3e-AtYUC6A | TTT | GAC | ATA | AAG | CCG | GAG | TTT | AAC | CAA | ACG | GTT | GAG | TCG | GCT | GCG | TTT | GAT | GAA | AAC | CTT |
| | F | D | I | N | P | K | F | N | E | T | V | Q | S | A | K | Y | D | E | T | F |
| Class-4f-AtYUC3 | TTT | GAC | ATC | AAC | CCT | AAG | TTC | AAC | GAG | ACA | GTC | CAG | TCC | GCT | AAA | TAT | GAT | GAG | ACG | TTC |
| | F | E | I | T | P | Q | F | N | E | C | V | Q | S | A | R | Y | D | E | T | S |
| Class-4f-AtYUC5 | TTC | GAA | ATT | ACT | CCT | CAG | TTC | AAC | GAG | TGT | GTC | CAG | TCT | GCT | CGA | TAC | GAC | GAG | ACA | AGT |
| | F | D | L | R | P | K | F | N | E | T | V | Q | S | A | K | Y | D | K | R | F |
| Class-4f-AtYUC7 | TTT | GAT | CTC | CGC | CCC | AAG | TTC | AAT | GAG | ACG | GTC | CAA | TCA | GCT | AAA | TAT | GAC | AAG | AGG | TTT |
| | F | E | I | N | P | K | F | N | E | C | V | Q | T | A | R | F | D | E | T | S |
| Class-4f-AtYUC8 | TTC | GAG | ATC | AAC | CCT | AAG | TTC | AAC | GAG | TGT | GTG | CAG | ACT | GCT | CGG | TTC | GAT | GAG | ACC | AGT |
| | F | D | I | K | P | E | F | N | K | S | V | E | S | A | R | F | D | E | T | S |
| Class-4f-AtYUC9 | TTC | GAT | ATT | AAA | CCG | GAG | TTT | AAT | AAA | AGC | GTT | GAG | TCT | GCT | CGG | TTT | GAT | GAA | ACT | AGC |
| | F | R | I | A | P | R | F | N | K | N | V | Q | N | A | A | Y | D | S | S | S |
| Class-5b-AtYUC1 | TTC | CGC | ATC | GCT | CCA | AGG | TTC | AAC | AAG | AAC | GTA | CAA | AAC | GCA | GCT | TAC | GAT | TCT | TCC | TCC |
| | F | N | I | K | P | V | F | N | Q | T | V | E | K | A | E | F | D | D | A | S |
| Class-5b-AtYUC4A | TTT | AAT | ATC | AAA | CCG | GTA | TTT | AAC | CAA | ACC | GTC | GAG | AAA | GCA | GAG | TTC | GAT | GAC | GCA | TCT |

| | 141 | 142 | 143 | 144 | 145 | 146 | 147 | 148 | 149 | 150 | 151 | 152 | 153 | 154 | 155 | 156 | 157 | 158 | 159 | 160 |
|---|---|---|---|---|---|---|---|---|---|---|---|---|---|---|---|---|---|---|---|---|
| | * | * | | | * | | * | | * | | | | | | | | | | * | |
| | G | Q | W | I | V | K | V | V | N | K | - | - | - | T | T | A | L | I | E | V |
| Class-1l-AtYUC11 | GGT | CAA | TGG | ATC | GTG | AAG | GTG | GTG | AAC | AAA | --- | --- | --- | ACC | ACG | GCG | TTG | ATA | GAA | GTT |
| | N | K | W | R | V | V | A | E | N | T | - | - | - | V | T | G | E | T | E | V |
| Class-1q-AtYUC10 | AAC | AAG | TGG | AGG | GTC | GTG | GCC | GAA | AAC | ACG | --- | --- | --- | GTA | ACC | GGA | GAA | ACC | GAG | GTG |
| | G | L | W | R | V | R | T | T | G | G | - | - | - | K | K | D | E | T | M | E |
| Class-3e-AtYUC2 | GGG | TTA | TGG | AGG | GTG | AGG | ACA | ACC | GGA | GGG | --- | --- | --- | AAG | AAG | GAT | GAG | ACA | ATG | GAG |
| | G | M | W | R | V | T | S | V | G | E | - | - | - | - | - | E | G | T | T | E |
| Class-3e-AtYUC6A | GGG | ATG | TGG | CGC | GTG | ACT | AGC | GTG | GGA | GAA | --- | --- | --- | --- | --- | GAA | GGC | ACG | ACG | GAG |
| | G | L | W | R | V | K | T | I | S | N | M | G | Q | L | G | S | C | E | F | E |
| Class-4f-AtYUC3 | GGG | CTG | TGG | CGG | GTT | AAG | ACC | ATT | TCA | AAC | ATG | GGT | CAG | CTC | GGT | TCT | TGC | GAG | TTT | GAG |
| | G | L | W | R | I | K | T | T | S | S | S | - | - | S | S | G | S | E | M | E |
| Class-4f-AtYUC5 | GGT | CTC | TGG | CGC | ATC | AAG | ACA | ACA | TCT | TCC | TCT | --- | --- | TCC | TCG | GGG | TCG | GAG | ATG | GAA |
| | G | L | W | R | V | Q | T | V | L | R | S | E | L | L | G | Y | C | E | F | E |
| Class-4f-AtYUC7 | GGG | CTG | TGG | CGG | GTC | CAG | ACT | GTT | TTG | AGA | AGC | GAG | CTG | CTC | GGT | TAT | TGT | GAG | TTC | GAG |
| | G | L | W | R | V | K | T | V | S | K | S | E | - | S | T | Q | T | E | V | E |
| Class-4f-AtYUC8 | GGC | TTG | TGG | CGA | GTC | AAG | ACC | GTG | TCG | AAG | AGT | GAG | --- | TCG | ACT | CAG | ACC | GAG | GTC | GAG |
| | G | L | W | R | V | R | T | T | - | - | - | - | - | S | D | G | E | E | M | E |
| Class-4f-AtYUC9 | GGG | CTA | TGG | AGG | GTT | AGA | ACA | ACC | --- | --- | --- | --- | --- | TCA | GAC | GGA | GAG | GAG | ATG | GAA |
| | G | F | W | R | V | K | T | - | - | - | - | - | - | - | - | H | D | N | T | E |
| Class-5b-AtYUC1 | GGT | TTC | TGG | AGA | GTA | AAG | ACT | --- | --- | --- | --- | --- | --- | --- | --- | CAT | GAT | AAC | ACA | GAG |
| | G | L | W | N | V | K | T | Q | D | G | - | - | - | - | - | - | - | - | - | V |
| Class-5b-AtYUC4A | GGT | CTA | TGG | AAT | GTG | AAG | ACG | CAA | GAC | GGT | --- | --- | --- | --- | --- | --- | --- | --- | --- | GTA |

(i)

| | 161 | 162 | 163 | 164 | 165 | 166 | 167 | 168 | 169 | 170 | 171 | 172 | 173 | 174 | 175 | 176 | 177 | 178 | 179 | 180 |
|---|---|---|---|---|---|---|---|---|---|---|---|---|---|---|---|---|---|---|---|---|
| | * | | | | | | | | | * | | | * | * | * | | | * | * | * |
| | Y | S | A | K | F | M | V | A | A | T | G | E | N | G | E | G | V | I | P | E |
| Class-1l-AtYUC11 | TAC | TCG | GCG | AAG | TTT | ATG | GTT | GCT | GCG | ACG | GGA | GAG | AAT | GGC | GAA | GGT | GTG | ATT | CCA | GAG |
| | Y | W | S | E | F | L | V | V | A | T | G | E | N | G | D | G | N | I | P | M |
| Class-1q-AtYUC10 | TAT | TGG | TCG | GAG | TTT | CTG | GTG | GTT | GCG | ACC | GGG | GAG | AAC | GGA | GAT | GGG | AAT | ATT | CCG | ATG |
| | Y | V | S | R | W | L | V | V | A | T | G | E | N | A | E | E | V | M | P | E |
| Class-3e-AtYUC2 | TAT | GTA | TCA | CGG | TGG | CTT | GTT | GTG | GCG | ACC | GGG | GAG | AAT | GCC | GAG | GAG | GTG | ATG | CCG | GAG |
| | Y | V | C | R | W | L | V | A | A | T | G | E | N | A | E | P | V | V | P | R |
| Class-3e-AtYUC6A | TAT | GTT | TGT | CGG | TGG | TTA | GTG | GCG | GCG | ACG | GGG | GAG | AAT | GCG | GAG | CCG | GTG | GTA | CCT | AGG |
| | Y | I | C | R | W | I | V | V | A | T | G | E | N | A | E | K | V | V | P | D |
| Class-4f-AtYUC3 | TAT | ATA | TGC | AGA | TGG | ATT | GTG | GTG | GCA | ACG | GGA | GAG | AAC | GCT | GAG | AAA | GTT | GTG | CCG | GAT |
| | Y | I | C | R | W | L | V | V | A | T | G | E | N | A | E | K | V | V | P | E |
| Class-4f-AtYUC5 | TAT | ATC | TGC | CGG | TGG | TTA | GTG | GTG | GCG | ACG | GGA | GAA | AAC | GCG | GAG | AAA | GTT | GTT | CCA | GAG |
| | Y | I | C | R | W | L | V | V | A | T | G | E | N | A | E | K | V | V | P | E |
| Class-4f-AtYUC7 | TAT | ATA | TGC | AGG | TGG | CTC | GTG | GTG | GCG | ACG | GGA | GAG | AAT | GCT | GAG | AAA | GTG | GTG | CCG | GAG |
| | Y | I | C | R | W | L | V | V | A | T | G | E | N | A | E | R | V | M | P | E |
| Class-4f-AtYUC8 | TAT | ATT | TGC | CGG | TGG | CTT | GTG | GTG | GCT | ACG | GGA | GAA | AAT | GCG | GAG | AGA | GTG | ATG | CCG | GAG |
| | Y | I | C | R | W | L | V | V | A | T | G | E | N | A | E | R | V | V | P | E |
| Class-4f-AtYUC9 | TAT | ATT | TGC | CGG | TGG | TTG | GTG | GTG | GCG | ACG | GGA | GAA | AAC | GCC | GAA | CGT | GTT | GTC | CCA | GAG |
| | Y | L | S | K | W | L | I | V | A | T | G | E | N | A | D | P | Y | F | P | E |
| Class-5b-AtYUC1 | TAC | CTC | TCC | AAA | TGG | CTT | ATC | GTA | GCC | ACC | GGT | GAG | AAC | GCA | GAT | CCA | TAC | TTC | CCC | GAG |
| | Y | T | S | T | W | L | V | V | A | T | G | E | N | A | E | P | V | F | P | N |
| Class-5b-AtYUC4A | TAC | ACA | TCC | ACG | TGG | CTG | GTG | GTT | GCC | ACA | GGG | GAA | AAC | GCT | GAA | CCG | GTG | TTT | CCG | AAC |

Modified PAL2NAL output. Each codon is numbered, and all motifs are designated according to the motif keys.

**(j)** FMO identifyng motif

| | 181 | 182 | 183 | 184 | 185 | 186 | 187 | 188 | 189 | 190 | 191 | 192 | 193 | 194 | 195 | 196 | 197 | 198 | 199 | 200 |
|---|---|---|---|---|---|---|---|---|---|---|---|---|---|---|---|---|---|---|---|---|
| | | | * | * | * | | | | | | | * | | | | | | | | * |
| | I | P | G | L | V | E | S | F | - | - | Q | G | K | Y | L | H | S | S | E | Y |
| Class-1l-AtYUC11 | ATT | CCG | GGG | CTT | GTA | GAG | AGC | TTT | --- | --- | CAA | GGA | AAG | TAT | TTG | CAC | TCA | AGT | GAG | TAC |
| | V | E | G | I | - | D | T | F | - | - | G | G | E | I | M | H | S | S | E | Y |
| Class-1q-AtYUC10 | GTG | GAG | GGG | ATT | --- | GAC | ACT | TTT | --- | --- | GGG | GGA | GAG | ATT | ATG | CAC | TCG | AGT | GAG | TAC |
| | I | D | G | I | - | P | D | F | - | - | G | G | P | I | L | H | T | S | S | Y |
| Class-3e-AtYUC2 | ATT | GAT | GGA | ATC | --- | CCG | GAT | TTT | --- | --- | GGT | GGA | CCT | ATC | CTC | CAC | ACA | AGC | TCC | TAT |
| | F | E | G | M | - | D | K | F | A | A | A | G | V | V | K | H | T | C | H | Y |
| Class-3e-AtYUC6A | TTT | GAG | GGG | ATG | --- | GAT | AAG | TTT | GCA | GCC | GCC | GGG | GTA | GTT | AAG | CAC | ACG | TGT | CAT | TAT |
| | F | E | G | L | - | E | D | F | - | - | G | G | D | V | L | H | A | G | D | Y |
| Class-4f-AtYUC3 | TTT | GAA | GGT | CTA | --- | GAG | GAT | TTT | --- | --- | GGC | GGC | GAT | GTT | CTC | CAT | GCT | GGT | GAC | TAT |
| | I | D | G | L | T | T | E | F | - | - | E | G | E | V | I | H | S | C | E | Y |
| Class-4f-AtYUC5 | ATC | GAT | GGG | CTC | ACA | ACG | GAG | TTT | --- | --- | GAA | GGC | GAG | GTG | ATC | CAC | TCG | TGT | GAG | TAC |
| | F | E | G | L | - | E | D | F | - | - | G | G | D | V | L | H | A | G | D | Y |
| Class-4f-AtYUC7 | TTT | GAG | GGT | TTA | --- | GAA | GAT | TTT | --- | --- | GGT | GGT | GAT | GTT | CTT | CAC | GCT | GGA | GAT | TAT |
| | I | D | G | L | - | S | E | F | - | - | S | G | E | V | I | H | A | C | D | Y |
| Class-4f-AtYUC8 | ATT | GAT | GGT | CTT | --- | TCT | GAG | TTT | --- | --- | TCC | GGT | GAG | GTG | ATT | CAC | GCT | TGT | GAT | TAC |
| | I | N | G | L | M | T | E | F | - | - | D | G | E | V | I | H | A | C | E | Y |
| Class-4f-AtYUC9 | ATT | AAT | GGT | CTT | ATG | ACG | GAG | TTT | --- | --- | GAC | GGA | GAA | GTG | ATT | CAC | GCT | TGT | GAG | TAT |
| | I | P | G | R | - | K | K | F | S | - | G | G | K | I | V | H | A | S | E | Y |
| Class-5b-AtYUC1 | ATT | CCA | GGG | AGA | --- | AAG | AAG | TTT | TCC | --- | GGC | GGA | AAA | ATC | GTT | CAC | GCG | AGT | GAG | TAC |
| | I | P | G | L | - | K | K | F | - | - | T | G | P | V | V | H | T | S | A | Y |
| Class-5b-AtYUC4A | ATA | CCC | GGT | TTA | --- | AAG | AAG | TTT | --- | --- | ACT | GGA | CCG | GTT | GTT | CAC | ACC | AGT | GCG | TAC |

**(k)** GXGXXG

| | 201 | 202 | 203 | 204 | 205 | 206 | 207 | 208 | 209 | 210 | 211 | 212 | 213 | 214 | 215 | 216 | 217 | 218 | 219 | 220 |
|---|---|---|---|---|---|---|---|---|---|---|---|---|---|---|---|---|---|---|---|---|
| | * | * | * | | | | | | * | | | * | * | * | | | | * | | * |
| | K | N | G | E | K | F | A | G | K | D | V | L | V | V | G | C | G | N | S | G |
| Class-1l-AtYUC11 | AAG | AAC | GGA | GAG | AAG | TTC | GCT | GGA | AAA | GAT | GTT | TTG | GTC | GTC | GGA | TGT | GGA | AAT | TCT | GGC |
| | K | S | G | R | D | F | K | D | K | N | V | L | V | V | G | G | G | N | S | G |
| Class-1q-AtYUC10 | AAG | TCC | GGT | CGT | GAT | TTT | AAA | GAT | AAA | AAT | GTT | CTT | GTG | GTC | GGA | GGT | GGA | AAT | TCC | GGT |
| | K | S | G | E | I | F | S | E | K | K | I | L | V | V | G | C | G | N | S | G |
| Class-3e-AtYUC2 | AAG | AGC | GGT | GAA | ATA | TTT | AGT | GAG | AAG | AAG | ATT | TTG | GTT | GTA | GGA | TGT | GGA | AAC | TCC | GGG |
| | K | T | G | G | D | F | A | G | K | R | V | L | V | V | G | C | G | N | S | G |
| Class-3e-AtYUC6A | AAA | ACC | GGT | GGA | GAT | TTC | GCC | GGA | AAA | AGG | GTT | CTT | GTC | GTC | GGA | TGT | GGA | AAC | TCC | GGT |
| | K | S | G | G | R | Y | Q | G | K | K | V | L | V | V | G | C | G | N | S | G |
| Class-4f-AtYUC3 | AAA | TCC | GGT | GGA | AGG | TAC | CAA | GGG | AAA | AAG | GTT | TTG | GTG | GTG | GGA | TGT | GGA | AAC | TCC | GGT |
| | K | S | G | E | K | Y | R | G | K | S | V | L | V | V | G | C | G | N | S | G |
| Class-4f-AtYUC5 | AAA | TCC | GGC | GAG | AAA | TAC | AGA | GGA | AAG | AGT | GTT | CTT | GTC | GTC | GGA | TGT | GGA | AAC | TCC | GGC |
| | K | S | G | E | R | Y | R | G | K | R | V | L | V | V | G | C | G | N | S | G |
| Class-4f-AtYUC7 | AAA | TCC | GGC | GAG | AGG | TAC | CGC | GGA | AAA | CGA | GTT | CTC | GTA | GTT | GGA | TGT | GGA | AAC | TCA | GGC |
| | K | S | G | E | K | F | A | G | K | K | V | L | V | V | G | C | G | N | S | G |
| Class-4f-AtYUC8 | AAG | TCC | GGC | GAG | AAA | TTC | GCC | GGG | AAA | AAA | GTT | CTC | GTC | GTT | GGT | TGT | GGA | AAC | TCC | GGC |
| | K | S | G | E | K | F | R | G | K | R | V | L | V | V | G | C | G | N | S | G |
| Class-4f-AtYUC9 | AAG | TCC | GGC | GAG | AAA | TTC | AGA | GGA | AAG | AGA | GTT | CTT | GTC | GTC | GGA | TGT | GGA | AAC | TCA | GGC |
| | K | S | G | E | E | F | R | R | Q | K | V | L | V | V | G | C | G | N | S | G |
| Class-5b-AtYUC1 | AAA | AGC | GGC | GAA | GAG | TTC | CGG | CGG | CAG | AAA | GTT | TTG | GTT | GTC | GGA | TGT | GGA | AAT | TCC | GGC |
| | K | S | G | S | A | F | A | N | R | K | V | L | V | V | G | C | G | N | S | G |
| Class-5b-AtYUC4A | AAG | TCC | GGT | TCG | GCA | TTC | GCA | AAC | CGG | AAG | GTT | TTG | GTG | GTT | GGT | TGT | GGA | AAT | TCC | GGT |

**(l)**

| | 221 | 222 | 223 | 224 | 225 | 226 | 227 | 228 | 229 | 230 | 231 | 232 | 233 | 234 | 235 | 236 | 237 | 238 | 239 | 240 |
|---|---|---|---|---|---|---|---|---|---|---|---|---|---|---|---|---|---|---|---|---|
| | | * | | | | * | * | * | * | * | * | * | | | | * | * | | * | |
| | M | E | I | A | Y | D | L | S | K | C | N | A | N | V | S | I | V | V | R | S |
| Class-1l-AtYUC11 | ATG | GAG | ATT | GCT | TAT | GAT | CTA | TCT | AAG | TGC | AAC | GCT | AAT | GTC | TCC | ATC | GTT | GTT | CGT | AGC |
| | M | E | I | S | F | D | L | C | N | F | G | A | N | T | T | I | L | I | R | T |
| Class-1q-AtYUC10 | ATG | GAG | ATT | AGT | TTT | GAT | CTT | TGC | AAC | TTC | GGA | GCT | AAT | ACG | ACC | ATT | CTA | ATT | AGA | ACT |
| | M | E | V | C | L | D | L | C | N | F | N | A | L | P | S | L | V | V | R | D |
| Class-3e-AtYUC2 | ATG | GAA | GTT | TGT | TTA | GAC | CTT | TGC | AAC | TTC | AAT | GCT | CTT | CCT | TCT | CTT | GTG | GTT | CGT | GAC |
| | M | E | V | C | L | D | L | C | N | F | G | A | Q | P | S | L | V | V | R | D |
| Class-3e-AtYUC6A | ATG | GAG | GTT | TGT | TTG | GAT | CTC | TGC | AAC | TTC | GGT | GCT | CAG | CCT | TCT | CTC | GTT | GTC | AGA | GAC |
| | M | E | V | S | L | D | L | Y | N | H | G | A | N | P | S | M | V | V | R | S |
| Class-4f-AtYUC3 | ATG | GAA | GTC | TCT | CTC | GAT | CTC | TAC | AAT | CAC | GGA | GCA | AAC | CCA | TCG | ATG | GTC | GTT | CGT | AGC |
| | M | E | V | S | L | D | L | A | N | H | N | A | N | A | S | M | V | V | R | S |
| Class-4f-AtYUC5 | ATG | GAA | GTC | TCT | CTT | GAT | CTC | GCA | AAT | CAC | AAT | GCT | AAT | GCA | TCC | ATG | GTT | GTT | CGT | AGC |
| | M | E | V | S | L | D | L | C | N | H | D | A | S | P | S | M | V | V | R | S |
| Class-4f-AtYUC7 | ATG | GAA | GTC | TCT | CTT | GAT | CTA | TGC | AAT | CAT | GAT | GCG | AGC | CCA | TCA | ATG | GTC | GTT | CGA | AGC |
| | M | E | V | S | L | D | L | A | N | H | F | A | K | P | S | M | V | V | R | S |
| Class-4f-AtYUC8 | ATG | GAA | GTT | TCT | CTT | GAC | CTA | GCA | AAC | CAT | TTC | GCT | AAG | CCT | TCC | ATG | GTC | GTG | AGA | AGC |
| | M | E | V | S | L | D | L | A | N | H | N | A | I | T | S | M | V | V | R | S |
| Class-4f-AtYUC9 | ATG | GAA | GTC | TCT | CTT | GAT | CTT | GCT | AAC | CAC | AAT | GCA | ATT | ACT | TCC | ATG | GTC | GTT | AGA | AGC |
| | M | E | I | S | L | D | L | V | R | H | N | A | S | P | H | L | V | V | R | N |
| Class-5b-AtYUC1 | ATG | GAA | ATT | AGC | TTA | GAC | CTC | GTC | CGA | CAT | AAC | GCA | TCT | CCT | CAT | CTT | GTT | GTC | CGG | AAC |
| | M | E | V | S | L | D | L | C | R | Y | N | A | L | P | H | M | V | V | R | N |
| Class-5b-AtYUC4A | ATG | GAG | GTC | AGC | TTG | GAT | CTT | TGT | AGA | TAC | AAT | GCT | CTG | CCT | CAT | ATG | GTC | GTT | AGA | AAC |

Modified PAL2NAL output. Each codon is numbered, and all motifs are designated according to the motif keys.

| | 241 | 242 | 243 | 244 | 245 | 246 | 247 | 248 | 249 | 250 | 251 | 252 | 253 | 254 | 255 | 256 | 257 | 258 | 259 | 260 |
|---|---|---|---|---|---|---|---|---|---|---|---|---|---|---|---|---|---|---|---|---|
| Motif | | (l) | | | | | | | | (m) | | | | | | | | | | |
| | | | * | * | | * | | * | | | * | | * | | * | | | | * | |
| | Q | V | H | V | L | T | R | C | I | V | - | - | - | - | - | R | I | G | M | S |
| Class-1l-AtYUC11 | CAG | GTG | CAC | GTG | TTA | ACG | AGA | TGT | ATC | GTA | --- | --- | --- | --- | --- | CGG | ATA | GGA | ATG | TCG |
| | P | R | H | V | V | T | K | E | V | I | - | - | - | - | - | H | L | G | M | T |
| Class-1q-AtYUC10 | CCA | AGA | CAC | GTG | GTG | ACT | AAA | GAA | GTG | ATA | --- | --- | --- | --- | --- | CAC | TTG | GGG | ATG | ACA |
| | S | V | H | V | L | P | Q | E | M | L | G | I | S | T | F | G | I | S | T | S |
| Class-3e-AtYUC2 | TCG | GTA | CAC | GTA | TTA | CCT | CAA | GAA | ATG | CTA | GGT | ATA | TCG | ACT | TTC | GGG | ATA | TCC | ACG | AGC |
| | A | V | H | V | L | P | R | E | M | L | G | T | S | T | F | G | L | S | M | F |
| Class-3e-AtYUC6A | GCT | GTG | CAC | GTC | CTA | CCA | CGA | GAG | ATG | TTG | GGT | ACT | TCA | ACT | TTT | GGG | CTG | TCC | ATG | TTC |
| | A | V | H | V | L | P | R | E | I | F | G | K | S | T | F | E | L | G | V | T |
| Class-4f-AtYUC3 | GCT | GTT | CAT | GTT | TTA | CCA | AGA | GAG | ATT | TTT | GGG | AAA | TCA | ACG | TTT | GAA | TTA | GGA | GTT | ACG |
| | S | V | H | V | L | P | R | E | I | L | G | K | S | S | F | E | I | S | M | M |
| Class-4f-AtYUC5 | TCG | GTT | CAT | GTG | TTA | CCA | AGA | GAG | ATT | TTA | GGC | AAA | TCT | AGT | TTT | GAA | ATC | TCC | ATG | ATG |
| | S | V | H | V | L | P | R | E | V | L | G | K | S | T | F | E | L | S | V | T |
| Class-4f-AtYUC7 | TCC | GTT | CAT | GTA | TTA | CCG | AGA | GAA | GTT | CTT | GGG | AAG | TCG | ACG | TTC | GAG | TTA | AGT | GTA | ACA |
| | S | L | H | V | M | P | R | E | V | M | G | K | S | T | F | E | L | A | M | K |
| Class-4f-AtYUC8 | TCT | CTT | CAC | GTG | ATG | CCG | AGG | GAA | GTA | ATG | GGT | AAA | TCA | ACG | TTT | GAG | CTT | GCA | ATG | AAG |
| | S | V | H | V | L | P | R | E | I | M | G | K | S | T | F | G | I | S | V | M |
| Class-4f-AtYUC9 | TCG | GTT | CAT | GTT | TTA | CCG | AGG | GAG | ATT | ATG | GGG | AAA | TCA | ACA | TTT | GGA | ATC | TCA | GTG | ATG |
| | T | V | H | V | L | P | R | E | I | L | G | V | S | T | F | G | V | G | M | T |
| Class-5b-AtYUC1 | ACC | GTT | CAT | GTG | TTG | CCA | AGG | GAG | ATA | CTT | GGG | GTA | TCA | ACA | TTT | GGA | GTT | GGA | ATG | ACA |
| | S | V | H | V | L | P | R | D | F | F | G | L | S | T | F | G | I | A | M | T |
| Class-5b-AtYUC4A | TCT | GTA | CAT | GTA | TTA | CCA | AGA | GAC | TTT | TTT | GGT | CTA | TCA | ACA | TTT | GGA | ATA | GCC | ATG | ACA |

| | 261 | 262 | 263 | 264 | 265 | 266 | 267 | 268 | 269 | 270 | 271 | 272 | 273 | 274 | 275 | 276 | 277 | 278 | 279 | 280 |
|---|---|---|---|---|---|---|---|---|---|---|---|---|---|---|---|---|---|---|---|---|
| Motif | | | | | | | | | | | | | (n) | | | | | | | |
| | * | | * | | | * | * | | * | * | | * | | * | * | | * | * | | |
| | L | L | R | F | F | P | V | K | L | V | D | R | L | C | L | L | L | A | E | L |
| Class-1l-AtYUC11 | TTG | CTC | AGG | TTT | TTC | CCG | GTA | AAA | TTA | GTT | GAC | CGT | TTG | TGT | CTA | TTA | CTA | GCG | GAG | CTG |
| | L | L | K | Y | A | P | V | A | M | V | D | T | L | V | T | T | M | A | K | I |
| Class-1q-AtYUC10 | CTT | CTG | AAG | TAT | GCT | CCA | GTG | GCG | ATG | GTC | GAC | ACA | TTG | GTG | ACG | ACT | ATG | GCA | AAA | ATT |
| | L | L | K | W | F | P | V | H | V | V | D | R | F | L | L | R | M | S | R | L |
| Class-3e-AtYUC2 | CTG | CTC | AAG | TGG | TTT | CCA | GTG | CAC | GTG | GTG | GAC | CGG | TTC | TTG | TTA | CGT | ATG | TCT | CGG | TTG |
| | L | L | K | W | L | P | I | R | L | V | D | R | F | L | L | V | V | S | R | F |
| Class-3e-AtYUC6A | TTA | CTG | AAA | TGG | CTG | CCC | ATC | CGG | CTT | GTT | GAC | CGT | TTC | CTT | TTG | GTT | GTT | TCC | CGG | TTC |
| | M | M | K | Y | M | P | V | W | L | A | D | K | T | I | L | F | L | A | R | I |
| Class-4f-AtYUC3 | ATG | ATG | AAG | TAT | ATG | CCC | GTT | TGG | CTC | GCG | GAC | AAG | ACT | ATA | CTC | TTT | CTG | GCG | AGG | ATT |
| | L | M | K | W | F | P | L | W | L | V | D | K | I | L | L | I | L | A | W | L |
| Class-4f-AtYUC5 | TTG | ATG | AAG | TGG | TTT | CCT | CTG | TGG | CTA | GTA | GAC | AAG | ATT | CTA | CTG | ATT | CTT | GCA | TGG | CTG |
| | M | M | K | W | M | P | V | W | L | V | D | K | T | L | L | V | L | T | R | L |
| Class-4f-AtYUC7 | ATG | ATG | AAA | TGG | ATG | CCG | GTT | TGG | CTC | GTG | GAC | AAG | ACT | CTT | CTC | GTT | CTC | ACA | AGG | TTA |
| | M | L | R | W | F | P | L | W | L | V | D | K | I | L | L | V | L | S | W | M |
| Class-4f-AtYUC8 | ATG | TTA | AGA | TGG | TTT | CCT | CTA | TGG | TTA | GTC | GAC | AAG | ATA | TTG | TTG | GTT | TTA | AGT | TGG | ATG |
| | M | M | K | W | L | P | L | W | L | V | D | K | L | L | L | I | L | S | W | L |
| Class-4f-AtYUC9 | ATG | ATG | AAG | TGG | CTA | CCT | TTA | TGG | CTC | GTA | GAC | AAG | CTT | CTG | CTT | ATC | TTA | TCG | TGG | TTG |
| | L | L | K | C | L | P | L | R | L | V | D | K | F | L | L | L | M | A | N | L |
| Class-5b-AtYUC1 | CTT | CTC | AAA | TGC | TTA | CCC | TTA | AGG | CTC | GTT | GAC | AAG | TTC | TTG | TTA | TTG | ATG | GCC | AAT | CTT |
| | L | L | K | W | F | P | L | K | L | V | D | K | F | L | L | L | L | A | N | S |
| Class-5b-AtYUC4A | CTA | CTG | AAA | TGG | TTT | CCT | CTA | AAG | CTG | GTG | GAC | AAA | TTT | CTC | CTA | CTT | CTT | GCT | AAT | TCT |

| | 281 | 282 | 283 | 284 | 285 | 286 | 287 | 288 | 289 | 290 | 291 | 292 | 293 | 294 | 295 | 296 | 297 | 298 | 299 | 300 |
|---|---|---|---|---|---|---|---|---|---|---|---|---|---|---|---|---|---|---|---|---|
| Motif | | | | | | | | | | (o) | | | | | | | | | | |
| | | * | * | * | | | * | * | * | * | * | * | * | * | * | * | * | * | | |
| | R | F | R | N | T | S | R | Y | G | L | V | R | P | N | N | G | P | F | L | N |
| Class-1l-AtYUC11 | AGG | TTT | CGG | AAT | ACT | TCG | AGA | TAT | GGG | CTT | GTA | AGA | CCC | AAT | AAT | GGC | CCA | TTT | CTG | AAC |
| | L | Y | G | D | L | S | K | Y | G | L | F | R | P | K | Q | G | P | F | A | T |
| Class-1q-AtYUC10 | TTG | TAC | GGA | GAT | CTC | TCC | AAG | TAC | GGA | CTC | TTC | CGA | CCA | AAA | CAA | GGT | CCT | TTC | GCC | ACC |
| | V | L | G | D | T | D | R | L | G | L | V | R | P | K | L | G | P | L | E | R |
| Class-3e-AtYUC2 | GTT | CTT | GGT | GAC | ACG | GAT | CGG | TTA | GGG | TTA | GTT | CGA | CCA | AAA | CTT | GGC | CCT | CTT | GAA | CGC |
| | I | L | G | D | T | T | L | L | G | L | N | R | P | R | L | G | P | L | E | L |
| Class-3e-AtYUC6A | ATC | CTC | GGG | GAT | ACT | ACC | CTT | TTA | GGT | CTT | AAC | AGG | CCC | CGG | TTA | GGT | CCA | CTC | GAG | CTC |
| | I | L | G | N | T | D | K | Y | G | L | K | R | P | K | I | G | P | L | E | L |
| Class-4f-AtYUC3 | ATC | TTG | GGG | AAT | ACC | GAT | AAA | TAC | GGT | CTA | AAA | AGG | CCG | AAA | ATT | GGA | CCG | TTA | GAG | CTA |
| | I | L | G | N | L | T | K | Y | G | L | K | R | P | T | M | G | P | M | E | L |
| Class-4f-AtYUC5 | ATT | TTA | GGA | AAT | TTG | ACA | AAG | TAT | GGC | CTA | AAG | AGG | CCC | ACG | ATG | GGC | CCA | ATG | GAG | CTC |
| | L | L | G | N | T | D | K | Y | G | L | K | R | P | E | I | G | P | L | E | L |
| Class-4f-AtYUC7 | CTT | CTA | GGG | AAC | ACC | GAT | AAG | TAT | GGG | CTC | AAG | AGG | CCT | GAA | ATT | GGA | CCG | TTA | GAG | CTG |
| | V | L | G | N | I | E | K | Y | G | L | K | R | P | E | M | G | P | M | E | L |
| Class-4f-AtYUC8 | GTT | CTT | GGA | AAC | ATC | GAG | AAA | TAC | GGT | TTG | AAA | CGA | CCA | GAG | ATG | GGT | CCA | ATG | GAG | CTA |
| | V | L | G | S | L | S | N | Y | G | L | K | R | P | D | I | G | P | M | E | L |
| Class-4f-AtYUC9 | GTT | CTA | GGG | AGC | TTA | TCA | AAC | TAT | GGG | CTT | AAA | AGG | CCT | GAC | ATC | GGC | CCA | ATG | GAG | CTT |
| | S | F | G | N | T | D | R | L | G | L | R | R | P | K | T | G | P | L | E | L |
| Class-5b-AtYUC1 | TCG | TTT | GGA | AAT | ACC | GAC | CGG | TTG | GGC | CTT | CGC | CGA | CCA | AAA | ACG | GGT | CCG | CTT | GAG | CTG |
| | T | L | G | N | T | D | L | L | G | L | R | R | P | K | T | G | P | I | E | L |
| Class-5b-AtYUC4A | ACG | TTG | GGC | AAT | ACC | GAC | CTT | TTA | GGC | CTT | CGA | CGA | CCT | AAG | ACC | GGA | CCA | ATT | GAG | CTA |

Modified PAL2NAL output. Each codon is numbered, and all motifs are designated according to the motif keys.

| | 301 | 302 | 303 | 304 | 305 | 306 | 307 | 308 | 309 | 310 | 311 | 312 | 313 | 314 | 315 | 316 | 317 | 318 | 319 | 320 |
|---|---|---|---|---|---|---|---|---|---|---|---|---|---|---|---|---|---|---|---|---|
| | * | * | | | (o) * | * | | * | | * | * | | * | | | * (p) | * | * | * | |
| | K | L | I | T | G | R | S | A | T | I | D | V | G | C | V | G | E | I | K | S |
| Class-1l-AtYUC11 | AAG | CTA | ATC | ACC | GGC | CGG | TCA | GCT | ACC | ATT | GAC | GTC | GGC | TGC | GTC | GGA | GAG | ATA | AAG | TCC |
| | K | L | F | T | G | K | A | P | V | I | D | V | G | T | V | E | K | I | R | D |
| Class-1q-AtYUC10 | AAA | CTC | TTT | ACC | GGA | AAA | GCT | CCT | GTC | ATT | GAT | GTT | GGA | ACT | GTC | GAG | AAG | ATT | CGC | GAT |
| | K | I | K | C | G | K | T | P | V | L | D | V | G | T | L | A | K | I | R | S |
| Class-3e-AtYUC2 | AAG | ATC | AAA | TGC | GGA | AAG | ACT | CCT | GTT | TTG | GAC | GTT | GGC | ACT | CTT | GCC | AAA | ATC | CGA | AGT |
| | K | N | I | S | G | K | T | P | V | L | D | V | G | T | L | A | K | I | K | T |
| Class-3e-AtYUC6A | AAA | AAT | ATC | TCC | GGT | AAA | ACT | CCG | GTT | CTC | GAC | GTT | GGC | ACG | CTA | GCC | AAA | ATC | AAA | ACC |
| | K | N | K | E | G | K | T | P | V | L | D | I | G | A | L | P | K | I | R | S |
| Class-4f-AtYUC3 | AAG | AAC | AAG | GAA | GGC | AAA | ACT | CCG | GTT | CTT | GAC | ATC | GGA | GCG | TTA | CCC | AAA | ATC | AGA | TCG |
| | K | I | V | S | G | K | T | P | V | L | D | I | G | A | M | E | K | I | K | S |
| Class-4f-AtYUC5 | AAG | ATT | GTT | TCA | GGT | AAG | ACT | CCA | GTT | CTA | GAC | ATC | GGA | GCT | ATG | GAG | AAA | ATC | AAA | TCC |
| | K | N | T | A | G | K | T | P | V | L | D | I | G | A | I | S | M | I | K | S |
| Class-4f-AtYUC7 | AAG | AAC | ACC | GCA | GGT | AAA | ACT | CCC | GTG | CTA | GAC | ATC | GGA | GCC | ATT | TCA | ATG | ATC | AAA | TCA |
| | K | S | V | K | G | K | T | P | V | L | D | I | G | A | I | E | K | I | R | L |
| Class-4f-AtYUC8 | AAA | AGC | GTG | AAA | GGC | AAG | ACA | CCG | GTG | CTT | GAC | ATC | GGA | GCT | ATA | GAG | AAG | ATC | CGG | TTA |
| | K | S | M | T | G | K | T | P | V | L | D | I | G | A | L | E | K | I | K | S |
| Class-4f-AtYUC9 | AAA | AGC | ATG | ACA | GGA | AAG | ACG | CCG | GTT | CTT | GAC | ATC | GGT | GCT | CTG | GAG | AAG | ATA | AAA | TCC |
| | K | N | V | T | G | K | S | P | V | L | D | V | G | A | M | S | L | I | R | S |
| Class-5b-AtYUC1 | AAA | AAT | GTC | ACC | GGC | AAA | AGT | CCG | GTT | CTC | GAT | GTC | GGA | GCT | ATG | TCT | CTC | ATC | AGA | TCC |
| | K | N | V | T | G | K | T | P | V | L | D | V | G | A | I | S | L | I | R | S |
| Class-5b-AtYUC4A | AAG | AAC | GTC | ACC | GGT | AAA | ACT | CCC | GTT | CTT | GAT | GTC | GGT | GCC | ATT | TCT | TTA | ATC | CGA | TCA |

| | 321 | 322 | 323 | 324 | 325 | 326 | 327 | 328 | 329 | 330 | 331 | 332 | 333 | 334 | 335 | 336 | 337 | 338 | 339 | 340 |
|---|---|---|---|---|---|---|---|---|---|---|---|---|---|---|---|---|---|---|---|---|
| | * (p) | | | | | * | | | | * | | | * | | | | | | | * |
| | G | K | I | Q | V | V | T | - | S | I | K | R | I | E | G | K | T | V | E | F |
| Class-1l-AtYUC11 | GGC | AAG | ATT | CAG | GTT | GTG | ACG | --- | TCG | ATT | AAG | CGT | ATA | GAA | GGG | AAG | ACA | GTA | GAA | TTC |
| | G | E | I | Q | V | I | N | G | G | I | G | S | I | N | G | K | T | L | T | F |
| Class-1q-AtYUC10 | GGC | GAG | ATT | CAG | GTT | ATC | AAT | GGT | GGG | ATT | GGA | AGC | ATC | AAC | GGG | AAG | ACC | TTG | ACG | TTC |
| | G | H | I | K | V | Y | P | - | E | L | K | R | V | M | H | Y | S | A | E | F |
| Class-3e-AtYUC2 | GGA | CAC | ATC | AAG | GTG | TAT | CCG | --- | GAG | TTG | AAA | CGG | GTA | ATG | CAT | TAT | TCG | GCA | GAG | TTT |
| | G | D | I | K | V | C | S | - | G | I | R | R | L | K | R | H | E | V | E | F |
| Class-3e-AtYUC6A | GGA | GAC | ATT | AAG | GTG | TGT | TCG | --- | GGG | ATA | AGA | AGG | TTA | AAA | CGA | CAT | GAA | GTT | GAG | TTC |
| | G | K | I | K | I | V | P | - | G | I | I | K | F | G | K | G | K | V | E | L |
| Class-4f-AtYUC3 | GGA | AAG | ATC | AAA | ATC | GTC | CCC | --- | GGA | ATC | ATA | AAG | TTC | GGC | AAA | GGC | AAA | GTT | GAG | CTA |
| | G | E | V | E | I | V | P | - | G | I | K | R | F | S | R | S | H | V | E | L |
| Class-4f-AtYUC5 | GGT | GAA | GTA | GAA | ATC | GTC | CCG | --- | GGA | ATT | AAA | CGG | TTC | TCT | CGT | AGT | CAC | GTG | GAG | CTA |
| | G | K | I | K | I | V | A | - | G | I | A | K | F | G | P | G | K | V | E | L |
| Class-4f-AtYUC7 | GGA | AAG | ATC | AAA | ATC | GTA | GCG | --- | GGA | ATA | GCT | AAG | TTT | GGT | CCG | GGA | AAG | GTC | GAA | CTC |
| | G | K | I | N | V | V | P | - | G | I | K | R | F | N | G | N | K | V | E | L |
| Class-4f-AtYUC8 | GGA | AAG | ATC | AAC | GTG | GTT | CCA | --- | GGG | ATC | AAA | AGG | TTT | AAC | GGA | AAC | AAA | GTC | GAA | CTT |
| | G | D | V | E | I | V | P | - | A | I | K | Q | F | S | R | H | H | V | E | L |
| Class-4f-AtYUC9 | GGC | GAC | GTA | GAA | ATT | GTA | CCT | --- | GCA | ATC | AAA | CAG | TTC | TCG | CGT | CAC | CAC | GTG | GAG | CTC |
| | G | M | I | Q | I | M | E | - | G | V | K | E | I | T | K | K | G | A | K | F |
| Class-5b-AtYUC1 | GGC | ATG | ATT | CAG | ATA | ATG | GAA | --- | GGT | GTA | AAG | GAA | ATA | ACA | AAG | AAA | GGA | GCA | AAG | TTT |
| | G | Q | I | K | V | T | Q | - | A | V | K | E | I | T | R | N | G | A | K | F |
| Class-5b-AtYUC4A | GGA | CAA | ATT | AAA | GTG | ACG | CAA | --- | GCC | GTG | AAA | GAA | ATA | ACG | AGG | AAC | GGG | GCA | AAG | TTT |

| | 341 | 342 | 343 | 344 | 345 | 346 | 347 | 348 | 349 | 350 | 351 | 352 | 353 | 354 | 355 | 356 | 357 | 358 | 359 | 360 |
|---|---|---|---|---|---|---|---|---|---|---|---|---|---|---|---|---|---|---|---|---|
| | | * | * | | | | | * | * | * | * | | * (q) F/LATGY | * | | | * | * | | |
| | I | D | G | N | T | K | N | V | D | S | I | V | F | A | T | G | Y | K | S | S |
| Class-1l-AtYUC11 | ATC | GAT | GGA | AAC | ACC | AAA | AAT | GTG | GAC | TCC | ATT | GTC | TTC | GCA | ACT | GGT | TAC | AAA | AGC | AGC |
| | E | N | G | H | K | Q | D | F | D | A | I | V | F | A | T | G | Y | K | S | S |
| Class-1q-AtYUC10 | GAA | AAC | GGA | CAT | AAA | CAA | GAT | TTC | GAT | GCG | ATT | GTG | TTT | GCT | ACT | GGT | TAC | AAA | AGC | TCC |
| | V | D | G | R | V | D | N | F | D | A | I | I | L | A | T | G | Y | K | S | N |
| Class-3e-AtYUC2 | GTT | GAT | GGG | AGA | GTA | GAT | AAC | TTC | GAC | GCC | ATT | ATA | CTC | GCC | ACG | GGT | TAC | AAA | AGC | AAC |
| | D | N | G | K | T | E | R | F | D | A | I | I | L | A | T | G | Y | K | S | N |
| Class-3e-AtYUC6A | GAT | AAC | GGA | AAA | ACA | GAG | AGA | TTT | GAC | GCC | ATT | ATA | TTA | GCA | ACT | GGC | TAC | AAA | AGC | AAC |
| | I | D | G | R | V | L | E | I | D | S | V | I | L | A | T | G | Y | R | S | N |
| Class-4f-AtYUC3 | ATA | GAT | GGC | CGT | GTT | CTT | GAG | ATT | GAT | TCC | GTT | ATT | CTA | GCT | ACT | GGA | TAC | AGA | AGC | AAC |
| | V | D | G | Q | R | L | D | L | D | A | V | V | L | A | T | G | Y | R | S | N |
| Class-4f-AtYUC5 | GTA | GAC | GGT | CAG | AGA | TTA | GAT | CTC | GAT | GCG | GTG | GTT | CTG | GCC | ACG | GGT | TAC | CGT | AGC | AAC |
| | V | D | G | R | V | L | Q | I | D | S | V | I | L | A | T | G | Y | R | S | N |
| Class-4f-AtYUC7 | GTG | GAT | GGT | CGA | GTT | CTG | CAG | ATT | GAT | TCA | GTG | ATT | CTT | GCC | ACT | GGT | TAC | AGA | AGC | AAT |
| | V | N | G | E | Q | L | D | V | D | S | V | V | L | A | T | G | Y | R | S | N |
| Class-4f-AtYUC8 | GTC | AAT | GGA | GAA | CAA | CTA | GAC | GTA | GAC | TCA | GTT | GTT | CTA | GCT | ACT | GGT | TAC | CGC | AGC | AAC |
| | V | D | G | Q | K | L | D | I | D | A | V | V | L | A | T | G | Y | R | S | N |
| Class-4f-AtYUC9 | GTA | GAT | GGT | CAG | AAG | CTA | GAT | ATC | GAC | GCT | GTG | GTT | CTC | GCC | ACC | GGT | TAT | CGT | AGC | AAC |
| | M | D | G | Q | E | K | D | F | D | S | I | I | F | A | T | G | Y | K | S | N |
| Class-5b-AtYUC1 | ATG | GAT | GGT | CAA | GAA | AAG | GAC | TTT | GAC | TCT | ATC | ATA | TTT | GCC | ACT | GGT | TAC | AAA | AGC | AAC |
| | L | N | G | K | E | I | E | F | D | S | I | I | L | A | T | G | Y | K | S | N |
| Class-5b-AtYUC4A | CTG | AAT | GGG | AAA | GAA | ATT | GAA | TTC | GAC | TCC | ATA | ATA | TTA | GCG | ACT | GGG | TAT | AAG | AGT | AAT |

Modified PAL2NAL output. Each codon is numbered, and all motifs are designated according to the motif keys.

| | 361 | 362 | 363 | 364 | 365 | 366 | 367 | 368 | 369 | 370 | 371 | 372 | 373 | 374 | 375 | 376 | 377 | 378 | 379 | 380 |
|---|---|---|---|---|---|---|---|---|---|---|---|---|---|---|---|---|---|---|---|---|
| | | * | | | * | | | | * | | | * | * | (r) | | | | * | * | * |
| | V | S | K | W | L | E | V | D | D | G | D | L | F | N | - | E | N | G | M | P |
| Class-1l-AtYUC11 | GTT | TCA | AAA | TGG | CTT | GAG | GTC | GAT | GAT | GGA | GAT | CTG | TTC | AAC | --- | GAG | AAT | GGG | ATG | CCG |
| | V | C | N | W | L | E | - | - | D | Y | E | Y | V | M | K | K | D | G | F | P |
| Class-1q-AtYUC10 | GTT | TGC | AAT | TGG | TTA | GAG | --- | --- | GAC | TAT | GAA | TAC | GTG | ATG | AAG | AAG | GAT | GGA | TTC | CCC |
| | V | P | M | W | L | K | - | - | G | V | N | M | F | S | E | K | D | G | F | P |
| Class-3e-AtYUC2 | GTA | CCC | ATG | TGG | CTA | AAG | --- | --- | GGA | GTG | AAC | ATG | TTT | TCT | GAG | AAA | GAT | GGA | TTT | CCG |
| | V | P | S | W | L | K | - | - | E | N | K | M | F | S | K | K | D | G | F | P |
| Class-3e-AtYUC6A | GTA | CCC | TCT | TGG | CTA | AAG | --- | --- | GAG | AAT | AAA | ATG | TTT | AGT | AAG | AAA | GAT | GGA | TTT | CCA |
| | V | P | S | W | L | K | - | - | D | N | D | F | F | S | - | D | D | G | I | P |
| Class-4f-AtYUC3 | GTC | CCT | TCA | TGG | CTT | AAG | --- | --- | GAC | AAC | GAC | TTC | TTC | TCC | --- | GAT | GAT | GGG | ATA | CCA |
| | V | P | S | W | L | Q | - | - | E | N | D | L | F | S | - | K | N | G | F | P |
| Class-4f-AtYUC5 | GTC | CCA | TCA | TGG | CTC | CAA | --- | --- | GAA | AAT | GAT | CTG | TTT | TCG | --- | AAA | AAC | GGG | TTC | CCG |
| | V | P | S | W | L | K | - | - | E | N | D | - | L | G | - | E | I | G | I | E |
| Class-4f-AtYUC7 | GTC | CCT | TCA | TGG | CTT | AAG | --- | --- | GAA | AAC | GAC | --- | TTG | GGT | --- | GAG | ATT | GGG | ATA | GAG |
| | V | P | Y | W | L | Q | - | - | E | N | E | F | F | A | - | K | N | G | F | P |
| Class-4f-AtYUC8 | GTC | CCA | TAT | TGG | CTA | CAA | --- | --- | GAG | AAT | GAG | TTC | TTT | GCA | --- | AAG | AAT | GGT | TTC | CCA |
| | V | P | S | W | L | Q | - | - | E | S | E | F | F | S | - | K | N | G | F | P |
| Class-4f-AtYUC9 | GTC | CCT | TCT | TGG | CTT | CAG | --- | --- | GAA | AGT | GAG | TTC | TTT | TCG | --- | AAA | AAT | GGG | TTT | CCG |
| | V | P | T | W | L | Q | - | - | G | G | D | F | F | T | - | D | D | G | M | P |
| Class-5b-AtYUC1 | GTG | CCT | ACT | TGG | CTT | CAG | --- | --- | GGA | GGT | GAT | TTT | TTC | ACG | --- | GAC | GAT | GGG | ATG | CCG |
| | V | P | D | W | L | K | - | - | E | N | S | F | F | T | - | K | E | G | M | P |
| Class-5b-AtYUC4A | GTA | CCC | GAT | TGG | CTT | AAG | --- | --- | GAG | AAT | AGT | TTT | TTC | ACA | --- | AAA | GAA | GGA | ATG | CCA |

| | 381 | 382 | 383 | 384 | 385 | 386 | 387 | 388 | 389 | 390 | 391 | 392 | 393 | 394 | 395 | 396 | 397 | 398 | 399 | 400 |
|---|---|---|---|---|---|---|---|---|---|---|---|---|---|---|---|---|---|---|---|---|
| | | | * | | * | | | * | | | * | | (s) | | * | * | | * | * | * |
| | K | R | E | F | P | - | D | H | W | K | G | K | N | G | L | Y | S | A | G | F |
| Class-1l-AtYUC11 | AAA | CGC | GAG | TTC | CCG | --- | GAT | CAC | TGG | AAG | GGG | AAG | AAT | GGT | TTG | TAT | AGT | GCT | GGC | TTT |
| | K | A | P | M | P | - | K | H | W | K | G | E | K | N | L | Y | C | A | G | F |
| Class-1q-AtYUC10 | AAA | GCA | CCG | ATG | CCA | --- | AAA | CAT | TGG | AAA | GGA | GAG | AAG | AAT | CTT | TAC | TGT | GCC | GGA | TTT |
| | H | K | P | F | P | - | N | G | W | K | G | E | S | G | L | Y | A | V | G | F |
| Class-3e-AtYUC2 | CAT | AAA | CCA | TTT | CCT | --- | AAC | GGT | TGG | AAA | GGC | GAA | AGC | GGA | TTG | TAT | GCA | GTC | GGT | TTC |
| | I | Q | E | F | P | - | E | G | W | R | G | E | C | G | L | Y | A | V | G | F |
| Class-3e-AtYUC6A | ATA | CAA | GAG | TTC | CCT | --- | GAG | GGA | TGG | AGA | GGG | GAA | TGT | GGG | CTA | TAC | GCG | GTC | GGA | TTC |
| | K | N | P | F | P | - | N | G | W | K | G | E | A | G | L | Y | A | V | G | F |
| Class-4f-AtYUC3 | AAA | AAC | CCG | TTT | CCA | --- | AAC | GGA | TGG | AAA | GGA | GAG | GCG | GGA | TTG | TAT | GCG | GTA | GGG | TTC |
| | K | S | P | F | P | - | N | A | W | K | G | K | S | G | L | Y | A | A | G | F |
| Class-4f-AtYUC5 | AAA | TCG | CCG | TTT | CCA | --- | AAC | GCG | TGG | AAA | GGG | AAA | TCG | GGA | CTT | TAC | GCG | GCT | GGA | TTC |
| | K | N | P | F | P | - | K | G | W | K | G | K | A | G | L | Y | A | V | G | F |
| Class-4f-AtYUC7 | AAA | AAC | CCA | TTT | CCA | --- | AAA | GGG | TGG | AAA | GGG | AAG | GCT | GGA | TTG | TAT | GCG | GTG | GGA | TTC |
| | K | T | V | A | D | N | N | G | W | K | G | R | T | G | L | Y | A | V | G | F |
| Class-4f-AtYUC8 | AAG | ACA | GTA | GCT | GAC | AAC | AAT | GGT | TGG | AAA | GGA | AGG | ACA | GGA | TTG | TAT | GCG | GTT | GGG | TTT |
| | K | S | P | F | P | - | N | A | W | K | G | K | S | G | L | Y | A | A | G | F |
| Class-4f-AtYUC9 | AAA | TCG | CCG | TTT | CCA | --- | AAC | GCG | TGG | AAA | GGG | AAA | TCG | GGA | TTA | TAC | GCG | GCC | GGA | TTC |
| | K | T | P | F | P | - | N | G | W | R | G | G | K | G | L | Y | T | V | G | F |
| Class-5b-AtYUC1 | AAA | ACG | CCG | TTT | CCT | --- | AAC | GGC | TGG | AGA | GGA | GGG | AAA | GGA | TTG | TAC | ACA | GTT | GGT | TTT |
| | K | T | P | F | P | - | N | G | W | K | G | E | K | G | L | Y | T | V | G | F |
| Class-5b-AtYUC4A | AAA | ACG | CCG | TTT | CCT | --- | AAC | GGA | TGG | AAA | GGA | GAG | AAG | GGA | CTA | TAC | ACG | GTG | GGT | TTC |

| | 401 | 402 | 403 | 404 | 405 | 406 | 407 | 408 | 409 | 410 | 411 | 412 | 413 | 414 | 415 | 416 | 417 | 418 | 419 | 420 |
|---|---|---|---|---|---|---|---|---|---|---|---|---|---|---|---|---|---|---|---|---|
| | | | (s) | * | * | | * | | | | * | * | * | | * | | | * | * | |
| | G | K | Q | G | L | A | G | I | S | R | D | A | R | N | I | A | R | D | I | D |
| Class-1l-AtYUC11 | GGG | AAA | CAA | GGC | TTG | GCT | GGT | ATT | TCA | AGA | GAT | GCT | CGG | AAC | ATT | GCT | AGA | GAC | ATT | GAT |
| | S | R | K | G | I | A | G | G | A | E | D | A | M | S | V | A | D | D | I | R |
| Class-1q-AtYUC10 | TCA | AGA | AAA | GGA | ATC | GCC | GGA | GGT | GCT | GAG | GAT | GCT | ATG | TCT | GTA | GCC | GAC | GAC | ATC | AGA |
| | T | K | L | G | L | L | G | A | A | I | D | A | K | K | I | A | E | D | I | E |
| Class-3e-AtYUC2 | ACA | AAG | CTT | GGA | TTG | CTT | GGT | GCA | GCC | ATT | GAT | GCC | AAG | AAG | ATC | GCT | GAG | GAT | ATT | GAG |
| | T | K | R | G | I | S | G | A | S | M | D | A | K | R | I | A | E | D | I | H |
| Class-3e-AtYUC6A | ACA | AAA | CGT | GGG | ATT | AGT | GGA | GCA | TCA | ATG | GAT | GCA | AAG | AGA | ATA | GCT | GAA | GAC | ATA | CAC |
| | T | R | K | G | L | F | G | A | S | L | D | A | M | S | V | A | H | D | I | A |
| Class-4f-AtYUC3 | ACG | AGA | AAA | GGA | TTG | TTT | GGT | GCG | TCG | CTT | GAT | GCT | ATG | AGT | GTG | GCT | CAT | GAT | ATT | GCT |
| | T | R | K | G | L | A | G | A | S | A | D | A | V | N | I | A | Q | D | I | G |
| Class-4f-AtYUC5 | ACA | AGG | AAA | GGA | TTG | GCT | GGA | GCA | TCA | GCA | GAC | GCC | GTT | AAC | ATC | GCT | CAA | GAT | ATT | GGA |
| | T | G | R | G | L | S | G | A | S | F | D | A | M | S | V | A | H | D | I | A |
| Class-4f-AtYUC7 | ACC | GGA | AGA | GGA | CTC | TCA | GGT | GCA | TCG | TTC | GAT | GCA | ATG | AGT | GTA | GCT | CAT | GAC | ATT | GCC |
| | T | R | K | G | L | S | G | A | S | M | D | A | V | K | I | A | Q | D | I | G |
| Class-4f-AtYUC8 | ACG | AGG | AAA | GGG | CTC | TCA | GGT | GCG | TCA | ATG | GAC | GCG | GTT | AAG | ATC | GCA | CAA | GAC | ATA | GGC |
| | T | R | K | G | L | A | G | A | S | V | D | A | V | N | I | A | Q | D | I | G |
| Class-4f-AtYUC9 | ACG | AGA | AAA | GGG | TTG | GCT | GGA | GCA | TCA | GTA | GAC | GCC | GTT | AAC | ATC | GCT | CAA | GAC | ATT | GGA |
| | T | R | R | G | L | L | G | T | A | S | D | A | V | K | I | A | G | E | I | G |
| Class-5b-AtYUC1 | ACG | AGA | AGA | GGA | CTC | CTT | GGA | ACG | GCG | TCT | GAC | GCC | GTT | AAG | ATC | GCT | GGC | GAA | ATT | GGT |
| | T | R | R | G | L | S | G | T | A | Y | D | A | V | K | I | A | E | D | I | T |
| Class-5b-AtYUC4A | ACG | AGA | AGA | GGA | CTT | TCG | GGA | ACA | GCC | TAT | GAC | GCC | GTT | AAG | ATC | GCT | GAG | GAT | ATA | ACA |

Modified PAL2NAL output. Each codon is numbered, and all motifs are designated according to the motif keys.

| | 421 | 422 | 423 | 424 | 425 | 426 | 427 | 428 * | 429 | 430 | 431 | 432 | 433 | 434 | 435 | 436 | 437 | 438 | 439 * | 440 |
|---|---|---|---|---|---|---|---|---|---|---|---|---|---|---|---|---|---|---|---|---|
| | S | L | V | C | G | R | S | S | K | - | N | K | L | S | K | - | - | - | - | - |
| Class-1l-AtYUC11 | TCT | CTG | GTT | TGT | GGG | AGA | AGC | AGC | AAG | --- | AAC | AAG | CTT | TCC | AAA | --- | --- | --- | --- | --- |
| | S | I | L | A | T | L | K | N | N | - | - | - | - | - | - | - | - | - | - | - |
| Class-1q-AtYUC10 | TCT | ATC | TTG | GCT | ACC | TTA | AAA | AAT | AAT | --- | --- | --- | --- | --- | --- | --- | --- | --- | --- | --- |
| | V | Q | - | - | - | - | - | - | - | - | R | H | F | L | P | L | A | R | P | Q |
| Class-3e-AtYUC2 | GTT | CAA | --- | --- | --- | --- | --- | --- | --- | --- | CGA | CAT | TTC | TTA | CCA | TTG | GCT | CGT | CCT | CAA |
| | K | C | W | K | Q | D | E | Q | V | - | K | K | I | - | - | - | - | - | - | - |
| Class-3e-AtYUC6A | AAG | TGT | TGG | AAA | CAA | GAC | GAG | CAA | GTA | --- | AAA | AAA | ATC | --- | --- | --- | --- | --- | --- | --- |
| | N | R | W | K | E | E | S | K | Q | - | Q | K | K | T | A | A | A | R | H | R |
| Class-4f-AtYUC3 | AAT | AGG | TGG | AAA | GAA | GAG | TCT | AAG | CAA | --- | CAG | AAG | AAA | ACT | GCT | GCT | GCT | CGT | CAT | CGT |
| | N | V | W | R | E | E | T | K | R | - | Q | K | M | R | T | R | V | G | H | R |
| Class-4f-AtYUC5 | AAC | GTG | TGG | AGA | GAA | GAA | ACC | AAA | CGA | --- | CAG | AAA | ATG | AGA | ACA | CGA | GTG | GGT | CAC | CGC |
| | N | S | W | K | E | E | T | K | Q | - | Q | I | K | T | V | A | T | R | H | R |
| Class-4f-AtYUC7 | AAT | AGC | TGG | AAA | GAA | GAA | ACT | AAG | CAA | --- | CAG | ATC | AAA | ACT | GTC | GCT | ACT | CGC | CAC | CGT |
| | S | V | W | Q | L | E | T | K | Q | - | P | T | K | R | S | R | G | S | L | R |
| Class-4f-AtYUC8 | TCT | GTT | TGG | CAA | CTT | GAA | ACA | AAA | CAA | --- | CCC | ACG | AAA | CGC | TCA | AGG | GGT | TCT | CTT | CGA |
| | N | V | W | R | E | E | T | K | R | - | Q | K | M | R | R | N | V | G | H | R |
| Class-4f-AtYUC9 | AAT | GTG | TGG | AGA | GAA | GAG | ACC | AAA | CGA | --- | CAG | AAG | ATG | AGA | AGA | AAT | GTG | GGT | CAC | CGA |
| | D | Q | W | R | D | E | I | K | G | - | S | T | R | N | M | - | - | - | - | - |
| Class-5b-AtYUC1 | GAC | CAG | TGG | AGA | GAC | GAA | ATC | AAG | GGG | --- | TCC | ACC | AGG | AAT | ATG | --- | --- | --- | --- | --- |
| | D | Q | W | M | K | F | N | G | P | L | S | C | R | N | I | - | - | - | - | - |
| Class-5b-AtYUC4A | GAC | CAA | TGG | ATG | AAA | TTT | AAC | GGT | CCG | TTG | AGT | TGT | AGG | AAT | ATT | --- | --- | --- | --- | --- |

(t) (u)

| | 441 | 442 * | 443 * | 444 * | 445 | 446 | 447 | 448 | 449 | 450 | 451 | 452 | 453 | 454 |
|---|---|---|---|---|---|---|---|---|---|---|---|---|---|---|
| | - | - | - | - | - | - | - | - | - | - | - | - | - | - |
| Class-1l-AtYUC11 | --- | --- | --- | --- | --- | --- | --- | --- | --- | --- | --- | --- | --- | --- |
| | - | - | - | - | - | - | - | - | - | - | - | - | - | - |
| Class-1q-AtYUC10 | --- | --- | --- | --- | --- | --- | --- | --- | --- | --- | --- | --- | --- | --- |
| | H | C | - | - | - | - | - | - | - | - | - | - | - | - |
| Class-3e-AtYUC2 | CAT | TGT | --- | --- | --- | --- | --- | --- | --- | --- | --- | --- | --- | --- |
| | - | - | - | - | - | - | - | - | - | - | - | - | - | - |
| Class-3e-AtYUC6A | --- | --- | --- | --- | --- | --- | --- | --- | --- | --- | --- | --- | --- | --- |
| | R | C | I | S | H | F | - | - | - | - | - | - | - | - |
| Class-4f-AtYUC3 | CGT | TGT | ATT | TCA | CAT | TTC | --- | --- | --- | --- | --- | --- | --- | --- |
| | R | C | I | S | V | A | - | - | - | - | - | - | - | - |
| Class-4f-AtYUC5 | AGA | TGC | ATT | TCA | GTT | GCT | --- | --- | --- | --- | --- | --- | --- | --- |
| | R | C | I | S | H | F | - | - | - | - | - | - | - | - |
| Class-4f-AtYUC7 | CGT | TGT | ATT | TCA | CAC | TTT | --- | --- | --- | --- | --- | --- | --- | --- |
| | R | C | I | S | Q | Q | F | - | - | - | - | - | - | - |
| Class-4f-AtYUC8 | AGG | TGT | ATC | TCT | CAA | CAG | TTC | --- | --- | --- | --- | --- | --- | --- |
| | R | C | I | S | V | A | - | - | - | - | - | - | - | - |
| Class-4f-AtYUC9 | AGA | TGC | ATC | TCA | GTT | GCT | --- | --- | --- | --- | --- | --- | --- | --- |
| | - | C | S | S | R | F | V | F | T | - | - | S | K | S |
| Class-5b-AtYUC1 | --- | TGC | AGT | TCT | CGT | TTT | GTC | TTT | ACC | --- | --- | TCT | AAA | TCC |
| | - | C | S | S | H | I | I | H | L | H | F | N | K | S |
| Class-5b-AtYUC4A | --- | TGT | AGT | TCC | CAT | ATT | ATT | CAT | CTT | CAT | TTC | AAT | AAA | TCC |

(u)

(PAL2NAL output from https://www.bork.embl.de/pal2nal/pal2nal.with_codeml.v14.cgi)

PAL2NAL Reference: -

Mikita Suyama, David Torrents, and Peer Bork (2006). PAL2NAL: robust conversion of protein sequence alignments into the corresponding codon alignments. Nucleic Acids Res. 34, W609-W612.

Modified PAL2NAL output. Each codon is numbered, and all motifs are designated according to the motif keys.
