## Supplementary Information (Supplementary Figures and Tables (S1-S2)) for "The auxin gatekeepers: Evolution and diversification of the YUCCA family"

**(All supplementary Figures and Tables are listed together here)**

#### INDEX

**Supplementary Figures:** pages 1-12


| No | Short title | Page no |
|---|---|---|
| Figure S1 | YUCs classification scheme | 2 |
| Figure S2 | Pairwise diversity between class B FMOs and within YUCs classes | 3 |
| Figure S3 | YUCs diversity in various plant groups | 4 |
| Figure S4 | Arabidopsis YUCs (AtYUCs) - motif and structural conservation patterns | 5 |
| Figure S5 | Arabidopsis YUCs – Protein topography and electrostatic surface potential | 6 |
| Figure S6 | Evolutionary selection pressure analysis on AtYUCs | 7 |
| Figure S7 | Comparison of subcellular localisation at the species level | 8 |
| Figure S8 | Predicted subcellular localisations and membrane type details | 9 |
| Figure S9 | Arabidopsis TAA/TARs - structural conservation pattern | 10 |
| Figure S10 | *A. thaliana* YUCs (AtYUCs) protein-protein interaction network | 11-12 |


**Supplementary Tables:** pages 13-14


| No | Short title | Page no |
|---|---|---|
| Table S1 | Overall view of motif representation across various studied FMOs | 13 |
| Table S2 | Overall view of motif representation in various YUCs across various plant classes | 14 |

**Figure S1**. **YUCCA (YUC) classification scheme** [Classification system referred to and according to the rules of CYP nomenclature system (Nelson 2006)]

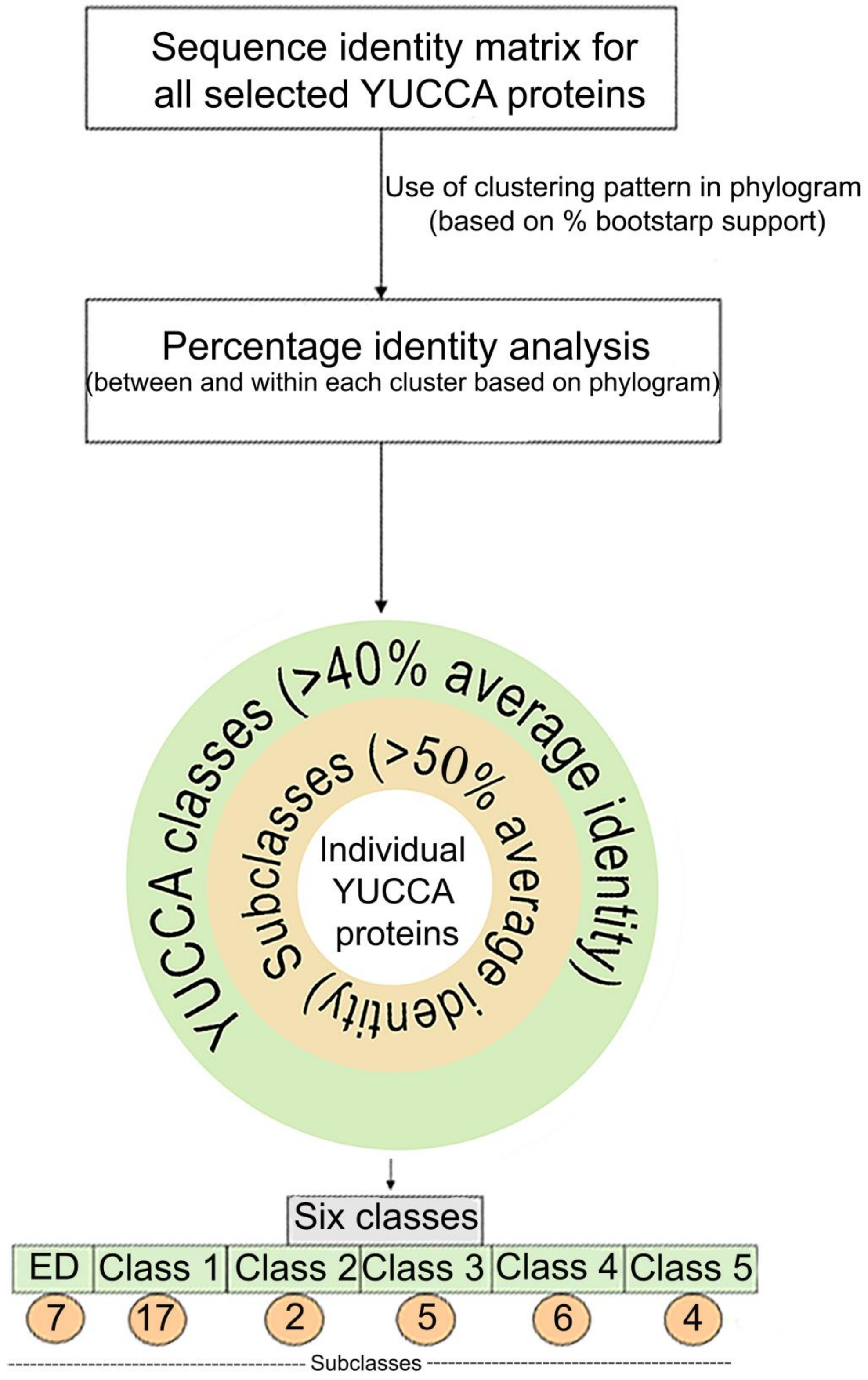

**Figure S2. Pairwise diversity between class B FMOs and within YUCs classes**. **A)** Pairwise distance (diversity) between various class B FMOs [YUCs (Clade III FMO), sYUCs (Clade details not available), N-OX (Clade II FMO) and GS-OX (Clade IV FMO)]. **B)** YUC diversity within various classes/subclasses – which further supports the YUC classification schema. **[**Pairwise-based diversity within the group is calculated in the DIVEIN webserver using the multiple sequence alignment fasta file (URL: https://indra.mullins.microbiol.washington.edu/DIVEIN/index.html, accessed on June 2024). More details are given in the **Data S2D-E**.]

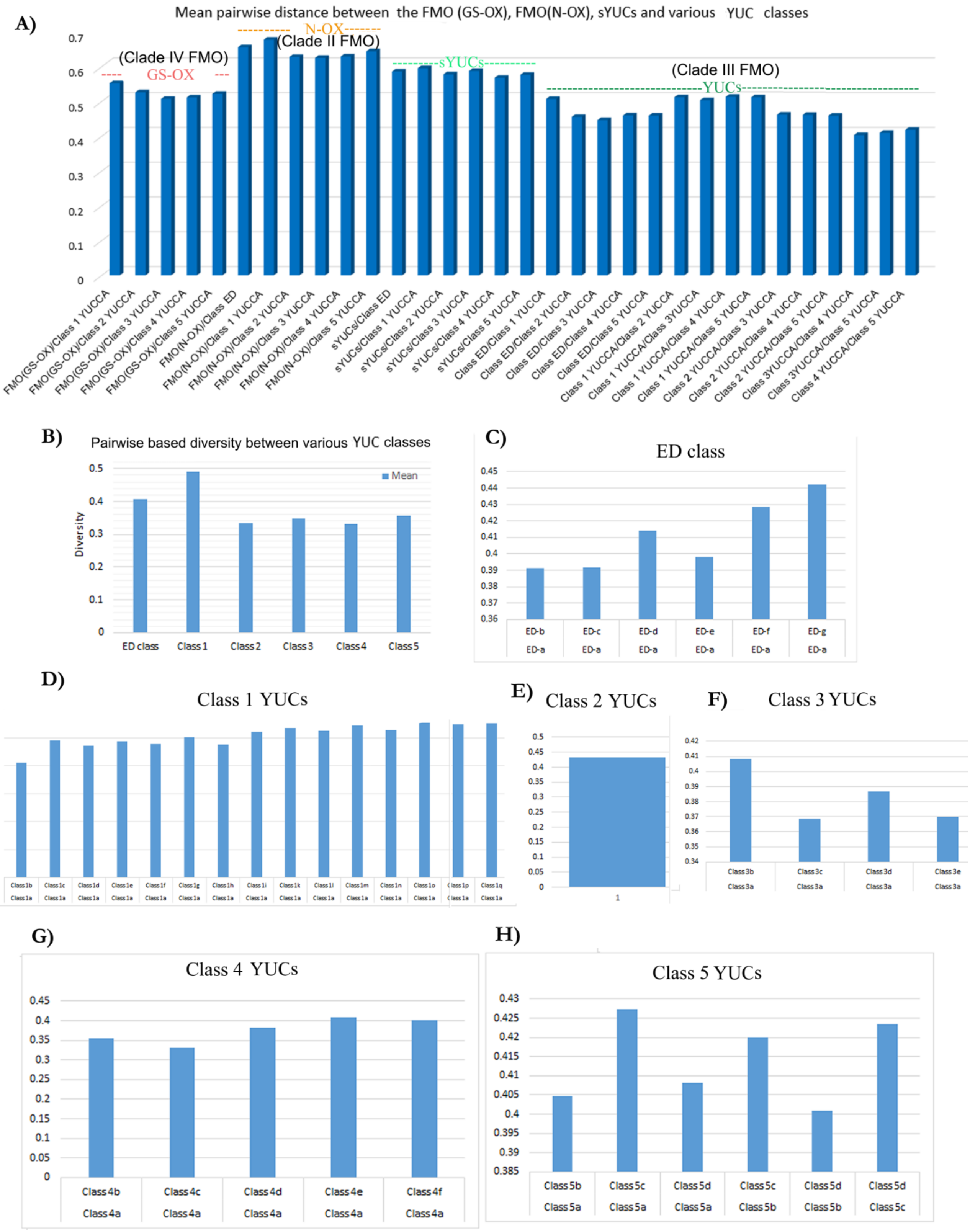

**Figure S3. YUCs diversity in various plant groups.** Pairwise based diversity within various taxa is calculated in the DIVEIN webserver based on the multiple sequence alignment fasta file (URL: https://indra.mullins.microbiol.washington.edu/DIVEIN/index.html, accessed on June 2024). More details in Data S2D. **Note**: This graphic is based on the analysis of 469 YUCs, which includes limited representation from Hornworts, Liverworts, and other Early diverging groups. However, all taxonomic groups show a trend of diversification and here angiosperms YUCs show considerably more diversity in comparison to other lineages.

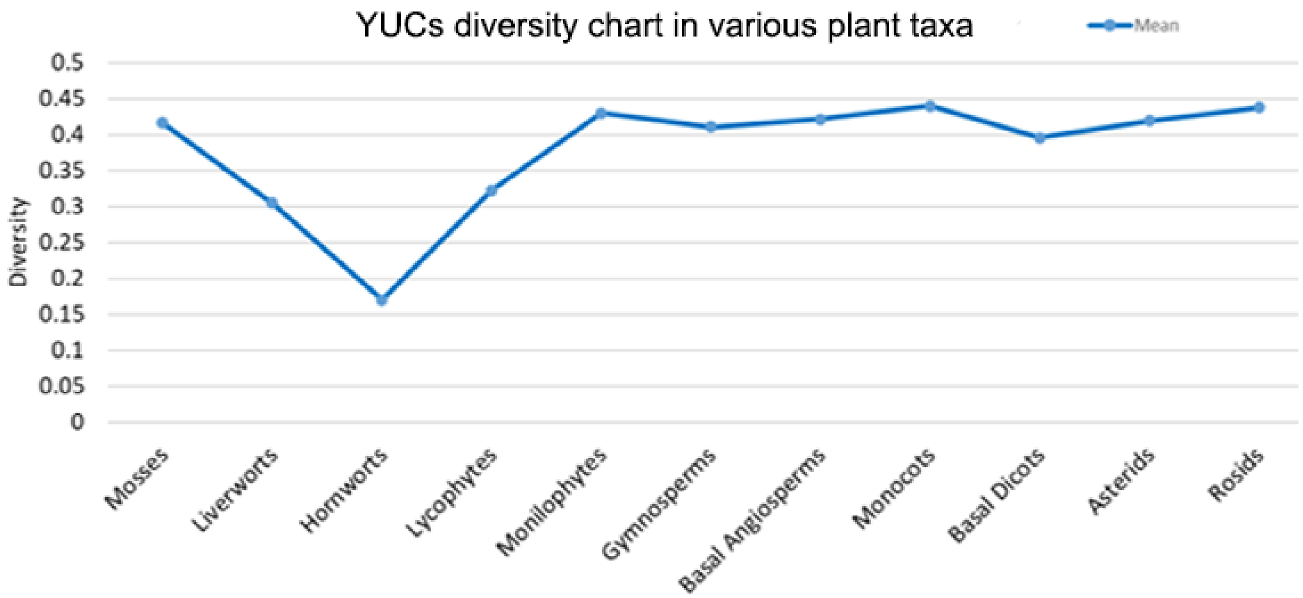

**Figure S4. Motif and structural conservation patterns**. **A)** Motif organization of AtYUCs based on their respective classes. Seven distinct motif pattern types are observed among all AtYUCs, highlighting their diversity and unique characteristics. **B**) Motif organization of Moss (*Physcomitrella patens*) PpYUCs. **C)** Phylogram representing AtYUCs AlphaFold structures. In the phylogram, major classes are highlighted in blue circles and numbered (No model for class 2, as no AtYUCs belong to that class). Subclasses if any (only for AtYUCs), is also depicted at the base of the respective branch. AtYUC6 with reported TR activity is highlighted in red star (*). Different motif areas are highlighted and marked in each structure. Common YUC motif patterns, like the signature motifs, are not highlighted. AtYUCs from Class 1 (AtYUC10-11) possess Type-5 motif type with GAXTSIVVRX(3)H motif while other class AtYUCs possess PREXXG containing motif in the corresponding position. *N*-terminal RRCVWV motif is confidently predicted in AtYUC2, AtYUC3,5,7,8,9 however their putative presence is predicted in AtYUC1, AtYUC4A-B., AtYUC6A-B as well- hence marked as 'p' (putative). Motif analysis was performed in the MEME suite (more details **Data S5**).

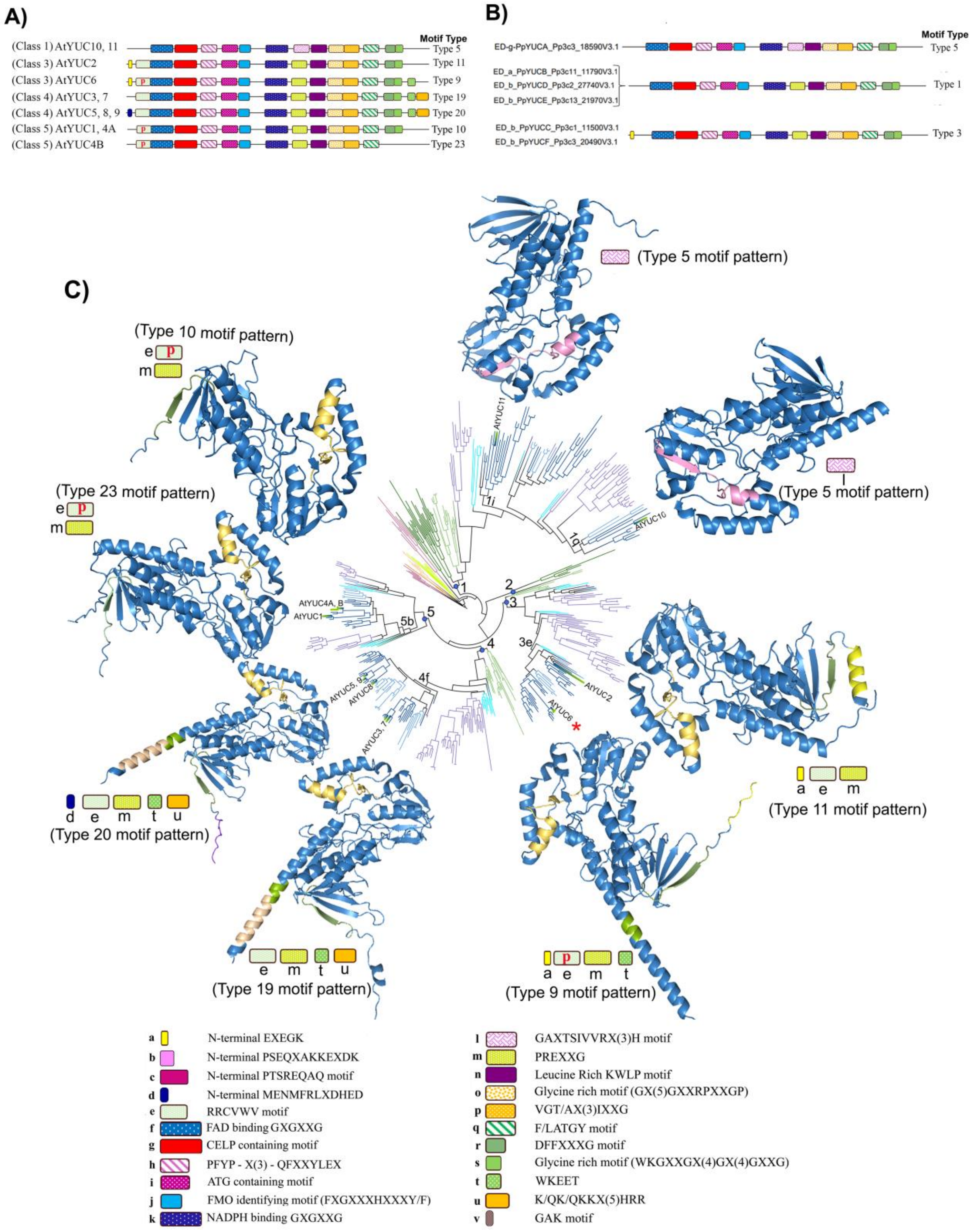

**Figure S5. Arabidopsis YUCs – Protein topography and electrostatic surface potential**. **A)** Substrate binding cavity details. **B)** Electrostatic surface potential (ESP) details of AtYUCs. The structures display electrostatic charge distribution (positively charged residues in blue and negatively charged residues in red). Substrate binding cavities in each protein are marked with an orange arrow. **C)** Cavity parameters. Color scale: Blue (high) - Red (low). (CASTpFold server was used for predicting cavity details (area and volume of substrate binding pocket), https://cfold.bme.uic.edu/castpfold/, accessed October 2024. ASPB electrostatics plugin in PyMOL was used for ESP calculations)

A)Protein topography with surface pocket details B)Electrostatic surface potential in AtYUCs

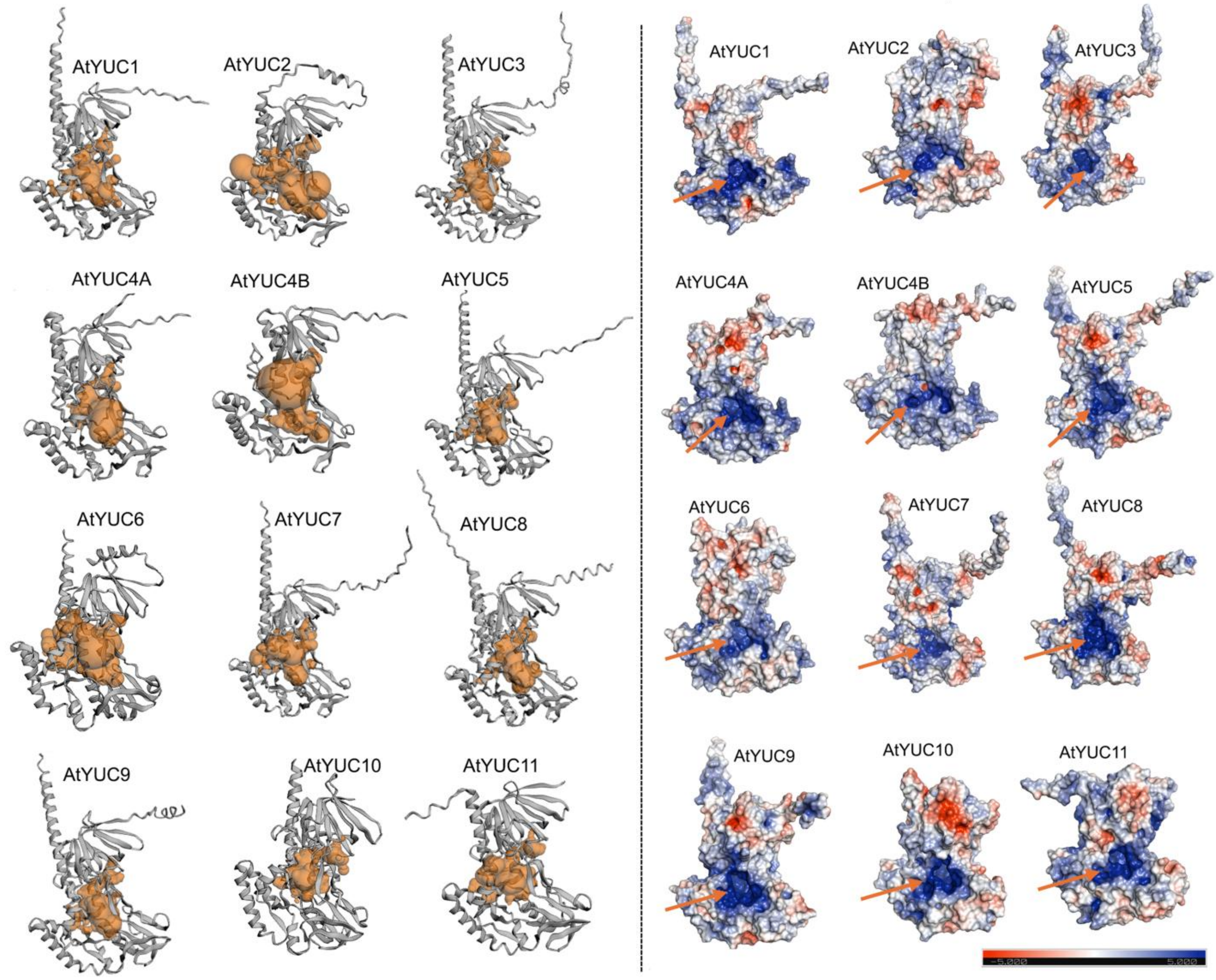


C)

| Protein | The surface area (SA) and volume (V) of substrate binding pocket | |
|---|---|---|
| | Area (SA) (Å2) | Volume (V)(Å3) |
| AtYUC1 | 1091.208 | 836.157 |
| AtYUC2 | 1908.96 | 1858.948 |
| AtYUC3 | 1118.449 | 808.676 |
| AtYUC4A | 1120.554 | 1084.734 |
| AtYUC4B | 1236.019 | 1228.993 |
| AtYUC5 | 1046.323 | 898.813 |
| AtYUC6 | 1844.782 | 1701.192 |
| AtYUC7 | 1503.857 | 1276.37 |
| AtYUC8 | 1141.429 | 951.058 |
| AtYUC9 | 1137.565 | 870.011 |
| AtYUC10 | 1001.705 | 758.633 |
| AtYUC11 | 1101.679 | 853.854 |
| **Mean** | **1271.0 4 Å** | **1093.95 Å** |

**Figure S6. Evolutionary selection pressure analysis on AtYUCs** (Done in Datamonkey server, used PAL2NAL output of AtYUCs nucleotide alignment)

A) **SLAC (Single-Likelihood Ancestor Counting) output displaying purifying selection for AtYCs.** **S**ynonymous and non-synonymous substitution rates based on 11 AtYUCs codon-aligned nucleotide sequences. (SLAC module used in Datamonkey server: https://www.datamonkey.org/slac, accessed on December 2024). Results suggested that most sites in AtYUCs are either under negative (purifying) selection (184 sites) or evolving neutrally.

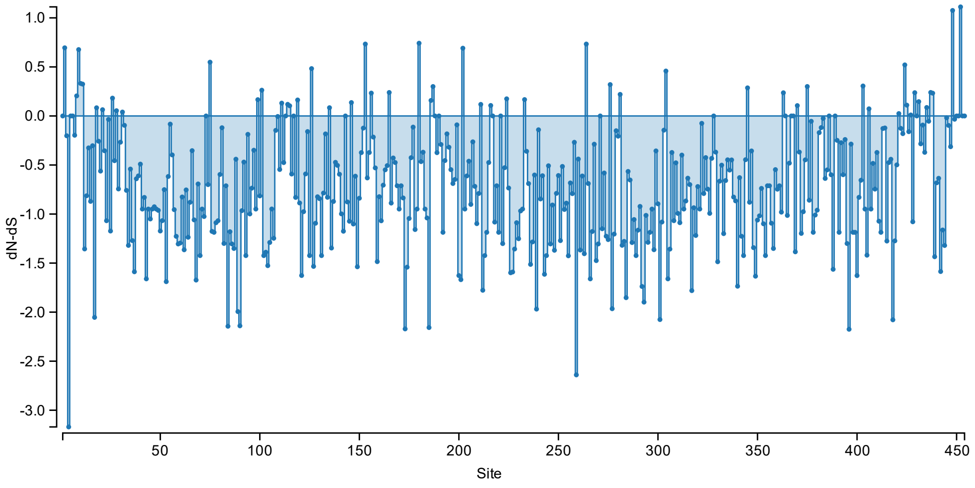


B) **MEME v4.0 (Mixed Effects Model of Evolution) estimates for positive selection.** Analysis shows 11 sites under diversifying selection (More codon-wise details and their respective values are given in **Data S7**)

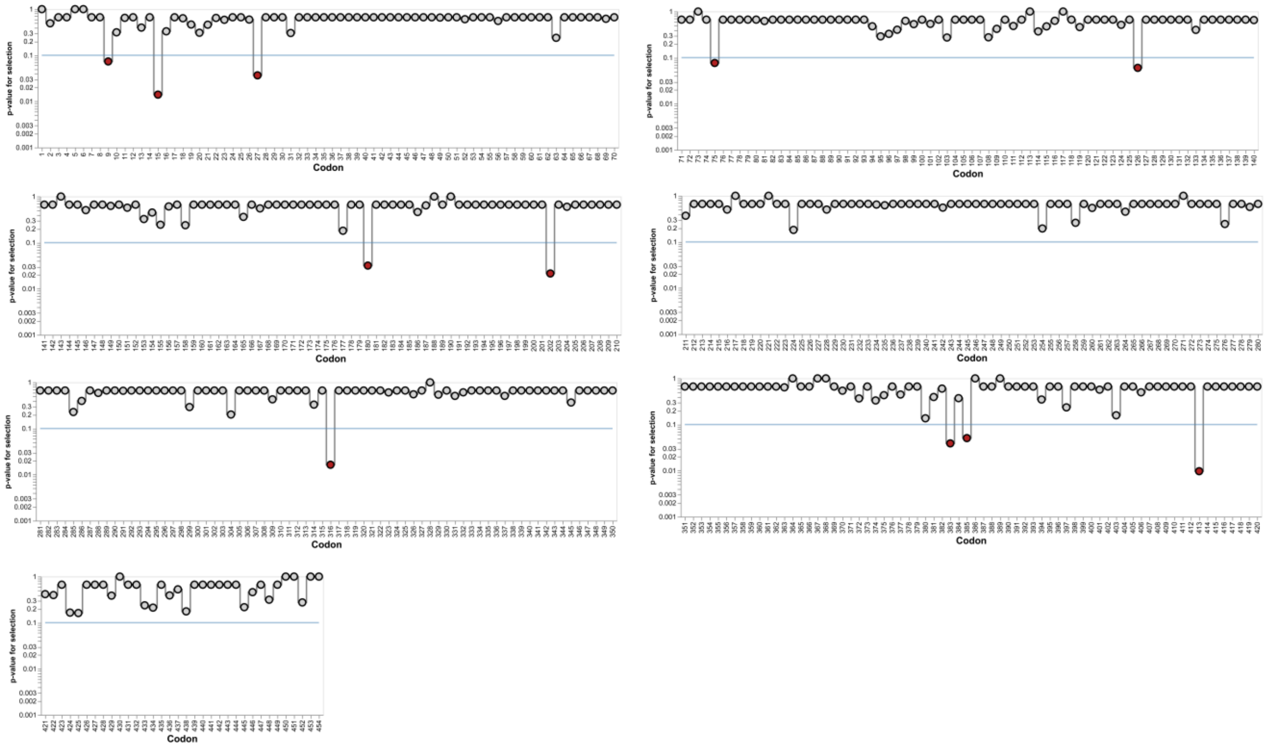

**Figure S7. Comparison of subcellular localisation at the species level** (for both TAA/TARs and YUCs) [prediction was done in DeepLoc-2.1 server, accessed on 17 October 2023: Web URL: https://services.healthtech.dtu.dk/services/DeepLoc-2.1/].

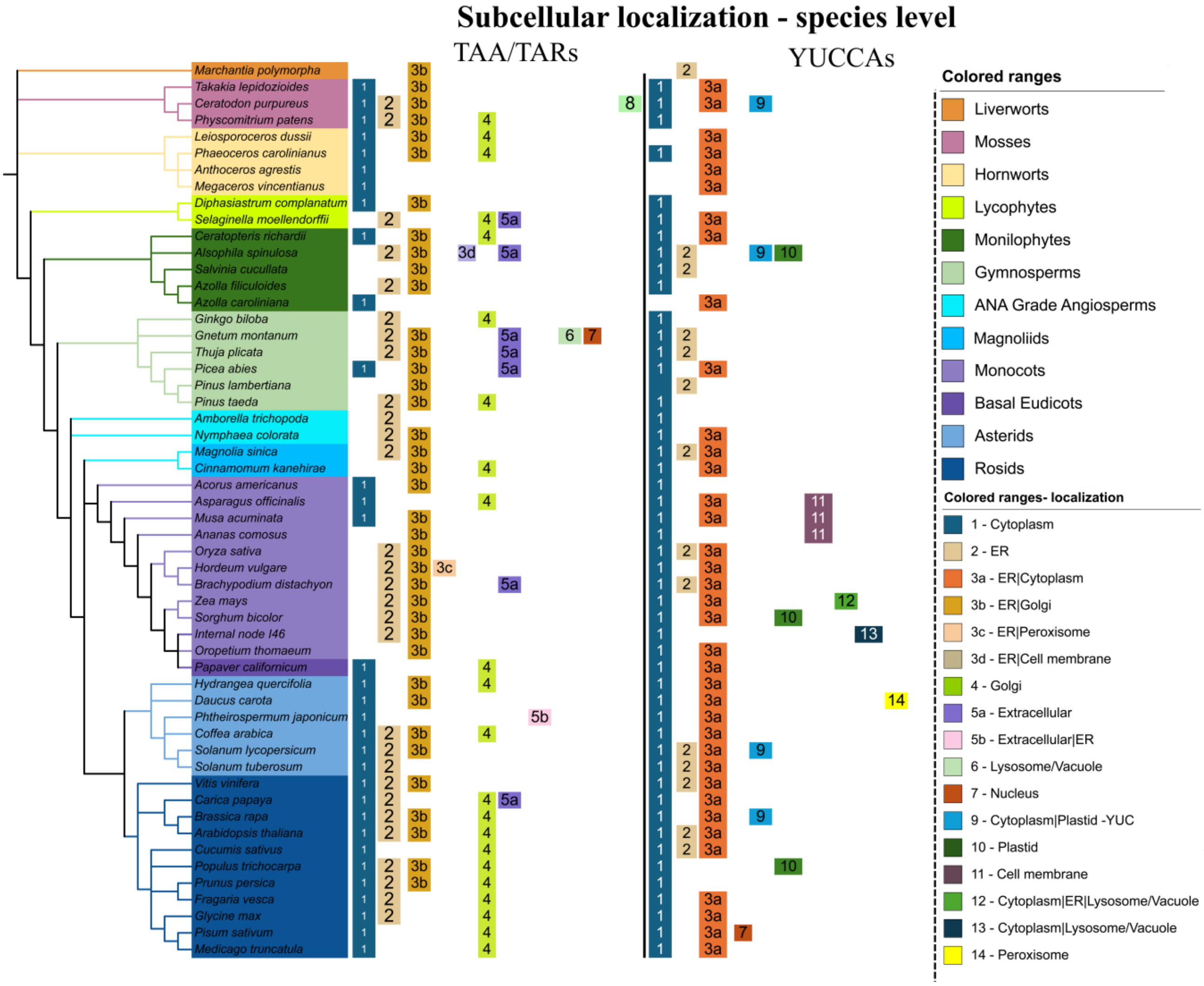

**Figure S8. Predicted subcellular localisations and membrane type. A) S**ubcellular localisation features of TAA/TAR and YUCs in general [DeepLoc 2.1 (Thumuluri et al., 2022)]. **B**) Protein membrane type comparison between TAA/TARs and YUCs. **C**) Class-based localization comparison. Those with less than 1% are not mapped. A single nuclear localization event was observed in class-4 YUCs. **D**) Comparison of observed signal sequences between the classes. More details with localization mapped against the corresponding protein are given **in Data S9**.

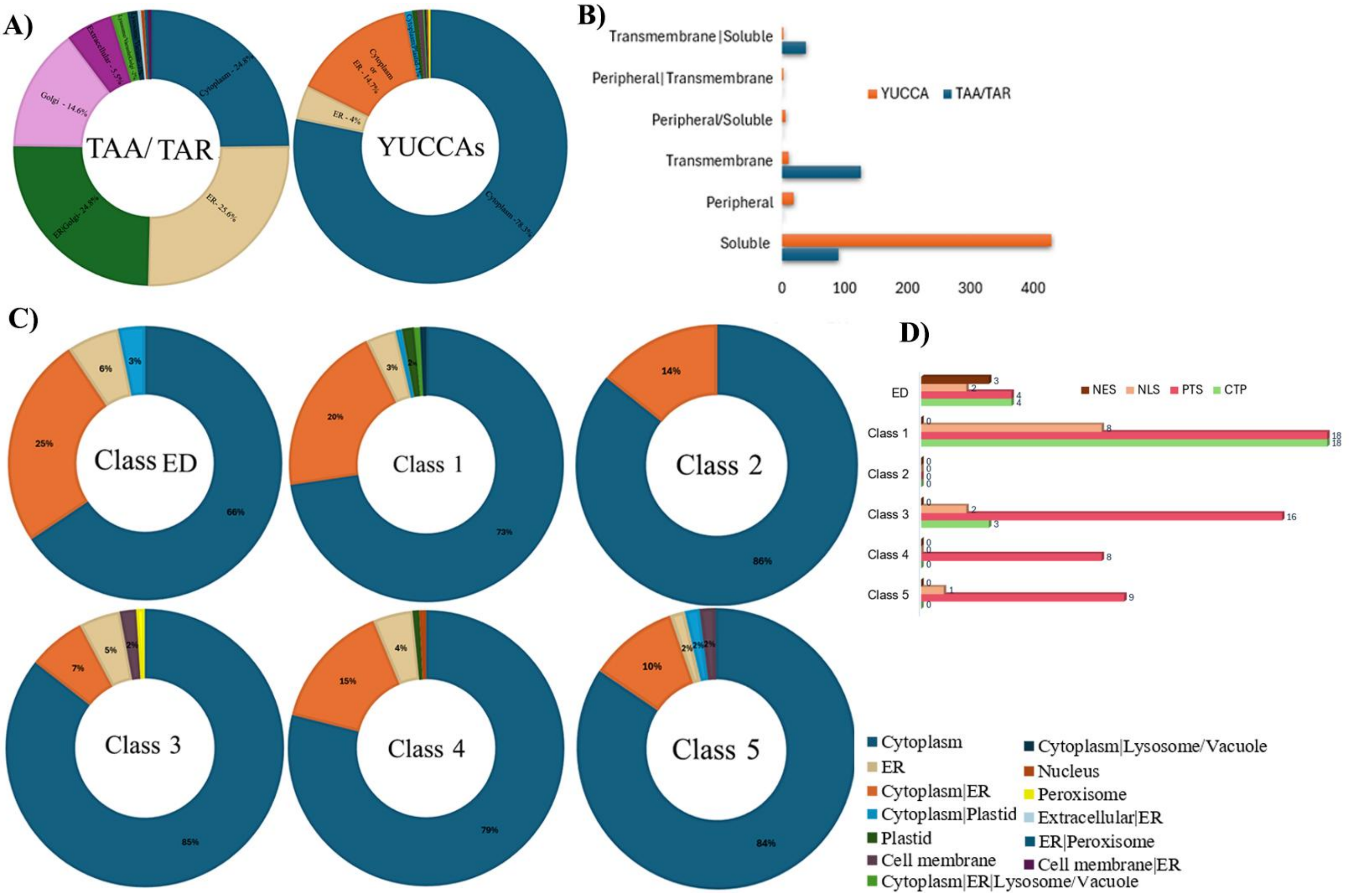

**Figure S9. Arabidopsis TAA/TARs - structural conservation pattern**. A) Sequence conservation pattern of TAA/TARs. all the catalytic sites (*), substrate binding sites (+) and the switch (T101, #) are highlighted. Signal peptide (labelled as **sp** - blue rectangle in alignment), detected by the SignalP v4.0 program in SMART. Transmembrane helix region (labelled as TM, green rectangle in the alignment), detected by the TMHMM v2.0 program in SMART (URL: http://smart.embl-heidelberg.de/smart, accessed on September 2024) B) Conservation pattern of clade 1 representative (AtTAR3) and clade-2 representative (AtTAA). Conservation patterns were analysed in Consurf and pictures were made with Pymol (Academic version). All the catalytic and substrate binding sites are represented as spheres. C) TAA/TAR domain organization pattern.

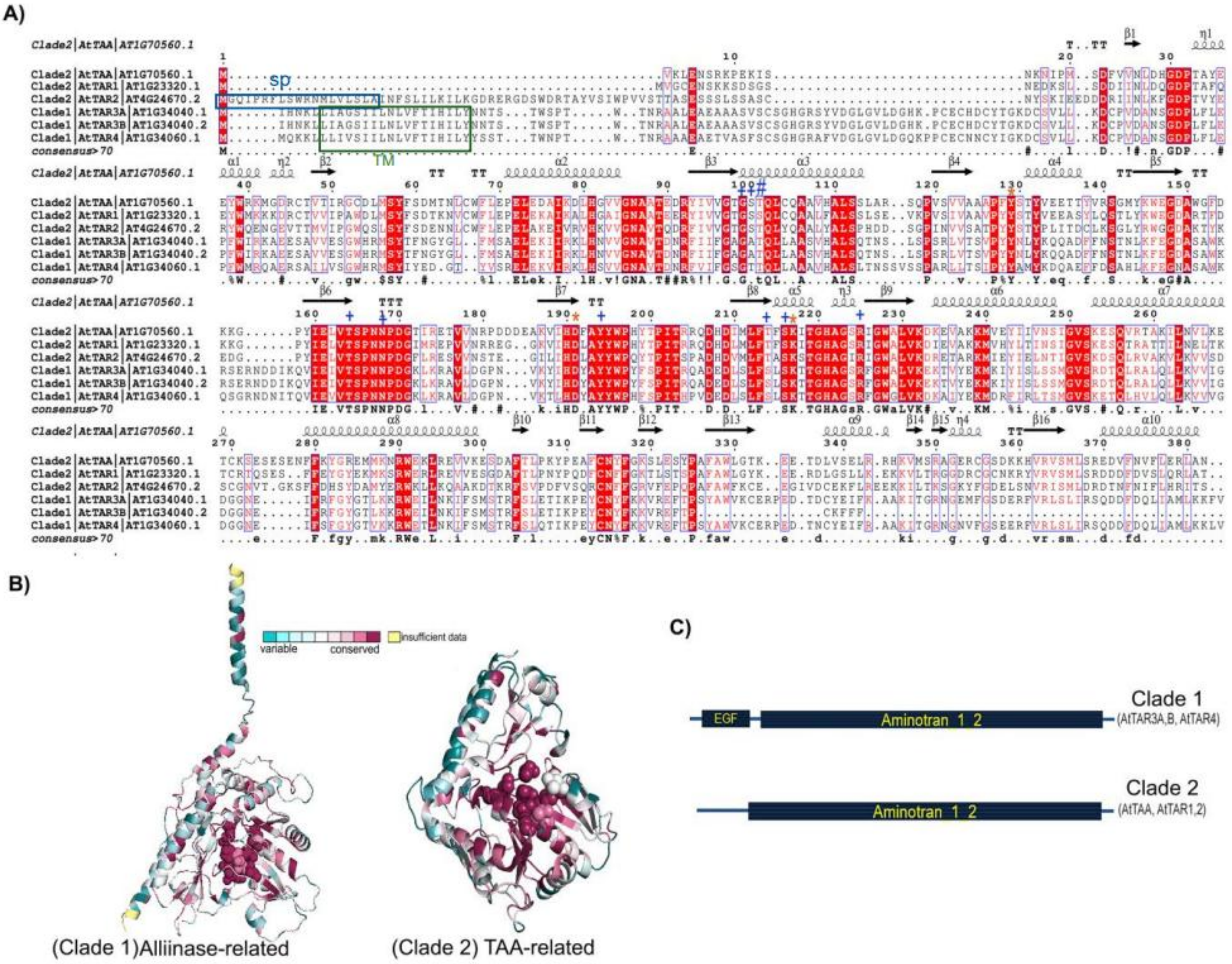

**Figure S10.** ***A. thaliana*** **YUCs (AtYUCs) interaction network.** The interaction data was obtained from the STRING database (Number of possible protein-protein interactions for each AtYUC is in bracket, confidence score is also given). The direct interacting partners (first shell of interactions) that are common to all AtYUCs are highlighted in orange. Six interaction partners are consistently associated with all AtYUCs, and all are playing crucial roles in auxin biosynthesis [i.e. L-tryptophan--pyruvate aminotransferase 1 (TAA1), Tryptophan aminotransferase-related protein 1 (TAR1), Tryptophan aminotransferase-related protein 2 (TAR2), Amidase 1(AMI1) and Aromatic aminotransferase ISS1 (ISS1), Indole-3-acetaldehyde oxidase or IAAld-oxidase (AAO2)]. Nitrilase (NIT) convert indole-3-acetonitrile to IAA and AAO1 are present in all class 3-5 AtYUCs. ISS1 involved in auxin homeostasis and AUX1 (Auxin transporter protein 1), is involved in proton-driven auxin influx. Aldehyde dehydrogenases (ALDHs) have a role in stress response and are predicted to interact and co-express with AtYUC11, AtYUC2, AtYUC3, AtYUC7, AtYUC8 and AtYUC4. PINs are auxin efflux carriers that have role in Auxin transport. Anthranilate synthase beta/alpha subunit 1 (ASB1/ASA1) catalyzes the first step of tryptophan biosynthesis and has a regulatory role in auxin biosynthesis. T7N22.1 has not been characterized yet. Other interacting partners mentioned are mainly transcription factors. Class 5 YUCs show many interacting factors and AtYUC1 display 45 high confidence interactions. More details of interaction (STRING output) are available at **Data S10(B)**

### AtYUCs protein- protein interactions

(No of interactions (confidence cut off > 0.7) is given in bracket)

Confidence score for each interacting protein is also mentioned in brackets

| Class 1 | | Class 3 | | Class 4 | | | | | Class 5 | |
|---|---|---|---|---|---|---|---|---|---|---|
| AtYUC10 (13) | AtYUC11 (10) | AtYUC2 (20) | AtYUC6 (18) | AtYUC3 (18) | AtYUC5 (18) | AtYUC7 (12) | AtYUC8 (26) | AtYUC9 (17) | AtYUC1 (45) | AtYUC4 (27) |
| TAR1 (0.963) | TAR1 (0.949) | TAA1 (0.963) | TAA1 (0.96) | TAA1 (0.962) | TAA1 (0.962) | TAA1 (0.925) | TAA1 (0.969) | TAA1 (0.958) | TAA1 (0.974) | TAA1 (0.963) |
| TAA1 (0.927) | TAA1 (0.925) | TAR2 (0.925) | TAR2 (0.936) | TAR1 (0.923) | TAR2 (0.924) | TAR1 (0.923) | ISS1 (0.931) | ISS1 (0.937) | TAR2 (0.966) | TAR2 (0.956) |
| TAR2 (0.921) | TAR2 (0.921) | TAR1 (0.924) | TAR1 (0.924) | TAR2 (0.921) | TAR1 (0.921) | TAR2 (0.912) | TAR2 (0.921) | TAR1 (0.911) | ISS1 (0.964) | TAR1 (0.934) |
| ISS1 (0.9) | ISS1 (0.9) | ISS1 (0.909) | ISS1 (0.903) | ISS1 (0.913) | ISS1 (0.913) | ISS1 (0.9) | TAR1 (0.911) | TAR2 (0.889) | TAR1 (0.962) | ISS1 (0.918) |
| T7N22.1 (0.785) | ALDH3F1 (0.746) | AMI1 (0.854) | NIT1 (0.828) | AMI1 (0.863) | IDD14 (0.852) | AAO1 (0.848) | IAA29 (0.902) | BHLH72 (0.784) | AMI1 (0.925) | SHI (0.874) |
| NFYB9 (0.785) | ALDH3H1 (0.725) | AAO1 (0.836) | AMI1 (0.824) | PIN2 (0.773) | TAR3 (0.796) | AMI1 (0.823) | PIF4 (0.901) | AMI1 (0.781) | AAO1 (0.907) | AMI1 (0.859) |
| AMI1 (0.759) | AAO2 (0.722) | PIN3 (0.796) | NIT3 (0.8) | AAO1 (0.756) | AAO1 (0.787) | AAO2 (0.721) | AMI1 (0.819) | IAA29 (0.758) | AUX1 (0.898) | AAO1 (0.836) |
| LEC2 (0.757) | ALDH2B7 (0.717) | AUX1 (0.793) | AUX1 (0.794) | AUX1 (0.755) | AMI1 (0.768) | TAR3 (0.719) | IAA19 (0.811) | PIF4 (0.739) | NIT1 (0.894) | SRS1 (0.826) |
| AUX1 (0.754) | AAO1 (0.715) | NIT3 (0.769) | ASB1 (0.783) | PIN7 (0.741) | NIT2 (0.762) | TAR4 (0.716) | BHLH72 (0.806) | AUX1 (0.726) | NIT2 (0.858) | NIT2 (0.788) |
| ORTH3 (0.739) | AMI1 (0.707) | TIR1 (0.758) | TIR1 (0.777) | TAR3 (0.741) | IDD16 (0.76) | NIT1 (0.714) | PIF5 (0.794) | ASB1 (0.72) | NIT3 (0.849) | LEC2 (0.785) |
| NIT2 (0.714) | | PIN7 (0.752) | AAO1 (0.776) | PIN4 (0.74) | PIN3 (0.76) | ALDH2B7 (0.714) | TAR3 (0.791) | ASA1 (0.718) | AAO2 (0.837) | PIN3 (0.785) |
| NIT3 (0.703) | | PIN4 (0.749) | LAX3 (0.776) | TAR4 (0.739) | NIT3 (0.758) | ALDH3H1 (0.701) | AUX1 (0.789) | AHL29 (0.716) | PIN3 (0.823) | AUX1 (0.784) |
| AAO2 (0.702) | | AAO2 (0.739) | PIN3 (0.775) | AAO2 (0.737) | AGL21 (0.74) | | PHYB (0.777) | AAO1 (0.709) | TIR1 (0.822) | PIN7 (0.771) |
| | | ARF8 (0.739) | NIT2 (0.763) | NIT2 (0.724) | NIT1 (0.726) | | AUF1 (0.777) | PIN3 (0.704) | CYP79B2 (0.796) | SRS2-2 (0.761) |
| | | TAR3 (0.738) | ABCB19 (0.738) | NIT3 (0.722) | SGR5 (0.719) | | PIN7 (0.757) | NIT2 (0.703) | NPY1 (0.796) | NPY1 (0.759) |
| | | NIT2 (0.737) | TAR4 (0.72) | PIN3 (0.721) | AUX1 (0.719) | | PIN3 (0.752) | AAO2 (0.702) | PIN7 (0.796) | NIT1 (0.755) |
| | | LAX3 (0.719) | SAG12 (0.718) | ALDH3H1 (0.72) | PIF4 (0.717) | | PIN4 (0.744) | PIN7 (0.7) | NPY5 (0.795) | AAO2 (0.743) |
| | | PIN2 (0.718) | AAO2 (0.714) | ALDH3F1 (0.7) | AAO2 (0.714) | | AAO1 (0.743) | | PIN4 (0.793) | AAO4 (0.723) |
| | | ALDH3F1 (0.709) | | | | | CYP79B2 (0.742) | | TAR3 (0.793) | ARF3 (0.719) |
| | | NIT1 (0.7) | | | | | HFR1 (0.74) | | SRS1 (0.788) | WOX11 (0.719) |
| | | | | | | | NIT3 (0.739) | | ASB1 (0.785) | PLT5 (0.719) |
| | | | | | | | NIT1 (0.721) | | TAR4 (0.785) | PIN4 (0.718) |
| | | | | | | | ELF3 (0.718) | | NPY3 (0.785) | SRS7 (0.718) |
| | | | | | | | ASB1 (0.718) | | PIN2 (0.785) | WUS (0.717) |
| | | | | | | | ALDH3F1 (0.705) | | LEC2 (0.784) | NIT3 (0.715) |
| | | | | | | | AAO2 (0.702) | | LAX3 (0.784) | ALDH3F1 (0.711) |
| | | | | | | | | | AFB2 (0.784) | ALDH3H1 (0.701) |
| | | | | | | | | | SHI (0.784) | |
| | | | | | | | | | AAO4 (0.762) | |
| | | | | | | | | | ASA1 (0.758) | |
| | | | | | | | | | WOX11 (0.758) | |
| | | | | | | | | | IAMT1 (0.758) | |
| | | | | | | | | | ARF7 (0.755) | |
| | | | | | | | | | ARF8 (0.755) | |
| | | | | | | | | | PIN8 (0.755) | |
| | | | | | | | | | ARF5 (0.753) | |
| | | | | | | | | | PIN5 (0.752) | |
| | | | | | | | | | LAX2 (0.74) | |
| | | | | | | | | | IAA17 (0.739) | |
| | | | | | | | | | WOX5 (0.729) | |
| | | | | | | | | | WUS (0.722) | |
| | | | | | | | | | ARF3 (0.719) | |
| | | | | | | | | | ARF6 (0.719) | |
| | | | | | | | | | PIF4 (0.717) | |
| | | | | | | | | | AFB3 (0.711) | |

#### Color code

- Common to all ATYUCs (TAA1, TAR1, TAR2, ISS1, AMI1, AAO2)
- AAO1
- ALDH
- Nitrilase
- PIN
- CYP79B2
- Transcription factors
- AUX1
- ARF
- ASB1/ASA1
- TIR-auxin receptor
- TAR3/4
- LAX
- NPY
- AAO4
- Uncharacterized protein
- Other proteins

#### Supplementary Tables

**Table S1**. Overall view of motif representation across various studied class B FMOs (Motif prediction was done in MEME suite: https://meme-suite.org/meme, accessed on August-September 2024). Motifs present in all class B FMOs are highlighted in red, and motifs that are common to YUCs and sYUCs are highlighted in green.

| Occurance of specific motifs in each classes | | | | | | | | | | | | |
|---|---|---|---|---|---|---|---|---|---|---|---|---|
| | **Motif type** | **FMO(S-OX) (Clade II*)** | **FMO(N-OX) (Clade IV*)** | **sYUC** | **YUC enzymes (Clade III FMO*)** | | | | | | | |
| | | | | | ED | 1(a-i) | 1 (j-q) | 2a | 2b | 3 | 4 | 5 |
| a | N-terminal EXEGK | | | | | | | | | | | |
| b | N-terminal PSEQXAKKEXDK | | | | | | | | | | | |
| c | N-terminal PTSREQAQ-his-rich | | | | | | | | | | | |
| d | N-terminal-MENMFRLXDHED | | | | | | | | | | | |
| e | N-terminal RRCVWVNGP | | | | | | | | | | | |
| **f** | **FAD binding GXGXXG** | | | | | | | | | | | |
| g | CELP | | | | | | | | | | | |
| g-1 | DHLP | | | | | | | | | | | |
| **h** | **PFYP-X(3)-QFXXYLEX (PFP--(X)n--Y/WLEX motif)** | | | | | | | | | | | |
| **i** | **ATG containg motif** | | | | | | | | | | | |
| i-1 | CXGXY/F (instead of ATG in FMOs (GS-OX and N-OX) | | | | | | | | | | | |
| **j** | **FMO-identifying FXGXXXHXXXY/F** | | | | | | | | | | | |
| **k** | **NADPH binding GXGXXG** | | | | | | | | | | | |
| l | GAXTSIVVRX(3)H | | | | | | | | | | | |
| m | PREXXGXSTF | | | | | | | | | | | |
| n | Leucine Rich KWLP motif | | | | | | | | | | | |
| o | Glycine Rich GX(5)GXXRPXXGP | | | | | | | | | | | |
| **p** | **VGT/AX(3)IXXG** | | | | | | | | | | | |
| **q** | **F/LATGY** | | | | | | | | | | | |
| r | DFFXXXG motif | | | | | | | | | | | |
| s | Glycine Rich WKGXXGX4GX4GXXG (not consistently present in all YUCs, but exists in all YUC classes) | | | | | | | | | | | |
| t | WKEET | | | | | | | | | | | |
| u | C-terminal K/QK/QKKX(5)HRR | | | | | | | | | | | |
| v | GAK motif | | | | | | | | | | | |
| | *FMO clades based on Nicoll and Mascotti, 2023, https://doi.org/10.1016/j.bbadva.2023.100108 | | | | Note: In YUCs alone 22 unique motifs | | | | | | | |

**Radar chart displaying a summary of the motif pattern types [based on the motif ('a - v') occurrence]**
(e.g., Type 5a motif pattern is mostly found in Class 1 YUCs, Type 10 is prevalent in Class-5 YUCs etc.)

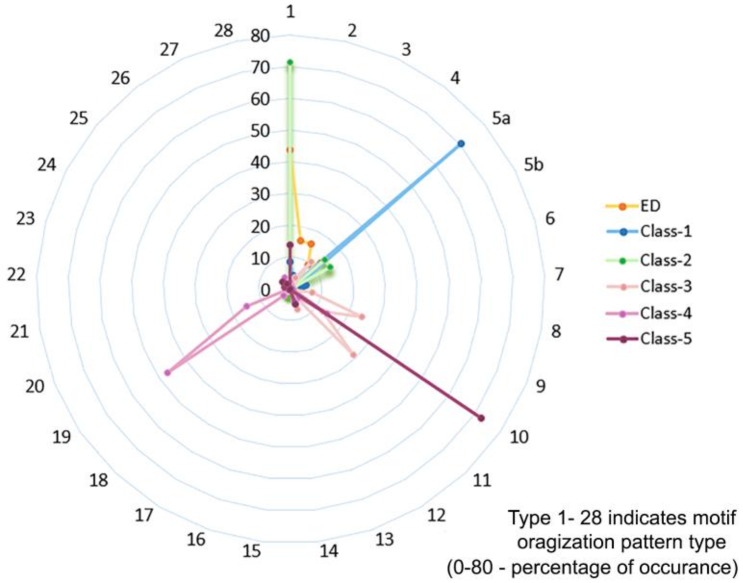

**Table S2.** Overall view of motif representation in various YUCs across various plant classes

| | Motif type | **Occurance of specific motifs in various plant groups** (Blue indicates presence and white indicates absence) | | | | | | | | | | |
|---|---|---|---|---|---|---|---|---|---|---|---|---|
| | | **Mosses** | **Liverworts** | **Hornworts** | **Lycophytes** | **Monilophytes** | **Gymnosperms** | **Basal angiosperms** | **Monocots** | **Basal dicots** | **Asterids** | **Rosids** |
| **a** | N-terminal EXEGK | ■ | ■ | | ■ | ■ | ■ | ■ | ■ | ■ | ■ | ■ |
| **b** | N-terminal PSEQXAKKEXDK | | | | | | ■ | | | | | |
| **c** | N-terminal PTSREQAQ-his-rich | | | | | | | | ■ | | | |
| **d** | N-terminal-MENMFRLXDHED | | | | | | | | | | | ■ |
| **e** | N-terminal RRCVWVNGP | | | | | | ■ | ■ | ■ | ■ | ■ | ■ |
| **f** | FAD binding GXGXXG | ■ | ■ | ■ | ■ | ■ | ■ | ■ | ■ | ■ | ■ | ■ |
| **g** | CELP | ■ | ■ | ■ | ■ | ■ | ■ | ■ | ■ | ■ | ■ | ■ |
| **h** | PFYP-X(3)-QFXXYLEX (PFP--(X)n--Y/WLEX motif) | ■ | ■ | ■ | ■ | ■ | ■ | ■ | ■ | ■ | ■ | ■ |
| **i** | ATG containg motif | ■ | ■ | ■ | ■ | ■ | ■ | ■ | ■ | ■ | ■ | ■ |
| **j** | FMO-identifying FXGXXXHXXXY/F | ■ | ■ | ■ | ■ | ■ | ■ | ■ | ■ | ■ | ■ | ■ |
| **k** | NADPH binding GXGXXG | ■ | ■ | ■ | ■ | ■ | ■ | ■ | ■ | ■ | ■ | ■ |
| **l** | GAXTSIVVRX(3)H | ■ | | | | ■ | ■ | ■ | ■ | ■ | ■ | ■ |
| **m** | PREXXGXSTF | ■ | ■ | ■ | ■ | ■ | ■ | ■ | ■ | ■ | ■ | ■ |
| **n** | Leucine Rich KWLP motif | ■ | ■ | ■ | ■ | ■ | ■ | ■ | ■ | ■ | ■ | ■ |
| **o** | Glycine Rich GX(5)GXXRPXXGP | ■ | ■ | ■ | ■ | ■ | ■ | ■ | ■ | ■ | ■ | ■ |
| **p** | VGT/AX(3)IXXG | ■ | ■ | ■ | ■ | ■ | ■ | ■ | ■ | ■ | ■ | ■ |
| **q** | F/LATGY | ■ | ■ | ■ | ■ | ■ | ■ | ■ | ■ | ■ | ■ | ■ |
| **r** | DFFXXXG motif | ■ | ■ | ■ | ■ | ■ | ■ | ■ | ■ | ■ | ■ | ■ |
| **s** | Glycine Rich WKGXXGX4GX4GXXG | ■ | ■ | ■ | ■ | ■ | ■ | ■ | ■ | ■ | ■ | ■ |
| **t** | WKEET | | | | | ■ | ■ | ■ | ■ | ■ | ■ | ■ |
| **u** | C-terminal K/QK/QKKX(5)HRR | | | | | | ■ | ■ | ■ | ■ | ■ | ■ |
| **v** | GAK motif | | | | | | | | ■ | | | ■ |
